# Designing of a next generation multiepitope based vaccine (MEV) against SARS-COV-2: Immunoinformatics and *in silico* approaches

**DOI:** 10.1101/2020.02.28.970343

**Authors:** Muhammad Tahir ul Qamar, Abdur Rehman, Usman Ali Ashfaq, Muhammad Qasim Awan, Israr Fatima, Farah Shahid, Ling-Ling Chen

**Affiliations:** College of Life Science and Technology, Guangxi University, Nanning, P. R. China; Department of Bioinformatics and Biotechnology, Government College University Faisalabad, Pakistan

**Keywords:** SARS-COV-2, COVID-19, Pneumonia, Epitopes, Vaccine, Linkers, Adjuvant

## Abstract

Coronavirus disease 2019 (COVID-19) associated pneumonia caused by severe acute respiratory coronavirus 2 (SARS-COV-2) was first reported in Wuhan, China in December 2019. Till date, no vaccine or completely effective drug is available to cure COVID-19. Therefore, an effective vaccine against SARS-COV-2 is crucially needed. This study was conducted to design an effective multiepitope based vaccine (MEV) against SARS-COV-2. Seven antigenic proteins were taken as targets and different epitopes (B-cell, T-cell and IFN-γ inducing) were predicted. Highly antigenic and overlapping epitopes were shortlisted. Selected epitopes indicated significant interactions with the HLA-binding alleles and 99.29% coverage of the world’s population. Finally, 505 amino acids long MEV was designed by connecting sixteen MHC class I and eleven MHC class II epitopes with suitable linkers and adjuvant. Linkers and adjuvant were added to enhance the immunogenicity response of the MEV. The antigenicity, allergenicity, physiochemical properties and structural details of MEV were analyzed in order to ensure safety and immunogenicity. MEV construct was non-allergenic, antigenic, stable and flexible. Molecular docking followed by molecular dynamics (MD) simulation analysis, demonstrated a stable and strong binding affinity of MEV with human pathogenic toll-like receptors (TLR), TLR3 and TLR8. Codon optimization and *in silico* cloning of MEV ensured increased expression in the *Escherichia coli* K-12 system. Designed MEV in present study could be a potential candidate for further vaccine production process against COVID-19. However, to ensure its safety and immunogenic profile, the proposed MEV needs to be experimentally validated.

## 1. Introduction

Viruses are dangerous pathogens and can cause irreversible losses to human lives and economy. The world hardly learns to deal with a virus when new emerges and threatens the future of humanity. A similar situation arises when a new strain of coronavirus (CoV) not previously identified in humans was reported last year (2019) [1]. Positive-sense RNA viruses called corona viruses belong to the Coronaviridae family that are distributed broadly among human and mammals. In the last two decades there have been more than 10,000 reported infections of two types of coronaviruses: severe acute respiratory coronavirus (SARS-COV) or Middle East Coronavirus (MERS-COV) [2]. Last year December (2019), few cases of unknown viral based pneumonia were reported in Wuhan city (Hubei province, P.R. China). Later it was identified that, this pneumonia is linked with a novel type of CoV outbreak. World Health Organization (WHO) named the virus SARS-CoV-2 and linked disease as coronavirus disease 2019 (COVID-19). The COVID-19 quickly widespread around the globe and claimed thousands of human lives. WHO announced global health emergency and recognized COVID-19 as pandemic [3-5]. SARS-CoV-2 was indicated root cause of COVID-19 through deep sequencing analysis from lower respiratory tract samples of patients [6]. SARS-COV-2 genome sequence is almost 70% similar to SARS-COV, and 40% similar to the MERS-COV [7]. Symptoms of SARS-COV-2 may arise within 2 days or up-to 14 days of exposure. Symptoms such as fever, diarrhea and respiratory disorder are found in infected patients [4]. According to latest research SARS-CoV-2 has an identical genomic organization as of beta-coronaviruses; 5’-untranslated region (UTR), orf1ab (replicas complex), nsps (encoding non-structural proteins), S (spike) protein, E (envelope) protein, M (membrane) protein, Oraf6, orf7a, orf8, N (nucleocapsid) protein, orf10, 3’-UTR and several unknown non-structural open reading frames [3, 8].

There is currently no vaccine or approved treatment for COVID-19. Few traditional Chinese medicine such as Shufengjiedu capsules and Lianhuaqingwen capsules were reported effective against COVID-19 [9, 10]. Nonetheless, no clinical trials support the safety and efficacy of these medicinal products. Similarly, clinical trials are in process for few experimental drugs, including remdesivir and chloroquine which were found effective *in vitro* against COVID-19 [11]. However, there is no clinical trial-based drug or vaccine reported yet. To prevent viral diseases vaccine is the most effective approach. Now a days, availability of genomic information, advance software and immunological data sets could greatly facilitate researchers to identify the effective epitopes from pathogens’ proteins that can be used to develop active sub-unit vaccines [12-15]. The subunit vaccine contains the fragments of antigenic proteins of pathogen that can trigger an immune response against the target pathogen [16, 17]. In recent studies, few candidate vaccine constructs were reported against MERS-CoV [18], Chikungunya virus [19], Ebola virus [20], Zika virus [21], HCV [22], Flavivirus [23] and Cytomegalovirus [24] with promising results. The *in silico* methods reduce the number of *in vitro* experiments and save time, overcome cost obstacles and increase the potential for successful vaccine design [25-27].

In present study, SARS-CoV-2 proteome was explored to determine the potent antigenic proteins and their further screening for B-cell, T-cell and IFN-γ inducing epitopes prediction with their MHC (major histocompatibility complex) alleles. Antigenicity, allergenicity and toxicity of predicted epitopes were analyzed. Multiepitope based vaccine (MEV) construct was designed by using the most potential and interacting epitopes, with the addition of suitable linkers and an adjuvant. Adjuvants are generally defined as molecules that may increase or modulate the intrinsic immunogenicity of an antigen [28]. Adjuvants are essential to reduce the amount of antigen and the number of injections, as they help to induce effective and persistent immune responses [29]. Several *in silico* approaches were utilized to validate the antigenicity, immunogenicity, allergenicity, structural stability/flexibility and physiochemical properties of designed MEV. Furthermore, molecular docking and MD simulations analyses were carried out to investigate binding interaction and stability of the MEV with human pathogenic receptors. At the end, the MEV codons were optimized for *E. coli* system and *in silico* cloning was performed to ensure its expression profiling (Figure 1).

**Figure 1.**
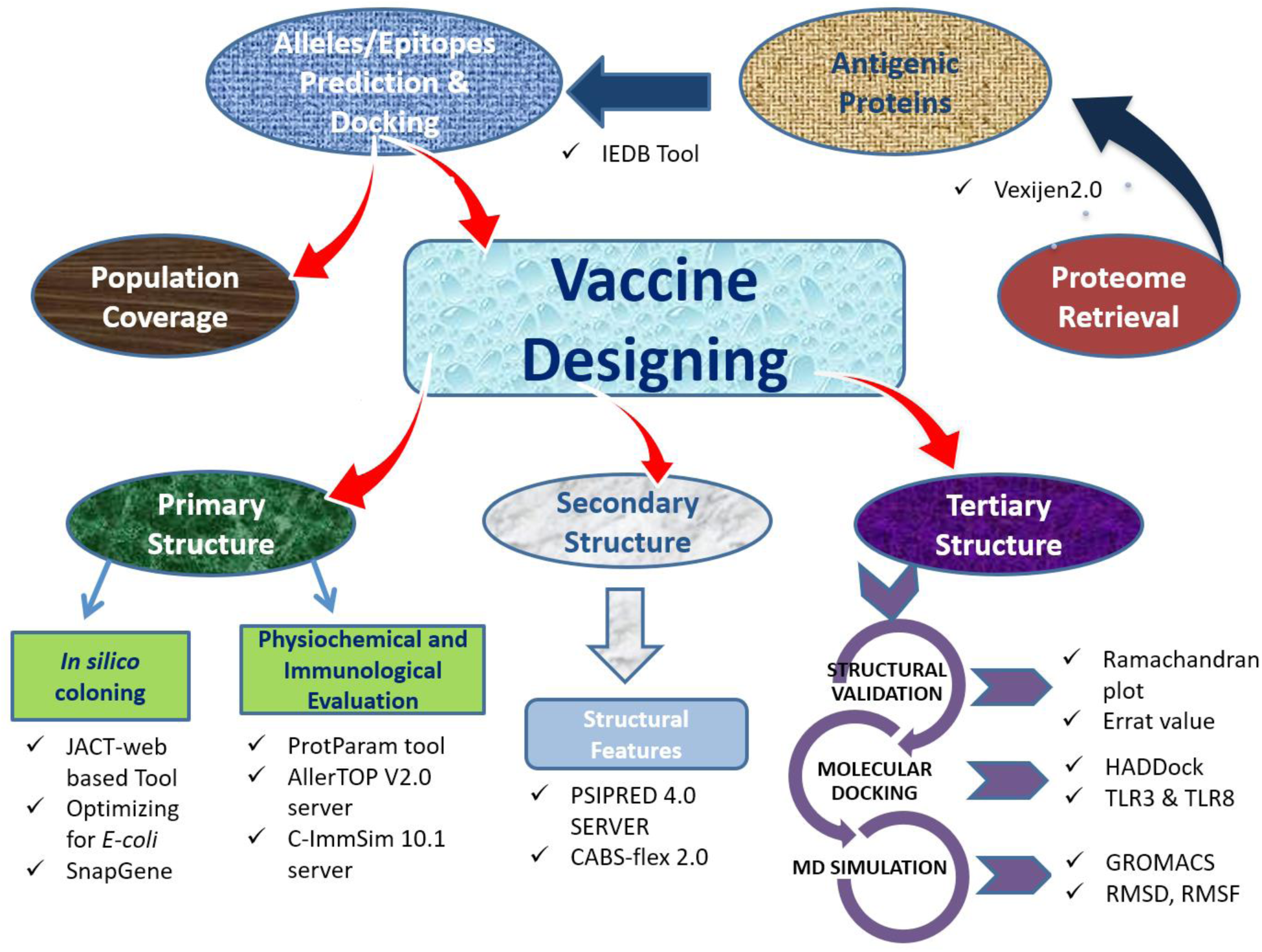
The overall experimental workflow used to develop MEV construct against SARS-COV-2.

## 2. Material and methods

### 2.1. Sequence retrieval and analysis of antigenic proteins

In the first step, whole proteome of SARS-CoV-2 was retrieved from GENBANK [30]. After that, individual protein sequences were extracted and stored as FASTA format for further analysis.

#### 2.1.1. Antigenicity and physiochemical properties evaluation

The Expassy Protparam tool was used to determine the physical and chemical properties of selected proteins [31]. To check protein antigenicity, the Vaxijen 2.0 software was used [32]. The threshold value was held at 0.5, and the secondary structure of proteins was predicted by using SOPMA tool [33].

#### 2.1.2. Tertiary structure prediction and refinement of target proteins

Tertiary stores of most of SARS-CoV-2 proteins are not reported yet. Therefore, combinations of different approaches were employed to predict good quality structures for further analysis. Online tools such as Swiss model which work on homology based modeling algorithms and Raptor X which work on deep learning modules, were primarily used for the tertiary structure prediction of SARS-COV-2 proteins [34-36]. Predicted Models were then refined by galaxy refine server and validated by Ramachandran plot analysis.

### 2.2. Epitopes prediction and validation

#### 2.2.1. B-cell epitope prediction

In immune system the B-Cell epitope helps to detect viral infection and activities. B-cells provide humoral immunity by secreting immunoglobulins which can neutralize antigen upon binding. A surface receptor of B-cell recognizes B-cell epitopes, resulting the generation of antigen-specific immunoglobulins [37]. B-cell epitopes are two types, linear (continuous) and conformational (discontinuous). An online database of ABCPred was used to predict linear B-cell epitopes [38, 39]. Conformational epitopes were predicted by Ellipro server [40].

#### 2.2.2. T-cell Epitope prediction

T-cells express TCRs (T-cell receptors), that could recognize specific antigens and can generate cellular and humoral immune response against them [41]. The Immune Epitope Database (IEDB) consensus method [42, 43] was used to predict MHC classes I and MHC II epitopes with consensus score less than 2 filter cut.

#### 2.2.3. Immunogenicity prediction of epitopes

To evaluate the antigenicity, allergenicity and toxicity of B-cell and T-cell epitope, Vaxijen v2.0 and AllerTOP v2.0 were used. The filtering criteria 0.5 was used. Only filtered, non-antigenic, non-allergenic and non-toxic epitopes were selected for further studies.

#### 2.2.4. Conservation analysis of epitopes

IEDB conservancy analysis tool [44] was used, to monitor the degree of conservation in the sequence of B-cell and T-cell epitopes. Epitopes showing 100 percent conservation were shortlisted for further analysis.

#### 2.2.5. Interferon-γ *inducing epitope prediction*

IFN-γ is acknowledged to elicit intrinsic safe responses and can directly detain viral duplication [45, 46]. Besides, they can trigger the versatile immune reactions by preparing cytotoxic T lymphocyte (CTL) and Helper T lymphocyte (HTL). IFN epitope server was used to calculate IFN-γ epitopes of selected proteins of SARS-CoV-2 using SVM hybrid algorithms along with Motif [47].

#### 2.2.6. Epitopes modeling and molecular docking

Epitopes which show favorable strong binding affinities with a common experimentally validated allele, are good choice to design MEV construct. Therefore, molecular docking between screened epitopes and human allele was performed. Molecular docking is an *in silico* approach which determined the binning affinity between ligand and its target proteins, and also highlight the important residues involve in the interaction. 3D structures of overlapping, highly antigenic and conserved epitopes with corresponding common alleles were predicted using PEPFOLD [48]. The X-ray crystallographic structure of a common human allele (HLA-B7) was retrieved from protein data bank (PDB ID: 3VCL). Molecular docking was performed using same protocol of our previously published studies [5, 18, 19, 49, 50]. To visualize the docked complexes and draw figures, the PyMOL molecular graphics system was used [51].

#### 2.2.7. Population coverage analysis of selected epitopes

The selected epitopes for MEV construct should effectively cover major populations across the globe. For population coverage, overlapping, antigenic, conserved and strongly interacting with HLA-B7 allele epitopes were selected and further analysed using the IEDB population coverage analysis tool by maintaining the default analysis parameters ^[44]^. This tool is designed to estimate the population coverage of epitopes from diverse counties based on the distribution of their MHC-binding alleles. As SARS-CoV-2 is global pandemic, therefore, worldwide analysis has been performed.

### 2.3. Construction of multi epitope vaccine

To construct a sub-unit vaccine, the epitopes with following properties are usually preferred: (a) 100% conserved, (b) overlapping, (c) highly immunogenic, (d) non-allergic, (e) non-toxic, (f) with significant population coverage, (g) having strong binding affinity with common human allele and (h) have no similarity with the human proteins. Therefore, only those CTLs, HTLs and B-cell epitope were further selected which passed all the eight above parameters to construct SARS-CoV-2 MEV. To boost the immune response an adjuvant was attached with the first CTL epitope with the EAAAK linker, while other epitopes were connected using AAY and GPGPG linkers to preserve their independent immunogenic activities after their inter-interaction compatibility validation using HADDOCK Guru interface [52].

#### 2.3.1. Immunogenic and physicochemical properties evaluation of the vaccine construct

The vaccine construct should be stable, antigenic and non-allergic. The ProtParam tool [53] was used to evaluate the physicochemical properties of MEV. It analyzes different physical and chemical features of proteins including grand average hydropathy, half-life, stability/instability index, theoretical pI, aliphatic index hydropathy and half-life. The MEV properties were further verified using VaxiJen 2.0 and AllerTOP V2.0 servers [54]. Moreover, PSIPRED was used to analysed the secondary structure of MEV [55].

#### 2.3.2. Tertiary structure prediction of vaccine construct

As vaccine construct is combination of different epitopes, therefore, the RaptorX server was used to develop MEV 3D tertiary structure. The RaptorX server use a multi-template threading approach to determine the tertiary structure of query protein [34].

#### 2.3.3. Refinement and validation of vaccine construct

Galaxy Refine server MD simulation approach was used to refine the MEV predicted 3D structure [56]. To verify refined MEV structure quality, Ramachandran plot analysis was performed using RAMPAGE server [57], followed by structural validation analysis using ProSA-web server [58]. The quality scores outside the normal range of natural proteins during structure validation process reveal potential defects in the protein structure model. Therefore, ERRAT server was further used to evaluate the statistics of non-bonded interactions in MEV construct [59]. Furthermore, MEV structural flexibility was also analysed using CABS-Flex 2.0 server [60]. The flexibility of vaccine is an important aspect for its functionating, and CABS-Flex server provides a detail overview of flexibility and stability of query protein by simulating its residues [61].

#### 2.3.4. Immunogenicity evaluation of the vaccine construct

An *in silico* immune simulation was performed using C-ImmSim 10.1 server [62], in order to validate immunological response of constructed MEV. This server simulates the three major functional mamal system components (bone marrow, thymus and lymph node) [62]. The MEV has been tested for the ability to simulate various types of immune cells such as HTL, CTL, B-cells, NK cells, dendritic cells, Immunoglobulins and cytokines. Clinically the minimum recommended interval between two doses of vaccines is four weeks [24, 41, 63, 64]. Consequently, three injections (each injection contain a thousand units of MEV) were administered using C-ImmSim immunostimulatory, with the recommended interval of four weeks (1, 84 and 168 time-steps parameters were set as 1 time-step is equal to eight hours of real life) for a total of 1000 steps of simulation. Other parameters were kept as default.

### 2.4. Molecular docking of vaccine construct with human immune receptor

The host produces an efficient immune response if an antigen / vaccine interacts properly with the target immune cells. Therefore, molecular docking analysis was performed to analyse integrations between the MEV and the human immune receptors. TLR3 and TLR8 have been extensively studied and studies found their vital roles in the generation of antiviral immune response. HADDOCK was used for the MEV docking with TLR3 (PDB ID: 1ZIW) and TLR8 (PDB ID: 3W3 G). To visualize the docked complex and draw figures, the PyMOL educational version was used [51]. In addition, the online database PDBsum was used to demonstrate the interacting residues of docked complexes [65].

### 2.5. Molecular dynamics simulation analysis of vaccine construct and receptors complexes

MD simulation is an important approach to analyse the stability of the receptor-ligand complex [41, 66]. Complexes of MEV with TLR3 and TLR8 were simulated at 20 ns using GROMACS 5.1.4 [67] by following the same protocol of our previously published studies [5, 49, 68]. The trajectories were saved for each complex after every 2 fs and root mean square deviation (RMSD) and root mean square fluctuations (RMSF) analysis were performed using GROMCAS tools.

### 2.6. Codon optimization and in silico cloning

Codon adaptation is a way of increasing the translation efficacy of external genes in the host if the use of codon in both species varies. After carefully evaluating MEV properties and immune response, finally its codon optimization was performed followed by *in silico* cloning. The Java Codon Adaptation Tool (JCAT) server [69] was used for codon optimization of MEV to make it compatible with widely used prokaryotic expression system; *E. coli* K12 [70]. The available additional options were selected to evade (a) rho-independent transcription termination, (b) prokaryote ribosome binding site and (c) restriction enzymes cleavage sites. The GC (guanine and cytosine) contents together with codon adaptation index (CAI) [71] were evaluated. Furthermore, to facilitate restriction and cloning, sticky ends restriction sites of XhoI and HindIII restriction enzymes were introduced at the start/N-terminal and end/C-terminal of the optimized MEV sequence, respectively. Finally, the adapted nucleotide sequence of MEV was cloned into the *E. Coli* pET28a(+) vector with SnapGene 4.2 tool (https:/snapgene.com/) to ensure the *in vitro* expression.

## 3. Results

### 3.1. Target proteins sequence and structural analysis

The amino acid sequences of SARS-CoV-2 important vaccine target proteins (ORF1 [QHD43415.1], S [QHD43416.1], ORF3a [QHD43417.1], E [QHD43418.1], M [QHD43419.1], ORF6 [QHD43420.1], ORF7a [QHD443421.1], ORF8 [QHD43422.1], N [QHD43423.2] and ORF10 [QHI42199.1]) were retrieved and saved in FASTA format. Vaxijen server was used to check the antigenicity of proteins. Total 7 highly antigenic proteins were detected. The most antigenic protein found was ORF10, followed by E, M, ORF6, ORF7a, ORF8, and N, with antigenic values 0.7185, 0.6502, 0.6441, 0.6131, 0.6025, 0.5102 and 0.5059 respectively. ORF1, S, and ORF3 proteins had antigenic values less than 0.5, therefore they were excluded from further analysis. Among selected 7 highly antigenic SARS-CoV-2 proteins for further analysis, 5 were non-structural proteins (M, N, ORF6, ORF7a, and ORF10) and 2 were structural proteins (E, ORF8). Other physicochemical characteristics of selected proteins including theoretical pi, molecular weight, half-life, stability profile, aliphatic index, etc were analysed using ProtParam server (Supplementary Table 1) and their secondary structures was predicted using SOPMA tool (Supplementary Table 2).

The 3D models of selected proteins tertiary structures were predicted using Swiss model tool and Raptor X tool, and in order to select best quality models, predicted structures were further refined by galaxy refine server followed by Ramachandran plot validation. The most of structures predicted using Swiss model were of better quality than Raptor X predicted structures, except for N and Orf7a proteins. Therefore, good quality models were selected for further analysis (Supplementary Figure 1). There was no suitable structure predicted for ORF10 because of small number of residues. So, its structure was predicted by PEPFOLD server [48] (Supplementary Table 3).

### 3.2. Prediction of B-cell, T-cell and IFN-γ inducing epitopes

Screened out B-cell epitopes were 100% conserved in all protein sequences and ere highly antigenic. All the target proteins were predicted to have a total 55 linear epitopes (E-4, M-12, ORF6-1, ORF7a-6, ORF8-9, N-22, and ORF10-1) (Supplementary Table 4). Moreover, a total of 24 (E-4, M-2, ORF6-3, ORF7a-4, ORF8-4, NC-4, and ORF10-3) conformational epitopes were predicted in all target proteins (Supplementary Table 5).

As mentioned before, epitopes that can bind to multiple alleles because of their strong defense capabilities are considered the most appropriate epitopes. Therefore, total 31 MHC class I (E-9, M-4, ORF6-2, ORF7a-3, ORF8-7, N-4, ORF10-2) (Supplementary Table 6) and 40 MHC class II (E-4, M-5, ORF6-4, ORF7a-7, ORF8-12, N-4, ORF10-4) epitopes, 100% conserved among target protein sequences and highly antigenic were chosen for further study (Supplementary Table 7).

The HTLs helps to activate CTLs together with other immune cells upon various types of cytokines release i.e., IFN-γ, interleukin-4 and interleukin-10 [17, 72]. Thus, HTL epitopes that induce cytokines are vital for the progress of vaccines or immunotherapy. A total of 988 IFN-γ inducing epitopes from target proteins (E-67, M-214, ORF6-53, ORF7a-100, ORF8-113, N-411, ORF10-30) were obtained (Supplementary Table 8).

### 3.3. Interaction analysis of epitopes with HLA-B7 allele

As stated before, to construct a sub-unit vaccine, the chosen epitopes should be 100% conserved, overlapping and antigenic. Therefore, total 50 conserved/antigenic epitopes from selected proteins overlapping in all 3 categories (B-cell, T-cell and IFN-γ) were selected for further validation of their interactions with a common human allele. The 3D structures of selected epitopes were predicted using PEPFOLD. The binding patterns of selected epitopes with a common conserved allele HLA-B7 were analyzed through molecular docking and it was found that only 27 epitopes bind deep inside in the HLA-B7 binding pocket (Supplementary Figure 2-3). Each bound epitope to HLA-B7 depict stronger than −10.00 Kcal/mol docking affinity. Docking binding energy scores together with their detail information is mentioned in Table 1. All the 27 selected epitopes ensured their binding efficiency as well as their suitability to be used in multiple epitope-based vaccine construct.

**Table 1.**
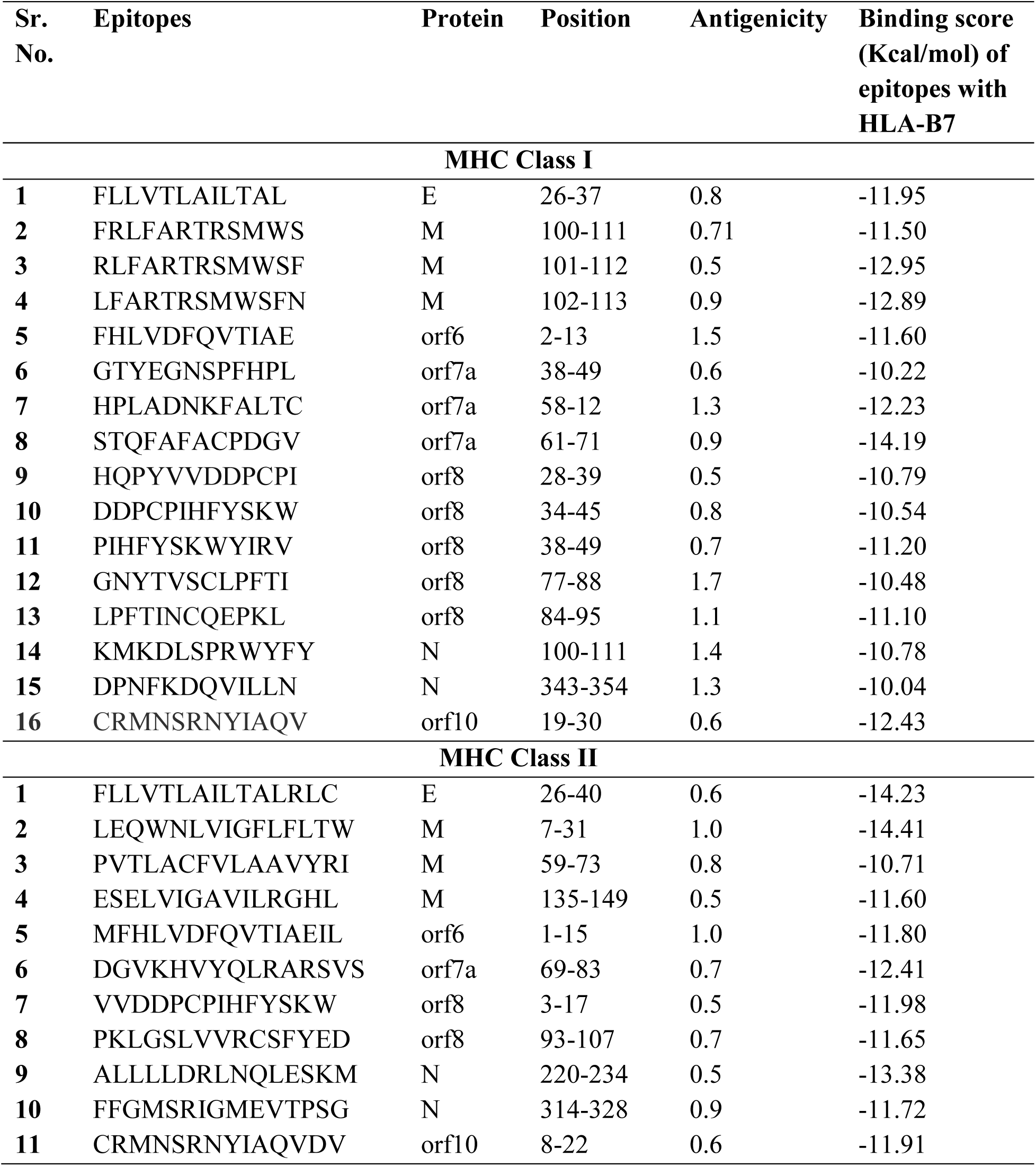
Final selected epitopes from SARS-CoV-2 antigenic proteins used to design MEV construct.

### 3.4. Population coverage

The distribution and expression of HLA alleles vary by ethnic groups and regions of the world. Therefore, it affects the successful development of an epitope-based vaccine. The IEDB population tool was used to check the population coverage of selected epitopes and their suitability for vaccine construct. Selected epitopes showed the 99.29% world population coverage. The highest coverage of population found within a country Sweden 99.79%. The population coverage for worst COVID-19 hit countries; China, Italy, Spain and Singapore were 84.51%, 96.59%, 99.26% and 85.23%, respectively (Figure 2). Likewise, significant population coverage was identified for other seriously affected countries with SARS-CoV-2 including Australia, USA and France. The result suggested that our MEV could help to combat against COVID-19 in most of the regions round the globe.

**Figure 2.**
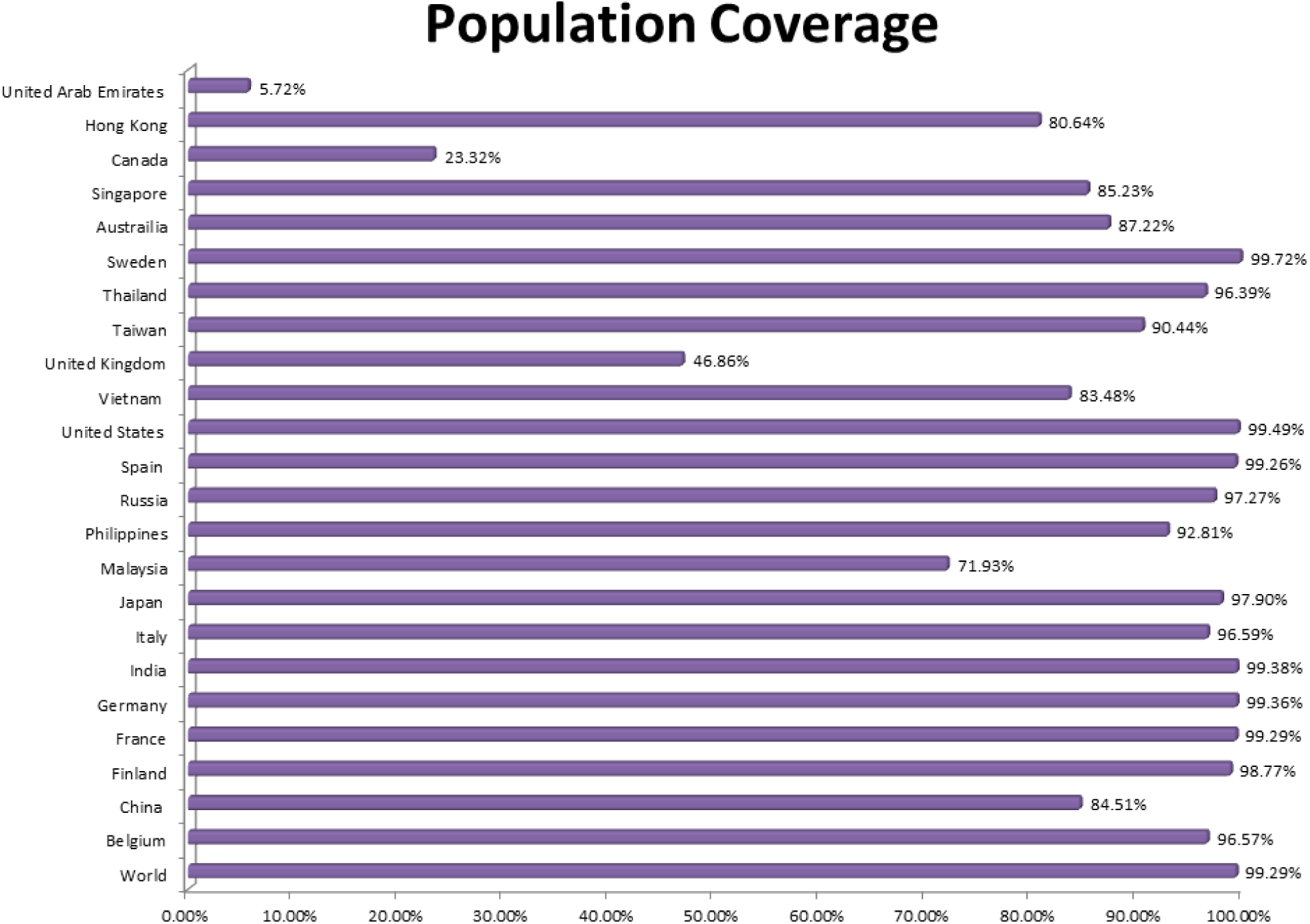
Worldwide population coverage of MEV epitopes based on their respective HLA binding alleles.

### 3.5. Construction of multiepitope based vaccine

All 27 selected epitopes (E-2, M-15, ORF6-2, ORF7a-4, ORF8-7, N-5, ORF10-2) were validated for their inter-interactions and further used to develop an MEV construct. An adjuvant (45 amino acid long β-defensin) was linked with the help of EAAAK linker at the start (to the N-terminal of the MEV). EAAAK linker reduce interaction with other protein regions with efficient separation and increase stability [73, 74]. The vaccine’s immunogenicity may increase with an adjuvant. Epitopes were merge together based on their interaction’s compatibility in sequential manner with AAY and GPGPG linkers respectively. AAY and GPGPG prevents the generation of junctional epitopes, which is a major concern in the design of multiepitope vaccines; On the other hand, they facilitates the immunization and presentation of epitopes [75, 76]. The final vaccine construct comprises 505 amino acid residues (Figure 3).

**Figure 3.**
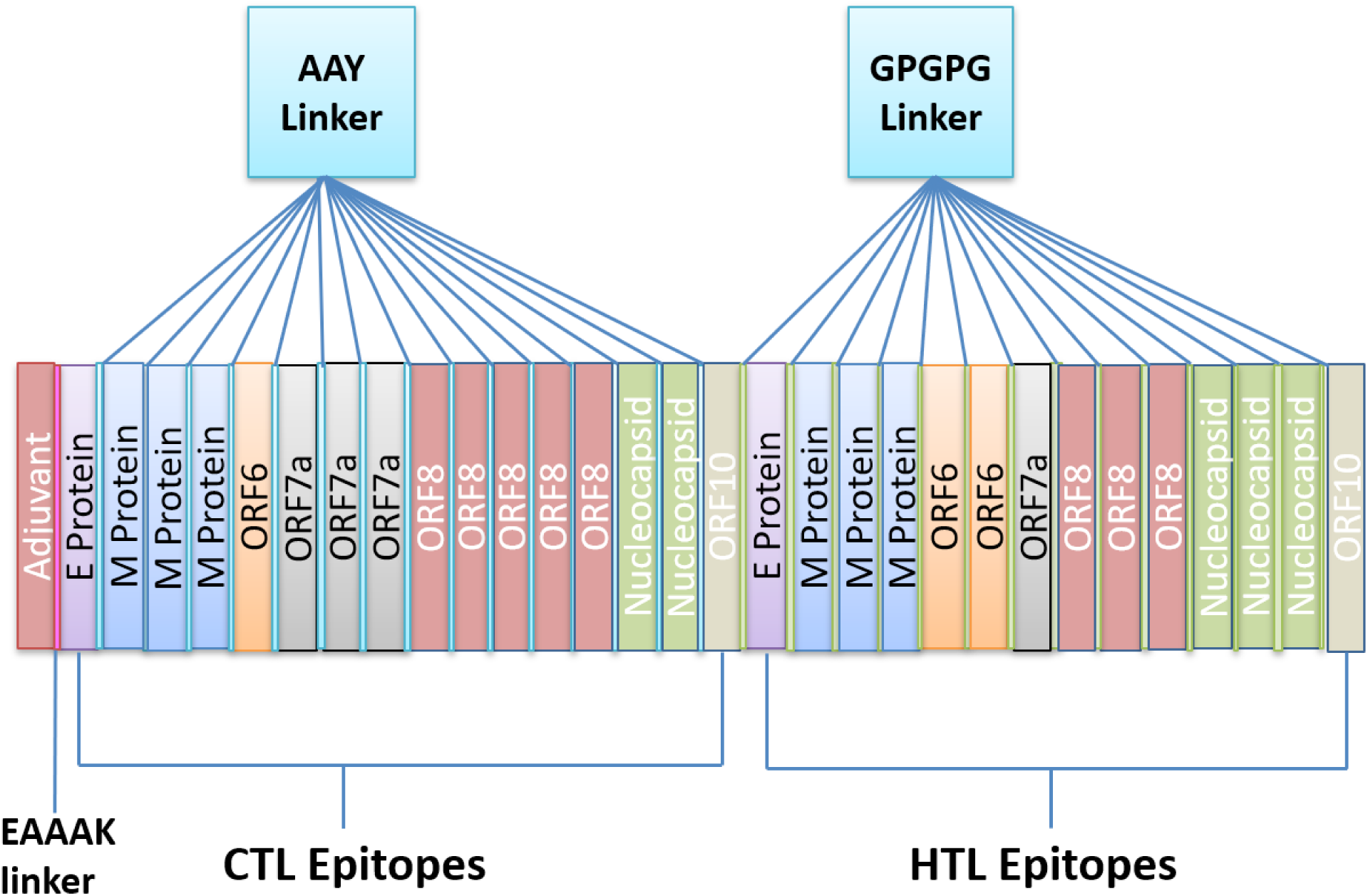
Schematic diagram of MEV construct: an adjuvant (Maroon) linked at N-terminal with the help of EAAAK linker (Pink). CTL epitopes are joined by AAY linkers (Blue) while HTL epitopes are joined by GPGPG linkers (Green).

### 3.6. Evaluation of multiepitope based vaccine

First, Blast-p analysis was performed against *Homo sapiens* proteome with default parameters to validate that MEV is non-homologous. Protein with less than 37% identity generally considered as non-homologous protein. However, MEV showed no similarity (higher or equal to 37%) with the human proteins.

Next, allergenicity, antigenicity and toxicity of the vaccine construct were evaluated. Results described that MEV is highly antigenic (0.6741 at 0.5% threshold), non-allergenic and non-toxic.

Further, the physiochemical properties of the SARS-CoV-2 MEV construct were determined using Protparam. It contains 505 amino acids with 55426.35 KDa molecular weight, indicating good antigenic nature. The isoelectric point (PI) of MEV was 9.12 which show its negative. Negatively charged MEV showed the value of isoelectric point (pI) less than 7. MEV was categorized as stable as instability index computed by the Protparam was 33.41. The aliphatic index was 82.75, which represents the idea of the proportional volume of the aliphatic side chains. The protein sequence has a GRAVY value of 0.105, indicating the hydrophobic nature of the MEV. Total time taken for a protein to disappear after it has been synthesized in cell depicted as the half-life of the protein which was computed as > 20 h for yeast, 30 h for mammalian-reticulocytes and > 10 h for *Escherichia coli*. MEV computed formula wa C2549H3850N666O669S28, depicting the cumulative numbers of Carbon (C), Oxygen (O), Nitrogen (N), Hydrogen (H) and Sulfur (S). Above results indicated MEV as a suitable potential vaccine.

### 3.7. Structural analysis of multiepitope based vaccine

Secondary structure of MEV was predicted by PSIPRED. Among the 505 amino acids, the formation of α-helix is comprised of 176 amino acids representing 35.20%, 109 amino acids in β-strands representing 21.59% and 215 amino acids forms the coils which are 42.58% of the whole MEV construct (Supplementary Figure 4).

To determine the tertiary structure of vaccine RaptorX server was used. Structure was refined by Galaxy refine server (Figure 4). The improved model depicted that 95% amino acids are in favorable region, 96.3% of residues in most favorable region, 3.7% of residues in permitted region and 0.0% in outer region according to Ramachandran plot analysis. Further analyses revealed qRMSD is 0.428, poor rotamers are 0%, MolProbity is 1.889, clash score is 13.6, and Z score is −2.25 Z. In addition, the refined model showed 0 errors with PROCHECK validation. The refined model score was 85.7143 in quality check analysis through ERRAT. These results show that the refined model is of good quality.

**Figure 4.**
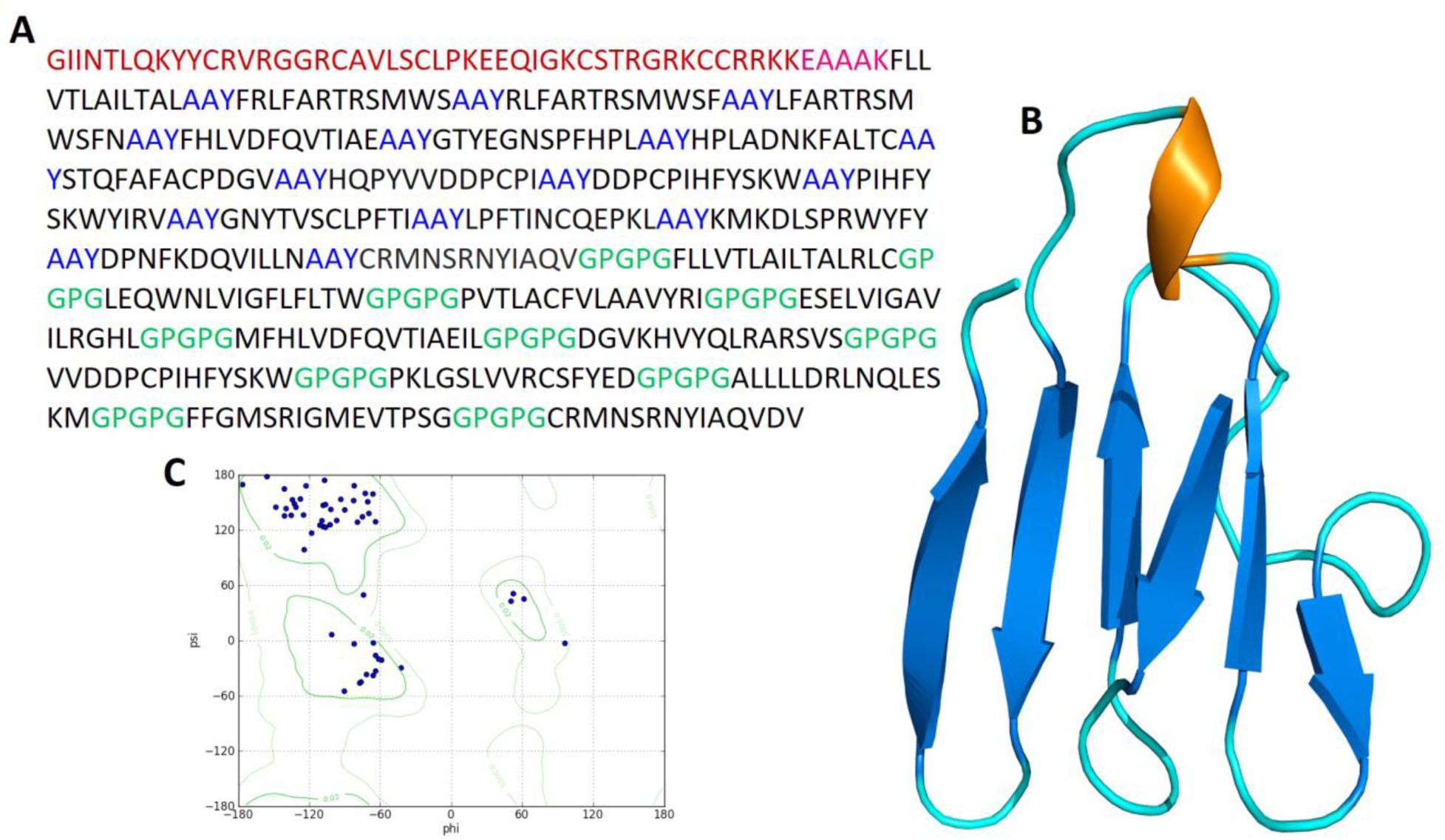
(**A**) A 505 amino acid long MEV sequence consisting an adjuvant (Maroon) linked at N-terminal with the help of EAAAK linker (Pink). CTL epitopes are joined by AAY linkers (Blue) while HTL epitopes are joined by GPGPG linkers (Green). (**B**) The predicted three-dimensional structure of the MEV vaccine construct (Alpha helixes: Brown; Beta sheets: Blue; Loops: Cyan). (**C**) Ramachandran plot analysis result of predicted MEV structure where 96.3% of residues present in the most favorable region.

In addition, flexibility of MEV structure was evaluated using CABS-flex 2.0 server with 50 cycles simulation at 1.4 °C temperature. Among the 10 final retrieved 3D structures, regions near to N-terminal depicted lesser fluctuation compared with the regions near the C-terminal (Supplementary Figure 5A). Resultant contact-map presented the residue-residue interaction pattern for all the 10 final retrieved models (Supplementary Figure 5B). Finally, root mean square fluctuation (RMSF) plot revealed the fluctuation of each of the amino acid of MEV model ranging from 0.0 Å to 3.5 Å. The presence of fluctuations in the MEV structure, indicated its high flexibility and endorse it as a potential vaccine construct (Supplementary Figure 5B).

### 3.8. Prediction of B-cell epitopes in multiepitope based vaccine

B-lymphocytes besides secreting cytokines, also produce antigens, which in return provide humoral immunity [77]. Therefore, MEV ideally should have B-cell epitopes with its domains. Three conformational/discontinuous and 92 linear/continuous B-cell epitopes from the MEV were predicted without altering the prediction parameters of Ellipro and ABCPred 2.0 (Supplementary Tables 9-10).

### 3.9. Immunogenicity evaluation of multiepitope based vaccine

The *in silico* immune system simulation against MEV showed significant activity of B-cells and T-cells, which was consistent with the actual immune responses. Primary immune response was observed with the increased levels of IgM in the start. Later, secondary and tertiary immune responses were observed comparitively higher than the primary response. The immunoglobulin activity (i.e., IgG1 + IgG2, IgM, and IgG + IgM antibodies) was normally at high levels with a consistent decrease in MEV concentration. The high levels of triggered B-cells and memory B-cell formation was observed, that is indicator of an effective long-lasting immune response produced by the MEV (Figure 5A-D). Similarly, higher levels of HTL and CTL as well as development of memory Th and Tc were observed which is vital for triggering the immune response (Figure 5E-H). The significant levels of T regulatory cells and the continuous and increased proliferation of macrophages and dendritic cells were observed during introduction of MEV. The higher levels of cytokines like IL-2, and IFN-γ were also observed (Figure I-L). These observations indicated the generation of promising antiviral response by the proposed MEV construct.

**Figure 5.**
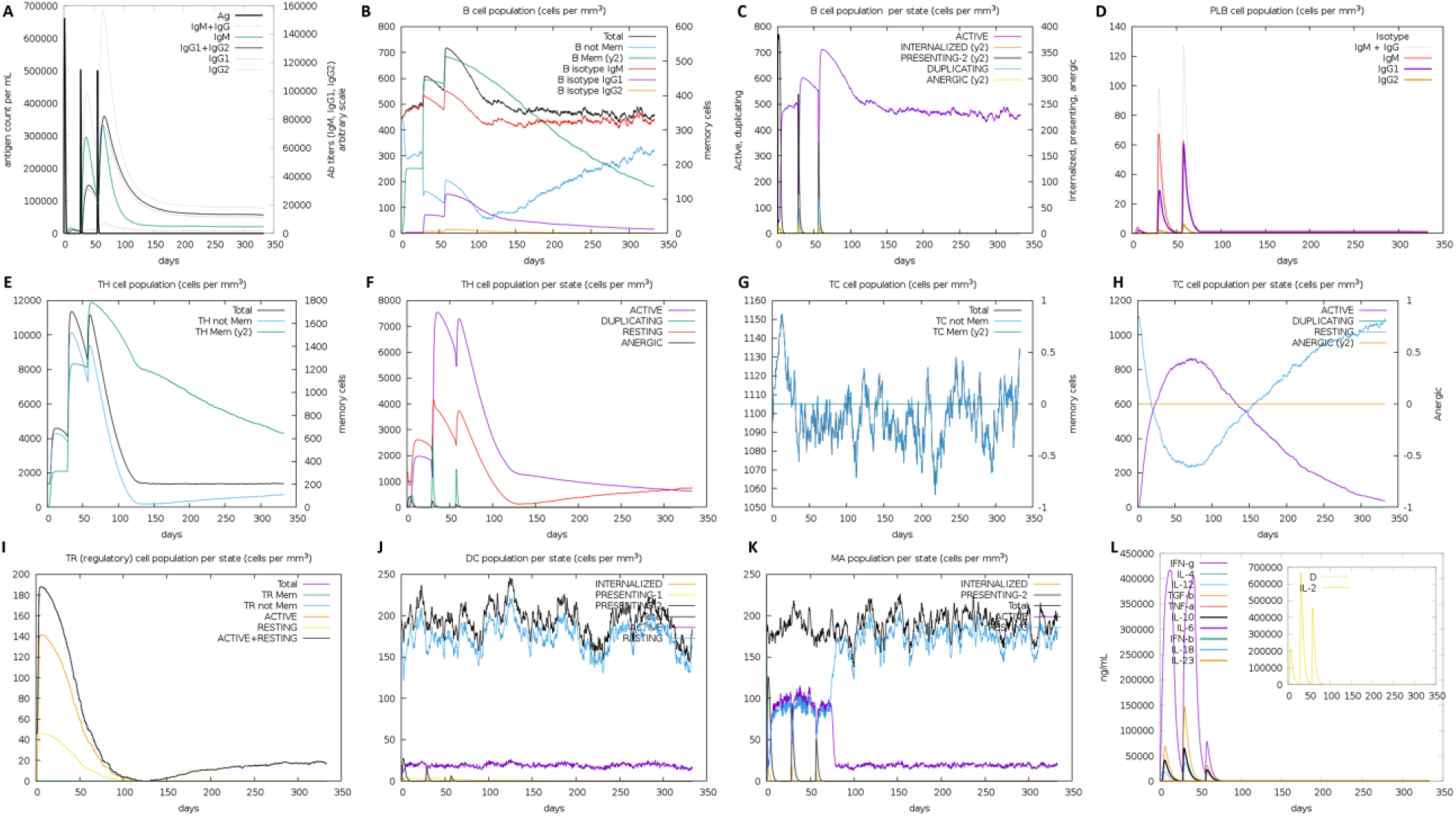
*In silico* simulation of immune response against MEV as an antigen: (**A**) Antigen and immunoglobulins, (**B**) B-cell population, (**C**) B-cell population per state, (**D**) Plasma B-cell population, (**E**) Helper T-cell population, (**F**) Helper T-cell population per state, (**G**) Cytotoxic T-cell population, (**H**) Cytotoxic T-cell population per state, (**I**) Reduced levels of T regulatory cells, (**J**) Dendritic cell population per state, (**K**) Macrophage population per state, and (**L**) Different levels of cytokine and interleukins with Simpson index (D) of immune response.

### 3.10. Molecular docking of multiepitope based vaccine with TLR3 and TLR8

An appropriate association between immune receptor molecules and the antigen molecule is essential to activate an immune responsiveness. HADDOCK server has thus been used to perform the docking of the MEV with human immune receptors TLR3 and TLR8. TLR3 and TLR8 can efficiently induce the immune response after virus recognition. The docking analysis showed good interactions between the MEV and TLR3/TLR8. The binding scores of MEV-TLR3 and MEV-TLR8 were −293.90 Kcal/mol and −283.20 Kcal/mol, respectively. TLR3 is shown in the orange color, while the MEV is shown in the blue color, respectively, in Figure 6A. It was observed that MEV made 11 hydrogen bond interactions within range of 3.00 Å with TLR3 Figure 6B-C. MEV interacting amino acids with hydrogen bonding to TLR3 shown in green color stick representation, while similarly TLR3 amino acids interacting through hydrogen bonding with MEV shown in red color stick representation.

**Figure 6.**
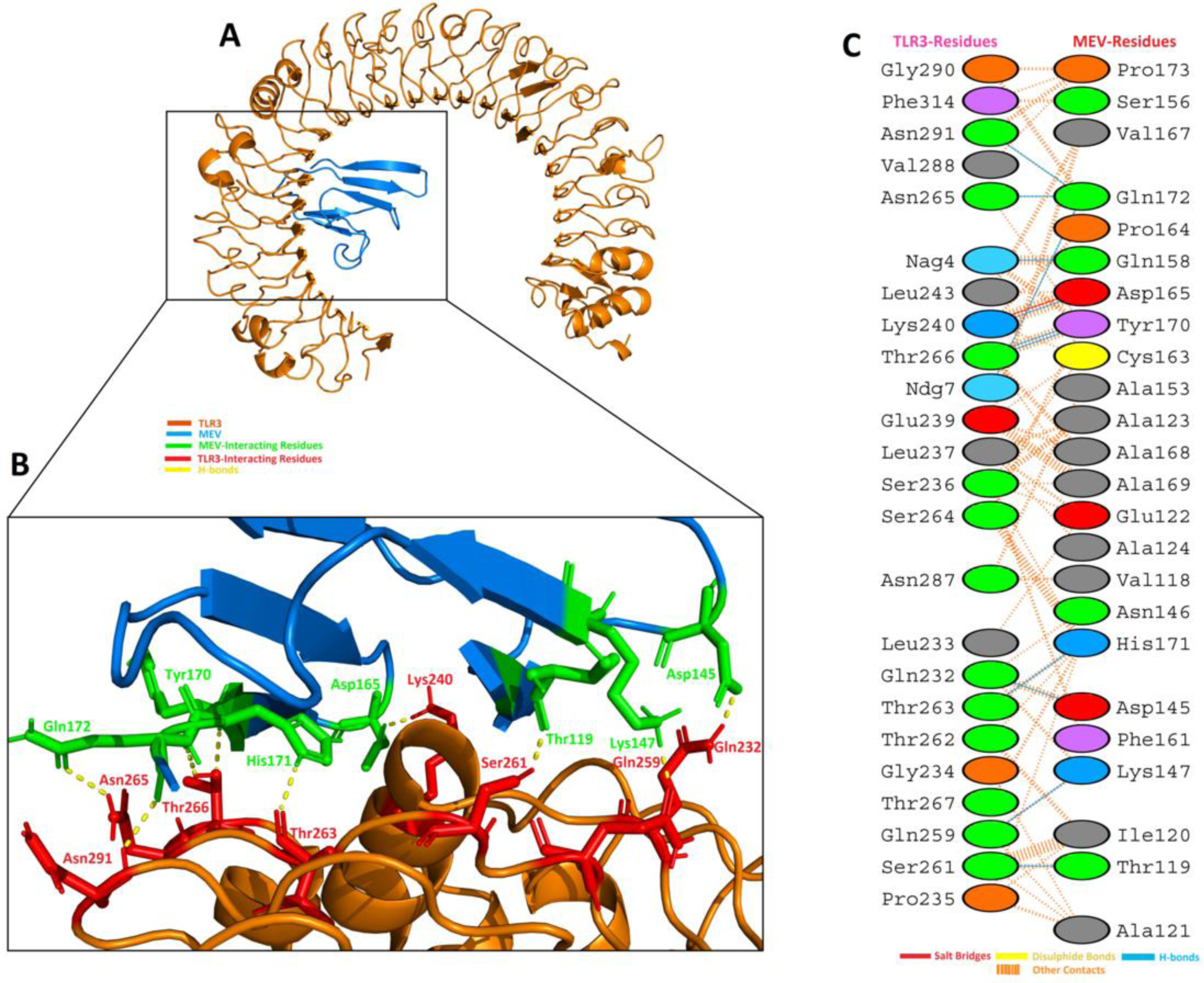
MEV construct docking with human TLR3: (**A**) MEV-TLR3 docked complex in cartoon representation. TLR3 displayed with brown color and MEV vaccine construct displayed with blue color. (**B**) Interacting residues illustration between MEV and TLR3 complex. Interacting residues of MEV are highlighted with green color stick representation, while interacting residues of TLR3 are highlighted with red color stick representation. Hydrogen bonds are represented with yellow color dotted lines. (**C**) All interacting residues of MEV and TLR3. Hydrogen bonds are shown with blue color lines. The colors of interacting residues are representing properties of amino acids (Positive: Blue, Negative: Red, Neutral: Green, Aliphatic: Grey, Aromatic: Pink, Pro&Gly: Orange and Cys: Yellow).

In case of TLR8, it is shown in the grey color, while the MEV is shown in the blue color, respectively, in Figure 7A. It was observed that MEV made 9 hydrogen bond interactions within range of 3.00 Å with TLR8 Figure 6B-C. Similar to TLR3, MEV interacting amino acids with hydrogen bonding to TLR8 shown in green color stick representation, while TLR8 amino acids interacting through hydrogen bonding with MEV shown in red color stick representation. These results indicated that this MEV is best suitable candidate for vaccine production.

**Figure 7.**
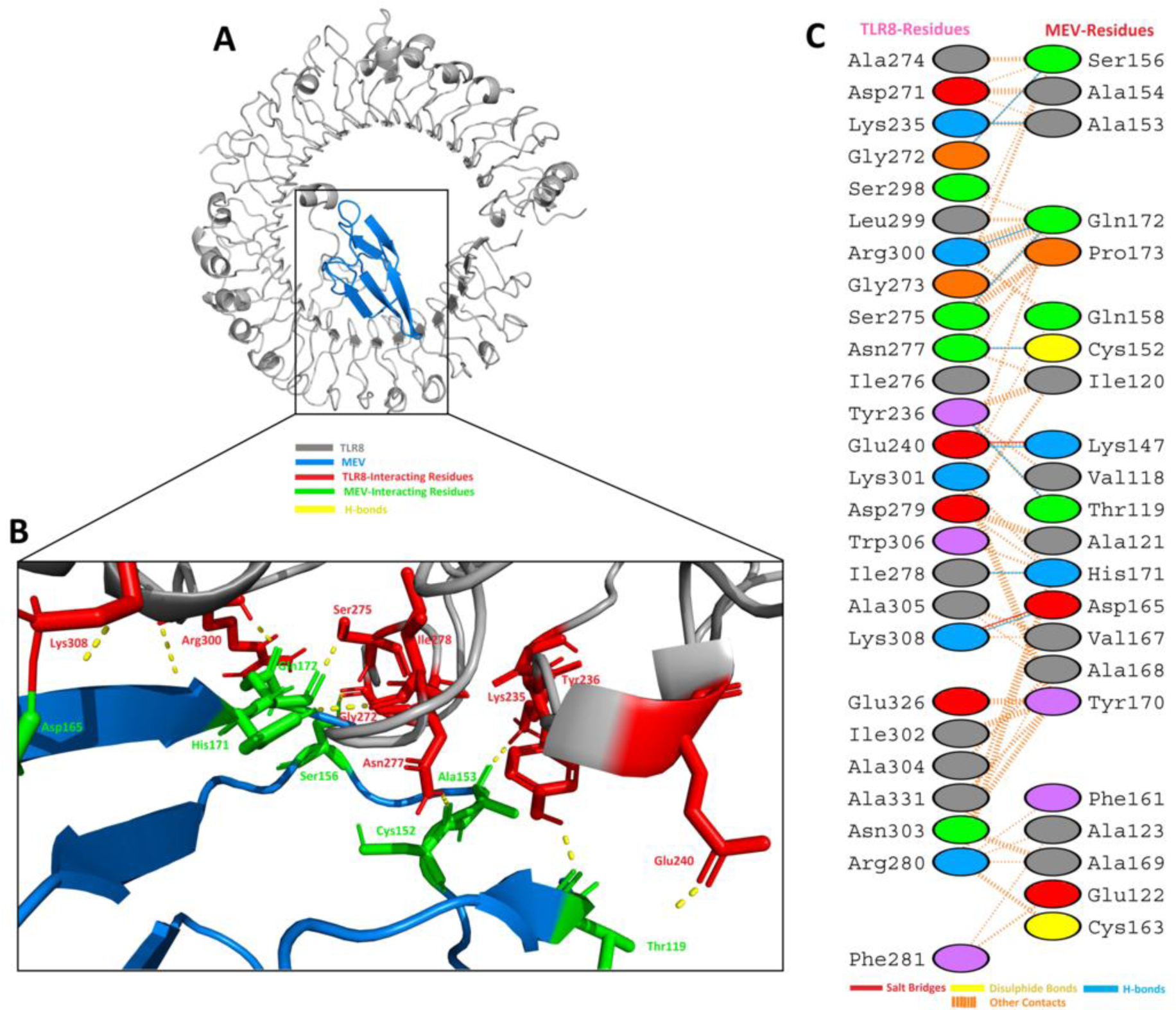
MEV construct docking with human TLR8: (**A**) MEV-TLR8 docked complex in cartoon representation. TLR8 displayed with grey color and MEV vaccine construct displayed with blue color. (**B**) Interacting residues illustration between MEV and TLR8 complex. Interacting residues of MEV are highlighted with green color stick representation, while interacting residues of TLR8 are highlighted with red color stick representation. Hydrogen bonds are represented with yellow color dotted lines. (**C**) All interacting residues of MEV and TLR8. Hydrogen bonds are shown with blue color lines. The colors of interacting residues are representing properties of amino acids (Positive: Blue, Negative: Red, Neutral: Green, Aliphatic: Grey, Aromatic: Pink, Pro&Gly: Orange and Cys: Yellow).

### 3.11. MD simulation of TLR3-MEV and TLR8-MEV complexes

MD simulation is a common approach used to analyse the micro-interactions between ligand/vaccine and protein/receptor structures [66, 78]. In order to further assess MEV dynamics and stability, its docking complexes with TLR3 and TLR8 were simulated by 20 ns MD followed by Root Mean Square Deviations (RMSD) and Root Mean Square Fluctuations (RMSF) analysis. The values of RMSD of backbone atoms were computed to monitor the structural stability of MEV and immune receptor complexes (Figure 8A). The RMSD average values for both complexes TLR3-MEV and TLR8-MEV are 4.5 ± 0.02 Å and 5.6 ± 0.02 Å, respectively. There were no significant variations noticed in both docked complexes. Both the systems remain stable throughout 20 ns simulation. To further compute, the residual and side-chain flexibility, RMSF over 20 ns time were computed. Little fluctuations at the terminal constituent residues of both complexes were observed, while middle residues depicted stable behavior with an average 1.5 ± 0.02 Å RMSF (Figure 8B). In general, both complexes were stable without obvious fluctuations. These results validate the docking interaction analysis and endorsed that MEV can strongly bind with immune receptors to generate significant immune response against SARS-CoV-2.

**Figure 8.**
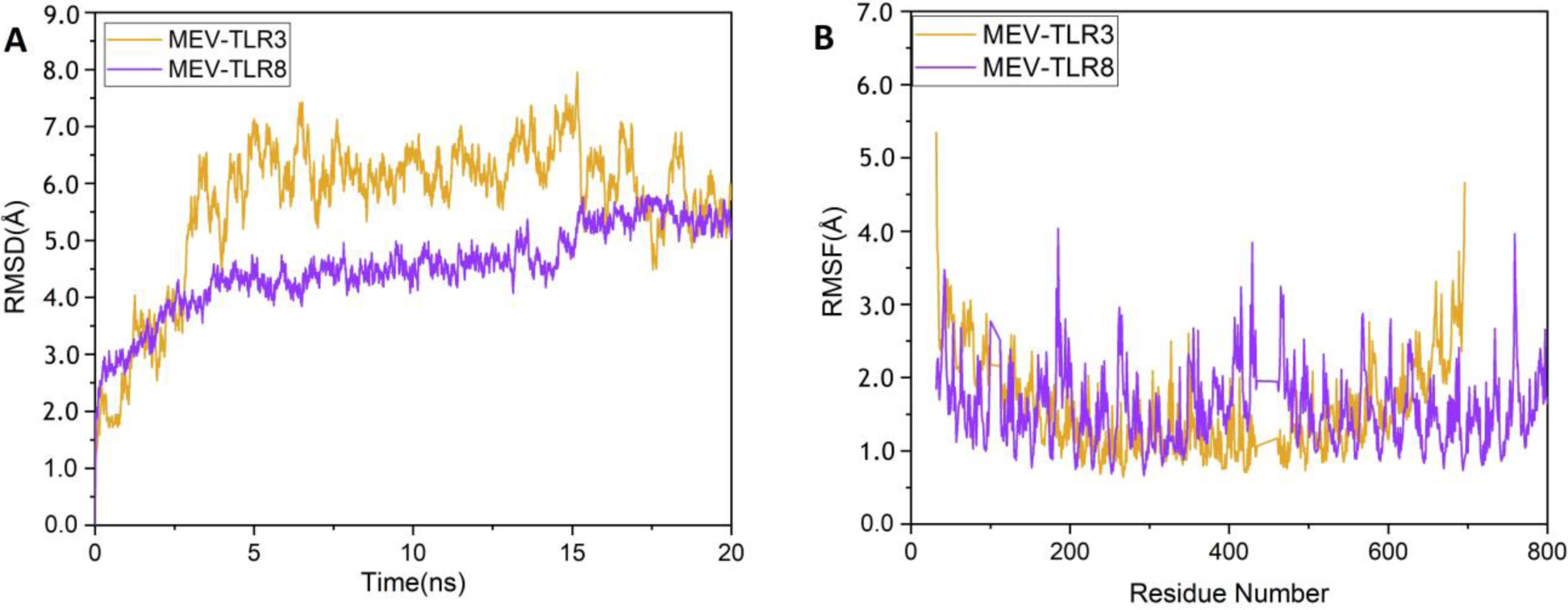
MD simulation at 20 ns results: (A) The RMSD plot of the MEV-TLR3 and MEV-TLR8 complexes. MEV-TLR3 complex RMSD represented with purple color line while MEV-TLR8 complex RMSD is represented with orange color line. (B) The RMSF plot of the MEV-TLR3 and MEV-TLR8 complexes. MEV-TLR3 complex RMSF represented with purple color line while MEV-TLR8 complex RMSF is represented with orange color line.

### 3.12. In silico cloning

*In silico* cloning was performed to ensure expression of SARS-CoV-2 derived MEV in commonly used *E. coli* host. First, codons of MEV construct were adapted as per codon usage of *E. coli* expression system. JCAT server was used to optimize the MEV codons according to *E. coli* (strain K12). The optimized MEV construct contained 1515 nucleotides (Supplementary Table 11), an ideal range of GC content 56.30% (30-70%) and CAI value 0.93 (0.8-1.0), and showing the high possibility of positive protein expression and reliability. In next step, buffer compatible restriction enzymes XhoI and HindIII restriction sites were added to the both ends of MEV optimized nucleotide sequence to aid the cloning/purification process. Finally, refined MEV sequence was cloned between XhoI and HindIII restriction sites at the multiple cloning site of pET28a (+) vector (Figure 9). The total length of the clone was 6.875 kbp.

**Figure 9.**
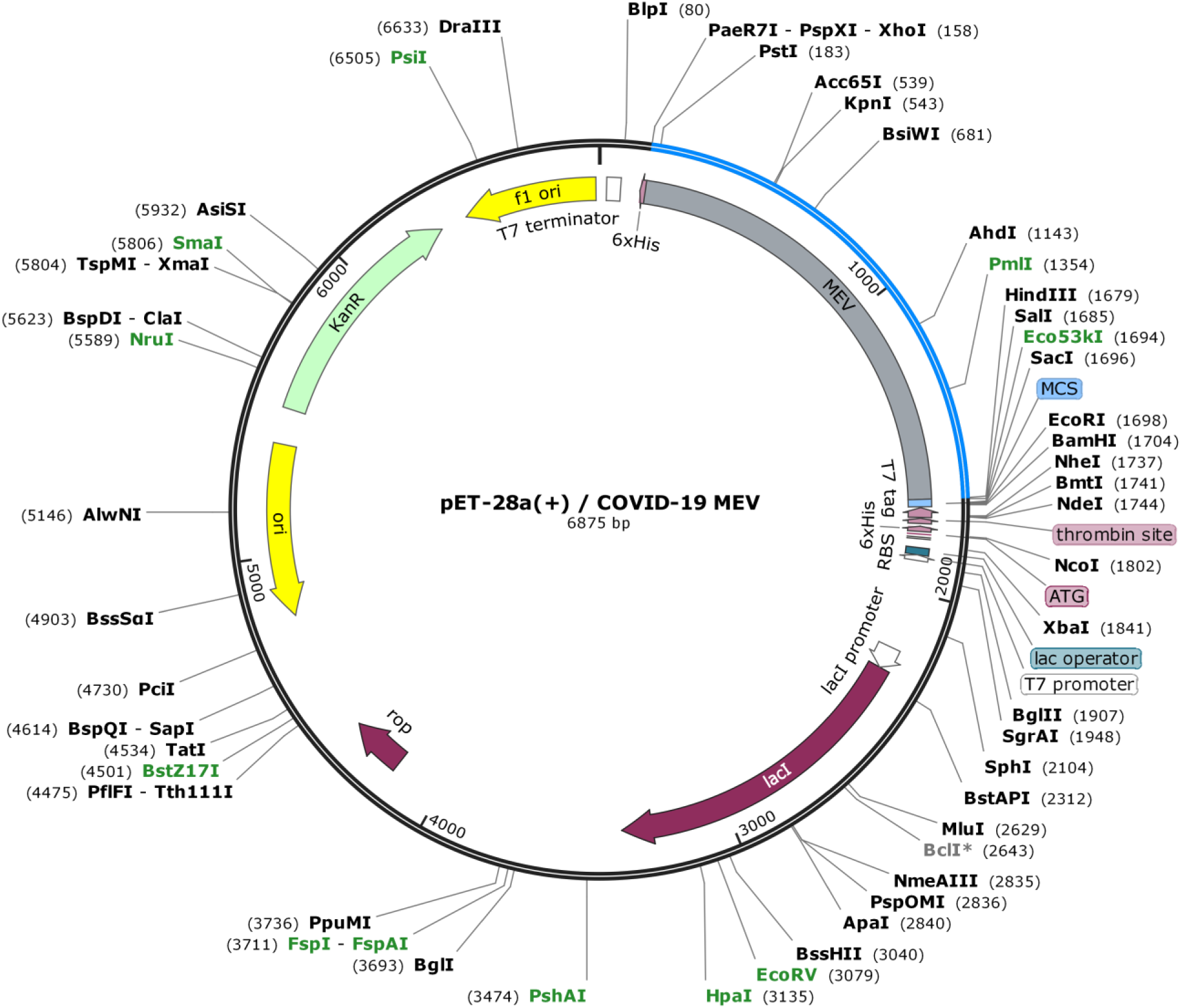
MEV codon optimized nucleotide sequence for *E. coli* expression system, *in silico* cloned at multiple cloning site (MCS) of pET28a(+) vector, shown with blue color in vector backbone (black color) and with grey color functional annotation. MEV cloned sequence is surrounded by XhoI (158) and HindIII (1679) restriction enzyme site, and 6-histidine residues.

## 4. Discussion

Vaccination has many useful effects for improving people health at a low-cost and best aid to inhibit transmission of diseases around the world. However, the development and production of the vaccine is labour-intensive and costly. Immunoinformatics approaches can reduce this burden. Today researchers are searching methods for the development of subunit vaccines from complete genome/proteome of pathogens [79]. Epitope prediction for antibodies becomes more significant with the advancement of the computational tools for designing a vaccine [80]. In the field of bioinformatics, Immuno-informatics is a sub-branch that includes a lot of tools & databases. Immunological datasets prediction and *in silico* analysis are done with the help of those tools. With the advancement of tools and a variety of data availability like genomic, proteomic, and different algorithms made it more effective for scientists to accurately predict epitopes that are much effective in the development of the sub-unit vaccines [12, 13, 81].

An outbreak of SARS-COV-2 in late December 2019 that later turned into a global pandemic resulted in thousands of deaths around the globe [6]. After the outbreak, remarkable progress has been made towards the structural genomics, proteomics and drug repurposing against the SARS-CoV-2, yet there is no exact cure has been identified for COVID-19. Till date, no effective vaccine is available against SARS-CoV-2. Therefore, COVID-19 vaccine need to be designed so that global pandemic situation could be controlled effectively.

The recent study was conducted to design a multiepitope based vaccine (MEV) against SARS-COV-2 by using immunoinformatics and *in silico* approaches. MEV is advantageous compared to monovalent vaccine because it can elicit humoral, innate and cellular immunity responses together [82]. Three different studies have been reported yet, which reported lists of potential epitopes using Surface glycoprotein, Envelope protein, and Membrane glycoprotein [10, 47]. However, no other sub-unit vaccine construct has been reported against SARS-CoV-2 to date. The current MEV construct is therefore is very important to curb COVID-19. Amino acid sequences of ten proteins of SARS-COV-2 were taken from Genbank and their antigenicity was checked. Highly antigenic proteins were selected for further analysis. After the complete physiochemical analysis of antigenic proteins, B-cell, T-cell and IFN-γ inducing epitopes were predicted. Antigenic and overlapped T-cell epitopes with B-cells and IFN-γ epitopes were further validated using molecular docking with HLA-B7 allele and their population coverage was estimated. Finally, MEV was designed with the help of linkers and adjuvant. An adjuvant was added to the N-terminal of the MEV and epitopes were linked with the help of AAY and GPGPG linkers. Adjuvant was added to increase the immunogenicity of the vaccine [83]. Linkers were added to help maintain the function of each epitope so that after being imported into the human body they can function independently/properly [35, 84].

MEV was found to be highly antigenic, immunogenic, non-toxic and non-allergenic indicating an epitope-based vaccine’s potential to cause robust immune responses. During *in silico* immunological validation analysis, higher B and T-cell activity was observed that is rationally identical with typical immune responses. B and T-cell memory formation was evident, and results depicted it last for several months. Molecular docking analysis followed by MD simulation revealed that the MEV is potentially able to properly/firmly occupy the TLR3/TLR8 receptors with minimal energy. Furthermore, the expression of MEV construct inside the host *E. coli* K12 was improved through codon optimization. The results of recent study suggest that the MEV being designed is a potential candidate to undergo *in vivo* and *in vitro* experimental analysis to develop a potential vaccine against COVID-19.

## 5. Conclusion

Recent global pandemic of SARS-CoV-2 claimed hundreds of precious lives in various regions of the world and crumble the economies of several countries. There is no drug or vaccine reported against SARS-CoV-2 yet. Few antiviral drugs and vaccines are in clinical trials, but none have yet been declared as clinically approved anti-COVID-19 therapeutic. In this study, a successful attempt was made to design a sub-unit MEV against SARS-COV-2. Immunoinformatics and *in silico* approaches were used to develop a potential and safe MEV that could trigger three types of the immune responses: humoral, innate and cellular. A highly immunogenic, safe, stable and strongly interacting with human receptors, MEV has been reported in present study that could be a potential candidate for vaccine production. However, current research is the result of an integrated vaccinomics approach. Therefore, further experimental research by the vaccinologists is required to demonstrate the efficacy of the designed vaccine.

## Acknowledgements

This work was supported by the Starting Research Grant for High-level Talents from Guangxi University and Postdoctoral research platform grant of Guangxi University.

## Authors’ contributions

MTQ, LLC and UAA conceived and designed this study; MTQ and AR performed the experiments; MQA, IF and FS analyse the results; MTQ and AR wrote the manuscript; UAA and LLC improved and revised the manuscript, and all the authors approved the final version.

## Supplementary data

**Supplementary Figure 1**. Cartoon 3D structural representation of SARS-CoV-2 antigenic proteins. (**A**) E protein tertiary structure, (**B**) M protein tertiary structure, (**C**) N protein tertiary structure, (**D**) ORF6 protein tertiary structure, (**E**) ORF7a protein tertiary structure, (**F**) ORF8 protein tertiary structure and (**G**) ORF10 protein tertiary structure.

**Supplementary Figure 2**. The 3D binding pattern of the selected 16 MHC Class I epitopes (meshed-pink) docked with HLA-B7 allele (cyan). Hydrogen bond interactions are highlighted with yellow color dotted lines. Numbering 1-16 is consistent with the Table 1 of main text.

**Supplementary Figure 3**. The 3D binding pattern of the selected 11 MHC Class II epitopes (purple blue) docked with HLA-B7 allele (cyan). Hydrogen bond interactions are highlighted with yellow color dotted lines. Numbering 1-11 is consistent with the Table 1 of main text.

**Supplementary Figure 4**. Secondary structure depiction of MEV construct.

**Supplementary Figure 5**. MEV structural flexibility results. (**A**) Cartoon representation of top 10 final models showing obvious fluctuation throughout. (**B**) MEV residue-residue interaction/contact map. The interactive area is represented in the central panel. (**C**) RMSF plot representing the obvious fluctuations of MEV residues during simulation.

**Supplementary Table 1.** Physiochemical properties of the SARS-COV-2 proteins.

**Supplementary Table 2.** Secondary structure of the SARS-COV-2 proteins.

**Supplementary Table 3.** 3D structural details of selected SARS-COV-2 proteins.

**Supplementary Table 4.** Complete list of predicted potential linear B-cell epitopes of SARS-COV-2 proteins.

**Supplementary Table 5.** List of predicted potential conformational B-cell epitopes of SARS-COV-2 proteins.

**Supplementary Table 6.** List of predicted potential MHC Class I epitopes of SARS-COV-2 proteins.

**Supplementary Table 7.** List of predicted potential MHC Class II epitopes of SARS-COV-2 proteins.

**Supplementary Table 8.** List of predicted potential IFN-γ inducing epitopes of SARS-COV-2 proteins.

**Supplementary Table 9.** Linear B-cell epitopes in the final MEV construct.

**Supplementary Table 10.** Conformational B-cell epitopes in the final MEV construct.

**Supplementary Table 11.** Codon optimized nucleotide sequence of MEV construct for cloning in *E. coli* strain K12. Green color bold sequence at 5’ site (N-terminal) is representing XhoI restriction enzyme site, while maroon color bold sequence at 3’ site (C-terminal) is representing HindIII restriction enzyme site.

