## Supplementary material for "Designing of a next generation multiepitope based vaccine (MEV) against SARS-COV-2: Immunoinformatics and *in silico* approaches": Supplementary Table 1.docx

Supplementary Table 1. Physiochemical properties of the SARS-COV-2 proteins.

| Proteins | Molecular Weight | | | Theoretical pI | Instability index | Half‐life | Stability Profiling | Aliphatic Index | Grand Average of Hydropathy |
| --- | --- | --- | --- | --- | --- | --- | --- | --- | --- |
| **Envelope protein** | 8365.04 | | | 8.57 | 38.68 | 30 hours (mammalian reticulocytes, in vitro). >20 hours (yeast, in vivo). >10 hours (Escherichia coli, in vivo). | stable | 144.00 | 1.128 |
| **Membrane glycoprotein** | 25146.62 | | | 9.51 | 39.14 | 30 hours (mammalian reticulocytes, in vitro). >20 hours (yeast, in vivo). >10 hours (Escherichia coli, in vivo). | stable | 120.86 | 0.446 |
| **ORF6 protein** | | 7272.54 | | 4.60 | 31.16 | 30 hours (mammalian reticulocytes, in vitro). >20 hours (yeast, in vivo). >10 hours (Escherichia coli, in vivo). | stable | 130.98 | 0.233 |
| **ORF7a** | | 13744.17 | | 8.23 | 48.66 | 30 hours (mammalian reticulocytes, in vitro). >20 hours (yeast, in vivo). >10 hours (Escherichia coli, in vivo) | unstable | 100.74 | 0.318 |
| **ORF8 protein** | | 13831.01 | | 5.42 | 45.79 | 30 hours (mammalian reticulocytes, in vitro). >20 hours (yeast, in vivo) >10 hours (Escherichia coli, in vivo). | unstable | 97.36 | 0.219 |
| **Nucleocapsid phosphoprotein** | | | 45625.70 | 10.07 | 55.09 | 30 hours (mammalian reticulocytes, in vitro). >20 hours (yeast, in vivo). >10 hours (Escherichia coli, in vivo). | unstable | 52.53 | -0.971 |
| ORF10 protein | | | 4449.23 | 7.93 | 16.06 | 30 hours (mammalian reticulocytes, in vitro).  >20 hours (yeast, in vivo). >10 hours (Escherichia coli, in vivo). | stable | 107.63 | 0.637 |
