## Supplementary material for "Designing of a next generation multiepitope based vaccine (MEV) against SARS-COV-2: Immunoinformatics and *in silico* approaches": Supplementary Table 2.docx

Supplementary Table 2. Secondary structure of the SARS-COV-2 proteins.

| **Proteins** | **Sequence Length** | **α-helix** | **β-Turn** | **Random Coils** |
| --- | --- | --- | --- | --- |
| **Envelope** | 75 | 44.00% | 9.33% | 20.00% |
| **M glycoprotein** | 222 | 34.68% | 6.76% | 37.39% |
| **ORF6 protein** | 61 | 70.49% | 8.20% | 11.48% |
| **ORF7a** | 121 | 42.98% | 9.92% | 28.10% |
| **ORF8 protein** | 121 | 19.83% | 4.96% | 39.67% |
| **Nucleocapsid** | 419 | 21.24% | 6.92% | 55.13% |
| **ORF10** | 38 | 28.95% | 5.26% | 28.95% |
