## Supplementary material for "Designing of a next generation multiepitope based vaccine (MEV) against SARS-COV-2: Immunoinformatics and *in silico* approaches": Supplementary Table 3.docx

Supplementary Table 3. 3D structural details of selected SARS-COV-2 proteins.

| Proteins | Tool utilized for Modeling | Template | Ramachandran plot | | | Errat Results |
| --- | --- | --- | --- | --- | --- | --- |
|  |  |  | **Favored Region** | **Allowed Region** | **Disallowed Region** | **Quality Factor** |
| E | Swiss model | 5x29.1 | 92.9% | 3.9% | 3.2% | 61.22 |
| M | Swiss model | 6ck1.1 | 98.0% | 2.0% | 0.0% | 100 |
| N | Raptor x | - | 80.2% | 16.0% | 3.7% | 77.46 |
| Orf8 | Swiss model | 1xak.1 | 86.1% | 9.7% | 4.2% | 82.61 |
| Orf7a | Raptor x | - | 87.4% | 7.6% | 5.0% | 88.56 |
| Orf6 | Swiss model | 5vyj.1 | 97.1% | 2.9% | 0.0% | 100 |
| Orf10 | Pep-fold | - | 97.2% | 0.0% | 2.8% | 100 |
