## Supplementary material for "Designing of a next generation multiepitope based vaccine (MEV) against SARS-COV-2: Immunoinformatics and *in silico* approaches": Supplementary Table 4.docx

Supplementary Table 4. Complete list of predicted potential linear B-cell epitopes of SARS-COV-2 proteins.

| Protein | Peptide (Position) | Antigenicity |
| --- | --- | --- |
| E protein | SLVKPSFYVYSRVK (50) | 0.6 |
|  | LCAYCCNIVNVSLV (39) | 1.1 |
|  | ILTALRLCAYCCNI (33) | 0.7 |
|  | FLLVTLAILTALRL (26) | 0.8 |
| Membrane protein | SELVIGAVILRGHL (136) | 0.54 |
|  | TRPLLESELVIGAV (130) | 0.58 |
|  | DIKDLPKEITVATS (160) | O.67 |
|  | PVTLACFVLAAVYR (59) | O.97 |
|  | FLTWICLLQFAYAN (28) | 0.60 |
|  | AAVYRINWITGGIA (68) | 0.88 |
|  | ATSRTLSYYKLGAS (171) | 0.64 |
|  | GGIAIAMACLVGLM (78) | 0.87 |
|  | LEQWNLVIGFLFLT (17) | 0.94 |
|  | MADSNGTITVEELK (1) | 0.62 |
|  | VIGFLFLTWICLLQ (23) | 0.93 |
|  | FRLFARTRSMWSFN (100) | 0.71 |
| ORF6 | FHLVDFQVTIAEIL (2) | 1.2 |
| ORF7a | SSGTYEGNSPFHPL (36) | 0.5 |
|  | FALTCFSTQFAFAC (54) | 1.6 |
|  | VYQLRARSVSPKLF (74) | 0.6 |
|  | PFHPLADNKFALTC (45) | 1.2 |
|  | FSTQFAFACPDGVK (59) | 0.8 |
|  | DGVKHVYQLRARSV (69) | 0.6 |
| ORF8 | EAGSKSPIQYIDIG (64) | 1.2 |
|  | GNYTVSCLPFTINC (77) | 1.6 |
|  | PIHFYSKWYIRVGA (38) | 1.09 |
|  | SKWYIRVGARKSAP (43) | 0.75 |
|  | FLVFLGIITTVAAF (3) | 0.50 |
|  | TQHQPYVVDDPCPI (26) | 0.69 |
|  | VVDDPCPIHFYSKW (32) | 0.50 |
|  | LPFTINCQEPKLGS (84) | 1.34 |
|  | KLGSLVVRCSFYED (94) | 0.79 |
| Nucleocapsid | TNSSPDDQIGYYRR (76) | 0.68 |
|  | GSRGGSQASSRSSS (175) | 0.89 |
|  | DGKMKDLSPRWYFY (98) | 1.13 |
|  | PQNQRNAPRITFGG (6) | 0.73 |
|  | GNFGDQELIRQGTD (284) | 0.55 |
|  | GQTVTKKSAAEASK (243) | 0.53 |
|  | FGMSRIGMEVTPSG (315) | 1.01 |
|  | SQASSRSSSRSRNS (180) | 1.07 |
|  | KAYNVTQAFGRRGP (266) | 0.62 |
|  | LLLLDRLNQLESKM (221) | 0.55 |
|  | LSPRWYFYYLGTGP (104) | 1.37 |
|  | NGERSGARSKQRRP (29) | 0.52 |
|  | FTALTQHGKEDLKF (53) | 1.24 |
|  | KDPNFKDQVILLNK (342) | 1.16 |
|  | EVTPSGTWLTYTGA (323) | 0.69 |
|  | GQQQQGQTVTKKSA (238) | 0.59 |
|  | NSTPGSSRGTSPAR (196) | 0.58 |
|  | GPEQTQGNFGDQEL (278) | 1.19 |
|  | TGAIKLDDKDPNFK (334) | 1.80 |
|  | NQLESKMSGKGQQQ (228) | 1.09 |
|  | GTDYKHWPQIAQFA (295) | 0.65 |
|  | SSSRSRNSSRNSTP (186) | 0.80 |
| ORF10 | CRMNSRNYIAQVDV (19) | 0.67 |
