## Supplementary material for "Designing of a next generation multiepitope based vaccine (MEV) against SARS-COV-2: Immunoinformatics and *in silico* approaches": Supplementary Table 5.docx

Supplementary Table 5. List of predicted potential conformational B-cell epitopes of SARS-COV-2 proteins.

| Protein | Conformational B-cell epitopes | Position of epitopes on protein structure |
| --- | --- | --- |
| E-protein | **A:S60, A:R61, A:V62, A:K63, A:N64, A:L65** | 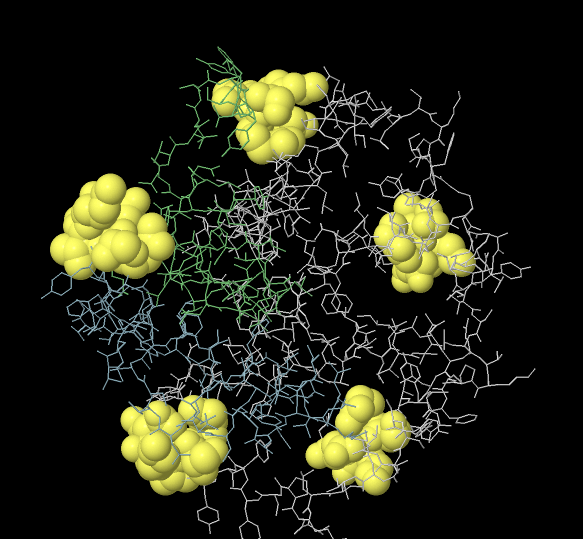 |
|  | A:E8, A:T9, A:G10, A:T11, A:L12, A:I13, A:V14, A:S16 | 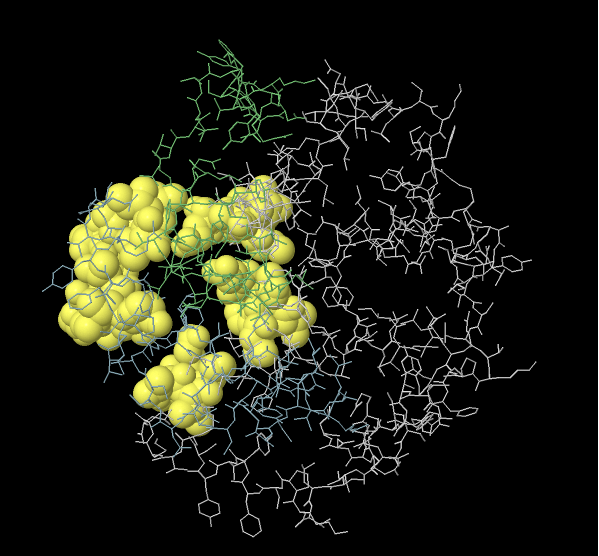 |
|  | **A: L51, A:V52, A:K53, A:S55, A:F56, A:Y59** | 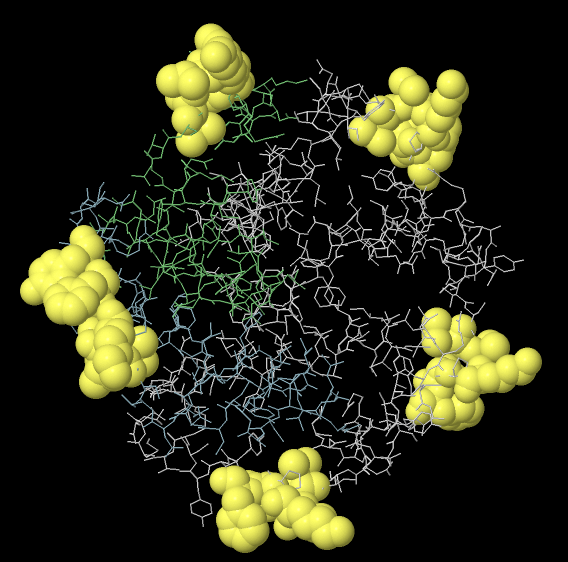 |
|  | A: A32, A:I33, A:T35, A:A36, A:L37, A:R38, A:L39, A:C40, A:A41, A:Y42 | 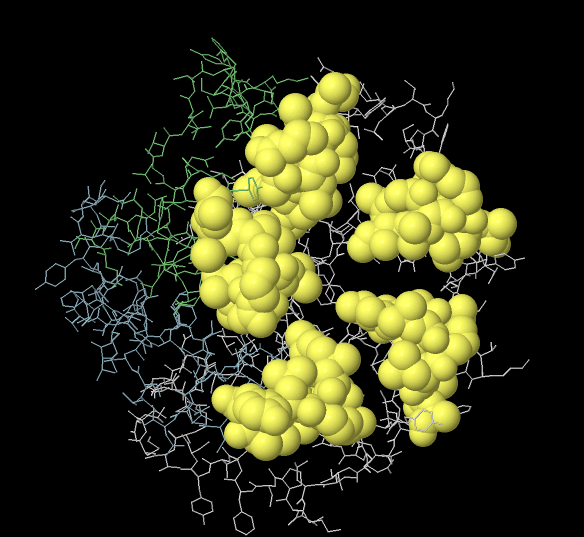 |
| M-protein | **A:F100, A:F103, A:A104, A:R105, A:T106, A:R107, A:S108, A:M109** | 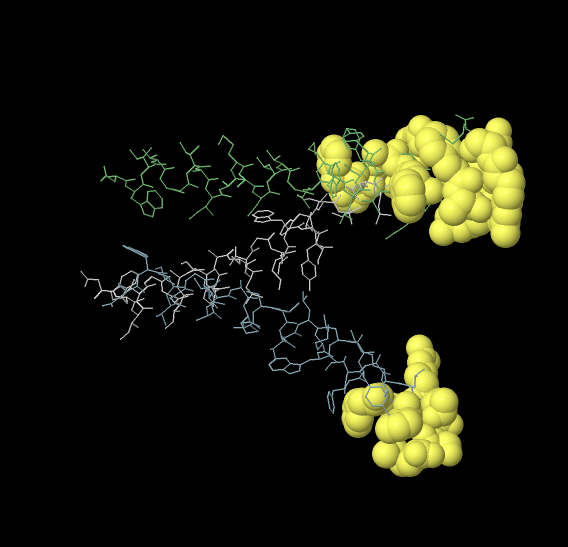 |
|  | A:I76, A:T77, A:G78, A:I80 | 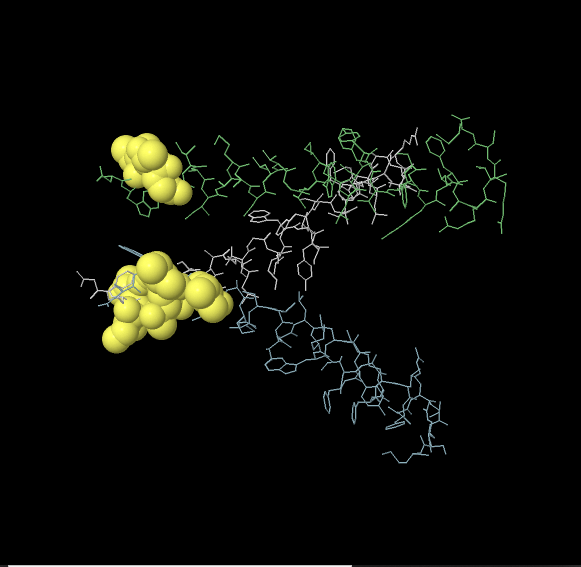 |
| N-protein | **A:R93, A:R95, A:G96, A:G97, A:D98, A:G99, A:K100, A:M101, A:D103, A:L104** | 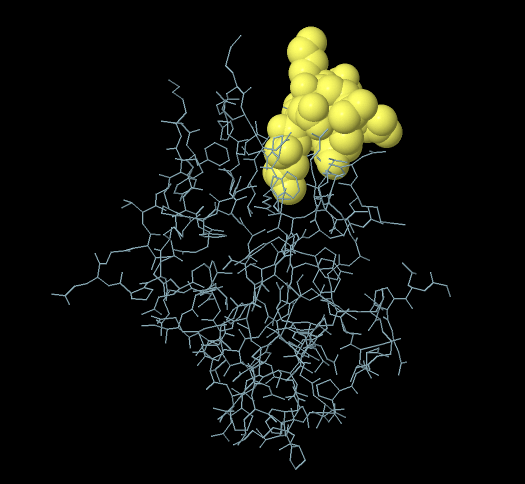 |
|  | **A:Q58, A:H59, A:G60, A:K61, A:E62, A:D63, A:F66, A:P67, A:L161, A:P162, A:Q163, A:G164, A:T165, A:T166, A:L167, A:P168, A:K169, A:G170, A:F171** | 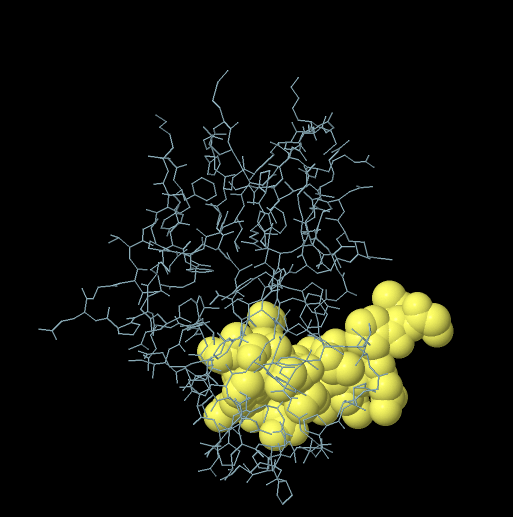 |
|  | A:G69, A:Q70, A:T76, A:N77, A:S78, A:S79, A:P80, A:D81, A:D82, A:G114, A:T115, A:G116, A:P117, A:E118, A:A119, A:G120, A:L121, A:P122, A:G124, A:A125, A:N126, A:K127, A:D128, A:G129, A:T135, A:E136, A:G137, A:A138, A:L139, A:N140, A:T141, A:P142, A:K143, A:D144, A:H145, A:I146 | 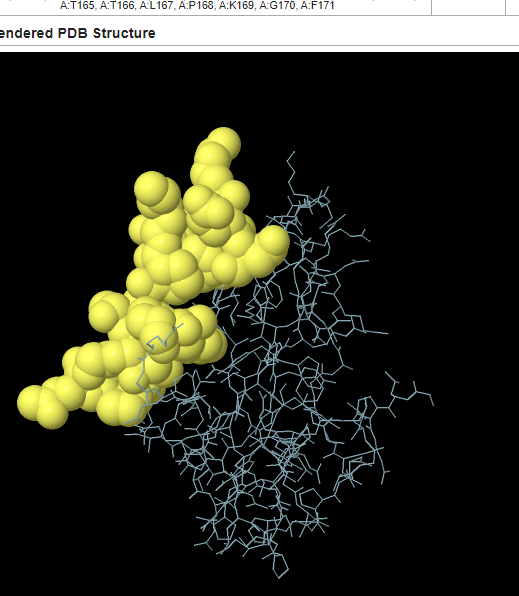 |
|  | A:G44, A:L45, A:P46, A:N47, A:N48, A:T49, A:W52, A:G147, A:T148, A:R149, A:N150, A:P151, A:A152, A:N153, A:N154, A:A155, A:A156 | 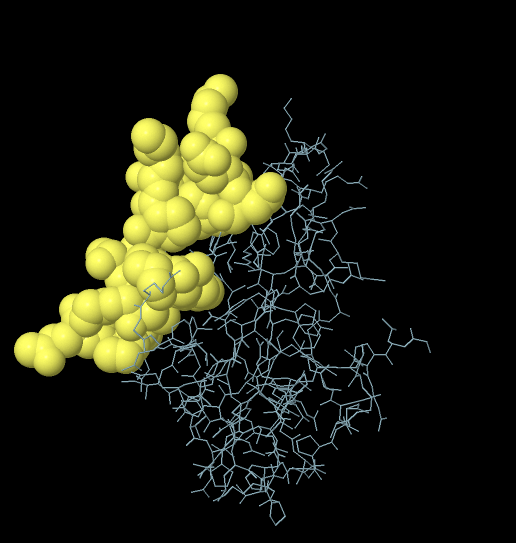 |
| Orf6 | **A:Y49, A:S50, A:D53** | 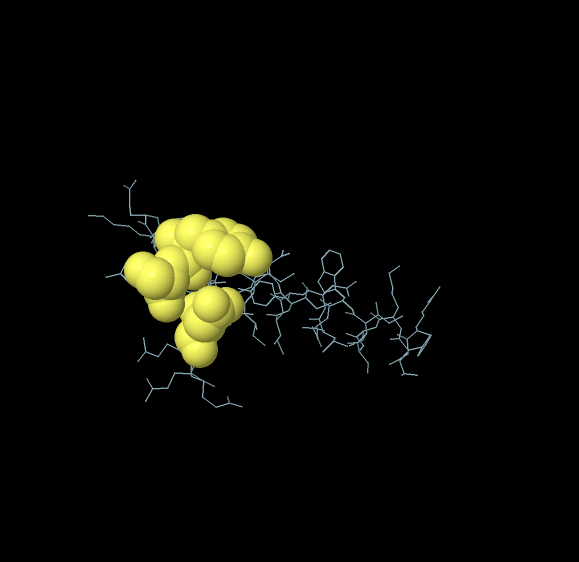 |
|  | A:T21, A:F22, A:K23, A:V24, A:S25 | 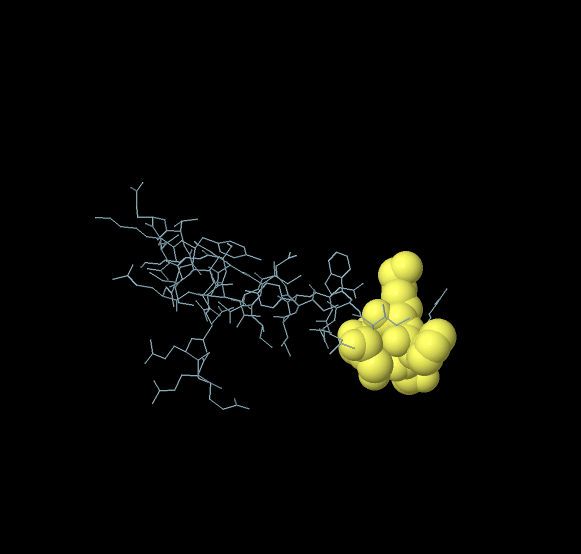 |
|  | **A:T45, A:E46, A:N47** | 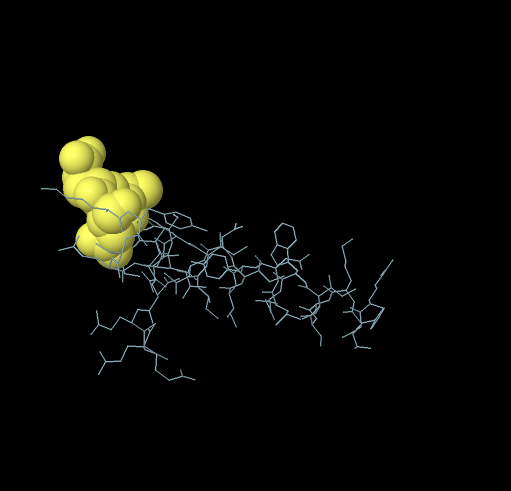 |
| Orf7a | A:S81, A:V82, A:S83, A:P84, A:K85, A:L86, A:F87, A:I88, A:R89, A:Q90, A:E91, A:E92, A:V93, A:Q94 | 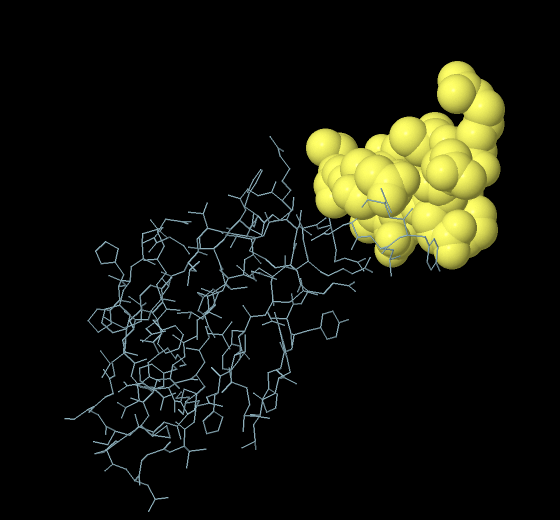 |
|  | A:E16, A:L17, A:C35, A:S36, A:S37, A:G38, A:C67, A:P68, A:D69, A:G70, A:V71, A:K72 | 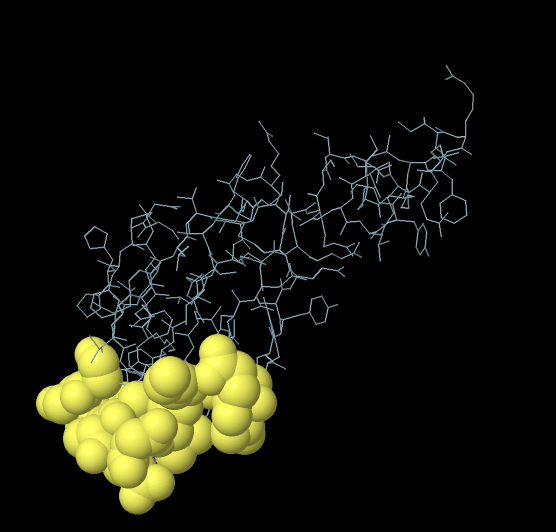 |
|  | A:E33, A:F46, A:H47, A:P48, A:L49, A:A50, A:D51, A:N52, A:K53 | 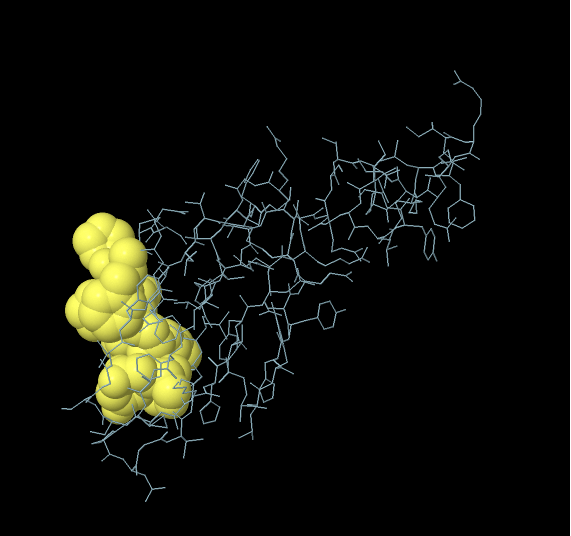 |
|  | A:E95, A:L96, A:Y97 | 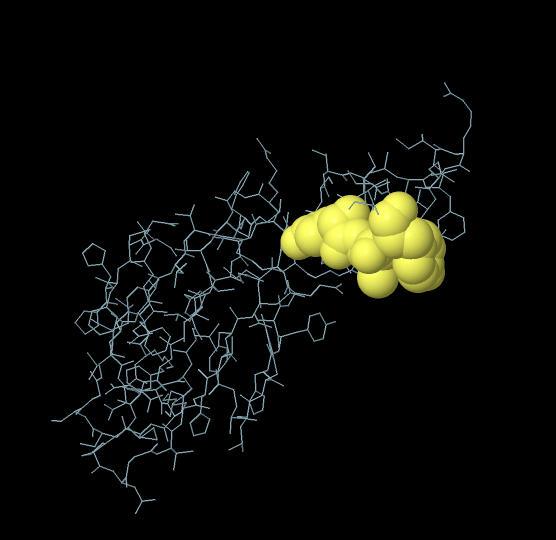 |
| Orf8 | A:L22, A:Q23, A:S24, A:C25, A:T26, A:Q27, A:H28, A:Q29, A:P30 | 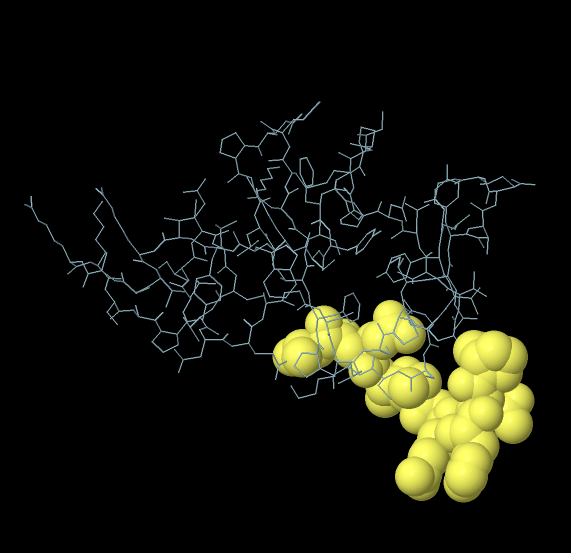 |
|  | A:Y46, A:I47, A:R48, A:V49, A:G50, A:A51, A:R52, A:S54, A:A55, A:P56, A:L57, A:I58, A:V62 | 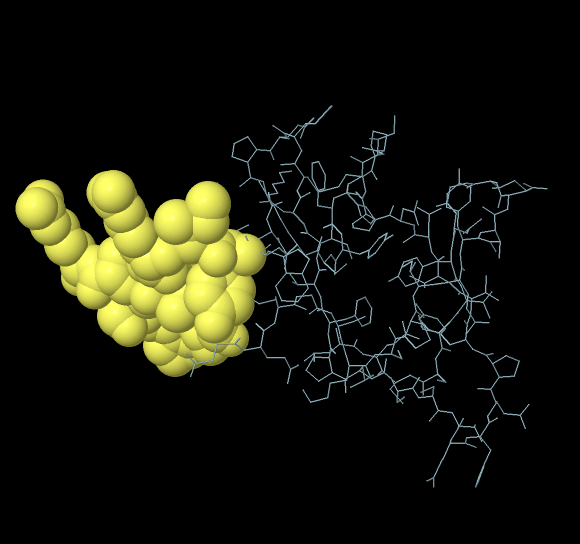 |
|  | A:C90, A:E92, A:P93, A:K94 | 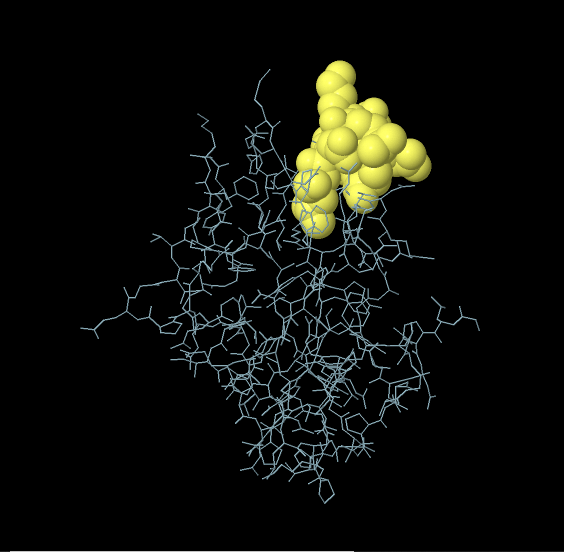 |
|  | A:D34, A:D35, A:Y73, A:I74, A:D75, A:I76, A:G77, A:N78 | 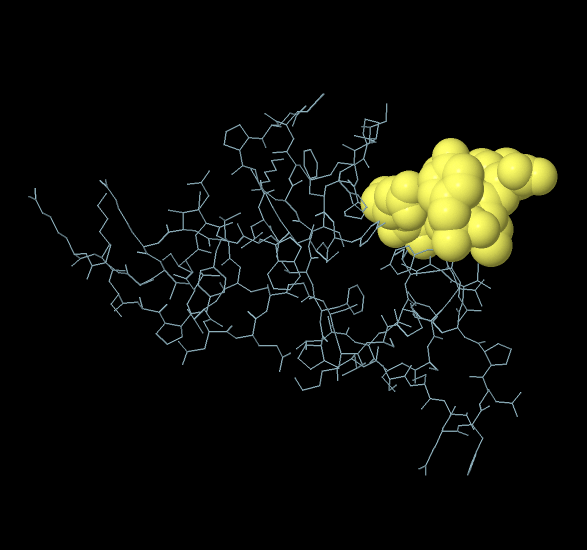 |
| Orf10 | :G2, _:Y3, _:Y14, _:L18, _:M21, _:N22, _:S23, _:R24, _:N25, _:Y26, _:I27, _:A28, _:Q29 | 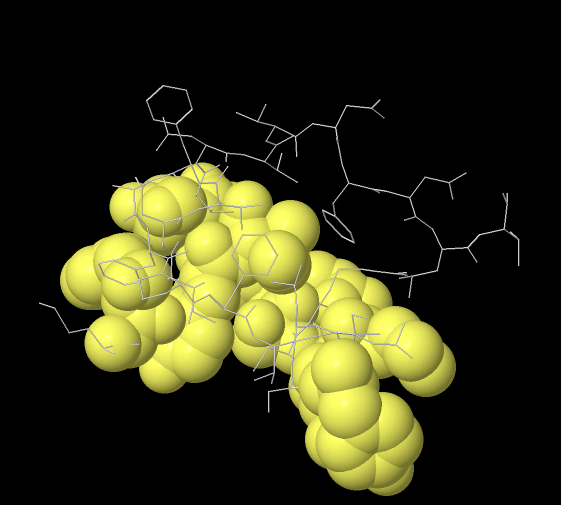 |
|  | _:N5, _:F7, _:A8, _:F9, _:P10, _:F11 | 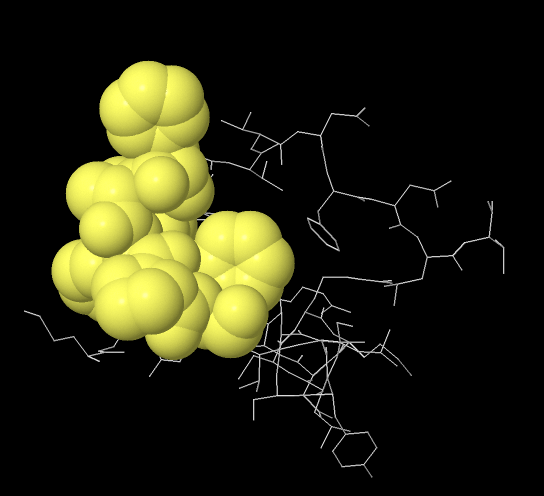 |
|  | _:V6, _:V32, _:V33, _:N34, _:F35, _:N36, _:L37, _:T38 | 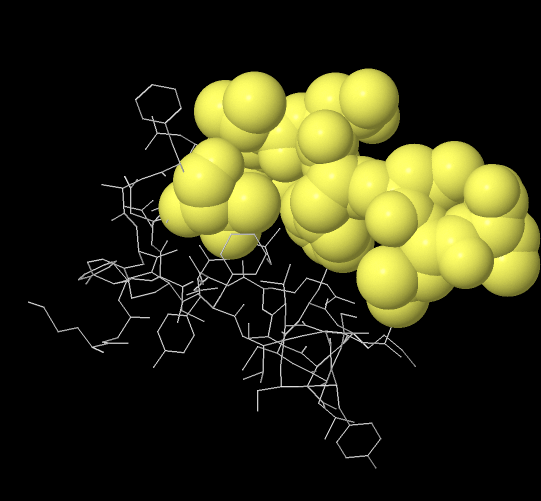 |
