## Supplementary material for "Designing of a next generation multiepitope based vaccine (MEV) against SARS-COV-2: Immunoinformatics and *in silico* approaches": Supplementary Table 6.docx

Supplementary Table 6. List of predicted potential MHC Class I epitopes of SARS-COV-2 proteins.

| Proteins | Epitopes | Position | Allele | Antigenicity |
| --- | --- | --- | --- | --- |
| Envelope Protein | FLLVTLAILTAL | 26-37 | HLA-A*02:01  HLA-A*02:06 | 0.6 |
|  | LLFLAFVVFLLV | 18-29 | HLA-A*02:01  HLA-B*51:01 | 0.6 |
|  | FLAFVVFLLVTL | 20-31 | HLA-A*02:01  HLA-B*51:01 | 0.7 |
|  | NVSLVKPSFYVY | 48-59 | HLA-A*30:02  HLA-A*29:02 | 0.6 |
|  | AILTALRLCAYC | 32-43 | HLA-A*25:01  HLA-A*01:01 | 0.7 |
|  | LFLAFVVFLLVT | 19-30 | HLA-A*02:01  HLA-B*51:01 | 0.6 |
|  | VYSRVKNLNSSR | 58-69 | HLA-A*31:01 | 0.5 |
|  | VSLVKPSFYVYS | 49-60 | HLA-A*30:02 | 0.5 |
|  | PSFYVYSRVKNL | 54-65 | HLA-C*14:02  HLA-C*07:01  HLA-C*06:02 | 0.8 |
| Membrane Protein | FRLFARTRSMWS | 100-111 | HLA-C*07:01  HLA-B*27:05  HLA-B*14:02  HLA-A*32:01 |  |
|  | RLFARTRSMWSF | 101-112 | HLA-A*32:01  HLA-B*08:01  HLA-B*46:01  HLA-B*27:05  HLA-B*15:01  HLA-B*57:01  HLA-A*24:02  HLA-A*23:01 | 0.5 |
|  | LFARTRSMWSFN | 102-113 | HLA-B*46:01  HLA-B*08:01  HLA-A*24:02 | 0.9 |
|  | ITVATSRTLSYY | 168-179 | HLA-A*01:01  HLA-A*30:02  HLA-A*26:01  HLA-A*29:02  HLA-B*57:01 | 0.7 |
| Orf6 | FKVSIWNLDYII | 22-33 | HLA-A*32:01 | 0.5 |
|  | FHLVDFQVTIAE | 2-13 | HLA-B*38:01 | 1.5 |
| Orf7a | HPLADNKFALTC | 58-12 | HLA-B*35:03  HLA-B*39:01  HLA-B*07:02 | 1.3 |
|  | GTYEGNSPFHPL | 38-49 | HLA-B*40:01  HLA-B*40:02  HLA-B*15:02 | 0.6 |
|  | STQFAFACPDGV | 61-71 | HLA-A*68:02 | 0.9 |
| Orf8 | HQPYVVDDPCPI | 28-39 | HLA-B*51:01 | 0.5 |
|  | GNYTVSCLPFTI | 77-88 | HLA-A*24:02 | 1.7 |
|  | DDPCPIHFYSKW | 34-45 | HLA-B*53:01 | 0.8 |
|  | LPFTINCQEPKL | 84-95 | HLA-B*35:03  HLA-B*51:01 | 1.1 |
|  | DFLEYHDVRVVL | 107-118 | HLA-B*40:01  HLA-B*18:01  HLA-C*07:02 | 0.6 |
|  | PIHFYSKWYIRV | 38-49 | HLA-A*31:01 | 0.7 |
|  | LEYHDVRVVLDF | 109-120 | HLA-B*18:01  HLA-B*44:02 | 1.0 |
| Nucleocapsid Protein | TATKAYNVTQAF | 263-274 | HLA-B*46:01 | 0.5 |
|  | KMKDLSPRWYFY | 100-111 | HLA-A*29:02  HLA-E*01:01  HLA-A*32:01  HLA-A*01:01  HLA-A*03:01  HLA-A*31:01  HLA-B*15:01 | 1.4 |
|  | MEVTPSGTWLTY | 322-333 | HLA-B*35:01  HLA-B*53:01  HLA-B*44:03  HLA-B*18:01  HLA-A*01:01  HLA-B*44:02  HLA-A*26:01  HLA-A*29:02 | 0.7 |
|  | DPNFKDQVILLN | 343-354 | HLA-B*35:03 | 1.3 |
| Orf10 | CRMNSRNYIAQV | 19-30 | HLA-C*06:02  HLA-B*27:05 | 0.6 |
|  | FAFPFTIYSLLL | 7-18 | HLA-C*12:03  HLA-C*03:03  HLA-B*46:01  HLA-B*51:01  HLA-B*53:01  HLA-C*06:02  HLA-C*07:01  HLA-A*68:02  HLA-B*38:01 | 0.7 |
