## Supplementary material for "Designing of a next generation multiepitope based vaccine (MEV) against SARS-COV-2: Immunoinformatics and *in silico* approaches": Supplementary Table 7.docx

Supplementary Table 7. List of predicted potential MHC Class II epitopes of SARS-COV-2 proteins.

| Protein | | Peptide (Position) | Alleles | Antigenicity |
| --- | --- | --- | --- | --- |
| E Protein | | FLLVTLAILTALRLC (26-40) | HLA-DRB1*11:04  HLA-DRB1*11:06  HLA-DRB1*13:11  HLA-DRB1*01:01  HLA-DRB1*01:02  HLA-DPA1*03:01/  DPB1*04:02  HLA-DRB1*07:01  HLA-DRB1*11:28  HLA-DRB1*13:05  HLA-DRB1*13:07 | 0.6311 |
|  |  | LLFLAFVVFLLVTLA (18-32) | HLA-DPA1*03:01/  DPB1*04:02  HLA-DPA1*01:03/  DPB1*02:01  HLA-DPA1*01/  DPB1*04:01  HLA-DPA1*02:01/  DPB1*01:01  HLA-DRB1*15:02  HLA-DRB1*04:23  HLA-DRB1*04:04  HLA-DRB1*04:08  HLA-DRB1*04:10  HLA-DQA1*05:01/  DQB1*02:01  HLA-DRB1*08:13  HLA-DRB1*07:03  HLA-DRB1*01:02  HLA-DRB1*04:05 | 0.8122 |
|  |  | VLLFLAFVVFLLVTL (17-31) | HLA-DPA1*03:01/  DPB1*04:02  HLA-DPA1*01:03/  DPB1*02:01  HLA-DPA1*01/  DPB1*04:01  HLA-DPA1*02:01/  DPB1*01:01  HLA-DRB1*15:02  HLA-DQA1*05:01/  DQB1*02:01  HLA-DRB1*15:01  HLA-DRB1*15:06  HLA-DRB1*07:03 | 0.6386 |
|  |  | LFLAFVVFLLVTLAI (19-33) | HLA-DPA1*03:01/  DPB1*04:02  HLA-DPA1*01:03/  DPB1*02:01  HLA-DPA1*01/  DPB1*04:01  HLA-DPA1*02:01/  DPB1*01:01  HLA-DRB1*15:02  HLA-DRB1*04:23  HLA-DRB1*11:04  HLA-DRB1*04:08  HLA-DRB1*13:11  HLA-DRB1*04:08  HLA-DRB1*04:10  HLA-DRB1*11:28  HLA-DRB1*13:05  HLA-DRB1*04:21  HLA-DRB1*08:13  HLA-DRB1*04:26  HLA-DRB1*07:03  HLA-DRB1*01:02  HLA-DRB1*04:04  HLA-DRB1*04:05 | 0.7471 |
| M Protein | | ESELVIGAVILRGHL (135-149) | HLA-DRB1*03:09  HLA-DRB1*11:07  HLA-DRB1*03:06  HLA-DRB1*03:07  HLA-DRB1*03:08  HLA-DQA1*01:02/  DQB1*06:02 | 0.5735 |
|  |  | PVTLACFVLAAVYRI (59-73) | HLA-DRB1*07:03  HLA-DRB1*11:20  HLA-DRB1*01:02  HLA-DRB1*07:01  HLA-DRB1*11:14  HLA-DRB1*13:23  HLA-DRB1*03:09  HLA-DRB1*13:07  HLA-DRB1*11:28  HLA-DRB1*13:05  HLA-DRB1*03:05  HLA-DRB1*04:08 | 0.8548 |
|  |  | LEQWNLVIGFLFLTW (17-31) | HLA-DPA1*01:03/  DPB1*02:01  HLA-DPA1*01/  DPB1*04:01  HLA-DRB5*01:05 | 1.0231 |
|  |  | RNRFLYIIKLIFLWL (42-56) | HLA-DRB1*11:28  HLA-DRB1*13:05  HLA-DRB1*13:21  HLA-DRB1*11:01  HLA-DPA1*02:01/  DPB1*01:01  HLA-DRB1*08:13  HLA-DRB1*08:01  HLA-DRB1*07:03  HLA-DPA1*01/  DPB1*04:01  HLA-DRB1*11:14  HLA-DRB1*13:23  HLA-DRB1*03:09  HLA-DRB4*01:01  HLA-DPA1*01:03/  DPB1*02:01  HLA-DRB1*08:17  HLA-DRB1*13:07  HLA-DRB1*15:01  HLA-DRB1*15:06 |  |
|  |  | SFRLFARTRSMWSFN (99-113) | HLA-DRB1*08:13  HLA-DRB1*11:14  HLA-DRB1*13:23  HLA-DRB1*15:02  HLA-DRB1*11:20  HLA-DRB1*11:01  HLA-DRB1*13:07  HLA-DRB1*15:06  HLA-DRB1*11:28  HLA-DRB1*13:05  HLA-DRB1*04:01  HLA-DRB1*04:26  HLA-DRB1*11:02  HLA-DRB1*11:21  HLA-DRB1*13:22  HLA-DRB1*03:05 | 0.7955 |
| Orf6 | | MFHLVDFQVTIAEIL (1-15) | HLA-DRB1*07:01  HLA-DRB1*07:03  HLA-DQA1*04:01  DQB1*04:02  HLA-DPA1*03:01  DPB1*04:02 | 1.0366 |
|  |  | FHLVDFQVTIAEILL (2-16) | HLA-DPA1*03:01/  DPB1*04:02  HLA-DRB1*07:03  HLA-DQA1*04:01/  DQB1*04:02  HLA-DRB1*07:01 | 1.1567 |
|  |  | DFQVTIAEILLIIMR (6-20) | HLA-DRB1*07:03  HLA-DPA1*03:01/  DPB1*04:02  HLA-DRB1*07:01 | 0.8778 |
|  |  | VDFQVTIAEILLIIM(5-19) | HLA-DPA1*03:01/  DPB1*04:02  HLA-DRB1*07:03  HLA-DRB1*07:01  HLA-DPA1*02:01/  DPB1*01:01 | 0.9754 |
| Orf7A | | DGVKHVYQLRARSVS (69-83) | HLA-DRB1*08:01  HLA-DRB1*08:13  HLA-DRB1*01:01  HLA-DRB1*08:04 | 0.7457 |
|  |  | HVYQLRARSVSPKLF (73-87) | HLA-DRB1*08:13  HLA-DRB1*08:01  HLA-DRB1*08:04 | 0.5654 |
|  |  | GVKHVYQLRARSVSP (70-84) | HLA-DRB1*08:13  HLA-DRB1*08:01  HLA-DRB1*01:01  HLA-DRB1*08:04 | 1.2761 |
|  |  | KHVYQLRARSVSPKL (72-86) | HLA-DRB1*08:13  HLA-DRB1*08:01  HLA-DRB1*01:01  HLA-DRB1*08:04 | 0.8381 |
|  |  | VKHVYQLRARSVSPK (71-85) | HLA-DRB1*08:13  HLA-DRB1*08:01  HLA-DRB1*01:01  HLA-DRB1*08:04 | 1.0865 |
|  |  | YQLRARSVSPKLFIR (75-89) | HLA-DRB1*08:13  HLA-DRB1*08:01  HLA-DRB1*08:04 | 0.7805 |
|  |  | AAIVFITLCFTLKRK (105-119) | HLA-DRB1*07:03  HLA-DPA1*02:01  DPB1*05:01 | 1.8170 |
| Orf8 | | CTQHQPYVVDDPCPI (25-39) | HLA-DRB3*01:01  HLA-DRB1*03:09  HLA-DRB1*03:05  HLA-DRB1*04:21 | 0.5165 |
|  |  | HQPYVVDDPCPIHFY (28-42) | HLA-DRB3*01:01  HLA-DRB1*03:09  HLA-DRB1*03:05  HLA-DRB1*04:21  HLA-DRB1*03:01 | 0.5587 |
|  |  | QHQPYVVDDPCPIHF (27-41) | HLA-DRB3*01:01  HLA-DRB1*03:09  HLA-DRB1*03:05  HLA-DRB1*04:21  HLA-DRB1*03:01 | 0.8637 |
|  |  | TQHQPYVVDDPCPIH (26-40) | HLA-DRB3*01:01  HLA-DRB1*03:09  HLA-DRB1*03:05  HLA-DRB1*04:21 | 0.6706 |
|  |  | PYVVDDPCPIHFYSK (30-44) | HLA-DRB1*03:09  HLA-DRB1*03:05  HLA-DRB3*01:01  HLA-DRB1*04:21 | 0.6766 |
|  |  | FLGIITTVAAFHQEC (6-20) | HLA-DRB1*04:08  HLA-DRB1*04:23  HLA-DRB1*04:10  HLA-DRB1*04:02  HLA-DRB1*04:26 | 0.6904 |
|  |  | VVDDPCPIHFYSKW (3-17) | HLA-DRB1*04:08  HLA-DRB1*15:02  HLA-DRB1*04:23  HLA-DRB1*04:10  HLA-DRB1*11:04  HLA-DRB1*11:06  HLA-DRB1*13:11  HLA-DRB1*04:05  HLA-DRB1*01:01  HLA-DRB1*04:02  HLA-DRB1*04:01  HLA-DRB1*04:26 | 0.5754 |
|  |  | LVFLGIITTVAAFHQ (4-18) | HLA-DRB1*04:08  HLA-DRB1*04:23  HLA-DRB1*04:10  HLA-DRB1*11:04  HLA-DRB1*11:06  HLA-DRB1*13:11  HLA-DRB1*08:02  HLA-DRB1*04:05  HLA-DRB1*04:02  HLA-DRB1*04:04  HLA-DRB1*04:01  HLA-DRB1*04:26 | 0.6791 |
|  |  | MKFLVFLGIITTVAA (1-15) | HLA-DRB1*04:08  HLA-DRB1*15:02  HLA-DRB1*04:23  HLA-DRB1*04:05  HLA-DRB1*04:04  HLA-DRB1*01:01  HLA-DRB1*04:10  HLA-DRB1*11:04  HLA-DRB1*11:06  HLA-DRB1*13:11  HLA-DRB1*04:02  HLA-DRB1*04:01  HLA-DRB1*04:26 | 0.5366 |
|  |  | LGIITTVAAFHQECS (7-21) | HLA-DRB1*04:23  HLA-DRB1*04:08  HLA-DRB1*04:10  HLA-DRB1*04:02  HLA-DRB1*04:26 | 0.7908 |
|  | | VFLGIITTVAAFHQE (5-19) | HLA-DRB1*04:08  HLA-DRB1*04:23  HLA-DRB1*04:10  HLA-DRB1*04:23  HLA-DRB1*04:08  HLA-DRB1*04:26  HLA-DRB1*04:01 | 0.6382 |
|  | | PKLGSLVVRCSFYED (93-107) | HLA-DRB1*08:06  HLA-DRB1*11:02  HLA-DRB1*11:21  HLA-DRB1*13:22  HLA-DRB1*13:01  HLA-DRB1*13:27  HLA-DRB1*13:28  HLA-DRB1*08:01  HLA-DRB1*13:04 | 0.7902 |
| Nc Protein | | FFGMSRIGMEVTPSG (314-328) | HLA-DRB1*11:28  HLA-DRB1*13:05  HLA-DRB1*13:21 | 0.9397 |
|  |  | FFGMSRIGMEVTPSG (314-328) | HLA-DRB1*11:28  HLA-DRB1*13:05  HLA-DRB1*13:21 | 0.9397 |
|  |  | KAYNVTQAFGRRGPE (266-280) | HLA-DRB5*01:05  HLA-DRB5*01:01 | 0.6104 |
|  |  | ALLLLDRLNQLESKM (220-234) | HLA-DRB1*11:04  HLA-DRB1*11:06  HLA-DRB1*13:11  HLA-DRB1*13:21  HLA-DRB1*13:07  HLA-DRB1*11:02  HLA-DRB1*11:21  HLA-DRB1*13:22  HLA-DRB1*13:04  HLA-DRB1*08:17  HLA-DPA1*03:01/  DPB1*04:02  HLA-DRB1*08:06  HLA-DRB1*11:28  HLA-DRB1*13:05  HLA-DRB1*08:04  HLA-DRB1*11:14  HLA-DRB1*13:23 | 0.5669 |
| Orf10 | CRMNSRNYIAQVDV (8-22) | | HLA-DRB1*15:02 | 0.6757 |
|  | FAFPFTIYSLLLCRM (7-21) | | HLA-DRB1*15:02  HLA-DPA1*01/  DPB1*04:01  HLA-DPA1*02:01/  DPB1*01:01  HLA-DPA1*03:01/  DPB1*04:02  HLA-DRB1*08:17  HLA-DPA1*01:03/  DPB1*02:01  HLA-DRB1*08:13 | 0.5718 |
|  | FPFTIYSLLLCRMNS (9-23) | | HLA-DRB1*15:02  HLA-DRB1*08:17  HLA-DPA1*01/  DPB1*04:01  HLA-DRB1*08:13 | 0.6560 |
|  | PFTIYSLLLCRMNSR (10-24) | | HLA-DRB1*15:02  HLA-DRB1*08:17  HLA-DRB1*08:13 | 0.9445 |
