## Supplementary material for "Designing of a next generation multiepitope based vaccine (MEV) against SARS-COV-2: Immunoinformatics and *in silico* approaches": Supplementary Table 8.docx

Supplementary Table 8. List of predicted potential IFN-γ inducing epitopes of SARS-COV-2 proteins.

| No. | Position | Epitope | Method | Result | Score |
| --- | --- | --- | --- | --- | --- |
| E Protein | | | | | |
| 1 | 36-51 | [LRLCAYCCNIVNVSL](http://crdd.osdd.net/raghava/ifnepitope/pep_design.php?sequence=LRLCAYCCNIVNVSL&method=hybrid&model=main) | MERCI | POSITIVE | 1 |
| 2 | 37-52 | [RLCAYCCNIVNVSLV](http://crdd.osdd.net/raghava/ifnepitope/pep_design.php?sequence=RLCAYCCNIVNVSLV&method=hybrid&model=main) | MERCI | POSITIVE | 1 |
| 3 | 38-53 | [LCAYCCNIVNVSLVK](http://crdd.osdd.net/raghava/ifnepitope/pep_design.php?sequence=LCAYCCNIVNVSLVK&method=hybrid&model=main) | MERCI | POSITIVE | 1 |
| 4 | 39-54 | [CAYCCNIVNVSLVKP](http://crdd.osdd.net/raghava/ifnepitope/pep_design.php?sequence=CAYCCNIVNVSLVKP&method=hybrid&model=main) | MERCI | POSITIVE | 1 |
| 5 | 40-55 | [AYCCNIVNVSLVKPS](http://crdd.osdd.net/raghava/ifnepitope/pep_design.php?sequence=AYCCNIVNVSLVKPS&method=hybrid&model=main) | MERCI | POSITIVE | 1 |
| 6 | 41-56 | [YCCNIVNVSLVKPSF](http://crdd.osdd.net/raghava/ifnepitope/pep_design.php?sequence=YCCNIVNVSLVKPSF&method=hybrid&model=main) | MERCI | POSITIVE | 1 |
| 7 | 42-57 | [CCNIVNVSLVKPSFY](http://crdd.osdd.net/raghava/ifnepitope/pep_design.php?sequence=CCNIVNVSLVKPSFY&method=hybrid&model=main) | MERCI | POSITIVE | 1 |
| 8 | 7-22 | [ETGTLIVNSVLLFLA](http://crdd.osdd.net/raghava/ifnepitope/pep_design.php?sequence=ETGTLIVNSVLLFLA&method=hybrid&model=main) | MERCI | NEGATIVE | 1 |
| 9 | 8-23 | [TGTLIVNSVLLFLAF](http://crdd.osdd.net/raghava/ifnepitope/pep_design.php?sequence=TGTLIVNSVLLFLAF&method=hybrid&model=main) | MERCI | NEGATIVE | 1 |
| 10 | 9-24 | [GTLIVNSVLLFLAFV](http://crdd.osdd.net/raghava/ifnepitope/pep_design.php?sequence=GTLIVNSVLLFLAFV&method=hybrid&model=main) | MERCI | NEGATIVE | 1 |
| 11 | 10-25 | [TLIVNSVLLFLAFVV](http://crdd.osdd.net/raghava/ifnepitope/pep_design.php?sequence=TLIVNSVLLFLAFVV&method=hybrid&model=main) | MERCI | NEGATIVE | 2 |
| 12 | 11-26 | [LIVNSVLLFLAFVVF](http://crdd.osdd.net/raghava/ifnepitope/pep_design.php?sequence=LIVNSVLLFLAFVVF&method=hybrid&model=main) | MERCI | NEGATIVE | 2 |
| 13 | 12-27 | [IVNSVLLFLAFVVFL](http://crdd.osdd.net/raghava/ifnepitope/pep_design.php?sequence=IVNSVLLFLAFVVFL&method=hybrid&model=main) | MERCI | NEGATIVE | 2 |
| 14 | 13-28 | [VNSVLLFLAFVVFLL](http://crdd.osdd.net/raghava/ifnepitope/pep_design.php?sequence=VNSVLLFLAFVVFLL&method=hybrid&model=main) | MERCI | NEGATIVE | 2 |
| 15 | 14-29 | [NSVLLFLAFVVFLLV](http://crdd.osdd.net/raghava/ifnepitope/pep_design.php?sequence=NSVLLFLAFVVFLLV&method=hybrid&model=main) | MERCI | NEGATIVE | 1 |
| 16 | 15-30 | [SVLLFLAFVVFLLVT](http://crdd.osdd.net/raghava/ifnepitope/pep_design.php?sequence=SVLLFLAFVVFLLVT&method=hybrid&model=main) | MERCI | NEGATIVE | 1 |
| 17 | 16-31 | [VLLFLAFVVFLLVTL](http://crdd.osdd.net/raghava/ifnepitope/pep_design.php?sequence=VLLFLAFVVFLLVTL&method=hybrid&model=main) | MERCI | NEGATIVE | 1 |
| 18 | 17-32 | [LLFLAFVVFLLVTLA](http://crdd.osdd.net/raghava/ifnepitope/pep_design.php?sequence=LLFLAFVVFLLVTLA&method=hybrid&model=main) | MERCI | NEGATIVE | 2 |
| 19 | 18-33 | [LFLAFVVFLLVTLAI](http://crdd.osdd.net/raghava/ifnepitope/pep_design.php?sequence=LFLAFVVFLLVTLAI&method=hybrid&model=main) | MERCI | NEGATIVE | 4 |
| 20 | 19-34 | [FLAFVVFLLVTLAIL](http://crdd.osdd.net/raghava/ifnepitope/pep_design.php?sequence=FLAFVVFLLVTLAIL&method=hybrid&model=main) | MERCI | NEGATIVE | 7 |
| 21 | 20-35 | [LAFVVFLLVTLAILT](http://crdd.osdd.net/raghava/ifnepitope/pep_design.php?sequence=LAFVVFLLVTLAILT&method=hybrid&model=main) | MERCI | NEGATIVE | 7 |
| 22 | 21-36 | [AFVVFLLVTLAILTA](http://crdd.osdd.net/raghava/ifnepitope/pep_design.php?sequence=AFVVFLLVTLAILTA&method=hybrid&model=main) | MERCI | NEGATIVE | 9 |
| 23 | 22-37 | [FVVFLLVTLAILTAL](http://crdd.osdd.net/raghava/ifnepitope/pep_design.php?sequence=FVVFLLVTLAILTAL&method=hybrid&model=main) | MERCI | NEGATIVE | 10 |
| 24 | 23-38 | [VVFLLVTLAILTALR](http://crdd.osdd.net/raghava/ifnepitope/pep_design.php?sequence=VVFLLVTLAILTALR&method=hybrid&model=main) | MERCI | NEGATIVE | 10 |
| 25 | 24-39 | [VFLLVTLAILTALRL](http://crdd.osdd.net/raghava/ifnepitope/pep_design.php?sequence=VFLLVTLAILTALRL&method=hybrid&model=main) | MERCI | NEGATIVE | 9 |
| 26 | 25-40 | [FLLVTLAILTALRLC](http://crdd.osdd.net/raghava/ifnepitope/pep_design.php?sequence=FLLVTLAILTALRLC&method=hybrid&model=main) | MERCI | NEGATIVE | 9 |
| 27 | 26-41 | [LLVTLAILTALRLCA](http://crdd.osdd.net/raghava/ifnepitope/pep_design.php?sequence=LLVTLAILTALRLCA&method=hybrid&model=main) | MERCI | NEGATIVE | 5 |
| 28 | 27-42 | [LVTLAILTALRLCAY](http://crdd.osdd.net/raghava/ifnepitope/pep_design.php?sequence=LVTLAILTALRLCAY&method=hybrid&model=main) | MERCI | NEGATIVE | 3 |
| 29 | 28-43 | [VTLAILTALRLCAYC](http://crdd.osdd.net/raghava/ifnepitope/pep_design.php?sequence=VTLAILTALRLCAYC&method=hybrid&model=main) | MERCI | NEGATIVE | 2 |
| 30 | 29-44 | [TLAILTALRLCAYCC](http://crdd.osdd.net/raghava/ifnepitope/pep_design.php?sequence=TLAILTALRLCAYCC&method=hybrid&model=main) | MERCI | NEGATIVE | 1 |
| 31 | 0-15 | [MYSFVSEETGTLIVN](http://crdd.osdd.net/raghava/ifnepitope/pep_design.php?sequence=MYSFVSEETGTLIVN&method=hybrid&model=main) | SVM | NEGATIVE | -1.3119431 |
| 32 | 1-16 | [YSFVSEETGTLIVNS](http://crdd.osdd.net/raghava/ifnepitope/pep_design.php?sequence=YSFVSEETGTLIVNS&method=hybrid&model=main) | SVM | NEGATIVE | -1.3881082 |
| 33 | 2-17 | [SFVSEETGTLIVNSV](http://crdd.osdd.net/raghava/ifnepitope/pep_design.php?sequence=SFVSEETGTLIVNSV&method=hybrid&model=main) | SVM | NEGATIVE | -1.3238486 |
| 34 | 3-18 | [FVSEETGTLIVNSVL](http://crdd.osdd.net/raghava/ifnepitope/pep_design.php?sequence=FVSEETGTLIVNSVL&method=hybrid&model=main) | SVM | NEGATIVE | -1.1118721 |
| 35 | 4-19 | [VSEETGTLIVNSVLL](http://crdd.osdd.net/raghava/ifnepitope/pep_design.php?sequence=VSEETGTLIVNSVLL&method=hybrid&model=main) | SVM | NEGATIVE | -0.98029675 |
| 36 | 5-20 | [SEETGTLIVNSVLLF](http://crdd.osdd.net/raghava/ifnepitope/pep_design.php?sequence=SEETGTLIVNSVLLF&method=hybrid&model=main) | SVM | NEGATIVE | -0.92370224 |
| 37 | 6-21 | [EETGTLIVNSVLLFL](http://crdd.osdd.net/raghava/ifnepitope/pep_design.php?sequence=EETGTLIVNSVLLFL&method=hybrid&model=main) | SVM | NEGATIVE | -0.72918894 |
| 38 | 30-45 | [LAILTALRLCAYCCN](http://crdd.osdd.net/raghava/ifnepitope/pep_design.php?sequence=LAILTALRLCAYCCN&method=hybrid&model=main) | SVM | NEGATIVE | -0.25662442 |
| 39 | 31-46 | [AILTALRLCAYCCNI](http://crdd.osdd.net/raghava/ifnepitope/pep_design.php?sequence=AILTALRLCAYCCNI&method=hybrid&model=main) | SVM | NEGATIVE | -0.041707531 |
| 40 | 32-47 | [ILTALRLCAYCCNIV](http://crdd.osdd.net/raghava/ifnepitope/pep_design.php?sequence=ILTALRLCAYCCNIV&method=hybrid&model=main) | SVM | NEGATIVE | -0.067419289 |
| 41 | 33-48 | [LTALRLCAYCCNIVN](http://crdd.osdd.net/raghava/ifnepitope/pep_design.php?sequence=LTALRLCAYCCNIVN&method=hybrid&model=main) | SVM | NEGATIVE | -0.11543448 |
| 42 | 34-49 | [TALRLCAYCCNIVNV](http://crdd.osdd.net/raghava/ifnepitope/pep_design.php?sequence=TALRLCAYCCNIVNV&method=hybrid&model=main) | SVM | POSITIVE | 0.057072828 |
| 43 | 35-50 | [ALRLCAYCCNIVNVS](http://crdd.osdd.net/raghava/ifnepitope/pep_design.php?sequence=ALRLCAYCCNIVNVS&method=hybrid&model=main) | SVM | NEGATIVE | -0.050255252 |
| 44 | 43-58 | [CNIVNVSLVKPSFYV](http://crdd.osdd.net/raghava/ifnepitope/pep_design.php?sequence=CNIVNVSLVKPSFYV&method=hybrid&model=main) | SVM | NEGATIVE | -0.68425525 |
| 45 | 44-59 | [NIVNVSLVKPSFYVY](http://crdd.osdd.net/raghava/ifnepitope/pep_design.php?sequence=NIVNVSLVKPSFYVY&method=hybrid&model=main) | SVM | NEGATIVE | -0.53035908 |
| 46 | 45-60 | [IVNVSLVKPSFYVYS](http://crdd.osdd.net/raghava/ifnepitope/pep_design.php?sequence=IVNVSLVKPSFYVYS&method=hybrid&model=main) | SVM | NEGATIVE | -0.70036781 |
| 47 | 46-61 | [VNVSLVKPSFYVYSR](http://crdd.osdd.net/raghava/ifnepitope/pep_design.php?sequence=VNVSLVKPSFYVYSR&method=hybrid&model=main) | SVM | NEGATIVE | -0.55006124 |
| 48 | 47-62 | [NVSLVKPSFYVYSRV](http://crdd.osdd.net/raghava/ifnepitope/pep_design.php?sequence=NVSLVKPSFYVYSRV&method=hybrid&model=main) | SVM | NEGATIVE | -0.50087928 |
| 49 | 48-63 | [VSLVKPSFYVYSRVK](http://crdd.osdd.net/raghava/ifnepitope/pep_design.php?sequence=VSLVKPSFYVYSRVK&method=hybrid&model=main) | SVM | NEGATIVE | -0.064637075 |
| 50 | 49-64 | [SLVKPSFYVYSRVKN](http://crdd.osdd.net/raghava/ifnepitope/pep_design.php?sequence=SLVKPSFYVYSRVKN&method=hybrid&model=main) | SVM | NEGATIVE | -0.28654698 |
| 51 | 50-65 | [LVKPSFYVYSRVKNL](http://crdd.osdd.net/raghava/ifnepitope/pep_design.php?sequence=LVKPSFYVYSRVKNL&method=hybrid&model=main) | SVM | NEGATIVE | -0.52227889 |
| 52 | 51-66 | [VKPSFYVYSRVKNLN](http://crdd.osdd.net/raghava/ifnepitope/pep_design.php?sequence=VKPSFYVYSRVKNLN&method=hybrid&model=main) | SVM | NEGATIVE | -0.4094391 |
| 53 | 52-67 | [KPSFYVYSRVKNLNS](http://crdd.osdd.net/raghava/ifnepitope/pep_design.php?sequence=KPSFYVYSRVKNLNS&method=hybrid&model=main) | SVM | NEGATIVE | -0.77512283 |
| 54 | 53-68 | [PSFYVYSRVKNLNSS](http://crdd.osdd.net/raghava/ifnepitope/pep_design.php?sequence=PSFYVYSRVKNLNSS&method=hybrid&model=main) | SVM | NEGATIVE | -0.63066447 |
| 55 | 54-69 | [SFYVYSRVKNLNSSR](http://crdd.osdd.net/raghava/ifnepitope/pep_design.php?sequence=SFYVYSRVKNLNSSR&method=hybrid&model=main) | SVM | NEGATIVE | -0.69311639 |
| 56 | 55-70 | [FYVYSRVKNLNSSRV](http://crdd.osdd.net/raghava/ifnepitope/pep_design.php?sequence=FYVYSRVKNLNSSRV&method=hybrid&model=main) | SVM | NEGATIVE | -0.33015493 |
| 57 | 56-71 | [YVYSRVKNLNSSRVP](http://crdd.osdd.net/raghava/ifnepitope/pep_design.php?sequence=YVYSRVKNLNSSRVP&method=hybrid&model=main) | SVM | NEGATIVE | -0.22463037 |
| 58 | 57-72 | [VYSRVKNLNSSRVPD](http://crdd.osdd.net/raghava/ifnepitope/pep_design.php?sequence=VYSRVKNLNSSRVPD&method=hybrid&model=main) | SVM | NEGATIVE | -0.35609024 |
| 59 | 58-73 | [YSRVKNLNSSRVPDL](http://crdd.osdd.net/raghava/ifnepitope/pep_design.php?sequence=YSRVKNLNSSRVPDL&method=hybrid&model=main) | SVM | NEGATIVE | -0.45329667 |
| 60 | 59-74 | [SRVKNLNSSRVPDLL](http://crdd.osdd.net/raghava/ifnepitope/pep_design.php?sequence=SRVKNLNSSRVPDLL&method=hybrid&model=main) | SVM | NEGATIVE | -0.33181769 |
| 61 | 60-75 | [RVKNLNSSRVPDLLV](http://crdd.osdd.net/raghava/ifnepitope/pep_design.php?sequence=RVKNLNSSRVPDLLV&method=hybrid&model=main) | SVM | NEGATIVE | -0.44780857 |
| 62 | 61-75 | [VKNLNSSRVPDLLV](http://crdd.osdd.net/raghava/ifnepitope/pep_design.php?sequence=VKNLNSSRVPDLLV&method=hybrid&model=main) | SVM | NEGATIVE | -0.99405478 |
| 63 | 62-75 | [KNLNSSRVPDLLV](http://crdd.osdd.net/raghava/ifnepitope/pep_design.php?sequence=KNLNSSRVPDLLV&method=hybrid&model=main) | SVM | NEGATIVE | -0.99531368 |
| 64 | 63-75 | [NLNSSRVPDLLV](http://crdd.osdd.net/raghava/ifnepitope/pep_design.php?sequence=NLNSSRVPDLLV&method=hybrid&model=main) | SVM | NEGATIVE | -1.0513826 |
| 65 | 64-75 | [LNSSRVPDLLV](http://crdd.osdd.net/raghava/ifnepitope/pep_design.php?sequence=LNSSRVPDLLV&method=hybrid&model=main) | SVM | NEGATIVE | -0.92410842 |
| 66 | 65-75 | [NSSRVPDLLV](http://crdd.osdd.net/raghava/ifnepitope/pep_design.php?sequence=NSSRVPDLLV&method=hybrid&model=main) | SVM | NEGATIVE | -0.7408137 |
| 67 | 66-75 | [SSRVPDLLV](http://crdd.osdd.net/raghava/ifnepitope/pep_design.php?sequence=SSRVPDLLV&method=hybrid&model=main) | SVM | NEGATIVE | -0.54102816 |
| M Protein | | | | | |
| 1 | 165-180 | [KEITVATSRTLSYYK](http://crdd.osdd.net/raghava/ifnepitope/pep_design.php?sequence=KEITVATSRTLSYYK&method=hybrid&model=main) | MERCI | POSITIVE | 1 |
| 2 | 166-181 | [EITVATSRTLSYYKL](http://crdd.osdd.net/raghava/ifnepitope/pep_design.php?sequence=EITVATSRTLSYYKL&method=hybrid&model=main) | MERCI | POSITIVE | 1 |
| 3 | 167-182 | [ITVATSRTLSYYKLG](http://crdd.osdd.net/raghava/ifnepitope/pep_design.php?sequence=ITVATSRTLSYYKLG&method=hybrid&model=main) | MERCI | POSITIVE | 1 |
| 4 | 168-183 | [TVATSRTLSYYKLGA](http://crdd.osdd.net/raghava/ifnepitope/pep_design.php?sequence=TVATSRTLSYYKLGA&method=hybrid&model=main) | MERCI | POSITIVE | 1 |
| 5 | 169-184 | [VATSRTLSYYKLGAS](http://crdd.osdd.net/raghava/ifnepitope/pep_design.php?sequence=VATSRTLSYYKLGAS&method=hybrid&model=main) | MERCI | POSITIVE | 1 |
| 6 | 170-185 | [ATSRTLSYYKLGASQ](http://crdd.osdd.net/raghava/ifnepitope/pep_design.php?sequence=ATSRTLSYYKLGASQ&method=hybrid&model=main) | MERCI | POSITIVE | 1 |
| 7 | 171-186 | [TSRTLSYYKLGASQR](http://crdd.osdd.net/raghava/ifnepitope/pep_design.php?sequence=TSRTLSYYKLGASQR&method=hybrid&model=main) | MERCI | POSITIVE | 1 |
| 8 | 172-187 | [SRTLSYYKLGASQRV](http://crdd.osdd.net/raghava/ifnepitope/pep_design.php?sequence=SRTLSYYKLGASQRV&method=hybrid&model=main) | MERCI | POSITIVE | 1 |
| 9 | 9-24 | [VEELKKLLEQWNLVI](http://crdd.osdd.net/raghava/ifnepitope/pep_design.php?sequence=VEELKKLLEQWNLVI&method=hybrid&model=main) | MERCI | NEGATIVE | 1 |
| 10 | 10-25 | [EELKKLLEQWNLVIG](http://crdd.osdd.net/raghava/ifnepitope/pep_design.php?sequence=EELKKLLEQWNLVIG&method=hybrid&model=main) | MERCI | NEGATIVE | 1 |
| 11 | 11-26 | [ELKKLLEQWNLVIGF](http://crdd.osdd.net/raghava/ifnepitope/pep_design.php?sequence=ELKKLLEQWNLVIGF&method=hybrid&model=main) | MERCI | NEGATIVE | 1 |
| 12 | 12-27 | [LKKLLEQWNLVIGFL](http://crdd.osdd.net/raghava/ifnepitope/pep_design.php?sequence=LKKLLEQWNLVIGFL&method=hybrid&model=main) | MERCI | NEGATIVE | 4 |
| 13 | 13-28 | [KKLLEQWNLVIGFLF](http://crdd.osdd.net/raghava/ifnepitope/pep_design.php?sequence=KKLLEQWNLVIGFLF&method=hybrid&model=main) | MERCI | NEGATIVE | 4 |
| 14 | 14-29 | [KLLEQWNLVIGFLFL](http://crdd.osdd.net/raghava/ifnepitope/pep_design.php?sequence=KLLEQWNLVIGFLFL&method=hybrid&model=main) | MERCI | NEGATIVE | 4 |
| 15 | 15-30 | [LLEQWNLVIGFLFLT](http://crdd.osdd.net/raghava/ifnepitope/pep_design.php?sequence=LLEQWNLVIGFLFLT&method=hybrid&model=main) | MERCI | NEGATIVE | 5 |
| 16 | 16-31 | [LEQWNLVIGFLFLTW](http://crdd.osdd.net/raghava/ifnepitope/pep_design.php?sequence=LEQWNLVIGFLFLTW&method=hybrid&model=main) | MERCI | NEGATIVE | 4 |
| 17 | 17-32 | [EQWNLVIGFLFLTWI](http://crdd.osdd.net/raghava/ifnepitope/pep_design.php?sequence=EQWNLVIGFLFLTWI&method=hybrid&model=main) | MERCI | NEGATIVE | 6 |
| 18 | 18-33 | [QWNLVIGFLFLTWIC](http://crdd.osdd.net/raghava/ifnepitope/pep_design.php?sequence=QWNLVIGFLFLTWIC&method=hybrid&model=main) | MERCI | NEGATIVE | 6 |
| 19 | 19-34 | [WNLVIGFLFLTWICL](http://crdd.osdd.net/raghava/ifnepitope/pep_design.php?sequence=WNLVIGFLFLTWICL&method=hybrid&model=main) | MERCI | NEGATIVE | 7 |
| 20 | 20-35 | [NLVIGFLFLTWICLL](http://crdd.osdd.net/raghava/ifnepitope/pep_design.php?sequence=NLVIGFLFLTWICLL&method=hybrid&model=main) | MERCI | NEGATIVE | 14 |
| 21 | 21-36 | [LVIGFLFLTWICLLQ](http://crdd.osdd.net/raghava/ifnepitope/pep_design.php?sequence=LVIGFLFLTWICLLQ&method=hybrid&model=main) | MERCI | NEGATIVE | 12 |
| 22 | 22-37 | [VIGFLFLTWICLLQF](http://crdd.osdd.net/raghava/ifnepitope/pep_design.php?sequence=VIGFLFLTWICLLQF&method=hybrid&model=main) | MERCI | NEGATIVE | 12 |
| 23 | 23-38 | [IGFLFLTWICLLQFA](http://crdd.osdd.net/raghava/ifnepitope/pep_design.php?sequence=IGFLFLTWICLLQFA&method=hybrid&model=main) | MERCI | NEGATIVE | 11 |
| 24 | 24-39 | [GFLFLTWICLLQFAY](http://crdd.osdd.net/raghava/ifnepitope/pep_design.php?sequence=GFLFLTWICLLQFAY&method=hybrid&model=main) | MERCI | NEGATIVE | 11 |
| 25 | 25-40 | [FLFLTWICLLQFAYA](http://crdd.osdd.net/raghava/ifnepitope/pep_design.php?sequence=FLFLTWICLLQFAYA&method=hybrid&model=main) | MERCI | NEGATIVE | 11 |
| 26 | 26-41 | [LFLTWICLLQFAYAN](http://crdd.osdd.net/raghava/ifnepitope/pep_design.php?sequence=LFLTWICLLQFAYAN&method=hybrid&model=main) | MERCI | NEGATIVE | 9 |
| 27 | 27-42 | [FLTWICLLQFAYANR](http://crdd.osdd.net/raghava/ifnepitope/pep_design.php?sequence=FLTWICLLQFAYANR&method=hybrid&model=main) | MERCI | NEGATIVE | 9 |
| 28 | 28-43 | [LTWICLLQFAYANRN](http://crdd.osdd.net/raghava/ifnepitope/pep_design.php?sequence=LTWICLLQFAYANRN&method=hybrid&model=main) | MERCI | NEGATIVE | 9 |
| 29 | 29-44 | [TWICLLQFAYANRNR](http://crdd.osdd.net/raghava/ifnepitope/pep_design.php?sequence=TWICLLQFAYANRNR&method=hybrid&model=main) | MERCI | NEGATIVE | 7 |
| 30 | 30-45 | [WICLLQFAYANRNRF](http://crdd.osdd.net/raghava/ifnepitope/pep_design.php?sequence=WICLLQFAYANRNRF&method=hybrid&model=main) | MERCI | NEGATIVE | 4 |
| 31 | 31-46 | [ICLLQFAYANRNRFL](http://crdd.osdd.net/raghava/ifnepitope/pep_design.php?sequence=ICLLQFAYANRNRFL&method=hybrid&model=main) | MERCI | NEGATIVE | 1 |
| 32 | 36-51 | [FAYANRNRFLYIIKL](http://crdd.osdd.net/raghava/ifnepitope/pep_design.php?sequence=FAYANRNRFLYIIKL&method=hybrid&model=main) | MERCI | NEGATIVE | 2 |
| 33 | 37-52 | [AYANRNRFLYIIKLI](http://crdd.osdd.net/raghava/ifnepitope/pep_design.php?sequence=AYANRNRFLYIIKLI&method=hybrid&model=main) | MERCI | NEGATIVE | 5 |
| 34 | 38-53 | [YANRNRFLYIIKLIF](http://crdd.osdd.net/raghava/ifnepitope/pep_design.php?sequence=YANRNRFLYIIKLIF&method=hybrid&model=main) | MERCI | NEGATIVE | 12 |
| 35 | 39-54 | [ANRNRFLYIIKLIFL](http://crdd.osdd.net/raghava/ifnepitope/pep_design.php?sequence=ANRNRFLYIIKLIFL&method=hybrid&model=main) | MERCI | NEGATIVE | 16 |
| 36 | 40-55 | [NRNRFLYIIKLIFLW](http://crdd.osdd.net/raghava/ifnepitope/pep_design.php?sequence=NRNRFLYIIKLIFLW&method=hybrid&model=main) | MERCI | NEGATIVE | 22 |
| 37 | 41-56 | [RNRFLYIIKLIFLWL](http://crdd.osdd.net/raghava/ifnepitope/pep_design.php?sequence=RNRFLYIIKLIFLWL&method=hybrid&model=main) | MERCI | NEGATIVE | 28 |
| 38 | 42-57 | [NRFLYIIKLIFLWLL](http://crdd.osdd.net/raghava/ifnepitope/pep_design.php?sequence=NRFLYIIKLIFLWLL&method=hybrid&model=main) | MERCI | NEGATIVE | 31 |
| 39 | 43-58 | [RFLYIIKLIFLWLLW](http://crdd.osdd.net/raghava/ifnepitope/pep_design.php?sequence=RFLYIIKLIFLWLLW&method=hybrid&model=main) | MERCI | NEGATIVE | 31 |
| 40 | 44-59 | [FLYIIKLIFLWLLWP](http://crdd.osdd.net/raghava/ifnepitope/pep_design.php?sequence=FLYIIKLIFLWLLWP&method=hybrid&model=main) | MERCI | NEGATIVE | 30 |
| 41 | 45-60 | [LYIIKLIFLWLLWPV](http://crdd.osdd.net/raghava/ifnepitope/pep_design.php?sequence=LYIIKLIFLWLLWPV&method=hybrid&model=main) | MERCI | NEGATIVE | 27 |
| 42 | 46-61 | [YIIKLIFLWLLWPVT](http://crdd.osdd.net/raghava/ifnepitope/pep_design.php?sequence=YIIKLIFLWLLWPVT&method=hybrid&model=main) | MERCI | NEGATIVE | 19 |
| 43 | 47-62 | [IIKLIFLWLLWPVTL](http://crdd.osdd.net/raghava/ifnepitope/pep_design.php?sequence=IIKLIFLWLLWPVTL&method=hybrid&model=main) | MERCI | NEGATIVE | 15 |
| 44 | 48-63 | [IKLIFLWLLWPVTLA](http://crdd.osdd.net/raghava/ifnepitope/pep_design.php?sequence=IKLIFLWLLWPVTLA&method=hybrid&model=main) | MERCI | NEGATIVE | 5 |
| 45 | 49-64 | [KLIFLWLLWPVTLAC](http://crdd.osdd.net/raghava/ifnepitope/pep_design.php?sequence=KLIFLWLLWPVTLAC&method=hybrid&model=main) | MERCI | NEGATIVE | 2 |
| 46 | 50-65 | [LIFLWLLWPVTLACF](http://crdd.osdd.net/raghava/ifnepitope/pep_design.php?sequence=LIFLWLLWPVTLACF&method=hybrid&model=main) | MERCI | NEGATIVE | 1 |
| 47 | 51-66 | [IFLWLLWPVTLACFV](http://crdd.osdd.net/raghava/ifnepitope/pep_design.php?sequence=IFLWLLWPVTLACFV&method=hybrid&model=main) | MERCI | NEGATIVE | 1 |
| 48 | 67-82 | [AAVYRINWITGGIAI](http://crdd.osdd.net/raghava/ifnepitope/pep_design.php?sequence=AAVYRINWITGGIAI&method=hybrid&model=main) | MERCI | NEGATIVE | 2 |
| 49 | 68-83 | [AVYRINWITGGIAIA](http://crdd.osdd.net/raghava/ifnepitope/pep_design.php?sequence=AVYRINWITGGIAIA&method=hybrid&model=main) | MERCI | NEGATIVE | 4 |
| 50 | 69-84 | [VYRINWITGGIAIAM](http://crdd.osdd.net/raghava/ifnepitope/pep_design.php?sequence=VYRINWITGGIAIAM&method=hybrid&model=main) | MERCI | NEGATIVE | 4 |
| 51 | 70-85 | [YRINWITGGIAIAMA](http://crdd.osdd.net/raghava/ifnepitope/pep_design.php?sequence=YRINWITGGIAIAMA&method=hybrid&model=main) | MERCI | NEGATIVE | 5 |
| 52 | 71-86 | [RINWITGGIAIAMAC](http://crdd.osdd.net/raghava/ifnepitope/pep_design.php?sequence=RINWITGGIAIAMAC&method=hybrid&model=main) | MERCI | NEGATIVE | 5 |
| 53 | 72-87 | [INWITGGIAIAMACL](http://crdd.osdd.net/raghava/ifnepitope/pep_design.php?sequence=INWITGGIAIAMACL&method=hybrid&model=main) | MERCI | NEGATIVE | 10 |
| 54 | 73-88 | [NWITGGIAIAMACLV](http://crdd.osdd.net/raghava/ifnepitope/pep_design.php?sequence=NWITGGIAIAMACLV&method=hybrid&model=main) | MERCI | NEGATIVE | 6 |
| 55 | 74-89 | [WITGGIAIAMACLVG](http://crdd.osdd.net/raghava/ifnepitope/pep_design.php?sequence=WITGGIAIAMACLVG&method=hybrid&model=main) | MERCI | NEGATIVE | 6 |
| 56 | 75-90 | [ITGGIAIAMACLVGL](http://crdd.osdd.net/raghava/ifnepitope/pep_design.php?sequence=ITGGIAIAMACLVGL&method=hybrid&model=main) | MERCI | NEGATIVE | 6 |
| 57 | 76-91 | [TGGIAIAMACLVGLM](http://crdd.osdd.net/raghava/ifnepitope/pep_design.php?sequence=TGGIAIAMACLVGLM&method=hybrid&model=main) | MERCI | NEGATIVE | 1 |
| 58 | 77-92 | [GGIAIAMACLVGLMW](http://crdd.osdd.net/raghava/ifnepitope/pep_design.php?sequence=GGIAIAMACLVGLMW&method=hybrid&model=main) | MERCI | NEGATIVE | 4 |
| 59 | 78-93 | [GIAIAMACLVGLMWL](http://crdd.osdd.net/raghava/ifnepitope/pep_design.php?sequence=GIAIAMACLVGLMWL&method=hybrid&model=main) | MERCI | NEGATIVE | 4 |
| 60 | 79-94 | [IAIAMACLVGLMWLS](http://crdd.osdd.net/raghava/ifnepitope/pep_design.php?sequence=IAIAMACLVGLMWLS&method=hybrid&model=main) | MERCI | NEGATIVE | 3 |
| 61 | 123-138 | [LHGTILTRPLLESEL](http://crdd.osdd.net/raghava/ifnepitope/pep_design.php?sequence=LHGTILTRPLLESEL&method=hybrid&model=main) | MERCI | NEGATIVE | 1 |
| 62 | 125-140 | [GTILTRPLLESELVI](http://crdd.osdd.net/raghava/ifnepitope/pep_design.php?sequence=GTILTRPLLESELVI&method=hybrid&model=main) | MERCI | NEGATIVE | 1 |
| 63 | 126-141 | [TILTRPLLESELVIG](http://crdd.osdd.net/raghava/ifnepitope/pep_design.php?sequence=TILTRPLLESELVIG&method=hybrid&model=main) | MERCI | NEGATIVE | 1 |
| 64 | 127-142 | [ILTRPLLESELVIGA](http://crdd.osdd.net/raghava/ifnepitope/pep_design.php?sequence=ILTRPLLESELVIGA&method=hybrid&model=main) | MERCI | NEGATIVE | 2 |
| 65 | 128-143 | [LTRPLLESELVIGAV](http://crdd.osdd.net/raghava/ifnepitope/pep_design.php?sequence=LTRPLLESELVIGAV&method=hybrid&model=main) | MERCI | NEGATIVE | 2 |
| 66 | 129-144 | [TRPLLESELVIGAVI](http://crdd.osdd.net/raghava/ifnepitope/pep_design.php?sequence=TRPLLESELVIGAVI&method=hybrid&model=main) | MERCI | NEGATIVE | 5 |
| 67 | 130-145 | [RPLLESELVIGAVIL](http://crdd.osdd.net/raghava/ifnepitope/pep_design.php?sequence=RPLLESELVIGAVIL&method=hybrid&model=main) | MERCI | NEGATIVE | 10 |
| 68 | 131-146 | [PLLESELVIGAVILR](http://crdd.osdd.net/raghava/ifnepitope/pep_design.php?sequence=PLLESELVIGAVILR&method=hybrid&model=main) | MERCI | NEGATIVE | 10 |
| 69 | 132-147 | [LLESELVIGAVILRG](http://crdd.osdd.net/raghava/ifnepitope/pep_design.php?sequence=LLESELVIGAVILRG&method=hybrid&model=main) | MERCI | NEGATIVE | 14 |
| 70 | 133-148 | [LESELVIGAVILRGH](http://crdd.osdd.net/raghava/ifnepitope/pep_design.php?sequence=LESELVIGAVILRGH&method=hybrid&model=main) | MERCI | NEGATIVE | 12 |
| 71 | 134-149 | [ESELVIGAVILRGHL](http://crdd.osdd.net/raghava/ifnepitope/pep_design.php?sequence=ESELVIGAVILRGHL&method=hybrid&model=main) | MERCI | NEGATIVE | 16 |
| 72 | 135-150 | [SELVIGAVILRGHLR](http://crdd.osdd.net/raghava/ifnepitope/pep_design.php?sequence=SELVIGAVILRGHLR&method=hybrid&model=main) | MERCI | NEGATIVE | 16 |
| 73 | 136-151 | [ELVIGAVILRGHLRI](http://crdd.osdd.net/raghava/ifnepitope/pep_design.php?sequence=ELVIGAVILRGHLRI&method=hybrid&model=main) | MERCI | NEGATIVE | 20 |
| 74 | 137-152 | [LVIGAVILRGHLRIA](http://crdd.osdd.net/raghava/ifnepitope/pep_design.php?sequence=LVIGAVILRGHLRIA&method=hybrid&model=main) | MERCI | NEGATIVE | 19 |
| 75 | 138-153 | [VIGAVILRGHLRIAG](http://crdd.osdd.net/raghava/ifnepitope/pep_design.php?sequence=VIGAVILRGHLRIAG&method=hybrid&model=main) | MERCI | NEGATIVE | 13 |
| 76 | 139-154 | [IGAVILRGHLRIAGH](http://crdd.osdd.net/raghava/ifnepitope/pep_design.php?sequence=IGAVILRGHLRIAGH&method=hybrid&model=main) | MERCI | NEGATIVE | 4 |
| 77 | 148-163 | [LRIAGHHLGRCDIKD](http://crdd.osdd.net/raghava/ifnepitope/pep_design.php?sequence=LRIAGHHLGRCDIKD&method=hybrid&model=main) | MERCI | NEGATIVE | 1 |
| 78 | 149-164 | [RIAGHHLGRCDIKDL](http://crdd.osdd.net/raghava/ifnepitope/pep_design.php?sequence=RIAGHHLGRCDIKDL&method=hybrid&model=main) | MERCI | NEGATIVE | 1 |
| 79 | 150-165 | [IAGHHLGRCDIKDLP](http://crdd.osdd.net/raghava/ifnepitope/pep_design.php?sequence=IAGHHLGRCDIKDLP&method=hybrid&model=main) | MERCI | NEGATIVE | 1 |
| 80 | 151-166 | [AGHHLGRCDIKDLPK](http://crdd.osdd.net/raghava/ifnepitope/pep_design.php?sequence=AGHHLGRCDIKDLPK&method=hybrid&model=main) | MERCI | NEGATIVE | 1 |
| 81 | 152-167 | [GHHLGRCDIKDLPKE](http://crdd.osdd.net/raghava/ifnepitope/pep_design.php?sequence=GHHLGRCDIKDLPKE&method=hybrid&model=main) | MERCI | NEGATIVE | 1 |
| 82 | 153-168 | [HHLGRCDIKDLPKEI](http://crdd.osdd.net/raghava/ifnepitope/pep_design.php?sequence=HHLGRCDIKDLPKEI&method=hybrid&model=main) | MERCI | NEGATIVE | 1 |
| 83 | 154-169 | [HLGRCDIKDLPKEIT](http://crdd.osdd.net/raghava/ifnepitope/pep_design.php?sequence=HLGRCDIKDLPKEIT&method=hybrid&model=main) | MERCI | NEGATIVE | 1 |
| 84 | 205-220 | [LNTDHSSSSDNIALL](http://crdd.osdd.net/raghava/ifnepitope/pep_design.php?sequence=LNTDHSSSSDNIALL&method=hybrid&model=main) | MERCI | NEGATIVE | 3 |
| 85 | 206-221 | [NTDHSSSSDNIALLV](http://crdd.osdd.net/raghava/ifnepitope/pep_design.php?sequence=NTDHSSSSDNIALLV&method=hybrid&model=main) | MERCI | NEGATIVE | 4 |
| 86 | 207-222 | [TDHSSSSDNIALLVQ](http://crdd.osdd.net/raghava/ifnepitope/pep_design.php?sequence=TDHSSSSDNIALLVQ&method=hybrid&model=main) | MERCI | NEGATIVE | 4 |
| 87 | 208-222 | [DHSSSSDNIALLVQ](http://crdd.osdd.net/raghava/ifnepitope/pep_design.php?sequence=DHSSSSDNIALLVQ&method=hybrid&model=main) | MERCI | NEGATIVE | 3 |
| 88 | 209-222 | [HSSSSDNIALLVQ](http://crdd.osdd.net/raghava/ifnepitope/pep_design.php?sequence=HSSSSDNIALLVQ&method=hybrid&model=main) | MERCI | NEGATIVE | 2 |
| 89 | 210-222 | [SSSSDNIALLVQ](http://crdd.osdd.net/raghava/ifnepitope/pep_design.php?sequence=SSSSDNIALLVQ&method=hybrid&model=main) | MERCI | NEGATIVE | 2 |
| 90 | 211-222 | [SSSDNIALLVQ](http://crdd.osdd.net/raghava/ifnepitope/pep_design.php?sequence=SSSDNIALLVQ&method=hybrid&model=main) | MERCI | NEGATIVE | 2 |
| 91 | 212-222 | [SSDNIALLVQ](http://crdd.osdd.net/raghava/ifnepitope/pep_design.php?sequence=SSDNIALLVQ&method=hybrid&model=main) | MERCI | NEGATIVE | 2 |
| 92 | 213-222 | [SDNIALLVQ](http://crdd.osdd.net/raghava/ifnepitope/pep_design.php?sequence=SDNIALLVQ&method=hybrid&model=main) | MERCI | NEGATIVE | 2 |
| 93 | 0-15 | [MADSNGTITVEELKK](http://crdd.osdd.net/raghava/ifnepitope/pep_design.php?sequence=MADSNGTITVEELKK&method=hybrid&model=main) | SVM | NEGATIVE | -0.27187013 |
| 94 | 1-16 | [ADSNGTITVEELKKL](http://crdd.osdd.net/raghava/ifnepitope/pep_design.php?sequence=ADSNGTITVEELKKL&method=hybrid&model=main) | SVM | NEGATIVE | -0.20234548 |
| 95 | 2-17 | [DSNGTITVEELKKLL](http://crdd.osdd.net/raghava/ifnepitope/pep_design.php?sequence=DSNGTITVEELKKLL&method=hybrid&model=main) | SVM | NEGATIVE | -0.17293036 |
| 96 | 3-18 | [SNGTITVEELKKLLE](http://crdd.osdd.net/raghava/ifnepitope/pep_design.php?sequence=SNGTITVEELKKLLE&method=hybrid&model=main) | SVM | NEGATIVE | -0.021610978 |
| 97 | 4-19 | [NGTITVEELKKLLEQ](http://crdd.osdd.net/raghava/ifnepitope/pep_design.php?sequence=NGTITVEELKKLLEQ&method=hybrid&model=main) | SVM | NEGATIVE | -0.17112456 |
| 98 | 5-20 | [GTITVEELKKLLEQW](http://crdd.osdd.net/raghava/ifnepitope/pep_design.php?sequence=GTITVEELKKLLEQW&method=hybrid&model=main) | SVM | POSITIVE | 0.19018912 |
| 99 | 6-21 | [TITVEELKKLLEQWN](http://crdd.osdd.net/raghava/ifnepitope/pep_design.php?sequence=TITVEELKKLLEQWN&method=hybrid&model=main) | SVM | POSITIVE | 0.27590852 |
| 100 | 7-22 | [ITVEELKKLLEQWNL](http://crdd.osdd.net/raghava/ifnepitope/pep_design.php?sequence=ITVEELKKLLEQWNL&method=hybrid&model=main) | SVM | NEGATIVE | -0.1961259 |
| 101 | 8-23 | [TVEELKKLLEQWNLV](http://crdd.osdd.net/raghava/ifnepitope/pep_design.php?sequence=TVEELKKLLEQWNLV&method=hybrid&model=main) | SVM | NEGATIVE | -0.34207592 |
| 102 | 32-47 | [CLLQFAYANRNRFLY](http://crdd.osdd.net/raghava/ifnepitope/pep_design.php?sequence=CLLQFAYANRNRFLY&method=hybrid&model=main) | SVM | NEGATIVE | -0.6467492 |
| 103 | 33-48 | [LLQFAYANRNRFLYI](http://crdd.osdd.net/raghava/ifnepitope/pep_design.php?sequence=LLQFAYANRNRFLYI&method=hybrid&model=main) | SVM | NEGATIVE | -0.59982611 |
| 104 | 34-49 | [LQFAYANRNRFLYII](http://crdd.osdd.net/raghava/ifnepitope/pep_design.php?sequence=LQFAYANRNRFLYII&method=hybrid&model=main) | SVM | NEGATIVE | -0.49947018 |
| 105 | 35-50 | [QFAYANRNRFLYIIK](http://crdd.osdd.net/raghava/ifnepitope/pep_design.php?sequence=QFAYANRNRFLYIIK&method=hybrid&model=main) | SVM | NEGATIVE | -0.39135003 |
| 106 | 52-67 | [FLWLLWPVTLACFVL](http://crdd.osdd.net/raghava/ifnepitope/pep_design.php?sequence=FLWLLWPVTLACFVL&method=hybrid&model=main) | SVM | NEGATIVE | -0.0084299978 |
| 107 | 53-68 | [LWLLWPVTLACFVLA](http://crdd.osdd.net/raghava/ifnepitope/pep_design.php?sequence=LWLLWPVTLACFVLA&method=hybrid&model=main) | SVM | POSITIVE | 0.11575149 |
| 108 | 54-69 | [WLLWPVTLACFVLAA](http://crdd.osdd.net/raghava/ifnepitope/pep_design.php?sequence=WLLWPVTLACFVLAA&method=hybrid&model=main) | SVM | POSITIVE | 0.18299498 |
| 109 | 55-70 | [LLWPVTLACFVLAAV](http://crdd.osdd.net/raghava/ifnepitope/pep_design.php?sequence=LLWPVTLACFVLAAV&method=hybrid&model=main) | SVM | POSITIVE | 0.18782542 |
| 110 | 56-71 | [LWPVTLACFVLAAVY](http://crdd.osdd.net/raghava/ifnepitope/pep_design.php?sequence=LWPVTLACFVLAAVY&method=hybrid&model=main) | SVM | POSITIVE | 0.10214626 |
| 111 | 57-72 | [WPVTLACFVLAAVYR](http://crdd.osdd.net/raghava/ifnepitope/pep_design.php?sequence=WPVTLACFVLAAVYR&method=hybrid&model=main) | SVM | POSITIVE | 0.23467354 |
| 112 | 58-73 | [PVTLACFVLAAVYRI](http://crdd.osdd.net/raghava/ifnepitope/pep_design.php?sequence=PVTLACFVLAAVYRI&method=hybrid&model=main) | SVM | POSITIVE | 0.26259643 |
| 113 | 59-74 | [VTLACFVLAAVYRIN](http://crdd.osdd.net/raghava/ifnepitope/pep_design.php?sequence=VTLACFVLAAVYRIN&method=hybrid&model=main) | SVM | POSITIVE | 0.12017983 |
| 114 | 60-75 | [TLACFVLAAVYRINW](http://crdd.osdd.net/raghava/ifnepitope/pep_design.php?sequence=TLACFVLAAVYRINW&method=hybrid&model=main) | SVM | POSITIVE | 0.04691397 |
| 115 | 61-76 | [LACFVLAAVYRINWI](http://crdd.osdd.net/raghava/ifnepitope/pep_design.php?sequence=LACFVLAAVYRINWI&method=hybrid&model=main) | SVM | POSITIVE | 0.24828858 |
| 116 | 62-77 | [ACFVLAAVYRINWIT](http://crdd.osdd.net/raghava/ifnepitope/pep_design.php?sequence=ACFVLAAVYRINWIT&method=hybrid&model=main) | SVM | NEGATIVE | -0.0024856237 |
| 117 | 63-78 | [CFVLAAVYRINWITG](http://crdd.osdd.net/raghava/ifnepitope/pep_design.php?sequence=CFVLAAVYRINWITG&method=hybrid&model=main) | SVM | NEGATIVE | -0.17932007 |
| 118 | 64-79 | [FVLAAVYRINWITGG](http://crdd.osdd.net/raghava/ifnepitope/pep_design.php?sequence=FVLAAVYRINWITGG&method=hybrid&model=main) | SVM | NEGATIVE | -0.4667889 |
| 119 | 65-80 | [VLAAVYRINWITGGI](http://crdd.osdd.net/raghava/ifnepitope/pep_design.php?sequence=VLAAVYRINWITGGI&method=hybrid&model=main) | SVM | NEGATIVE | -0.40173741 |
| 120 | 66-81 | [LAAVYRINWITGGIA](http://crdd.osdd.net/raghava/ifnepitope/pep_design.php?sequence=LAAVYRINWITGGIA&method=hybrid&model=main) | SVM | NEGATIVE | -0.23185459 |
| 121 | 80-95 | [AIAMACLVGLMWLSY](http://crdd.osdd.net/raghava/ifnepitope/pep_design.php?sequence=AIAMACLVGLMWLSY&method=hybrid&model=main) | SVM | NEGATIVE | -0.68589084 |
| 122 | 81-96 | [IAMACLVGLMWLSYF](http://crdd.osdd.net/raghava/ifnepitope/pep_design.php?sequence=IAMACLVGLMWLSYF&method=hybrid&model=main) | SVM | NEGATIVE | -0.41772627 |
| 123 | 82-97 | [AMACLVGLMWLSYFI](http://crdd.osdd.net/raghava/ifnepitope/pep_design.php?sequence=AMACLVGLMWLSYFI&method=hybrid&model=main) | SVM | NEGATIVE | -0.35362142 |
| 124 | 83-98 | [MACLVGLMWLSYFIA](http://crdd.osdd.net/raghava/ifnepitope/pep_design.php?sequence=MACLVGLMWLSYFIA&method=hybrid&model=main) | SVM | NEGATIVE | -0.42566017 |
| 125 | 84-99 | [ACLVGLMWLSYFIAS](http://crdd.osdd.net/raghava/ifnepitope/pep_design.php?sequence=ACLVGLMWLSYFIAS&method=hybrid&model=main) | SVM | NEGATIVE | -0.33975575 |
| 126 | 85-100 | [CLVGLMWLSYFIASF](http://crdd.osdd.net/raghava/ifnepitope/pep_design.php?sequence=CLVGLMWLSYFIASF&method=hybrid&model=main) | SVM | NEGATIVE | -0.25586126 |
| 127 | 86-101 | [LVGLMWLSYFIASFR](http://crdd.osdd.net/raghava/ifnepitope/pep_design.php?sequence=LVGLMWLSYFIASFR&method=hybrid&model=main) | SVM | NEGATIVE | -0.13920717 |
| 128 | 87-102 | [VGLMWLSYFIASFRL](http://crdd.osdd.net/raghava/ifnepitope/pep_design.php?sequence=VGLMWLSYFIASFRL&method=hybrid&model=main) | SVM | NEGATIVE | -0.29912806 |
| 129 | 88-103 | [GLMWLSYFIASFRLF](http://crdd.osdd.net/raghava/ifnepitope/pep_design.php?sequence=GLMWLSYFIASFRLF&method=hybrid&model=main) | SVM | NEGATIVE | -0.49329048 |
| 130 | 89-104 | [LMWLSYFIASFRLFA](http://crdd.osdd.net/raghava/ifnepitope/pep_design.php?sequence=LMWLSYFIASFRLFA&method=hybrid&model=main) | SVM | NEGATIVE | -0.52425393 |
| 131 | 90-105 | [MWLSYFIASFRLFAR](http://crdd.osdd.net/raghava/ifnepitope/pep_design.php?sequence=MWLSYFIASFRLFAR&method=hybrid&model=main) | SVM | NEGATIVE | -0.46118468 |
| 132 | 91-106 | [WLSYFIASFRLFART](http://crdd.osdd.net/raghava/ifnepitope/pep_design.php?sequence=WLSYFIASFRLFART&method=hybrid&model=main) | SVM | NEGATIVE | -0.31622612 |
| 133 | 92-107 | [LSYFIASFRLFARTR](http://crdd.osdd.net/raghava/ifnepitope/pep_design.php?sequence=LSYFIASFRLFARTR&method=hybrid&model=main) | SVM | NEGATIVE | -0.48522679 |
| 134 | 93-108 | [SYFIASFRLFARTRS](http://crdd.osdd.net/raghava/ifnepitope/pep_design.php?sequence=SYFIASFRLFARTRS&method=hybrid&model=main) | SVM | NEGATIVE | -0.32433941 |
| 135 | 94-109 | [YFIASFRLFARTRSM](http://crdd.osdd.net/raghava/ifnepitope/pep_design.php?sequence=YFIASFRLFARTRSM&method=hybrid&model=main) | SVM | NEGATIVE | -0.5440244 |
| 136 | 95-110 | [FIASFRLFARTRSMW](http://crdd.osdd.net/raghava/ifnepitope/pep_design.php?sequence=FIASFRLFARTRSMW&method=hybrid&model=main) | SVM | NEGATIVE | -0.60146246 |
| 137 | 96-111 | [IASFRLFARTRSMWS](http://crdd.osdd.net/raghava/ifnepitope/pep_design.php?sequence=IASFRLFARTRSMWS&method=hybrid&model=main) | SVM | NEGATIVE | -0.57009791 |
| 138 | 97-112 | [ASFRLFARTRSMWSF](http://crdd.osdd.net/raghava/ifnepitope/pep_design.php?sequence=ASFRLFARTRSMWSF&method=hybrid&model=main) | SVM | NEGATIVE | -0.20273129 |
| 139 | 98-113 | [SFRLFARTRSMWSFN](http://crdd.osdd.net/raghava/ifnepitope/pep_design.php?sequence=SFRLFARTRSMWSFN&method=hybrid&model=main) | SVM | NEGATIVE | -0.41279467 |
| 140 | 99-114 | [FRLFARTRSMWSFNP](http://crdd.osdd.net/raghava/ifnepitope/pep_design.php?sequence=FRLFARTRSMWSFNP&method=hybrid&model=main) | SVM | NEGATIVE | -0.74619566 |
| 141 | 100-115 | [RLFARTRSMWSFNPE](http://crdd.osdd.net/raghava/ifnepitope/pep_design.php?sequence=RLFARTRSMWSFNPE&method=hybrid&model=main) | SVM | NEGATIVE | -0.98086443 |
| 142 | 101-116 | [LFARTRSMWSFNPET](http://crdd.osdd.net/raghava/ifnepitope/pep_design.php?sequence=LFARTRSMWSFNPET&method=hybrid&model=main) | SVM | NEGATIVE | -1.1275363 |
| 143 | 102-117 | [FARTRSMWSFNPETN](http://crdd.osdd.net/raghava/ifnepitope/pep_design.php?sequence=FARTRSMWSFNPETN&method=hybrid&model=main) | SVM | NEGATIVE | -0.96077571 |
| 144 | 103-118 | [ARTRSMWSFNPETNI](http://crdd.osdd.net/raghava/ifnepitope/pep_design.php?sequence=ARTRSMWSFNPETNI&method=hybrid&model=main) | SVM | NEGATIVE | -0.98619771 |
| 145 | 104-119 | [RTRSMWSFNPETNIL](http://crdd.osdd.net/raghava/ifnepitope/pep_design.php?sequence=RTRSMWSFNPETNIL&method=hybrid&model=main) | SVM | NEGATIVE | -0.77274485 |
| 146 | 105-120 | [TRSMWSFNPETNILL](http://crdd.osdd.net/raghava/ifnepitope/pep_design.php?sequence=TRSMWSFNPETNILL&method=hybrid&model=main) | SVM | NEGATIVE | -0.94998855 |
| 147 | 106-121 | [RSMWSFNPETNILLN](http://crdd.osdd.net/raghava/ifnepitope/pep_design.php?sequence=RSMWSFNPETNILLN&method=hybrid&model=main) | SVM | NEGATIVE | -1.1290357 |
| 148 | 107-122 | [SMWSFNPETNILLNV](http://crdd.osdd.net/raghava/ifnepitope/pep_design.php?sequence=SMWSFNPETNILLNV&method=hybrid&model=main) | SVM | NEGATIVE | -1.195545 |
| 149 | 108-123 | [MWSFNPETNILLNVP](http://crdd.osdd.net/raghava/ifnepitope/pep_design.php?sequence=MWSFNPETNILLNVP&method=hybrid&model=main) | SVM | NEGATIVE | -0.98769647 |
| 150 | 109-124 | [WSFNPETNILLNVPL](http://crdd.osdd.net/raghava/ifnepitope/pep_design.php?sequence=WSFNPETNILLNVPL&method=hybrid&model=main) | SVM | NEGATIVE | -0.93062955 |
| 151 | 110-125 | [SFNPETNILLNVPLH](http://crdd.osdd.net/raghava/ifnepitope/pep_design.php?sequence=SFNPETNILLNVPLH&method=hybrid&model=main) | SVM | NEGATIVE | -1.051891 |
| 152 | 111-126 | [FNPETNILLNVPLHG](http://crdd.osdd.net/raghava/ifnepitope/pep_design.php?sequence=FNPETNILLNVPLHG&method=hybrid&model=main) | SVM | NEGATIVE | -0.94887501 |
| 153 | 112-127 | [NPETNILLNVPLHGT](http://crdd.osdd.net/raghava/ifnepitope/pep_design.php?sequence=NPETNILLNVPLHGT&method=hybrid&model=main) | SVM | NEGATIVE | -0.67274843 |
| 154 | 113-128 | [PETNILLNVPLHGTI](http://crdd.osdd.net/raghava/ifnepitope/pep_design.php?sequence=PETNILLNVPLHGTI&method=hybrid&model=main) | SVM | NEGATIVE | -0.57944131 |
| 155 | 114-129 | [ETNILLNVPLHGTIL](http://crdd.osdd.net/raghava/ifnepitope/pep_design.php?sequence=ETNILLNVPLHGTIL&method=hybrid&model=main) | SVM | NEGATIVE | -0.24614503 |
| 156 | 115-130 | [TNILLNVPLHGTILT](http://crdd.osdd.net/raghava/ifnepitope/pep_design.php?sequence=TNILLNVPLHGTILT&method=hybrid&model=main) | SVM | POSITIVE | 0.038813 |
| 157 | 116-131 | [NILLNVPLHGTILTR](http://crdd.osdd.net/raghava/ifnepitope/pep_design.php?sequence=NILLNVPLHGTILTR&method=hybrid&model=main) | SVM | POSITIVE | 0.06386843 |
| 158 | 117-132 | [ILLNVPLHGTILTRP](http://crdd.osdd.net/raghava/ifnepitope/pep_design.php?sequence=ILLNVPLHGTILTRP&method=hybrid&model=main) | SVM | POSITIVE | 0.33047918 |
| 159 | 118-133 | [LLNVPLHGTILTRPL](http://crdd.osdd.net/raghava/ifnepitope/pep_design.php?sequence=LLNVPLHGTILTRPL&method=hybrid&model=main) | SVM | POSITIVE | 0.23166217 |
| 160 | 119-134 | [LNVPLHGTILTRPLL](http://crdd.osdd.net/raghava/ifnepitope/pep_design.php?sequence=LNVPLHGTILTRPLL&method=hybrid&model=main) | SVM | POSITIVE | 0.23166217 |
| 161 | 120-135 | [NVPLHGTILTRPLLE](http://crdd.osdd.net/raghava/ifnepitope/pep_design.php?sequence=NVPLHGTILTRPLLE&method=hybrid&model=main) | SVM | POSITIVE | 0.075180357 |
| 162 | 121-136 | [VPLHGTILTRPLLES](http://crdd.osdd.net/raghava/ifnepitope/pep_design.php?sequence=VPLHGTILTRPLLES&method=hybrid&model=main) | SVM | POSITIVE | 0.1993441 |
| 163 | 122-137 | [PLHGTILTRPLLESE](http://crdd.osdd.net/raghava/ifnepitope/pep_design.php?sequence=PLHGTILTRPLLESE&method=hybrid&model=main) | SVM | POSITIVE | 0.063406441 |
| 164 | 124-139 | [HGTILTRPLLESELV](http://crdd.osdd.net/raghava/ifnepitope/pep_design.php?sequence=HGTILTRPLLESELV&method=hybrid&model=main) | SVM | POSITIVE | 0.21914036 |
| 165 | 140-155 | [GAVILRGHLRIAGHH](http://crdd.osdd.net/raghava/ifnepitope/pep_design.php?sequence=GAVILRGHLRIAGHH&method=hybrid&model=main) | SVM | POSITIVE | 0.025155483 |
| 166 | 141-156 | [AVILRGHLRIAGHHL](http://crdd.osdd.net/raghava/ifnepitope/pep_design.php?sequence=AVILRGHLRIAGHHL&method=hybrid&model=main) | SVM | POSITIVE | 0.12415297 |
| 167 | 142-157 | [VILRGHLRIAGHHLG](http://crdd.osdd.net/raghava/ifnepitope/pep_design.php?sequence=VILRGHLRIAGHHLG&method=hybrid&model=main) | SVM | POSITIVE | 0.28233564 |
| 168 | 143-158 | [ILRGHLRIAGHHLGR](http://crdd.osdd.net/raghava/ifnepitope/pep_design.php?sequence=ILRGHLRIAGHHLGR&method=hybrid&model=main) | SVM | POSITIVE | 0.051671066 |
| 169 | 144-159 | [LRGHLRIAGHHLGRC](http://crdd.osdd.net/raghava/ifnepitope/pep_design.php?sequence=LRGHLRIAGHHLGRC&method=hybrid&model=main) | SVM | POSITIVE | 0.055865543 |
| 170 | 145-160 | [RGHLRIAGHHLGRCD](http://crdd.osdd.net/raghava/ifnepitope/pep_design.php?sequence=RGHLRIAGHHLGRCD&method=hybrid&model=main) | SVM | NEGATIVE | -0.3630421 |
| 171 | 146-161 | [GHLRIAGHHLGRCDI](http://crdd.osdd.net/raghava/ifnepitope/pep_design.php?sequence=GHLRIAGHHLGRCDI&method=hybrid&model=main) | SVM | NEGATIVE | -0.40307383 |
| 172 | 147-162 | [HLRIAGHHLGRCDIK](http://crdd.osdd.net/raghava/ifnepitope/pep_design.php?sequence=HLRIAGHHLGRCDIK&method=hybrid&model=main) | SVM | NEGATIVE | -0.50417831 |
| 173 | 155-170 | [LGRCDIKDLPKEITV](http://crdd.osdd.net/raghava/ifnepitope/pep_design.php?sequence=LGRCDIKDLPKEITV&method=hybrid&model=main) | SVM | NEGATIVE | -0.54044826 |
| 174 | 156-171 | [GRCDIKDLPKEITVA](http://crdd.osdd.net/raghava/ifnepitope/pep_design.php?sequence=GRCDIKDLPKEITVA&method=hybrid&model=main) | SVM | NEGATIVE | -0.59617535 |
| 175 | 157-172 | [RCDIKDLPKEITVAT](http://crdd.osdd.net/raghava/ifnepitope/pep_design.php?sequence=RCDIKDLPKEITVAT&method=hybrid&model=main) | SVM | NEGATIVE | -0.45025607 |
| 176 | 158-173 | [CDIKDLPKEITVATS](http://crdd.osdd.net/raghava/ifnepitope/pep_design.php?sequence=CDIKDLPKEITVATS&method=hybrid&model=main) | SVM | NEGATIVE | -0.48481219 |
| 177 | 159-174 | [DIKDLPKEITVATSR](http://crdd.osdd.net/raghava/ifnepitope/pep_design.php?sequence=DIKDLPKEITVATSR&method=hybrid&model=main) | SVM | NEGATIVE | -0.39440361 |
| 178 | 160-175 | [IKDLPKEITVATSRT](http://crdd.osdd.net/raghava/ifnepitope/pep_design.php?sequence=IKDLPKEITVATSRT&method=hybrid&model=main) | SVM | NEGATIVE | -0.51513849 |
| 179 | 161-176 | [KDLPKEITVATSRTL](http://crdd.osdd.net/raghava/ifnepitope/pep_design.php?sequence=KDLPKEITVATSRTL&method=hybrid&model=main) | SVM | NEGATIVE | -0.6154126 |
| 180 | 162-177 | [DLPKEITVATSRTLS](http://crdd.osdd.net/raghava/ifnepitope/pep_design.php?sequence=DLPKEITVATSRTLS&method=hybrid&model=main) | SVM | NEGATIVE | -0.60228056 |
| 181 | 163-178 | [LPKEITVATSRTLSY](http://crdd.osdd.net/raghava/ifnepitope/pep_design.php?sequence=LPKEITVATSRTLSY&method=hybrid&model=main) | SVM | NEGATIVE | -0.45709636 |
| 182 | 164-179 | [PKEITVATSRTLSYY](http://crdd.osdd.net/raghava/ifnepitope/pep_design.php?sequence=PKEITVATSRTLSYY&method=hybrid&model=main) | SVM | NEGATIVE | -0.21379522 |
| 183 | 173-188 | [RTLSYYKLGASQRVA](http://crdd.osdd.net/raghava/ifnepitope/pep_design.php?sequence=RTLSYYKLGASQRVA&method=hybrid&model=main) | SVM | POSITIVE | 0.21078571 |
| 184 | 174-189 | [TLSYYKLGASQRVAG](http://crdd.osdd.net/raghava/ifnepitope/pep_design.php?sequence=TLSYYKLGASQRVAG&method=hybrid&model=main) | SVM | NEGATIVE | -0.0025116004 |
| 185 | 175-190 | [LSYYKLGASQRVAGD](http://crdd.osdd.net/raghava/ifnepitope/pep_design.php?sequence=LSYYKLGASQRVAGD&method=hybrid&model=main) | SVM | NEGATIVE | -0.1153439 |
| 186 | 176-191 | [SYYKLGASQRVAGDS](http://crdd.osdd.net/raghava/ifnepitope/pep_design.php?sequence=SYYKLGASQRVAGDS&method=hybrid&model=main) | SVM | POSITIVE | 0.1701859 |
| 187 | 177-192 | [YYKLGASQRVAGDSG](http://crdd.osdd.net/raghava/ifnepitope/pep_design.php?sequence=YYKLGASQRVAGDSG&method=hybrid&model=main) | SVM | NEGATIVE | -0.093445753 |
| 188 | 178-193 | [YKLGASQRVAGDSGF](http://crdd.osdd.net/raghava/ifnepitope/pep_design.php?sequence=YKLGASQRVAGDSGF&method=hybrid&model=main) | SVM | POSITIVE | 0.061331393 |
| 189 | 179-194 | [KLGASQRVAGDSGFA](http://crdd.osdd.net/raghava/ifnepitope/pep_design.php?sequence=KLGASQRVAGDSGFA&method=hybrid&model=main) | SVM | POSITIVE | 0.1061902 |
| 190 | 180-195 | [LGASQRVAGDSGFAA](http://crdd.osdd.net/raghava/ifnepitope/pep_design.php?sequence=LGASQRVAGDSGFAA&method=hybrid&model=main) | SVM | NEGATIVE | -0.10291477 |
| 191 | 181-196 | [GASQRVAGDSGFAAY](http://crdd.osdd.net/raghava/ifnepitope/pep_design.php?sequence=GASQRVAGDSGFAAY&method=hybrid&model=main) | SVM | NEGATIVE | -0.019580299 |
| 192 | 182-197 | [ASQRVAGDSGFAAYS](http://crdd.osdd.net/raghava/ifnepitope/pep_design.php?sequence=ASQRVAGDSGFAAYS&method=hybrid&model=main) | SVM | NEGATIVE | -0.072637369 |
| 193 | 183-198 | [SQRVAGDSGFAAYSR](http://crdd.osdd.net/raghava/ifnepitope/pep_design.php?sequence=SQRVAGDSGFAAYSR&method=hybrid&model=main) | SVM | NEGATIVE | -0.33872072 |
| 194 | 184-199 | [QRVAGDSGFAAYSRY](http://crdd.osdd.net/raghava/ifnepitope/pep_design.php?sequence=QRVAGDSGFAAYSRY&method=hybrid&model=main) | SVM | NEGATIVE | -0.17490742 |
| 195 | 185-200 | [RVAGDSGFAAYSRYR](http://crdd.osdd.net/raghava/ifnepitope/pep_design.php?sequence=RVAGDSGFAAYSRYR&method=hybrid&model=main) | SVM | NEGATIVE | -0.26891132 |
| 196 | 186-201 | [VAGDSGFAAYSRYRI](http://crdd.osdd.net/raghava/ifnepitope/pep_design.php?sequence=VAGDSGFAAYSRYRI&method=hybrid&model=main) | SVM | NEGATIVE | -0.49654106 |
| 197 | 187-202 | [AGDSGFAAYSRYRIG](http://crdd.osdd.net/raghava/ifnepitope/pep_design.php?sequence=AGDSGFAAYSRYRIG&method=hybrid&model=main) | SVM | NEGATIVE | -0.69157708 |
| 198 | 188-203 | [GDSGFAAYSRYRIGN](http://crdd.osdd.net/raghava/ifnepitope/pep_design.php?sequence=GDSGFAAYSRYRIGN&method=hybrid&model=main) | SVM | NEGATIVE | -0.69193589 |
| 199 | 189-204 | [DSGFAAYSRYRIGNY](http://crdd.osdd.net/raghava/ifnepitope/pep_design.php?sequence=DSGFAAYSRYRIGNY&method=hybrid&model=main) | SVM | NEGATIVE | -0.68434897 |
| 200 | 190-205 | [SGFAAYSRYRIGNYK](http://crdd.osdd.net/raghava/ifnepitope/pep_design.php?sequence=SGFAAYSRYRIGNYK&method=hybrid&model=main) | SVM | NEGATIVE | -0.88971678 |
| 201 | 191-206 | [GFAAYSRYRIGNYKL](http://crdd.osdd.net/raghava/ifnepitope/pep_design.php?sequence=GFAAYSRYRIGNYKL&method=hybrid&model=main) | SVM | NEGATIVE | -0.6527623 |
| 202 | 192-207 | [FAAYSRYRIGNYKLN](http://crdd.osdd.net/raghava/ifnepitope/pep_design.php?sequence=FAAYSRYRIGNYKLN&method=hybrid&model=main) | SVM | NEGATIVE | -0.65338731 |
| 203 | 193-208 | [AAYSRYRIGNYKLNT](http://crdd.osdd.net/raghava/ifnepitope/pep_design.php?sequence=AAYSRYRIGNYKLNT&method=hybrid&model=main) | SVM | NEGATIVE | -0.69283836 |
| 204 | 194-209 | [AYSRYRIGNYKLNTD](http://crdd.osdd.net/raghava/ifnepitope/pep_design.php?sequence=AYSRYRIGNYKLNTD&method=hybrid&model=main) | SVM | NEGATIVE | -0.61576668 |
| 205 | 195-210 | [YSRYRIGNYKLNTDH](http://crdd.osdd.net/raghava/ifnepitope/pep_design.php?sequence=YSRYRIGNYKLNTDH&method=hybrid&model=main) | SVM | NEGATIVE | -0.67234922 |
| 206 | 196-211 | [SRYRIGNYKLNTDHS](http://crdd.osdd.net/raghava/ifnepitope/pep_design.php?sequence=SRYRIGNYKLNTDHS&method=hybrid&model=main) | SVM | NEGATIVE | -0.4747254 |
| 207 | 197-212 | [RYRIGNYKLNTDHSS](http://crdd.osdd.net/raghava/ifnepitope/pep_design.php?sequence=RYRIGNYKLNTDHSS&method=hybrid&model=main) | SVM | NEGATIVE | -0.3171153 |
| 208 | 198-213 | [YRIGNYKLNTDHSSS](http://crdd.osdd.net/raghava/ifnepitope/pep_design.php?sequence=YRIGNYKLNTDHSSS&method=hybrid&model=main) | SVM | NEGATIVE | -0.27227262 |
| 209 | 199-214 | [RIGNYKLNTDHSSSS](http://crdd.osdd.net/raghava/ifnepitope/pep_design.php?sequence=RIGNYKLNTDHSSSS&method=hybrid&model=main) | SVM | NEGATIVE | -0.10777627 |
| 210 | 200-215 | [IGNYKLNTDHSSSSD](http://crdd.osdd.net/raghava/ifnepitope/pep_design.php?sequence=IGNYKLNTDHSSSSD&method=hybrid&model=main) | SVM | NEGATIVE | -0.084715512 |
| 211 | 201-216 | [GNYKLNTDHSSSSDN](http://crdd.osdd.net/raghava/ifnepitope/pep_design.php?sequence=GNYKLNTDHSSSSDN&method=hybrid&model=main) | SVM | POSITIVE | 0.059113387 |
| 212 | 202-217 | [NYKLNTDHSSSSDNI](http://crdd.osdd.net/raghava/ifnepitope/pep_design.php?sequence=NYKLNTDHSSSSDNI&method=hybrid&model=main) | SVM | NEGATIVE | -0.020933113 |
| 213 | 203-218 | [YKLNTDHSSSSDNIA](http://crdd.osdd.net/raghava/ifnepitope/pep_design.php?sequence=YKLNTDHSSSSDNIA&method=hybrid&model=main) | SVM | NEGATIVE | -0.052051428 |
| 214 | 204-219 | [KLNTDHSSSSDNIAL](http://crdd.osdd.net/raghava/ifnepitope/pep_design.php?sequence=KLNTDHSSSSDNIAL&method=hybrid&model=main) | SVM | POSITIVE | 0.21377461 |
| 1 | 5-20 | [PQNQRNAPRITFGGP](http://crdd.osdd.net/raghava/ifnepitope/pep_design.php?sequence=PQNQRNAPRITFGGP&method=hybrid&model=main) | MERCI | POSITIVE | 1 |
| N Protein | | | | | |
| 2 | 8-23 | [QRNAPRITFGGPSDS](http://crdd.osdd.net/raghava/ifnepitope/pep_design.php?sequence=QRNAPRITFGGPSDS&method=hybrid&model=main) | MERCI | POSITIVE | 1 |
| 3 | 36-51 | [SKQRRPQGLPNNTAS](http://crdd.osdd.net/raghava/ifnepitope/pep_design.php?sequence=SKQRRPQGLPNNTAS&method=hybrid&model=main) | MERCI | POSITIVE | 2 |
| 4 | 37-52 | [KQRRPQGLPNNTASW](http://crdd.osdd.net/raghava/ifnepitope/pep_design.php?sequence=KQRRPQGLPNNTASW&method=hybrid&model=main) | MERCI | POSITIVE | 2 |
| 5 | 38-53 | [QRRPQGLPNNTASWF](http://crdd.osdd.net/raghava/ifnepitope/pep_design.php?sequence=QRRPQGLPNNTASWF&method=hybrid&model=main) | MERCI | POSITIVE | 2 |
| 6 | 39-54 | [RRPQGLPNNTASWFT](http://crdd.osdd.net/raghava/ifnepitope/pep_design.php?sequence=RRPQGLPNNTASWFT&method=hybrid&model=main) | MERCI | POSITIVE | 2 |
| 7 | 40-55 | [RPQGLPNNTASWFTA](http://crdd.osdd.net/raghava/ifnepitope/pep_design.php?sequence=RPQGLPNNTASWFTA&method=hybrid&model=main) | MERCI | POSITIVE | 2 |
| 8 | 41-56 | [PQGLPNNTASWFTAL](http://crdd.osdd.net/raghava/ifnepitope/pep_design.php?sequence=PQGLPNNTASWFTAL&method=hybrid&model=main) | MERCI | POSITIVE | 3 |
| 9 | 42-57 | [QGLPNNTASWFTALT](http://crdd.osdd.net/raghava/ifnepitope/pep_design.php?sequence=QGLPNNTASWFTALT&method=hybrid&model=main) | MERCI | POSITIVE | 1 |
| 10 | 43-58 | [GLPNNTASWFTALTQ](http://crdd.osdd.net/raghava/ifnepitope/pep_design.php?sequence=GLPNNTASWFTALTQ&method=hybrid&model=main) | MERCI | POSITIVE | 1 |
| 11 | 44-59 | [LPNNTASWFTALTQH](http://crdd.osdd.net/raghava/ifnepitope/pep_design.php?sequence=LPNNTASWFTALTQH&method=hybrid&model=main) | MERCI | POSITIVE | 1 |
| 12 | 45-60 | [PNNTASWFTALTQHG](http://crdd.osdd.net/raghava/ifnepitope/pep_design.php?sequence=PNNTASWFTALTQHG&method=hybrid&model=main) | MERCI | POSITIVE | 1 |
| 13 | 46-61 | [NNTASWFTALTQHGK](http://crdd.osdd.net/raghava/ifnepitope/pep_design.php?sequence=NNTASWFTALTQHGK&method=hybrid&model=main) | MERCI | POSITIVE | 1 |
| 14 | 47-62 | [NTASWFTALTQHGKE](http://crdd.osdd.net/raghava/ifnepitope/pep_design.php?sequence=NTASWFTALTQHGKE&method=hybrid&model=main) | MERCI | POSITIVE | 1 |
| 15 | 48-63 | [TASWFTALTQHGKED](http://crdd.osdd.net/raghava/ifnepitope/pep_design.php?sequence=TASWFTALTQHGKED&method=hybrid&model=main) | MERCI | POSITIVE | 1 |
| 16 | 68-83 | [GQGVPINTNSSPDDQ](http://crdd.osdd.net/raghava/ifnepitope/pep_design.php?sequence=GQGVPINTNSSPDDQ&method=hybrid&model=main) | MERCI | POSITIVE | 1 |
| 17 | 69-84 | [QGVPINTNSSPDDQI](http://crdd.osdd.net/raghava/ifnepitope/pep_design.php?sequence=QGVPINTNSSPDDQI&method=hybrid&model=main) | MERCI | POSITIVE | 2 |
| 18 | 291-306 | [IRQGTDYKHWPQIAQ](http://crdd.osdd.net/raghava/ifnepitope/pep_design.php?sequence=IRQGTDYKHWPQIAQ&method=hybrid&model=main) | MERCI | POSITIVE | 4 |
| 19 | 292-307 | [RQGTDYKHWPQIAQF](http://crdd.osdd.net/raghava/ifnepitope/pep_design.php?sequence=RQGTDYKHWPQIAQF&method=hybrid&model=main) | MERCI | POSITIVE | 4 |
| 20 | 293-308 | [QGTDYKHWPQIAQFA](http://crdd.osdd.net/raghava/ifnepitope/pep_design.php?sequence=QGTDYKHWPQIAQFA&method=hybrid&model=main) | MERCI | POSITIVE | 4 |
| 21 | 297-312 | [YKHWPQIAQFAPSAS](http://crdd.osdd.net/raghava/ifnepitope/pep_design.php?sequence=YKHWPQIAQFAPSAS&method=hybrid&model=main) | MERCI | POSITIVE | 1 |
| 22 | 298-313 | [KHWPQIAQFAPSASA](http://crdd.osdd.net/raghava/ifnepitope/pep_design.php?sequence=KHWPQIAQFAPSASA&method=hybrid&model=main) | MERCI | POSITIVE | 1 |
| 23 | 299-314 | [HWPQIAQFAPSASAF](http://crdd.osdd.net/raghava/ifnepitope/pep_design.php?sequence=HWPQIAQFAPSASAF&method=hybrid&model=main) | MERCI | POSITIVE | 1 |
| 24 | 300-315 | [WPQIAQFAPSASAFF](http://crdd.osdd.net/raghava/ifnepitope/pep_design.php?sequence=WPQIAQFAPSASAFF&method=hybrid&model=main) | MERCI | POSITIVE | 1 |
| 25 | 301-316 | [PQIAQFAPSASAFFG](http://crdd.osdd.net/raghava/ifnepitope/pep_design.php?sequence=PQIAQFAPSASAFFG&method=hybrid&model=main) | MERCI | POSITIVE | 1 |
| 26 | 302-317 | [QIAQFAPSASAFFGM](http://crdd.osdd.net/raghava/ifnepitope/pep_design.php?sequence=QIAQFAPSASAFFGM&method=hybrid&model=main) | MERCI | POSITIVE | 1 |
| 27 | 398-413 | [DLDDFSKQLQQSMSS](http://crdd.osdd.net/raghava/ifnepitope/pep_design.php?sequence=DLDDFSKQLQQSMSS&method=hybrid&model=main) | MERCI | POSITIVE | 1 |
| 28 | 399-414 | [LDDFSKQLQQSMSSA](http://crdd.osdd.net/raghava/ifnepitope/pep_design.php?sequence=LDDFSKQLQQSMSSA&method=hybrid&model=main) | MERCI | POSITIVE | 1 |
| 29 | 400-415 | [DDFSKQLQQSMSSAD](http://crdd.osdd.net/raghava/ifnepitope/pep_design.php?sequence=DDFSKQLQQSMSSAD&method=hybrid&model=main) | MERCI | POSITIVE | 1 |
| 30 | 401-416 | [DFSKQLQQSMSSADS](http://crdd.osdd.net/raghava/ifnepitope/pep_design.php?sequence=DFSKQLQQSMSSADS&method=hybrid&model=main) | MERCI | POSITIVE | 1 |
| 31 | 15-30 | [TFGGPSDSTGSNQNG](http://crdd.osdd.net/raghava/ifnepitope/pep_design.php?sequence=TFGGPSDSTGSNQNG&method=hybrid&model=main) | MERCI | NEGATIVE | 1 |
| 32 | 16-31 | [FGGPSDSTGSNQNGE](http://crdd.osdd.net/raghava/ifnepitope/pep_design.php?sequence=FGGPSDSTGSNQNGE&method=hybrid&model=main) | MERCI | NEGATIVE | 1 |
| 33 | 17-32 | [GGPSDSTGSNQNGER](http://crdd.osdd.net/raghava/ifnepitope/pep_design.php?sequence=GGPSDSTGSNQNGER&method=hybrid&model=main) | MERCI | NEGATIVE | 1 |
| 34 | 18-33 | [GPSDSTGSNQNGERS](http://crdd.osdd.net/raghava/ifnepitope/pep_design.php?sequence=GPSDSTGSNQNGERS&method=hybrid&model=main) | MERCI | NEGATIVE | 1 |
| 35 | 19-34 | [PSDSTGSNQNGERSG](http://crdd.osdd.net/raghava/ifnepitope/pep_design.php?sequence=PSDSTGSNQNGERSG&method=hybrid&model=main) | MERCI | NEGATIVE | 1 |
| 36 | 20-35 | [SDSTGSNQNGERSGA](http://crdd.osdd.net/raghava/ifnepitope/pep_design.php?sequence=SDSTGSNQNGERSGA&method=hybrid&model=main) | MERCI | NEGATIVE | 1 |
| 37 | 90-105 | [TRRIRGGDGKMKDLS](http://crdd.osdd.net/raghava/ifnepitope/pep_design.php?sequence=TRRIRGGDGKMKDLS&method=hybrid&model=main) | MERCI | NEGATIVE | 1 |
| 38 | 91-106 | [RRIRGGDGKMKDLSP](http://crdd.osdd.net/raghava/ifnepitope/pep_design.php?sequence=RRIRGGDGKMKDLSP&method=hybrid&model=main) | MERCI | NEGATIVE | 1 |
| 39 | 92-107 | [RIRGGDGKMKDLSPR](http://crdd.osdd.net/raghava/ifnepitope/pep_design.php?sequence=RIRGGDGKMKDLSPR&method=hybrid&model=main) | MERCI | NEGATIVE | 1 |
| 40 | 93-108 | [IRGGDGKMKDLSPRW](http://crdd.osdd.net/raghava/ifnepitope/pep_design.php?sequence=IRGGDGKMKDLSPRW&method=hybrid&model=main) | MERCI | NEGATIVE | 1 |
| 41 | 94-109 | [RGGDGKMKDLSPRWY](http://crdd.osdd.net/raghava/ifnepitope/pep_design.php?sequence=RGGDGKMKDLSPRWY&method=hybrid&model=main) | MERCI | NEGATIVE | 1 |
| 42 | 119-134 | [GLPYGANKDGIIWVA](http://crdd.osdd.net/raghava/ifnepitope/pep_design.php?sequence=GLPYGANKDGIIWVA&method=hybrid&model=main) | MERCI | NEGATIVE | 4 |
| 43 | 120-135 | [LPYGANKDGIIWVAT](http://crdd.osdd.net/raghava/ifnepitope/pep_design.php?sequence=LPYGANKDGIIWVAT&method=hybrid&model=main) | MERCI | NEGATIVE | 6 |
| 44 | 121-136 | [PYGANKDGIIWVATE](http://crdd.osdd.net/raghava/ifnepitope/pep_design.php?sequence=PYGANKDGIIWVATE&method=hybrid&model=main) | MERCI | NEGATIVE | 6 |
| 45 | 122-137 | [YGANKDGIIWVATEG](http://crdd.osdd.net/raghava/ifnepitope/pep_design.php?sequence=YGANKDGIIWVATEG&method=hybrid&model=main) | MERCI | NEGATIVE | 6 |
| 46 | 123-138 | [GANKDGIIWVATEGA](http://crdd.osdd.net/raghava/ifnepitope/pep_design.php?sequence=GANKDGIIWVATEGA&method=hybrid&model=main) | MERCI | NEGATIVE | 7 |
| 47 | 124-139 | [ANKDGIIWVATEGAL](http://crdd.osdd.net/raghava/ifnepitope/pep_design.php?sequence=ANKDGIIWVATEGAL&method=hybrid&model=main) | MERCI | NEGATIVE | 7 |
| 48 | 125-140 | [NKDGIIWVATEGALN](http://crdd.osdd.net/raghava/ifnepitope/pep_design.php?sequence=NKDGIIWVATEGALN&method=hybrid&model=main) | MERCI | NEGATIVE | 7 |
| 49 | 126-141 | [KDGIIWVATEGALNT](http://crdd.osdd.net/raghava/ifnepitope/pep_design.php?sequence=KDGIIWVATEGALNT&method=hybrid&model=main) | MERCI | NEGATIVE | 6 |
| 50 | 127-142 | [DGIIWVATEGALNTP](http://crdd.osdd.net/raghava/ifnepitope/pep_design.php?sequence=DGIIWVATEGALNTP&method=hybrid&model=main) | MERCI | NEGATIVE | 6 |
| 51 | 128-143 | [GIIWVATEGALNTPK](http://crdd.osdd.net/raghava/ifnepitope/pep_design.php?sequence=GIIWVATEGALNTPK&method=hybrid&model=main) | MERCI | NEGATIVE | 3 |
| 52 | 129-144 | [IIWVATEGALNTPKD](http://crdd.osdd.net/raghava/ifnepitope/pep_design.php?sequence=IIWVATEGALNTPKD&method=hybrid&model=main) | MERCI | NEGATIVE | 2 |
| 53 | 144-159 | [HIGTRNPANNAAIVL](http://crdd.osdd.net/raghava/ifnepitope/pep_design.php?sequence=HIGTRNPANNAAIVL&method=hybrid&model=main) | MERCI | NEGATIVE | 2 |
| 54 | 145-160 | [IGTRNPANNAAIVLQ](http://crdd.osdd.net/raghava/ifnepitope/pep_design.php?sequence=IGTRNPANNAAIVLQ&method=hybrid&model=main) | MERCI | NEGATIVE | 2 |
| 55 | 146-161 | [GTRNPANNAAIVLQL](http://crdd.osdd.net/raghava/ifnepitope/pep_design.php?sequence=GTRNPANNAAIVLQL&method=hybrid&model=main) | MERCI | NEGATIVE | 1 |
| 56 | 147-162 | [TRNPANNAAIVLQLP](http://crdd.osdd.net/raghava/ifnepitope/pep_design.php?sequence=TRNPANNAAIVLQLP&method=hybrid&model=main) | MERCI | NEGATIVE | 1 |
| 57 | 148-163 | [RNPANNAAIVLQLPQ](http://crdd.osdd.net/raghava/ifnepitope/pep_design.php?sequence=RNPANNAAIVLQLPQ&method=hybrid&model=main) | MERCI | NEGATIVE | 1 |
| 58 | 149-164 | [NPANNAAIVLQLPQG](http://crdd.osdd.net/raghava/ifnepitope/pep_design.php?sequence=NPANNAAIVLQLPQG&method=hybrid&model=main) | MERCI | NEGATIVE | 1 |
| 59 | 150-165 | [PANNAAIVLQLPQGT](http://crdd.osdd.net/raghava/ifnepitope/pep_design.php?sequence=PANNAAIVLQLPQGT&method=hybrid&model=main) | MERCI | NEGATIVE | 1 |
| 60 | 151-166 | [ANNAAIVLQLPQGTT](http://crdd.osdd.net/raghava/ifnepitope/pep_design.php?sequence=ANNAAIVLQLPQGTT&method=hybrid&model=main) | MERCI | NEGATIVE | 1 |
| 61 | 152-167 | [NNAAIVLQLPQGTTL](http://crdd.osdd.net/raghava/ifnepitope/pep_design.php?sequence=NNAAIVLQLPQGTTL&method=hybrid&model=main) | MERCI | NEGATIVE | 1 |
| 62 | 169-184 | [GFYAEGSRGGSQASS](http://crdd.osdd.net/raghava/ifnepitope/pep_design.php?sequence=GFYAEGSRGGSQASS&method=hybrid&model=main) | MERCI | NEGATIVE | 1 |
| 63 | 170-185 | [FYAEGSRGGSQASSR](http://crdd.osdd.net/raghava/ifnepitope/pep_design.php?sequence=FYAEGSRGGSQASSR&method=hybrid&model=main) | MERCI | NEGATIVE | 1 |
| 64 | 171-186 | [YAEGSRGGSQASSRS](http://crdd.osdd.net/raghava/ifnepitope/pep_design.php?sequence=YAEGSRGGSQASSRS&method=hybrid&model=main) | MERCI | NEGATIVE | 1 |
| 65 | 172-187 | [AEGSRGGSQASSRSS](http://crdd.osdd.net/raghava/ifnepitope/pep_design.php?sequence=AEGSRGGSQASSRSS&method=hybrid&model=main) | MERCI | NEGATIVE | 1 |
| 66 | 173-188 | [EGSRGGSQASSRSSS](http://crdd.osdd.net/raghava/ifnepitope/pep_design.php?sequence=EGSRGGSQASSRSSS&method=hybrid&model=main) | MERCI | NEGATIVE | 1 |
| 67 | 174-189 | [GSRGGSQASSRSSSR](http://crdd.osdd.net/raghava/ifnepitope/pep_design.php?sequence=GSRGGSQASSRSSSR&method=hybrid&model=main) | MERCI | NEGATIVE | 1 |
| 68 | 175-190 | [SRGGSQASSRSSSRS](http://crdd.osdd.net/raghava/ifnepitope/pep_design.php?sequence=SRGGSQASSRSSSRS&method=hybrid&model=main) | MERCI | NEGATIVE | 1 |
| 69 | 176-191 | [RGGSQASSRSSSRSR](http://crdd.osdd.net/raghava/ifnepitope/pep_design.php?sequence=RGGSQASSRSSSRSR&method=hybrid&model=main) | MERCI | NEGATIVE | 1 |
| 70 | 177-192 | [GGSQASSRSSSRSRN](http://crdd.osdd.net/raghava/ifnepitope/pep_design.php?sequence=GGSQASSRSSSRSRN&method=hybrid&model=main) | MERCI | NEGATIVE | 1 |
| 71 | 178-193 | [GSQASSRSSSRSRNS](http://crdd.osdd.net/raghava/ifnepitope/pep_design.php?sequence=GSQASSRSSSRSRNS&method=hybrid&model=main) | MERCI | NEGATIVE | 1 |
| 72 | 179-194 | [SQASSRSSSRSRNSS](http://crdd.osdd.net/raghava/ifnepitope/pep_design.php?sequence=SQASSRSSSRSRNSS&method=hybrid&model=main) | MERCI | NEGATIVE | 1 |
| 73 | 180-195 | [QASSRSSSRSRNSSR](http://crdd.osdd.net/raghava/ifnepitope/pep_design.php?sequence=QASSRSSSRSRNSSR&method=hybrid&model=main) | MERCI | NEGATIVE | 1 |
| 74 | 189-204 | [SRNSSRNSTPGSSRG](http://crdd.osdd.net/raghava/ifnepitope/pep_design.php?sequence=SRNSSRNSTPGSSRG&method=hybrid&model=main) | MERCI | NEGATIVE | 1 |
| 75 | 190-205 | [RNSSRNSTPGSSRGT](http://crdd.osdd.net/raghava/ifnepitope/pep_design.php?sequence=RNSSRNSTPGSSRGT&method=hybrid&model=main) | MERCI | NEGATIVE | 1 |
| 76 | 191-206 | [NSSRNSTPGSSRGTS](http://crdd.osdd.net/raghava/ifnepitope/pep_design.php?sequence=NSSRNSTPGSSRGTS&method=hybrid&model=main) | MERCI | NEGATIVE | 2 |
| 77 | 192-207 | [SSRNSTPGSSRGTSP](http://crdd.osdd.net/raghava/ifnepitope/pep_design.php?sequence=SSRNSTPGSSRGTSP&method=hybrid&model=main) | MERCI | NEGATIVE | 2 |
| 78 | 193-208 | [SRNSTPGSSRGTSPA](http://crdd.osdd.net/raghava/ifnepitope/pep_design.php?sequence=SRNSTPGSSRGTSPA&method=hybrid&model=main) | MERCI | NEGATIVE | 2 |
| 79 | 194-209 | [RNSTPGSSRGTSPAR](http://crdd.osdd.net/raghava/ifnepitope/pep_design.php?sequence=RNSTPGSSRGTSPAR&method=hybrid&model=main) | MERCI | NEGATIVE | 2 |
| 80 | 195-210 | [NSTPGSSRGTSPARM](http://crdd.osdd.net/raghava/ifnepitope/pep_design.php?sequence=NSTPGSSRGTSPARM&method=hybrid&model=main) | MERCI | NEGATIVE | 2 |
| 81 | 196-211 | [STPGSSRGTSPARMA](http://crdd.osdd.net/raghava/ifnepitope/pep_design.php?sequence=STPGSSRGTSPARMA&method=hybrid&model=main) | MERCI | NEGATIVE | 1 |
| 82 | 197-212 | [TPGSSRGTSPARMAG](http://crdd.osdd.net/raghava/ifnepitope/pep_design.php?sequence=TPGSSRGTSPARMAG&method=hybrid&model=main) | MERCI | NEGATIVE | 1 |
| 83 | 198-213 | [PGSSRGTSPARMAGN](http://crdd.osdd.net/raghava/ifnepitope/pep_design.php?sequence=PGSSRGTSPARMAGN&method=hybrid&model=main) | MERCI | NEGATIVE | 1 |
| 84 | 200-215 | [SSRGTSPARMAGNGG](http://crdd.osdd.net/raghava/ifnepitope/pep_design.php?sequence=SSRGTSPARMAGNGG&method=hybrid&model=main) | MERCI | NEGATIVE | 1 |
| 85 | 201-216 | [SRGTSPARMAGNGGD](http://crdd.osdd.net/raghava/ifnepitope/pep_design.php?sequence=SRGTSPARMAGNGGD&method=hybrid&model=main) | MERCI | NEGATIVE | 2 |
| 86 | 202-217 | [RGTSPARMAGNGGDA](http://crdd.osdd.net/raghava/ifnepitope/pep_design.php?sequence=RGTSPARMAGNGGDA&method=hybrid&model=main) | MERCI | NEGATIVE | 2 |
| 87 | 203-218 | [GTSPARMAGNGGDAA](http://crdd.osdd.net/raghava/ifnepitope/pep_design.php?sequence=GTSPARMAGNGGDAA&method=hybrid&model=main) | MERCI | NEGATIVE | 2 |
| 88 | 204-219 | [TSPARMAGNGGDAAL](http://crdd.osdd.net/raghava/ifnepitope/pep_design.php?sequence=TSPARMAGNGGDAAL&method=hybrid&model=main) | MERCI | NEGATIVE | 2 |
| 89 | 205-220 | [SPARMAGNGGDAALA](http://crdd.osdd.net/raghava/ifnepitope/pep_design.php?sequence=SPARMAGNGGDAALA&method=hybrid&model=main) | MERCI | NEGATIVE | 2 |
| 90 | 206-221 | [PARMAGNGGDAALAL](http://crdd.osdd.net/raghava/ifnepitope/pep_design.php?sequence=PARMAGNGGDAALAL&method=hybrid&model=main) | MERCI | NEGATIVE | 2 |
| 91 | 209-224 | [MAGNGGDAALALLLL](http://crdd.osdd.net/raghava/ifnepitope/pep_design.php?sequence=MAGNGGDAALALLLL&method=hybrid&model=main) | MERCI | NEGATIVE | 3 |
| 92 | 210-225 | [AGNGGDAALALLLLD](http://crdd.osdd.net/raghava/ifnepitope/pep_design.php?sequence=AGNGGDAALALLLLD&method=hybrid&model=main) | MERCI | NEGATIVE | 3 |
| 93 | 211-226 | [GNGGDAALALLLLDR](http://crdd.osdd.net/raghava/ifnepitope/pep_design.php?sequence=GNGGDAALALLLLDR&method=hybrid&model=main) | MERCI | NEGATIVE | 3 |
| 94 | 212-227 | [NGGDAALALLLLDRL](http://crdd.osdd.net/raghava/ifnepitope/pep_design.php?sequence=NGGDAALALLLLDRL&method=hybrid&model=main) | MERCI | NEGATIVE | 4 |
| 95 | 213-228 | [GGDAALALLLLDRLN](http://crdd.osdd.net/raghava/ifnepitope/pep_design.php?sequence=GGDAALALLLLDRLN&method=hybrid&model=main) | MERCI | NEGATIVE | 4 |
| 96 | 214-229 | [GDAALALLLLDRLNQ](http://crdd.osdd.net/raghava/ifnepitope/pep_design.php?sequence=GDAALALLLLDRLNQ&method=hybrid&model=main) | MERCI | NEGATIVE | 4 |
| 97 | 215-230 | [DAALALLLLDRLNQL](http://crdd.osdd.net/raghava/ifnepitope/pep_design.php?sequence=DAALALLLLDRLNQL&method=hybrid&model=main) | MERCI | NEGATIVE | 4 |
| 98 | 216-231 | [AALALLLLDRLNQLE](http://crdd.osdd.net/raghava/ifnepitope/pep_design.php?sequence=AALALLLLDRLNQLE&method=hybrid&model=main) | MERCI | NEGATIVE | 3 |
| 99 | 217-232 | [ALALLLLDRLNQLES](http://crdd.osdd.net/raghava/ifnepitope/pep_design.php?sequence=ALALLLLDRLNQLES&method=hybrid&model=main) | MERCI | NEGATIVE | 2 |
| 100 | 218-233 | [LALLLLDRLNQLESK](http://crdd.osdd.net/raghava/ifnepitope/pep_design.php?sequence=LALLLLDRLNQLESK&method=hybrid&model=main) | MERCI | NEGATIVE | 2 |
| 101 | 219-234 | [ALLLLDRLNQLESKM](http://crdd.osdd.net/raghava/ifnepitope/pep_design.php?sequence=ALLLLDRLNQLESKM&method=hybrid&model=main) | MERCI | NEGATIVE | 1 |
| 102 | 220-235 | [LLLLDRLNQLESKMS](http://crdd.osdd.net/raghava/ifnepitope/pep_design.php?sequence=LLLLDRLNQLESKMS&method=hybrid&model=main) | MERCI | NEGATIVE | 1 |
| 103 | 326-341 | [SGTWLTYTGAIKLDD](http://crdd.osdd.net/raghava/ifnepitope/pep_design.php?sequence=SGTWLTYTGAIKLDD&method=hybrid&model=main) | MERCI | NEGATIVE | 1 |
| 104 | 327-342 | [GTWLTYTGAIKLDDK](http://crdd.osdd.net/raghava/ifnepitope/pep_design.php?sequence=GTWLTYTGAIKLDDK&method=hybrid&model=main) | MERCI | NEGATIVE | 1 |
| 105 | 328-343 | [TWLTYTGAIKLDDKD](http://crdd.osdd.net/raghava/ifnepitope/pep_design.php?sequence=TWLTYTGAIKLDDKD&method=hybrid&model=main) | MERCI | NEGATIVE | 1 |
| 106 | 329-344 | [WLTYTGAIKLDDKDP](http://crdd.osdd.net/raghava/ifnepitope/pep_design.php?sequence=WLTYTGAIKLDDKDP&method=hybrid&model=main) | MERCI | NEGATIVE | 1 |
| 107 | 330-345 | [LTYTGAIKLDDKDPN](http://crdd.osdd.net/raghava/ifnepitope/pep_design.php?sequence=LTYTGAIKLDDKDPN&method=hybrid&model=main) | MERCI | NEGATIVE | 1 |
| 108 | 331-346 | [TYTGAIKLDDKDPNF](http://crdd.osdd.net/raghava/ifnepitope/pep_design.php?sequence=TYTGAIKLDDKDPNF&method=hybrid&model=main) | MERCI | NEGATIVE | 1 |
| 109 | 332-347 | [YTGAIKLDDKDPNFK](http://crdd.osdd.net/raghava/ifnepitope/pep_design.php?sequence=YTGAIKLDDKDPNFK&method=hybrid&model=main) | MERCI | NEGATIVE | 1 |
| 110 | 333-348 | [TGAIKLDDKDPNFKD](http://crdd.osdd.net/raghava/ifnepitope/pep_design.php?sequence=TGAIKLDDKDPNFKD&method=hybrid&model=main) | MERCI | NEGATIVE | 1 |
| 111 | 339-354 | [DDKDPNFKDQVILLN](http://crdd.osdd.net/raghava/ifnepitope/pep_design.php?sequence=DDKDPNFKDQVILLN&method=hybrid&model=main) | MERCI | NEGATIVE | 1 |
| 112 | 340-355 | [DKDPNFKDQVILLNK](http://crdd.osdd.net/raghava/ifnepitope/pep_design.php?sequence=DKDPNFKDQVILLNK&method=hybrid&model=main) | MERCI | NEGATIVE | 1 |
| 113 | 341-356 | [KDPNFKDQVILLNKH](http://crdd.osdd.net/raghava/ifnepitope/pep_design.php?sequence=KDPNFKDQVILLNKH&method=hybrid&model=main) | MERCI | NEGATIVE | 1 |
| 114 | 342-357 | [DPNFKDQVILLNKHI](http://crdd.osdd.net/raghava/ifnepitope/pep_design.php?sequence=DPNFKDQVILLNKHI&method=hybrid&model=main) | MERCI | NEGATIVE | 2 |
| 115 | 343-358 | [PNFKDQVILLNKHID](http://crdd.osdd.net/raghava/ifnepitope/pep_design.php?sequence=PNFKDQVILLNKHID&method=hybrid&model=main) | MERCI | NEGATIVE | 2 |
| 116 | 344-359 | [NFKDQVILLNKHIDA](http://crdd.osdd.net/raghava/ifnepitope/pep_design.php?sequence=NFKDQVILLNKHIDA&method=hybrid&model=main) | MERCI | NEGATIVE | 2 |
| 117 | 345-360 | [FKDQVILLNKHIDAY](http://crdd.osdd.net/raghava/ifnepitope/pep_design.php?sequence=FKDQVILLNKHIDAY&method=hybrid&model=main) | MERCI | NEGATIVE | 2 |
| 118 | 346-361 | [KDQVILLNKHIDAYK](http://crdd.osdd.net/raghava/ifnepitope/pep_design.php?sequence=KDQVILLNKHIDAYK&method=hybrid&model=main) | MERCI | NEGATIVE | 1 |
| 119 | 347-362 | [DQVILLNKHIDAYKT](http://crdd.osdd.net/raghava/ifnepitope/pep_design.php?sequence=DQVILLNKHIDAYKT&method=hybrid&model=main) | MERCI | NEGATIVE | 1 |
| 120 | 408-419 | [QSMSSADSTQA](http://crdd.osdd.net/raghava/ifnepitope/pep_design.php?sequence=QSMSSADSTQA&method=hybrid&model=main) | MERCI | NEGATIVE | 3 |
| 121 | 409-419 | [SMSSADSTQA](http://crdd.osdd.net/raghava/ifnepitope/pep_design.php?sequence=SMSSADSTQA&method=hybrid&model=main) | MERCI | NEGATIVE | 3 |
| 122 | 410-419 | [MSSADSTQA](http://crdd.osdd.net/raghava/ifnepitope/pep_design.php?sequence=MSSADSTQA&method=hybrid&model=main) | MERCI | NEGATIVE | 2 |
| 123 | 0-15 | [MSDNGPQNQRNAPRI](http://crdd.osdd.net/raghava/ifnepitope/pep_design.php?sequence=MSDNGPQNQRNAPRI&method=hybrid&model=main) | SVM | NEGATIVE | -0.37994588 |
| 124 | 1-16 | [SDNGPQNQRNAPRIT](http://crdd.osdd.net/raghava/ifnepitope/pep_design.php?sequence=SDNGPQNQRNAPRIT&method=hybrid&model=main) | SVM | NEGATIVE | -0.21717551 |
| 125 | 2-17 | [DNGPQNQRNAPRITF](http://crdd.osdd.net/raghava/ifnepitope/pep_design.php?sequence=DNGPQNQRNAPRITF&method=hybrid&model=main) | SVM | NEGATIVE | -0.46857518 |
| 126 | 3-18 | [NGPQNQRNAPRITFG](http://crdd.osdd.net/raghava/ifnepitope/pep_design.php?sequence=NGPQNQRNAPRITFG&method=hybrid&model=main) | SVM | NEGATIVE | -0.79499459 |
| 127 | 4-19 | [GPQNQRNAPRITFGG](http://crdd.osdd.net/raghava/ifnepitope/pep_design.php?sequence=GPQNQRNAPRITFGG&method=hybrid&model=main) | SVM | NEGATIVE | -0.78344502 |
| 128 | 6-21 | [QNQRNAPRITFGGPS](http://crdd.osdd.net/raghava/ifnepitope/pep_design.php?sequence=QNQRNAPRITFGGPS&method=hybrid&model=main) | SVM | NEGATIVE | -0.63050395 |
| 129 | 7-22 | [NQRNAPRITFGGPSD](http://crdd.osdd.net/raghava/ifnepitope/pep_design.php?sequence=NQRNAPRITFGGPSD&method=hybrid&model=main) | SVM | NEGATIVE | -0.55131968 |
| 130 | 9-24 | [RNAPRITFGGPSDST](http://crdd.osdd.net/raghava/ifnepitope/pep_design.php?sequence=RNAPRITFGGPSDST&method=hybrid&model=main) | SVM | NEGATIVE | -0.52683343 |
| 131 | 10-25 | [NAPRITFGGPSDSTG](http://crdd.osdd.net/raghava/ifnepitope/pep_design.php?sequence=NAPRITFGGPSDSTG&method=hybrid&model=main) | SVM | NEGATIVE | -0.62961772 |
| 132 | 11-26 | [APRITFGGPSDSTGS](http://crdd.osdd.net/raghava/ifnepitope/pep_design.php?sequence=APRITFGGPSDSTGS&method=hybrid&model=main) | SVM | NEGATIVE | -0.64104749 |
| 133 | 12-27 | [PRITFGGPSDSTGSN](http://crdd.osdd.net/raghava/ifnepitope/pep_design.php?sequence=PRITFGGPSDSTGSN&method=hybrid&model=main) | SVM | NEGATIVE | -0.56189558 |
| 134 | 13-28 | [RITFGGPSDSTGSNQ](http://crdd.osdd.net/raghava/ifnepitope/pep_design.php?sequence=RITFGGPSDSTGSNQ&method=hybrid&model=main) | SVM | NEGATIVE | -0.81929342 |
| 135 | 14-29 | [ITFGGPSDSTGSNQN](http://crdd.osdd.net/raghava/ifnepitope/pep_design.php?sequence=ITFGGPSDSTGSNQN&method=hybrid&model=main) | SVM | NEGATIVE | -0.81835869 |
| 136 | 21-36 | [DSTGSNQNGERSGAR](http://crdd.osdd.net/raghava/ifnepitope/pep_design.php?sequence=DSTGSNQNGERSGAR&method=hybrid&model=main) | SVM | NEGATIVE | -0.54180277 |
| 137 | 22-37 | [STGSNQNGERSGARS](http://crdd.osdd.net/raghava/ifnepitope/pep_design.php?sequence=STGSNQNGERSGARS&method=hybrid&model=main) | SVM | NEGATIVE | -0.53136111 |
| 138 | 23-38 | [TGSNQNGERSGARSK](http://crdd.osdd.net/raghava/ifnepitope/pep_design.php?sequence=TGSNQNGERSGARSK&method=hybrid&model=main) | SVM | NEGATIVE | -0.75193546 |
| 139 | 24-39 | [GSNQNGERSGARSKQ](http://crdd.osdd.net/raghava/ifnepitope/pep_design.php?sequence=GSNQNGERSGARSKQ&method=hybrid&model=main) | SVM | NEGATIVE | -0.88059957 |
| 140 | 25-40 | [SNQNGERSGARSKQR](http://crdd.osdd.net/raghava/ifnepitope/pep_design.php?sequence=SNQNGERSGARSKQR&method=hybrid&model=main) | SVM | NEGATIVE | -0.50373778 |
| 141 | 26-41 | [NQNGERSGARSKQRR](http://crdd.osdd.net/raghava/ifnepitope/pep_design.php?sequence=NQNGERSGARSKQRR&method=hybrid&model=main) | SVM | NEGATIVE | -0.49902873 |
| 142 | 27-42 | [QNGERSGARSKQRRP](http://crdd.osdd.net/raghava/ifnepitope/pep_design.php?sequence=QNGERSGARSKQRRP&method=hybrid&model=main) | SVM | NEGATIVE | -0.080823456 |
| 143 | 28-43 | [NGERSGARSKQRRPQ](http://crdd.osdd.net/raghava/ifnepitope/pep_design.php?sequence=NGERSGARSKQRRPQ&method=hybrid&model=main) | SVM | NEGATIVE | -0.50778428 |
| 144 | 29-44 | [GERSGARSKQRRPQG](http://crdd.osdd.net/raghava/ifnepitope/pep_design.php?sequence=GERSGARSKQRRPQG&method=hybrid&model=main) | SVM | NEGATIVE | -0.32171563 |
| 145 | 30-45 | [ERSGARSKQRRPQGL](http://crdd.osdd.net/raghava/ifnepitope/pep_design.php?sequence=ERSGARSKQRRPQGL&method=hybrid&model=main) | SVM | NEGATIVE | -0.020005202 |
| 146 | 31-46 | [RSGARSKQRRPQGLP](http://crdd.osdd.net/raghava/ifnepitope/pep_design.php?sequence=RSGARSKQRRPQGLP&method=hybrid&model=main) | SVM | POSITIVE | 0.072258402 |
| 147 | 32-47 | [SGARSKQRRPQGLPN](http://crdd.osdd.net/raghava/ifnepitope/pep_design.php?sequence=SGARSKQRRPQGLPN&method=hybrid&model=main) | SVM | POSITIVE | 0.050659937 |
| 148 | 33-48 | [GARSKQRRPQGLPNN](http://crdd.osdd.net/raghava/ifnepitope/pep_design.php?sequence=GARSKQRRPQGLPNN&method=hybrid&model=main) | SVM | POSITIVE | 0.10342223 |
| 149 | 34-49 | [ARSKQRRPQGLPNNT](http://crdd.osdd.net/raghava/ifnepitope/pep_design.php?sequence=ARSKQRRPQGLPNNT&method=hybrid&model=main) | SVM | NEGATIVE | -0.016681311 |
| 150 | 35-50 | [RSKQRRPQGLPNNTA](http://crdd.osdd.net/raghava/ifnepitope/pep_design.php?sequence=RSKQRRPQGLPNNTA&method=hybrid&model=main) | SVM | NEGATIVE | -0.055722297 |
| 151 | 49-64 | [ASWFTALTQHGKEDL](http://crdd.osdd.net/raghava/ifnepitope/pep_design.php?sequence=ASWFTALTQHGKEDL&method=hybrid&model=main) | SVM | NEGATIVE | -0.50009616 |
| 152 | 50-65 | [SWFTALTQHGKEDLK](http://crdd.osdd.net/raghava/ifnepitope/pep_design.php?sequence=SWFTALTQHGKEDLK&method=hybrid&model=main) | SVM | NEGATIVE | -0.43260126 |
| 153 | 51-66 | [WFTALTQHGKEDLKF](http://crdd.osdd.net/raghava/ifnepitope/pep_design.php?sequence=WFTALTQHGKEDLKF&method=hybrid&model=main) | SVM | NEGATIVE | -0.43474643 |
| 154 | 52-67 | [FTALTQHGKEDLKFP](http://crdd.osdd.net/raghava/ifnepitope/pep_design.php?sequence=FTALTQHGKEDLKFP&method=hybrid&model=main) | SVM | NEGATIVE | -0.47468428 |
| 155 | 53-68 | [TALTQHGKEDLKFPR](http://crdd.osdd.net/raghava/ifnepitope/pep_design.php?sequence=TALTQHGKEDLKFPR&method=hybrid&model=main) | SVM | NEGATIVE | -0.33286068 |
| 156 | 54-69 | [ALTQHGKEDLKFPRG](http://crdd.osdd.net/raghava/ifnepitope/pep_design.php?sequence=ALTQHGKEDLKFPRG&method=hybrid&model=main) | SVM | NEGATIVE | -0.25907233 |
| 157 | 55-70 | [LTQHGKEDLKFPRGQ](http://crdd.osdd.net/raghava/ifnepitope/pep_design.php?sequence=LTQHGKEDLKFPRGQ&method=hybrid&model=main) | SVM | NEGATIVE | -0.33344812 |
| 158 | 56-71 | [TQHGKEDLKFPRGQG](http://crdd.osdd.net/raghava/ifnepitope/pep_design.php?sequence=TQHGKEDLKFPRGQG&method=hybrid&model=main) | SVM | NEGATIVE | -0.46896914 |
| 159 | 57-72 | [QHGKEDLKFPRGQGV](http://crdd.osdd.net/raghava/ifnepitope/pep_design.php?sequence=QHGKEDLKFPRGQGV&method=hybrid&model=main) | SVM | NEGATIVE | -0.36969304 |
| 160 | 58-73 | [HGKEDLKFPRGQGVP](http://crdd.osdd.net/raghava/ifnepitope/pep_design.php?sequence=HGKEDLKFPRGQGVP&method=hybrid&model=main) | SVM | NEGATIVE | -0.53312389 |
| 161 | 59-74 | [GKEDLKFPRGQGVPI](http://crdd.osdd.net/raghava/ifnepitope/pep_design.php?sequence=GKEDLKFPRGQGVPI&method=hybrid&model=main) | SVM | NEGATIVE | -0.35608418 |
| 162 | 60-75 | [KEDLKFPRGQGVPIN](http://crdd.osdd.net/raghava/ifnepitope/pep_design.php?sequence=KEDLKFPRGQGVPIN&method=hybrid&model=main) | SVM | NEGATIVE | -0.62190466 |
| 163 | 61-76 | [EDLKFPRGQGVPINT](http://crdd.osdd.net/raghava/ifnepitope/pep_design.php?sequence=EDLKFPRGQGVPINT&method=hybrid&model=main) | SVM | NEGATIVE | -0.48091014 |
| 164 | 62-77 | [DLKFPRGQGVPINTN](http://crdd.osdd.net/raghava/ifnepitope/pep_design.php?sequence=DLKFPRGQGVPINTN&method=hybrid&model=main) | SVM | NEGATIVE | -0.33116071 |
| 165 | 63-78 | [LKFPRGQGVPINTNS](http://crdd.osdd.net/raghava/ifnepitope/pep_design.php?sequence=LKFPRGQGVPINTNS&method=hybrid&model=main) | SVM | NEGATIVE | -0.12039267 |
| 166 | 64-79 | [KFPRGQGVPINTNSS](http://crdd.osdd.net/raghava/ifnepitope/pep_design.php?sequence=KFPRGQGVPINTNSS&method=hybrid&model=main) | SVM | NEGATIVE | -0.21307067 |
| 167 | 65-80 | [FPRGQGVPINTNSSP](http://crdd.osdd.net/raghava/ifnepitope/pep_design.php?sequence=FPRGQGVPINTNSSP&method=hybrid&model=main) | SVM | NEGATIVE | -0.43891798 |
| 168 | 66-81 | [PRGQGVPINTNSSPD](http://crdd.osdd.net/raghava/ifnepitope/pep_design.php?sequence=PRGQGVPINTNSSPD&method=hybrid&model=main) | SVM | NEGATIVE | -0.49575571 |
| 169 | 67-82 | [RGQGVPINTNSSPDD](http://crdd.osdd.net/raghava/ifnepitope/pep_design.php?sequence=RGQGVPINTNSSPDD&method=hybrid&model=main) | SVM | NEGATIVE | -0.60834661 |
| 170 | 70-85 | [GVPINTNSSPDDQIG](http://crdd.osdd.net/raghava/ifnepitope/pep_design.php?sequence=GVPINTNSSPDDQIG&method=hybrid&model=main) | SVM | NEGATIVE | -0.85105303 |
| 171 | 71-86 | [VPINTNSSPDDQIGY](http://crdd.osdd.net/raghava/ifnepitope/pep_design.php?sequence=VPINTNSSPDDQIGY&method=hybrid&model=main) | SVM | NEGATIVE | -0.77886212 |
| 172 | 72-87 | [PINTNSSPDDQIGYY](http://crdd.osdd.net/raghava/ifnepitope/pep_design.php?sequence=PINTNSSPDDQIGYY&method=hybrid&model=main) | SVM | NEGATIVE | -0.58628818 |
| 173 | 73-88 | [INTNSSPDDQIGYYR](http://crdd.osdd.net/raghava/ifnepitope/pep_design.php?sequence=INTNSSPDDQIGYYR&method=hybrid&model=main) | SVM | NEGATIVE | -0.61893959 |
| 174 | 74-89 | [NTNSSPDDQIGYYRR](http://crdd.osdd.net/raghava/ifnepitope/pep_design.php?sequence=NTNSSPDDQIGYYRR&method=hybrid&model=main) | SVM | NEGATIVE | -0.59304811 |
| 175 | 75-90 | [TNSSPDDQIGYYRRA](http://crdd.osdd.net/raghava/ifnepitope/pep_design.php?sequence=TNSSPDDQIGYYRRA&method=hybrid&model=main) | SVM | NEGATIVE | -0.22791555 |
| 176 | 76-91 | [NSSPDDQIGYYRRAT](http://crdd.osdd.net/raghava/ifnepitope/pep_design.php?sequence=NSSPDDQIGYYRRAT&method=hybrid&model=main) | SVM | NEGATIVE | -0.096505833 |
| 177 | 77-92 | [SSPDDQIGYYRRATR](http://crdd.osdd.net/raghava/ifnepitope/pep_design.php?sequence=SSPDDQIGYYRRATR&method=hybrid&model=main) | SVM | POSITIVE | 0.16646905 |
| 178 | 78-93 | [SPDDQIGYYRRATRR](http://crdd.osdd.net/raghava/ifnepitope/pep_design.php?sequence=SPDDQIGYYRRATRR&method=hybrid&model=main) | SVM | POSITIVE | 0.58541752 |
| 179 | 79-94 | [PDDQIGYYRRATRRI](http://crdd.osdd.net/raghava/ifnepitope/pep_design.php?sequence=PDDQIGYYRRATRRI&method=hybrid&model=main) | SVM | POSITIVE | 0.77042851 |
| 180 | 80-95 | [DDQIGYYRRATRRIR](http://crdd.osdd.net/raghava/ifnepitope/pep_design.php?sequence=DDQIGYYRRATRRIR&method=hybrid&model=main) | SVM | POSITIVE | 0.66602777 |
| 181 | 81-96 | [DQIGYYRRATRRIRG](http://crdd.osdd.net/raghava/ifnepitope/pep_design.php?sequence=DQIGYYRRATRRIRG&method=hybrid&model=main) | SVM | POSITIVE | 0.77580404 |
| 182 | 82-97 | [QIGYYRRATRRIRGG](http://crdd.osdd.net/raghava/ifnepitope/pep_design.php?sequence=QIGYYRRATRRIRGG&method=hybrid&model=main) | SVM | POSITIVE | 0.55082319 |
| 183 | 83-98 | [IGYYRRATRRIRGGD](http://crdd.osdd.net/raghava/ifnepitope/pep_design.php?sequence=IGYYRRATRRIRGGD&method=hybrid&model=main) | SVM | POSITIVE | 0.52618964 |
| 184 | 84-99 | [GYYRRATRRIRGGDG](http://crdd.osdd.net/raghava/ifnepitope/pep_design.php?sequence=GYYRRATRRIRGGDG&method=hybrid&model=main) | SVM | POSITIVE | 0.319095 |
| 185 | 85-100 | [YYRRATRRIRGGDGK](http://crdd.osdd.net/raghava/ifnepitope/pep_design.php?sequence=YYRRATRRIRGGDGK&method=hybrid&model=main) | SVM | POSITIVE | 0.44190447 |
| 186 | 86-101 | [YRRATRRIRGGDGKM](http://crdd.osdd.net/raghava/ifnepitope/pep_design.php?sequence=YRRATRRIRGGDGKM&method=hybrid&model=main) | SVM | POSITIVE | 0.2967349 |
| 187 | 87-102 | [RRATRRIRGGDGKMK](http://crdd.osdd.net/raghava/ifnepitope/pep_design.php?sequence=RRATRRIRGGDGKMK&method=hybrid&model=main) | SVM | POSITIVE | 0.44086179 |
| 188 | 88-103 | [RATRRIRGGDGKMKD](http://crdd.osdd.net/raghava/ifnepitope/pep_design.php?sequence=RATRRIRGGDGKMKD&method=hybrid&model=main) | SVM | POSITIVE | 0.089196847 |
| 189 | 89-104 | [ATRRIRGGDGKMKDL](http://crdd.osdd.net/raghava/ifnepitope/pep_design.php?sequence=ATRRIRGGDGKMKDL&method=hybrid&model=main) | SVM | NEGATIVE | -0.12786695 |
| 190 | 95-110 | [GGDGKMKDLSPRWYF](http://crdd.osdd.net/raghava/ifnepitope/pep_design.php?sequence=GGDGKMKDLSPRWYF&method=hybrid&model=main) | SVM | NEGATIVE | -0.22085802 |
| 191 | 96-111 | [GDGKMKDLSPRWYFY](http://crdd.osdd.net/raghava/ifnepitope/pep_design.php?sequence=GDGKMKDLSPRWYFY&method=hybrid&model=main) | SVM | NEGATIVE | -0.25016199 |
| 192 | 97-112 | [DGKMKDLSPRWYFYY](http://crdd.osdd.net/raghava/ifnepitope/pep_design.php?sequence=DGKMKDLSPRWYFYY&method=hybrid&model=main) | SVM | POSITIVE | 0.13520678 |
| 193 | 98-113 | [GKMKDLSPRWYFYYL](http://crdd.osdd.net/raghava/ifnepitope/pep_design.php?sequence=GKMKDLSPRWYFYYL&method=hybrid&model=main) | SVM | POSITIVE | 0.45987756 |
| 194 | 99-114 | [KMKDLSPRWYFYYLG](http://crdd.osdd.net/raghava/ifnepitope/pep_design.php?sequence=KMKDLSPRWYFYYLG&method=hybrid&model=main) | SVM | POSITIVE | 0.39401059 |
| 195 | 100-115 | [MKDLSPRWYFYYLGT](http://crdd.osdd.net/raghava/ifnepitope/pep_design.php?sequence=MKDLSPRWYFYYLGT&method=hybrid&model=main) | SVM | POSITIVE | 0.078021587 |
| 196 | 101-116 | [KDLSPRWYFYYLGTG](http://crdd.osdd.net/raghava/ifnepitope/pep_design.php?sequence=KDLSPRWYFYYLGTG&method=hybrid&model=main) | SVM | NEGATIVE | -0.30296092 |
| 197 | 102-117 | [DLSPRWYFYYLGTGP](http://crdd.osdd.net/raghava/ifnepitope/pep_design.php?sequence=DLSPRWYFYYLGTGP&method=hybrid&model=main) | SVM | NEGATIVE | -0.70723389 |
| 198 | 103-118 | [LSPRWYFYYLGTGPE](http://crdd.osdd.net/raghava/ifnepitope/pep_design.php?sequence=LSPRWYFYYLGTGPE&method=hybrid&model=main) | SVM | NEGATIVE | -0.74612286 |
| 199 | 104-119 | [SPRWYFYYLGTGPEA](http://crdd.osdd.net/raghava/ifnepitope/pep_design.php?sequence=SPRWYFYYLGTGPEA&method=hybrid&model=main) | SVM | NEGATIVE | -0.62866172 |
| 200 | 105-120 | [PRWYFYYLGTGPEAG](http://crdd.osdd.net/raghava/ifnepitope/pep_design.php?sequence=PRWYFYYLGTGPEAG&method=hybrid&model=main) | SVM | NEGATIVE | -0.57413373 |
| 201 | 106-121 | [RWYFYYLGTGPEAGL](http://crdd.osdd.net/raghava/ifnepitope/pep_design.php?sequence=RWYFYYLGTGPEAGL&method=hybrid&model=main) | SVM | NEGATIVE | -0.39095777 |
| 202 | 107-122 | [WYFYYLGTGPEAGLP](http://crdd.osdd.net/raghava/ifnepitope/pep_design.php?sequence=WYFYYLGTGPEAGLP&method=hybrid&model=main) | SVM | NEGATIVE | -0.77713009 |
| 203 | 108-123 | [YFYYLGTGPEAGLPY](http://crdd.osdd.net/raghava/ifnepitope/pep_design.php?sequence=YFYYLGTGPEAGLPY&method=hybrid&model=main) | SVM | NEGATIVE | -1.1450368 |
| 204 | 109-124 | [FYYLGTGPEAGLPYG](http://crdd.osdd.net/raghava/ifnepitope/pep_design.php?sequence=FYYLGTGPEAGLPYG&method=hybrid&model=main) | SVM | NEGATIVE | -1.0742872 |
| 205 | 110-125 | [YYLGTGPEAGLPYGA](http://crdd.osdd.net/raghava/ifnepitope/pep_design.php?sequence=YYLGTGPEAGLPYGA&method=hybrid&model=main) | SVM | NEGATIVE | -0.54446841 |
| 206 | 111-126 | [YLGTGPEAGLPYGAN](http://crdd.osdd.net/raghava/ifnepitope/pep_design.php?sequence=YLGTGPEAGLPYGAN&method=hybrid&model=main) | SVM | NEGATIVE | -0.491731 |
| 207 | 112-127 | [LGTGPEAGLPYGANK](http://crdd.osdd.net/raghava/ifnepitope/pep_design.php?sequence=LGTGPEAGLPYGANK&method=hybrid&model=main) | SVM | NEGATIVE | -0.61884004 |
| 208 | 113-128 | [GTGPEAGLPYGANKD](http://crdd.osdd.net/raghava/ifnepitope/pep_design.php?sequence=GTGPEAGLPYGANKD&method=hybrid&model=main) | SVM | NEGATIVE | -0.57904035 |
| 209 | 114-129 | [TGPEAGLPYGANKDG](http://crdd.osdd.net/raghava/ifnepitope/pep_design.php?sequence=TGPEAGLPYGANKDG&method=hybrid&model=main) | SVM | NEGATIVE | -0.57302473 |
| 210 | 115-130 | [GPEAGLPYGANKDGI](http://crdd.osdd.net/raghava/ifnepitope/pep_design.php?sequence=GPEAGLPYGANKDGI&method=hybrid&model=main) | SVM | NEGATIVE | -0.52276027 |
| 211 | 116-131 | [PEAGLPYGANKDGII](http://crdd.osdd.net/raghava/ifnepitope/pep_design.php?sequence=PEAGLPYGANKDGII&method=hybrid&model=main) | SVM | NEGATIVE | -0.21845712 |
| 212 | 117-132 | [EAGLPYGANKDGIIW](http://crdd.osdd.net/raghava/ifnepitope/pep_design.php?sequence=EAGLPYGANKDGIIW&method=hybrid&model=main) | SVM | NEGATIVE | -0.0091701176 |
| 213 | 118-133 | [AGLPYGANKDGIIWV](http://crdd.osdd.net/raghava/ifnepitope/pep_design.php?sequence=AGLPYGANKDGIIWV&method=hybrid&model=main) | SVM | NEGATIVE | -0.021112347 |
| 214 | 130-145 | [IWVATEGALNTPKDH](http://crdd.osdd.net/raghava/ifnepitope/pep_design.php?sequence=IWVATEGALNTPKDH&method=hybrid&model=main) | SVM | NEGATIVE | -0.057436399 |
| 215 | 131-146 | [WVATEGALNTPKDHI](http://crdd.osdd.net/raghava/ifnepitope/pep_design.php?sequence=WVATEGALNTPKDHI&method=hybrid&model=main) | SVM | NEGATIVE | -0.16868551 |
| 216 | 132-147 | [VATEGALNTPKDHIG](http://crdd.osdd.net/raghava/ifnepitope/pep_design.php?sequence=VATEGALNTPKDHIG&method=hybrid&model=main) | SVM | NEGATIVE | -0.42145299 |
| 217 | 133-148 | [ATEGALNTPKDHIGT](http://crdd.osdd.net/raghava/ifnepitope/pep_design.php?sequence=ATEGALNTPKDHIGT&method=hybrid&model=main) | SVM | NEGATIVE | -0.50094912 |
| 218 | 134-149 | [TEGALNTPKDHIGTR](http://crdd.osdd.net/raghava/ifnepitope/pep_design.php?sequence=TEGALNTPKDHIGTR&method=hybrid&model=main) | SVM | NEGATIVE | -0.3531847 |
| 219 | 135-150 | [EGALNTPKDHIGTRN](http://crdd.osdd.net/raghava/ifnepitope/pep_design.php?sequence=EGALNTPKDHIGTRN&method=hybrid&model=main) | SVM | NEGATIVE | -0.45960819 |
| 220 | 136-151 | [GALNTPKDHIGTRNP](http://crdd.osdd.net/raghava/ifnepitope/pep_design.php?sequence=GALNTPKDHIGTRNP&method=hybrid&model=main) | SVM | NEGATIVE | -0.66299378 |
| 221 | 137-152 | [ALNTPKDHIGTRNPA](http://crdd.osdd.net/raghava/ifnepitope/pep_design.php?sequence=ALNTPKDHIGTRNPA&method=hybrid&model=main) | SVM | NEGATIVE | -0.56269751 |
| 222 | 138-153 | [LNTPKDHIGTRNPAN](http://crdd.osdd.net/raghava/ifnepitope/pep_design.php?sequence=LNTPKDHIGTRNPAN&method=hybrid&model=main) | SVM | NEGATIVE | -0.63069528 |
| 223 | 139-154 | [NTPKDHIGTRNPANN](http://crdd.osdd.net/raghava/ifnepitope/pep_design.php?sequence=NTPKDHIGTRNPANN&method=hybrid&model=main) | SVM | NEGATIVE | -0.59655895 |
| 224 | 140-155 | [TPKDHIGTRNPANNA](http://crdd.osdd.net/raghava/ifnepitope/pep_design.php?sequence=TPKDHIGTRNPANNA&method=hybrid&model=main) | SVM | NEGATIVE | -0.53052231 |
| 225 | 141-156 | [PKDHIGTRNPANNAA](http://crdd.osdd.net/raghava/ifnepitope/pep_design.php?sequence=PKDHIGTRNPANNAA&method=hybrid&model=main) | SVM | NEGATIVE | -0.38317236 |
| 226 | 142-157 | [KDHIGTRNPANNAAI](http://crdd.osdd.net/raghava/ifnepitope/pep_design.php?sequence=KDHIGTRNPANNAAI&method=hybrid&model=main) | SVM | NEGATIVE | -0.41984237 |
| 227 | 143-158 | [DHIGTRNPANNAAIV](http://crdd.osdd.net/raghava/ifnepitope/pep_design.php?sequence=DHIGTRNPANNAAIV&method=hybrid&model=main) | SVM | NEGATIVE | -0.49183112 |
| 228 | 153-168 | [NAAIVLQLPQGTTLP](http://crdd.osdd.net/raghava/ifnepitope/pep_design.php?sequence=NAAIVLQLPQGTTLP&method=hybrid&model=main) | SVM | NEGATIVE | -0.40771118 |
| 229 | 154-169 | [AAIVLQLPQGTTLPK](http://crdd.osdd.net/raghava/ifnepitope/pep_design.php?sequence=AAIVLQLPQGTTLPK&method=hybrid&model=main) | SVM | NEGATIVE | -0.65879926 |
| 230 | 155-170 | [AIVLQLPQGTTLPKG](http://crdd.osdd.net/raghava/ifnepitope/pep_design.php?sequence=AIVLQLPQGTTLPKG&method=hybrid&model=main) | SVM | NEGATIVE | -0.79681089 |
| 231 | 156-171 | [IVLQLPQGTTLPKGF](http://crdd.osdd.net/raghava/ifnepitope/pep_design.php?sequence=IVLQLPQGTTLPKGF&method=hybrid&model=main) | SVM | NEGATIVE | -0.87603568 |
| 232 | 157-172 | [VLQLPQGTTLPKGFY](http://crdd.osdd.net/raghava/ifnepitope/pep_design.php?sequence=VLQLPQGTTLPKGFY&method=hybrid&model=main) | SVM | NEGATIVE | -0.9904743 |
| 233 | 158-173 | [LQLPQGTTLPKGFYA](http://crdd.osdd.net/raghava/ifnepitope/pep_design.php?sequence=LQLPQGTTLPKGFYA&method=hybrid&model=main) | SVM | NEGATIVE | -0.8410899 |
| 234 | 159-174 | [QLPQGTTLPKGFYAE](http://crdd.osdd.net/raghava/ifnepitope/pep_design.php?sequence=QLPQGTTLPKGFYAE&method=hybrid&model=main) | SVM | NEGATIVE | -0.91920858 |
| 235 | 160-175 | [LPQGTTLPKGFYAEG](http://crdd.osdd.net/raghava/ifnepitope/pep_design.php?sequence=LPQGTTLPKGFYAEG&method=hybrid&model=main) | SVM | NEGATIVE | -0.80924099 |
| 236 | 161-176 | [PQGTTLPKGFYAEGS](http://crdd.osdd.net/raghava/ifnepitope/pep_design.php?sequence=PQGTTLPKGFYAEGS&method=hybrid&model=main) | SVM | NEGATIVE | -0.95142865 |
| 237 | 162-177 | [QGTTLPKGFYAEGSR](http://crdd.osdd.net/raghava/ifnepitope/pep_design.php?sequence=QGTTLPKGFYAEGSR&method=hybrid&model=main) | SVM | NEGATIVE | -0.73415706 |
| 238 | 163-178 | [GTTLPKGFYAEGSRG](http://crdd.osdd.net/raghava/ifnepitope/pep_design.php?sequence=GTTLPKGFYAEGSRG&method=hybrid&model=main) | SVM | NEGATIVE | -0.65117748 |
| 239 | 164-179 | [TTLPKGFYAEGSRGG](http://crdd.osdd.net/raghava/ifnepitope/pep_design.php?sequence=TTLPKGFYAEGSRGG&method=hybrid&model=main) | SVM | NEGATIVE | -0.47916366 |
| 240 | 165-180 | [TLPKGFYAEGSRGGS](http://crdd.osdd.net/raghava/ifnepitope/pep_design.php?sequence=TLPKGFYAEGSRGGS&method=hybrid&model=main) | SVM | NEGATIVE | -0.33405755 |
| 241 | 166-181 | [LPKGFYAEGSRGGSQ](http://crdd.osdd.net/raghava/ifnepitope/pep_design.php?sequence=LPKGFYAEGSRGGSQ&method=hybrid&model=main) | SVM | NEGATIVE | -0.52156816 |
| 242 | 167-182 | [PKGFYAEGSRGGSQA](http://crdd.osdd.net/raghava/ifnepitope/pep_design.php?sequence=PKGFYAEGSRGGSQA&method=hybrid&model=main) | SVM | NEGATIVE | -0.18484711 |
| 243 | 168-183 | [KGFYAEGSRGGSQAS](http://crdd.osdd.net/raghava/ifnepitope/pep_design.php?sequence=KGFYAEGSRGGSQAS&method=hybrid&model=main) | SVM | NEGATIVE | -0.14875852 |
| 244 | 181-196 | [ASSRSSSRSRNSSRN](http://crdd.osdd.net/raghava/ifnepitope/pep_design.php?sequence=ASSRSSSRSRNSSRN&method=hybrid&model=main) | SVM | POSITIVE | 0.5604275 |
| 245 | 182-197 | [SSRSSSRSRNSSRNS](http://crdd.osdd.net/raghava/ifnepitope/pep_design.php?sequence=SSRSSSRSRNSSRNS&method=hybrid&model=main) | SVM | POSITIVE | 0.74507971 |
| 246 | 183-198 | [SRSSSRSRNSSRNST](http://crdd.osdd.net/raghava/ifnepitope/pep_design.php?sequence=SRSSSRSRNSSRNST&method=hybrid&model=main) | SVM | POSITIVE | 0.61963204 |
| 247 | 184-199 | [RSSSRSRNSSRNSTP](http://crdd.osdd.net/raghava/ifnepitope/pep_design.php?sequence=RSSSRSRNSSRNSTP&method=hybrid&model=main) | SVM | POSITIVE | 0.46761187 |
| 248 | 185-200 | [SSSRSRNSSRNSTPG](http://crdd.osdd.net/raghava/ifnepitope/pep_design.php?sequence=SSSRSRNSSRNSTPG&method=hybrid&model=main) | SVM | POSITIVE | 0.35247593 |
| 249 | 186-201 | [SSRSRNSSRNSTPGS](http://crdd.osdd.net/raghava/ifnepitope/pep_design.php?sequence=SSRSRNSSRNSTPGS&method=hybrid&model=main) | SVM | POSITIVE | 0.23130688 |
| 250 | 187-202 | [SRSRNSSRNSTPGSS](http://crdd.osdd.net/raghava/ifnepitope/pep_design.php?sequence=SRSRNSSRNSTPGSS&method=hybrid&model=main) | SVM | POSITIVE | 0.23130688 |
| 251 | 188-203 | [RSRNSSRNSTPGSSR](http://crdd.osdd.net/raghava/ifnepitope/pep_design.php?sequence=RSRNSSRNSTPGSSR&method=hybrid&model=main) | SVM | POSITIVE | 0.23130688 |
| 252 | 199-214 | [GSSRGTSPARMAGNG](http://crdd.osdd.net/raghava/ifnepitope/pep_design.php?sequence=GSSRGTSPARMAGNG&method=hybrid&model=main) | SVM | NEGATIVE | -0.21993723 |
| 253 | 207-222 | [ARMAGNGGDAALALL](http://crdd.osdd.net/raghava/ifnepitope/pep_design.php?sequence=ARMAGNGGDAALALL&method=hybrid&model=main) | SVM | NEGATIVE | -0.53658217 |
| 254 | 208-223 | [RMAGNGGDAALALLL](http://crdd.osdd.net/raghava/ifnepitope/pep_design.php?sequence=RMAGNGGDAALALLL&method=hybrid&model=main) | SVM | NEGATIVE | -0.3724267 |
| 255 | 221-236 | [LLLDRLNQLESKMSG](http://crdd.osdd.net/raghava/ifnepitope/pep_design.php?sequence=LLLDRLNQLESKMSG&method=hybrid&model=main) | SVM | NEGATIVE | -0.70765882 |
| 256 | 222-237 | [LLDRLNQLESKMSGK](http://crdd.osdd.net/raghava/ifnepitope/pep_design.php?sequence=LLDRLNQLESKMSGK&method=hybrid&model=main) | SVM | NEGATIVE | -0.71659976 |
| 257 | 223-238 | [LDRLNQLESKMSGKG](http://crdd.osdd.net/raghava/ifnepitope/pep_design.php?sequence=LDRLNQLESKMSGKG&method=hybrid&model=main) | SVM | NEGATIVE | -0.88468176 |
| 258 | 224-239 | [DRLNQLESKMSGKGQ](http://crdd.osdd.net/raghava/ifnepitope/pep_design.php?sequence=DRLNQLESKMSGKGQ&method=hybrid&model=main) | SVM | NEGATIVE | -0.803611 |
| 259 | 225-240 | [RLNQLESKMSGKGQQ](http://crdd.osdd.net/raghava/ifnepitope/pep_design.php?sequence=RLNQLESKMSGKGQQ&method=hybrid&model=main) | SVM | NEGATIVE | -0.73734158 |
| 260 | 226-241 | [LNQLESKMSGKGQQQ](http://crdd.osdd.net/raghava/ifnepitope/pep_design.php?sequence=LNQLESKMSGKGQQQ&method=hybrid&model=main) | SVM | NEGATIVE | -0.53176084 |
| 261 | 227-242 | [NQLESKMSGKGQQQQ](http://crdd.osdd.net/raghava/ifnepitope/pep_design.php?sequence=NQLESKMSGKGQQQQ&method=hybrid&model=main) | SVM | NEGATIVE | -0.18801115 |
| 262 | 228-243 | [QLESKMSGKGQQQQG](http://crdd.osdd.net/raghava/ifnepitope/pep_design.php?sequence=QLESKMSGKGQQQQG&method=hybrid&model=main) | SVM | NEGATIVE | -0.14547655 |
| 263 | 229-244 | [LESKMSGKGQQQQGQ](http://crdd.osdd.net/raghava/ifnepitope/pep_design.php?sequence=LESKMSGKGQQQQGQ&method=hybrid&model=main) | SVM | POSITIVE | 0.052063406 |
| 264 | 230-245 | [ESKMSGKGQQQQGQT](http://crdd.osdd.net/raghava/ifnepitope/pep_design.php?sequence=ESKMSGKGQQQQGQT&method=hybrid&model=main) | SVM | NEGATIVE | -0.079075535 |
| 265 | 231-246 | [SKMSGKGQQQQGQTV](http://crdd.osdd.net/raghava/ifnepitope/pep_design.php?sequence=SKMSGKGQQQQGQTV&method=hybrid&model=main) | SVM | POSITIVE | 0.13456666 |
| 266 | 232-247 | [KMSGKGQQQQGQTVT](http://crdd.osdd.net/raghava/ifnepitope/pep_design.php?sequence=KMSGKGQQQQGQTVT&method=hybrid&model=main) | SVM | POSITIVE | 0.28571522 |
| 267 | 233-248 | [MSGKGQQQQGQTVTK](http://crdd.osdd.net/raghava/ifnepitope/pep_design.php?sequence=MSGKGQQQQGQTVTK&method=hybrid&model=main) | SVM | POSITIVE | 0.37489685 |
| 268 | 234-249 | [SGKGQQQQGQTVTKK](http://crdd.osdd.net/raghava/ifnepitope/pep_design.php?sequence=SGKGQQQQGQTVTKK&method=hybrid&model=main) | SVM | POSITIVE | 0.3922792 |
| 269 | 235-250 | [GKGQQQQGQTVTKKS](http://crdd.osdd.net/raghava/ifnepitope/pep_design.php?sequence=GKGQQQQGQTVTKKS&method=hybrid&model=main) | SVM | POSITIVE | 0.43567523 |
| 270 | 236-251 | [KGQQQQGQTVTKKSA](http://crdd.osdd.net/raghava/ifnepitope/pep_design.php?sequence=KGQQQQGQTVTKKSA&method=hybrid&model=main) | SVM | POSITIVE | 0.46921866 |
| 271 | 237-252 | [GQQQQGQTVTKKSAA](http://crdd.osdd.net/raghava/ifnepitope/pep_design.php?sequence=GQQQQGQTVTKKSAA&method=hybrid&model=main) | SVM | POSITIVE | 0.43394659 |
| 272 | 238-253 | [QQQQGQTVTKKSAAE](http://crdd.osdd.net/raghava/ifnepitope/pep_design.php?sequence=QQQQGQTVTKKSAAE&method=hybrid&model=main) | SVM | POSITIVE | 0.20259606 |
| 273 | 239-254 | [QQQGQTVTKKSAAEA](http://crdd.osdd.net/raghava/ifnepitope/pep_design.php?sequence=QQQGQTVTKKSAAEA&method=hybrid&model=main) | SVM | POSITIVE | 0.082075984 |
| 274 | 240-255 | [QQGQTVTKKSAAEAS](http://crdd.osdd.net/raghava/ifnepitope/pep_design.php?sequence=QQGQTVTKKSAAEAS&method=hybrid&model=main) | SVM | NEGATIVE | -0.27772837 |
| 275 | 241-256 | [QGQTVTKKSAAEASK](http://crdd.osdd.net/raghava/ifnepitope/pep_design.php?sequence=QGQTVTKKSAAEASK&method=hybrid&model=main) | SVM | NEGATIVE | -0.56446879 |
| 276 | 242-257 | [GQTVTKKSAAEASKK](http://crdd.osdd.net/raghava/ifnepitope/pep_design.php?sequence=GQTVTKKSAAEASKK&method=hybrid&model=main) | SVM | NEGATIVE | -0.46861593 |
| 277 | 243-258 | [QTVTKKSAAEASKKP](http://crdd.osdd.net/raghava/ifnepitope/pep_design.php?sequence=QTVTKKSAAEASKKP&method=hybrid&model=main) | SVM | NEGATIVE | -0.43853919 |
| 278 | 244-259 | [TVTKKSAAEASKKPR](http://crdd.osdd.net/raghava/ifnepitope/pep_design.php?sequence=TVTKKSAAEASKKPR&method=hybrid&model=main) | SVM | NEGATIVE | -0.43579373 |
| 279 | 245-260 | [VTKKSAAEASKKPRQ](http://crdd.osdd.net/raghava/ifnepitope/pep_design.php?sequence=VTKKSAAEASKKPRQ&method=hybrid&model=main) | SVM | NEGATIVE | -0.45565206 |
| 280 | 246-261 | [TKKSAAEASKKPRQK](http://crdd.osdd.net/raghava/ifnepitope/pep_design.php?sequence=TKKSAAEASKKPRQK&method=hybrid&model=main) | SVM | NEGATIVE | -0.51387784 |
| 281 | 247-262 | [KKSAAEASKKPRQKR](http://crdd.osdd.net/raghava/ifnepitope/pep_design.php?sequence=KKSAAEASKKPRQKR&method=hybrid&model=main) | SVM | NEGATIVE | -0.60329591 |
| 282 | 248-263 | [KSAAEASKKPRQKRT](http://crdd.osdd.net/raghava/ifnepitope/pep_design.php?sequence=KSAAEASKKPRQKRT&method=hybrid&model=main) | SVM | NEGATIVE | -0.63787479 |
| 283 | 249-264 | [SAAEASKKPRQKRTA](http://crdd.osdd.net/raghava/ifnepitope/pep_design.php?sequence=SAAEASKKPRQKRTA&method=hybrid&model=main) | SVM | NEGATIVE | -0.58406246 |
| 284 | 250-265 | [AAEASKKPRQKRTAT](http://crdd.osdd.net/raghava/ifnepitope/pep_design.php?sequence=AAEASKKPRQKRTAT&method=hybrid&model=main) | SVM | NEGATIVE | -0.67285891 |
| 285 | 251-266 | [AEASKKPRQKRTATK](http://crdd.osdd.net/raghava/ifnepitope/pep_design.php?sequence=AEASKKPRQKRTATK&method=hybrid&model=main) | SVM | NEGATIVE | -0.77695571 |
| 286 | 252-267 | [EASKKPRQKRTATKA](http://crdd.osdd.net/raghava/ifnepitope/pep_design.php?sequence=EASKKPRQKRTATKA&method=hybrid&model=main) | SVM | NEGATIVE | -0.5907313 |
| 287 | 253-268 | [ASKKPRQKRTATKAY](http://crdd.osdd.net/raghava/ifnepitope/pep_design.php?sequence=ASKKPRQKRTATKAY&method=hybrid&model=main) | SVM | NEGATIVE | -0.76285046 |
| 288 | 254-269 | [SKKPRQKRTATKAYN](http://crdd.osdd.net/raghava/ifnepitope/pep_design.php?sequence=SKKPRQKRTATKAYN&method=hybrid&model=main) | SVM | NEGATIVE | -0.62479955 |
| 289 | 255-270 | [KKPRQKRTATKAYNV](http://crdd.osdd.net/raghava/ifnepitope/pep_design.php?sequence=KKPRQKRTATKAYNV&method=hybrid&model=main) | SVM | NEGATIVE | -0.49653899 |
| 290 | 256-271 | [KPRQKRTATKAYNVT](http://crdd.osdd.net/raghava/ifnepitope/pep_design.php?sequence=KPRQKRTATKAYNVT&method=hybrid&model=main) | SVM | NEGATIVE | -0.84130931 |
| 291 | 257-272 | [PRQKRTATKAYNVTQ](http://crdd.osdd.net/raghava/ifnepitope/pep_design.php?sequence=PRQKRTATKAYNVTQ&method=hybrid&model=main) | SVM | NEGATIVE | -0.77719048 |
| 292 | 258-273 | [RQKRTATKAYNVTQA](http://crdd.osdd.net/raghava/ifnepitope/pep_design.php?sequence=RQKRTATKAYNVTQA&method=hybrid&model=main) | SVM | NEGATIVE | -0.53480256 |
| 293 | 259-274 | [QKRTATKAYNVTQAF](http://crdd.osdd.net/raghava/ifnepitope/pep_design.php?sequence=QKRTATKAYNVTQAF&method=hybrid&model=main) | SVM | NEGATIVE | -0.53422098 |
| 294 | 260-275 | [KRTATKAYNVTQAFG](http://crdd.osdd.net/raghava/ifnepitope/pep_design.php?sequence=KRTATKAYNVTQAFG&method=hybrid&model=main) | SVM | NEGATIVE | -0.60429959 |
| 295 | 261-276 | [RTATKAYNVTQAFGR](http://crdd.osdd.net/raghava/ifnepitope/pep_design.php?sequence=RTATKAYNVTQAFGR&method=hybrid&model=main) | SVM | NEGATIVE | -0.39130288 |
| 296 | 262-277 | [TATKAYNVTQAFGRR](http://crdd.osdd.net/raghava/ifnepitope/pep_design.php?sequence=TATKAYNVTQAFGRR&method=hybrid&model=main) | SVM | NEGATIVE | -0.56767131 |
| 297 | 263-278 | [ATKAYNVTQAFGRRG](http://crdd.osdd.net/raghava/ifnepitope/pep_design.php?sequence=ATKAYNVTQAFGRRG&method=hybrid&model=main) | SVM | NEGATIVE | -0.59211136 |
| 298 | 264-279 | [TKAYNVTQAFGRRGP](http://crdd.osdd.net/raghava/ifnepitope/pep_design.php?sequence=TKAYNVTQAFGRRGP&method=hybrid&model=main) | SVM | NEGATIVE | -0.90076061 |
| 299 | 265-280 | [KAYNVTQAFGRRGPE](http://crdd.osdd.net/raghava/ifnepitope/pep_design.php?sequence=KAYNVTQAFGRRGPE&method=hybrid&model=main) | SVM | NEGATIVE | -1.0298071 |
| 300 | 266-281 | [AYNVTQAFGRRGPEQ](http://crdd.osdd.net/raghava/ifnepitope/pep_design.php?sequence=AYNVTQAFGRRGPEQ&method=hybrid&model=main) | SVM | NEGATIVE | -0.81878677 |
| 301 | 267-282 | [YNVTQAFGRRGPEQT](http://crdd.osdd.net/raghava/ifnepitope/pep_design.php?sequence=YNVTQAFGRRGPEQT&method=hybrid&model=main) | SVM | NEGATIVE | -0.79210435 |
| 302 | 268-283 | [NVTQAFGRRGPEQTQ](http://crdd.osdd.net/raghava/ifnepitope/pep_design.php?sequence=NVTQAFGRRGPEQTQ&method=hybrid&model=main) | SVM | NEGATIVE | -0.6086385 |
| 303 | 269-284 | [VTQAFGRRGPEQTQG](http://crdd.osdd.net/raghava/ifnepitope/pep_design.php?sequence=VTQAFGRRGPEQTQG&method=hybrid&model=main) | SVM | NEGATIVE | -0.53656268 |
| 304 | 270-285 | [TQAFGRRGPEQTQGN](http://crdd.osdd.net/raghava/ifnepitope/pep_design.php?sequence=TQAFGRRGPEQTQGN&method=hybrid&model=main) | SVM | NEGATIVE | -0.33791997 |
| 305 | 271-286 | [QAFGRRGPEQTQGNF](http://crdd.osdd.net/raghava/ifnepitope/pep_design.php?sequence=QAFGRRGPEQTQGNF&method=hybrid&model=main) | SVM | NEGATIVE | -0.39547133 |
| 306 | 272-287 | [AFGRRGPEQTQGNFG](http://crdd.osdd.net/raghava/ifnepitope/pep_design.php?sequence=AFGRRGPEQTQGNFG&method=hybrid&model=main) | SVM | NEGATIVE | -0.36765393 |
| 307 | 273-288 | [FGRRGPEQTQGNFGD](http://crdd.osdd.net/raghava/ifnepitope/pep_design.php?sequence=FGRRGPEQTQGNFGD&method=hybrid&model=main) | SVM | NEGATIVE | -0.50835136 |
| 308 | 274-289 | [GRRGPEQTQGNFGDQ](http://crdd.osdd.net/raghava/ifnepitope/pep_design.php?sequence=GRRGPEQTQGNFGDQ&method=hybrid&model=main) | SVM | NEGATIVE | -0.71361468 |
| 309 | 275-290 | [RRGPEQTQGNFGDQE](http://crdd.osdd.net/raghava/ifnepitope/pep_design.php?sequence=RRGPEQTQGNFGDQE&method=hybrid&model=main) | SVM | NEGATIVE | -0.60208568 |
| 310 | 276-291 | [RGPEQTQGNFGDQEL](http://crdd.osdd.net/raghava/ifnepitope/pep_design.php?sequence=RGPEQTQGNFGDQEL&method=hybrid&model=main) | SVM | NEGATIVE | -0.45746957 |
| 311 | 277-292 | [GPEQTQGNFGDQELI](http://crdd.osdd.net/raghava/ifnepitope/pep_design.php?sequence=GPEQTQGNFGDQELI&method=hybrid&model=main) | SVM | NEGATIVE | -0.57943858 |
| 312 | 278-293 | [PEQTQGNFGDQELIR](http://crdd.osdd.net/raghava/ifnepitope/pep_design.php?sequence=PEQTQGNFGDQELIR&method=hybrid&model=main) | SVM | NEGATIVE | -0.20900697 |
| 313 | 279-294 | [EQTQGNFGDQELIRQ](http://crdd.osdd.net/raghava/ifnepitope/pep_design.php?sequence=EQTQGNFGDQELIRQ&method=hybrid&model=main) | SVM | NEGATIVE | -0.17056653 |
| 314 | 280-295 | [QTQGNFGDQELIRQG](http://crdd.osdd.net/raghava/ifnepitope/pep_design.php?sequence=QTQGNFGDQELIRQG&method=hybrid&model=main) | SVM | POSITIVE | 0.11986752 |
| 315 | 281-296 | [TQGNFGDQELIRQGT](http://crdd.osdd.net/raghava/ifnepitope/pep_design.php?sequence=TQGNFGDQELIRQGT&method=hybrid&model=main) | SVM | NEGATIVE | -0.083898484 |
| 316 | 282-297 | [QGNFGDQELIRQGTD](http://crdd.osdd.net/raghava/ifnepitope/pep_design.php?sequence=QGNFGDQELIRQGTD&method=hybrid&model=main) | SVM | NEGATIVE | -0.056196918 |
| 317 | 283-298 | [GNFGDQELIRQGTDY](http://crdd.osdd.net/raghava/ifnepitope/pep_design.php?sequence=GNFGDQELIRQGTDY&method=hybrid&model=main) | SVM | NEGATIVE | -0.097001683 |
| 318 | 284-299 | [NFGDQELIRQGTDYK](http://crdd.osdd.net/raghava/ifnepitope/pep_design.php?sequence=NFGDQELIRQGTDYK&method=hybrid&model=main) | SVM | NEGATIVE | -0.32591643 |
| 319 | 285-300 | [FGDQELIRQGTDYKH](http://crdd.osdd.net/raghava/ifnepitope/pep_design.php?sequence=FGDQELIRQGTDYKH&method=hybrid&model=main) | SVM | NEGATIVE | -0.38390309 |
| 320 | 286-301 | [GDQELIRQGTDYKHW](http://crdd.osdd.net/raghava/ifnepitope/pep_design.php?sequence=GDQELIRQGTDYKHW&method=hybrid&model=main) | SVM | NEGATIVE | -0.3369466 |
| 321 | 287-302 | [DQELIRQGTDYKHWP](http://crdd.osdd.net/raghava/ifnepitope/pep_design.php?sequence=DQELIRQGTDYKHWP&method=hybrid&model=main) | SVM | NEGATIVE | -0.21013452 |
| 322 | 288-303 | [QELIRQGTDYKHWPQ](http://crdd.osdd.net/raghava/ifnepitope/pep_design.php?sequence=QELIRQGTDYKHWPQ&method=hybrid&model=main) | SVM | NEGATIVE | -0.32233915 |
| 323 | 289-304 | [ELIRQGTDYKHWPQI](http://crdd.osdd.net/raghava/ifnepitope/pep_design.php?sequence=ELIRQGTDYKHWPQI&method=hybrid&model=main) | SVM | NEGATIVE | -0.26726527 |
| 324 | 290-305 | [LIRQGTDYKHWPQIA](http://crdd.osdd.net/raghava/ifnepitope/pep_design.php?sequence=LIRQGTDYKHWPQIA&method=hybrid&model=main) | SVM | NEGATIVE | -0.18617189 |
| 325 | 294-309 | [GTDYKHWPQIAQFAP](http://crdd.osdd.net/raghava/ifnepitope/pep_design.php?sequence=GTDYKHWPQIAQFAP&method=hybrid&model=main) | SVM | NEGATIVE | -0.2579848 |
| 326 | 295-310 | [TDYKHWPQIAQFAPS](http://crdd.osdd.net/raghava/ifnepitope/pep_design.php?sequence=TDYKHWPQIAQFAPS&method=hybrid&model=main) | SVM | NEGATIVE | -0.049386579 |
| 327 | 296-311 | [DYKHWPQIAQFAPSA](http://crdd.osdd.net/raghava/ifnepitope/pep_design.php?sequence=DYKHWPQIAQFAPSA&method=hybrid&model=main) | SVM | POSITIVE | 0.26410613 |
| 328 | 303-318 | [IAQFAPSASAFFGMS](http://crdd.osdd.net/raghava/ifnepitope/pep_design.php?sequence=IAQFAPSASAFFGMS&method=hybrid&model=main) | SVM | POSITIVE | 1.0886801 |
| 329 | 304-319 | [AQFAPSASAFFGMSR](http://crdd.osdd.net/raghava/ifnepitope/pep_design.php?sequence=AQFAPSASAFFGMSR&method=hybrid&model=main) | SVM | POSITIVE | 0.99922081 |
| 330 | 305-320 | [QFAPSASAFFGMSRI](http://crdd.osdd.net/raghava/ifnepitope/pep_design.php?sequence=QFAPSASAFFGMSRI&method=hybrid&model=main) | SVM | POSITIVE | 0.74950829 |
| 331 | 306-321 | [FAPSASAFFGMSRIG](http://crdd.osdd.net/raghava/ifnepitope/pep_design.php?sequence=FAPSASAFFGMSRIG&method=hybrid&model=main) | SVM | POSITIVE | 0.69350449 |
| 332 | 307-322 | [APSASAFFGMSRIGM](http://crdd.osdd.net/raghava/ifnepitope/pep_design.php?sequence=APSASAFFGMSRIGM&method=hybrid&model=main) | SVM | POSITIVE | 0.76809868 |
| 333 | 308-323 | [PSASAFFGMSRIGME](http://crdd.osdd.net/raghava/ifnepitope/pep_design.php?sequence=PSASAFFGMSRIGME&method=hybrid&model=main) | SVM | POSITIVE | 0.8677712 |
| 334 | 309-324 | [SASAFFGMSRIGMEV](http://crdd.osdd.net/raghava/ifnepitope/pep_design.php?sequence=SASAFFGMSRIGMEV&method=hybrid&model=main) | SVM | POSITIVE | 0.59323664 |
| 335 | 310-325 | [ASAFFGMSRIGMEVT](http://crdd.osdd.net/raghava/ifnepitope/pep_design.php?sequence=ASAFFGMSRIGMEVT&method=hybrid&model=main) | SVM | NEGATIVE | -0.025297763 |
| 336 | 311-326 | [SAFFGMSRIGMEVTP](http://crdd.osdd.net/raghava/ifnepitope/pep_design.php?sequence=SAFFGMSRIGMEVTP&method=hybrid&model=main) | SVM | NEGATIVE | -0.054745866 |
| 337 | 312-327 | [AFFGMSRIGMEVTPS](http://crdd.osdd.net/raghava/ifnepitope/pep_design.php?sequence=AFFGMSRIGMEVTPS&method=hybrid&model=main) | SVM | NEGATIVE | -0.28960429 |
| 338 | 313-328 | [FFGMSRIGMEVTPSG](http://crdd.osdd.net/raghava/ifnepitope/pep_design.php?sequence=FFGMSRIGMEVTPSG&method=hybrid&model=main) | SVM | NEGATIVE | -0.49919675 |
| 339 | 314-329 | [FGMSRIGMEVTPSGT](http://crdd.osdd.net/raghava/ifnepitope/pep_design.php?sequence=FGMSRIGMEVTPSGT&method=hybrid&model=main) | SVM | NEGATIVE | -0.49068418 |
| 340 | 315-330 | [GMSRIGMEVTPSGTW](http://crdd.osdd.net/raghava/ifnepitope/pep_design.php?sequence=GMSRIGMEVTPSGTW&method=hybrid&model=main) | SVM | NEGATIVE | -0.40502252 |
| 341 | 316-331 | [MSRIGMEVTPSGTWL](http://crdd.osdd.net/raghava/ifnepitope/pep_design.php?sequence=MSRIGMEVTPSGTWL&method=hybrid&model=main) | SVM | NEGATIVE | -0.49087335 |
| 342 | 317-332 | [SRIGMEVTPSGTWLT](http://crdd.osdd.net/raghava/ifnepitope/pep_design.php?sequence=SRIGMEVTPSGTWLT&method=hybrid&model=main) | SVM | NEGATIVE | -0.28218267 |
| 343 | 318-333 | [RIGMEVTPSGTWLTY](http://crdd.osdd.net/raghava/ifnepitope/pep_design.php?sequence=RIGMEVTPSGTWLTY&method=hybrid&model=main) | SVM | NEGATIVE | -0.31234822 |
| 344 | 319-334 | [IGMEVTPSGTWLTYT](http://crdd.osdd.net/raghava/ifnepitope/pep_design.php?sequence=IGMEVTPSGTWLTYT&method=hybrid&model=main) | SVM | NEGATIVE | -0.34112841 |
| 345 | 320-335 | [GMEVTPSGTWLTYTG](http://crdd.osdd.net/raghava/ifnepitope/pep_design.php?sequence=GMEVTPSGTWLTYTG&method=hybrid&model=main) | SVM | NEGATIVE | -0.33484195 |
| 346 | 321-336 | [MEVTPSGTWLTYTGA](http://crdd.osdd.net/raghava/ifnepitope/pep_design.php?sequence=MEVTPSGTWLTYTGA&method=hybrid&model=main) | SVM | NEGATIVE | -0.052873528 |
| 347 | 322-337 | [EVTPSGTWLTYTGAI](http://crdd.osdd.net/raghava/ifnepitope/pep_design.php?sequence=EVTPSGTWLTYTGAI&method=hybrid&model=main) | SVM | NEGATIVE | -0.067629059 |
| 348 | 323-338 | [VTPSGTWLTYTGAIK](http://crdd.osdd.net/raghava/ifnepitope/pep_design.php?sequence=VTPSGTWLTYTGAIK&method=hybrid&model=main) | SVM | POSITIVE | 0.066403069 |
| 349 | 324-339 | [TPSGTWLTYTGAIKL](http://crdd.osdd.net/raghava/ifnepitope/pep_design.php?sequence=TPSGTWLTYTGAIKL&method=hybrid&model=main) | SVM | POSITIVE | 0.14513136 |
| 350 | 325-340 | [PSGTWLTYTGAIKLD](http://crdd.osdd.net/raghava/ifnepitope/pep_design.php?sequence=PSGTWLTYTGAIKLD&method=hybrid&model=main) | SVM | NEGATIVE | -0.17459442 |
| 351 | 334-349 | [GAIKLDDKDPNFKDQ](http://crdd.osdd.net/raghava/ifnepitope/pep_design.php?sequence=GAIKLDDKDPNFKDQ&method=hybrid&model=main) | SVM | NEGATIVE | -0.65898076 |
| 352 | 335-350 | [AIKLDDKDPNFKDQV](http://crdd.osdd.net/raghava/ifnepitope/pep_design.php?sequence=AIKLDDKDPNFKDQV&method=hybrid&model=main) | SVM | NEGATIVE | -0.54595864 |
| 353 | 336-351 | [IKLDDKDPNFKDQVI](http://crdd.osdd.net/raghava/ifnepitope/pep_design.php?sequence=IKLDDKDPNFKDQVI&method=hybrid&model=main) | SVM | NEGATIVE | -0.32988866 |
| 354 | 337-352 | [KLDDKDPNFKDQVIL](http://crdd.osdd.net/raghava/ifnepitope/pep_design.php?sequence=KLDDKDPNFKDQVIL&method=hybrid&model=main) | SVM | NEGATIVE | -0.31186955 |
| 355 | 338-353 | [LDDKDPNFKDQVILL](http://crdd.osdd.net/raghava/ifnepitope/pep_design.php?sequence=LDDKDPNFKDQVILL&method=hybrid&model=main) | SVM | NEGATIVE | -0.2567984 |
| 356 | 348-363 | [QVILLNKHIDAYKTF](http://crdd.osdd.net/raghava/ifnepitope/pep_design.php?sequence=QVILLNKHIDAYKTF&method=hybrid&model=main) | SVM | NEGATIVE | -0.13552403 |
| 357 | 349-364 | [VILLNKHIDAYKTFP](http://crdd.osdd.net/raghava/ifnepitope/pep_design.php?sequence=VILLNKHIDAYKTFP&method=hybrid&model=main) | SVM | NEGATIVE | -0.26705025 |
| 358 | 350-365 | [ILLNKHIDAYKTFPP](http://crdd.osdd.net/raghava/ifnepitope/pep_design.php?sequence=ILLNKHIDAYKTFPP&method=hybrid&model=main) | SVM | NEGATIVE | -0.56066857 |
| 359 | 351-366 | [LLNKHIDAYKTFPPT](http://crdd.osdd.net/raghava/ifnepitope/pep_design.php?sequence=LLNKHIDAYKTFPPT&method=hybrid&model=main) | SVM | NEGATIVE | -0.82019076 |
| 360 | 352-367 | [LNKHIDAYKTFPPTE](http://crdd.osdd.net/raghava/ifnepitope/pep_design.php?sequence=LNKHIDAYKTFPPTE&method=hybrid&model=main) | SVM | NEGATIVE | -0.77550579 |
| 361 | 353-368 | [NKHIDAYKTFPPTEP](http://crdd.osdd.net/raghava/ifnepitope/pep_design.php?sequence=NKHIDAYKTFPPTEP&method=hybrid&model=main) | SVM | NEGATIVE | -0.64090052 |
| 362 | 354-369 | [KHIDAYKTFPPTEPK](http://crdd.osdd.net/raghava/ifnepitope/pep_design.php?sequence=KHIDAYKTFPPTEPK&method=hybrid&model=main) | SVM | NEGATIVE | -0.81716417 |
| 363 | 355-370 | [HIDAYKTFPPTEPKK](http://crdd.osdd.net/raghava/ifnepitope/pep_design.php?sequence=HIDAYKTFPPTEPKK&method=hybrid&model=main) | SVM | NEGATIVE | -0.90479712 |
| 364 | 356-371 | [IDAYKTFPPTEPKKD](http://crdd.osdd.net/raghava/ifnepitope/pep_design.php?sequence=IDAYKTFPPTEPKKD&method=hybrid&model=main) | SVM | NEGATIVE | -1.1121354 |
| 365 | 357-372 | [DAYKTFPPTEPKKDK](http://crdd.osdd.net/raghava/ifnepitope/pep_design.php?sequence=DAYKTFPPTEPKKDK&method=hybrid&model=main) | SVM | NEGATIVE | -0.96900825 |
| 366 | 358-373 | [AYKTFPPTEPKKDKK](http://crdd.osdd.net/raghava/ifnepitope/pep_design.php?sequence=AYKTFPPTEPKKDKK&method=hybrid&model=main) | SVM | NEGATIVE | -0.79210183 |
| 367 | 359-374 | [YKTFPPTEPKKDKKK](http://crdd.osdd.net/raghava/ifnepitope/pep_design.php?sequence=YKTFPPTEPKKDKKK&method=hybrid&model=main) | SVM | NEGATIVE | -0.54867807 |
| 368 | 360-375 | [KTFPPTEPKKDKKKK](http://crdd.osdd.net/raghava/ifnepitope/pep_design.php?sequence=KTFPPTEPKKDKKKK&method=hybrid&model=main) | SVM | NEGATIVE | -0.18433032 |
| 369 | 361-376 | [TFPPTEPKKDKKKKA](http://crdd.osdd.net/raghava/ifnepitope/pep_design.php?sequence=TFPPTEPKKDKKKKA&method=hybrid&model=main) | SVM | POSITIVE | 0.0030940686 |
| 370 | 362-377 | [FPPTEPKKDKKKKAD](http://crdd.osdd.net/raghava/ifnepitope/pep_design.php?sequence=FPPTEPKKDKKKKAD&method=hybrid&model=main) | SVM | POSITIVE | 0.1229108 |
| 371 | 363-378 | [PPTEPKKDKKKKADE](http://crdd.osdd.net/raghava/ifnepitope/pep_design.php?sequence=PPTEPKKDKKKKADE&method=hybrid&model=main) | SVM | POSITIVE | 0.082799207 |
| 372 | 364-379 | [PTEPKKDKKKKADET](http://crdd.osdd.net/raghava/ifnepitope/pep_design.php?sequence=PTEPKKDKKKKADET&method=hybrid&model=main) | SVM | POSITIVE | 0.11708686 |
| 373 | 365-380 | [TEPKKDKKKKADETQ](http://crdd.osdd.net/raghava/ifnepitope/pep_design.php?sequence=TEPKKDKKKKADETQ&method=hybrid&model=main) | SVM | NEGATIVE | -0.078024331 |
| 374 | 366-381 | [EPKKDKKKKADETQA](http://crdd.osdd.net/raghava/ifnepitope/pep_design.php?sequence=EPKKDKKKKADETQA&method=hybrid&model=main) | SVM | NEGATIVE | -0.16706402 |
| 375 | 367-382 | [PKKDKKKKADETQAL](http://crdd.osdd.net/raghava/ifnepitope/pep_design.php?sequence=PKKDKKKKADETQAL&method=hybrid&model=main) | SVM | NEGATIVE | -0.19429611 |
| 376 | 368-383 | [KKDKKKKADETQALP](http://crdd.osdd.net/raghava/ifnepitope/pep_design.php?sequence=KKDKKKKADETQALP&method=hybrid&model=main) | SVM | NEGATIVE | -0.23179577 |
| 377 | 369-384 | [KDKKKKADETQALPQ](http://crdd.osdd.net/raghava/ifnepitope/pep_design.php?sequence=KDKKKKADETQALPQ&method=hybrid&model=main) | SVM | NEGATIVE | -0.70330412 |
| 378 | 370-385 | [DKKKKADETQALPQR](http://crdd.osdd.net/raghava/ifnepitope/pep_design.php?sequence=DKKKKADETQALPQR&method=hybrid&model=main) | SVM | NEGATIVE | -0.30613892 |
| 379 | 371-386 | [KKKKADETQALPQRQ](http://crdd.osdd.net/raghava/ifnepitope/pep_design.php?sequence=KKKKADETQALPQRQ&method=hybrid&model=main) | SVM | NEGATIVE | -0.31301639 |
| 380 | 372-387 | [KKKADETQALPQRQK](http://crdd.osdd.net/raghava/ifnepitope/pep_design.php?sequence=KKKADETQALPQRQK&method=hybrid&model=main) | SVM | NEGATIVE | -0.71768852 |
| 381 | 373-388 | [KKADETQALPQRQKK](http://crdd.osdd.net/raghava/ifnepitope/pep_design.php?sequence=KKADETQALPQRQKK&method=hybrid&model=main) | SVM | NEGATIVE | -0.71768852 |
| 382 | 374-389 | [KADETQALPQRQKKQ](http://crdd.osdd.net/raghava/ifnepitope/pep_design.php?sequence=KADETQALPQRQKKQ&method=hybrid&model=main) | SVM | NEGATIVE | -0.96618469 |
| 383 | 375-390 | [ADETQALPQRQKKQQ](http://crdd.osdd.net/raghava/ifnepitope/pep_design.php?sequence=ADETQALPQRQKKQQ&method=hybrid&model=main) | SVM | NEGATIVE | -1.1138755 |
| 384 | 376-391 | [DETQALPQRQKKQQT](http://crdd.osdd.net/raghava/ifnepitope/pep_design.php?sequence=DETQALPQRQKKQQT&method=hybrid&model=main) | SVM | NEGATIVE | -1.0429722 |
| 385 | 377-392 | [ETQALPQRQKKQQTV](http://crdd.osdd.net/raghava/ifnepitope/pep_design.php?sequence=ETQALPQRQKKQQTV&method=hybrid&model=main) | SVM | NEGATIVE | -0.92152785 |
| 386 | 378-393 | [TQALPQRQKKQQTVT](http://crdd.osdd.net/raghava/ifnepitope/pep_design.php?sequence=TQALPQRQKKQQTVT&method=hybrid&model=main) | SVM | NEGATIVE | -0.84387011 |
| 387 | 379-394 | [QALPQRQKKQQTVTL](http://crdd.osdd.net/raghava/ifnepitope/pep_design.php?sequence=QALPQRQKKQQTVTL&method=hybrid&model=main) | SVM | NEGATIVE | -0.59159268 |
| 388 | 380-395 | [ALPQRQKKQQTVTLL](http://crdd.osdd.net/raghava/ifnepitope/pep_design.php?sequence=ALPQRQKKQQTVTLL&method=hybrid&model=main) | SVM | NEGATIVE | -0.36907385 |
| 389 | 381-396 | [LPQRQKKQQTVTLLP](http://crdd.osdd.net/raghava/ifnepitope/pep_design.php?sequence=LPQRQKKQQTVTLLP&method=hybrid&model=main) | SVM | NEGATIVE | -0.23008436 |
| 390 | 382-397 | [PQRQKKQQTVTLLPA](http://crdd.osdd.net/raghava/ifnepitope/pep_design.php?sequence=PQRQKKQQTVTLLPA&method=hybrid&model=main) | SVM | NEGATIVE | -0.31553613 |
| 391 | 383-398 | [QRQKKQQTVTLLPAA](http://crdd.osdd.net/raghava/ifnepitope/pep_design.php?sequence=QRQKKQQTVTLLPAA&method=hybrid&model=main) | SVM | NEGATIVE | -0.058363837 |
| 392 | 384-399 | [RQKKQQTVTLLPAAD](http://crdd.osdd.net/raghava/ifnepitope/pep_design.php?sequence=RQKKQQTVTLLPAAD&method=hybrid&model=main) | SVM | NEGATIVE | -0.34911436 |
| 393 | 385-400 | [QKKQQTVTLLPAADL](http://crdd.osdd.net/raghava/ifnepitope/pep_design.php?sequence=QKKQQTVTLLPAADL&method=hybrid&model=main) | SVM | NEGATIVE | -0.14422474 |
| 394 | 386-401 | [KKQQTVTLLPAADLD](http://crdd.osdd.net/raghava/ifnepitope/pep_design.php?sequence=KKQQTVTLLPAADLD&method=hybrid&model=main) | SVM | NEGATIVE | -0.10288586 |
| 395 | 387-402 | [KQQTVTLLPAADLDD](http://crdd.osdd.net/raghava/ifnepitope/pep_design.php?sequence=KQQTVTLLPAADLDD&method=hybrid&model=main) | SVM | NEGATIVE | -0.42616623 |
| 396 | 388-403 | [QQTVTLLPAADLDDF](http://crdd.osdd.net/raghava/ifnepitope/pep_design.php?sequence=QQTVTLLPAADLDDF&method=hybrid&model=main) | SVM | NEGATIVE | -0.27974059 |
| 397 | 389-404 | [QTVTLLPAADLDDFS](http://crdd.osdd.net/raghava/ifnepitope/pep_design.php?sequence=QTVTLLPAADLDDFS&method=hybrid&model=main) | SVM | NEGATIVE | -0.20921521 |
| 398 | 390-405 | [TVTLLPAADLDDFSK](http://crdd.osdd.net/raghava/ifnepitope/pep_design.php?sequence=TVTLLPAADLDDFSK&method=hybrid&model=main) | SVM | NEGATIVE | -0.57932117 |
| 399 | 391-406 | [VTLLPAADLDDFSKQ](http://crdd.osdd.net/raghava/ifnepitope/pep_design.php?sequence=VTLLPAADLDDFSKQ&method=hybrid&model=main) | SVM | NEGATIVE | -0.84068754 |
| 400 | 392-407 | [TLLPAADLDDFSKQL](http://crdd.osdd.net/raghava/ifnepitope/pep_design.php?sequence=TLLPAADLDDFSKQL&method=hybrid&model=main) | SVM | NEGATIVE | -0.91195804 |
| 401 | 393-408 | [LLPAADLDDFSKQLQ](http://crdd.osdd.net/raghava/ifnepitope/pep_design.php?sequence=LLPAADLDDFSKQLQ&method=hybrid&model=main) | SVM | NEGATIVE | -1.101453 |
| 402 | 394-409 | [LPAADLDDFSKQLQQ](http://crdd.osdd.net/raghava/ifnepitope/pep_design.php?sequence=LPAADLDDFSKQLQQ&method=hybrid&model=main) | SVM | NEGATIVE | -1.0620777 |
| 403 | 395-410 | [PAADLDDFSKQLQQS](http://crdd.osdd.net/raghava/ifnepitope/pep_design.php?sequence=PAADLDDFSKQLQQS&method=hybrid&model=main) | SVM | NEGATIVE | -0.98182617 |
| 404 | 396-411 | [AADLDDFSKQLQQSM](http://crdd.osdd.net/raghava/ifnepitope/pep_design.php?sequence=AADLDDFSKQLQQSM&method=hybrid&model=main) | SVM | NEGATIVE | -1.1328703 |
| 405 | 397-412 | [ADLDDFSKQLQQSMS](http://crdd.osdd.net/raghava/ifnepitope/pep_design.php?sequence=ADLDDFSKQLQQSMS&method=hybrid&model=main) | SVM | NEGATIVE | -1.1370012 |
| 406 | 402-417 | [FSKQLQQSMSSADST](http://crdd.osdd.net/raghava/ifnepitope/pep_design.php?sequence=FSKQLQQSMSSADST&method=hybrid&model=main) | SVM | NEGATIVE | -0.92603454 |
| 407 | 403-418 | [SKQLQQSMSSADSTQ](http://crdd.osdd.net/raghava/ifnepitope/pep_design.php?sequence=SKQLQQSMSSADSTQ&method=hybrid&model=main) | SVM | NEGATIVE | -0.96414521 |
| 408 | 404-419 | [KQLQQSMSSADSTQA](http://crdd.osdd.net/raghava/ifnepitope/pep_design.php?sequence=KQLQQSMSSADSTQA&method=hybrid&model=main) | SVM | NEGATIVE | -0.77229272 |
| 409 | 405-419 | [QLQQSMSSADSTQA](http://crdd.osdd.net/raghava/ifnepitope/pep_design.php?sequence=QLQQSMSSADSTQA&method=hybrid&model=main) | SVM | NEGATIVE | -0.65661162 |
| 410 | 406-419 | [LQQSMSSADSTQA](http://crdd.osdd.net/raghava/ifnepitope/pep_design.php?sequence=LQQSMSSADSTQA&method=hybrid&model=main) | SVM | NEGATIVE | -0.75249777 |
| 411 | 407-419 | [QQSMSSADSTQA](http://crdd.osdd.net/raghava/ifnepitope/pep_design.php?sequence=QQSMSSADSTQA&method=hybrid&model=main) | SVM | NEGATIVE | -0.69025354 |
| ORF6 Protein | | | | | |
| 1 | 0-15 | [MFHLVDFQVTIAEIL](http://crdd.osdd.net/raghava/ifnepitope/pep_design.php?sequence=MFHLVDFQVTIAEIL&method=hybrid&model=main) | MERCI | NEGATIVE | 2 |
| 2 | 1-16 | [FHLVDFQVTIAEILL](http://crdd.osdd.net/raghava/ifnepitope/pep_design.php?sequence=FHLVDFQVTIAEILL&method=hybrid&model=main) | MERCI | NEGATIVE | 2 |
| 3 | 2-17 | [HLVDFQVTIAEILLI](http://crdd.osdd.net/raghava/ifnepitope/pep_design.php?sequence=HLVDFQVTIAEILLI&method=hybrid&model=main) | MERCI | NEGATIVE | 5 |
| 4 | 3-18 | [LVDFQVTIAEILLII](http://crdd.osdd.net/raghava/ifnepitope/pep_design.php?sequence=LVDFQVTIAEILLII&method=hybrid&model=main) | MERCI | NEGATIVE | 23 |
| 5 | 4-19 | [VDFQVTIAEILLIIM](http://crdd.osdd.net/raghava/ifnepitope/pep_design.php?sequence=VDFQVTIAEILLIIM&method=hybrid&model=main) | MERCI | NEGATIVE | 24 |
| 6 | 5-20 | [DFQVTIAEILLIIMR](http://crdd.osdd.net/raghava/ifnepitope/pep_design.php?sequence=DFQVTIAEILLIIMR&method=hybrid&model=main) | MERCI | NEGATIVE | 22 |
| 7 | 6-21 | [FQVTIAEILLIIMRT](http://crdd.osdd.net/raghava/ifnepitope/pep_design.php?sequence=FQVTIAEILLIIMRT&method=hybrid&model=main) | MERCI | NEGATIVE | 22 |
| 8 | 7-22 | [QVTIAEILLIIMRTF](http://crdd.osdd.net/raghava/ifnepitope/pep_design.php?sequence=QVTIAEILLIIMRTF&method=hybrid&model=main) | MERCI | NEGATIVE | 18 |
| 9 | 8-23 | [VTIAEILLIIMRTFK](http://crdd.osdd.net/raghava/ifnepitope/pep_design.php?sequence=VTIAEILLIIMRTFK&method=hybrid&model=main) | MERCI | NEGATIVE | 18 |
| 10 | 9-24 | [TIAEILLIIMRTFKV](http://crdd.osdd.net/raghava/ifnepitope/pep_design.php?sequence=TIAEILLIIMRTFKV&method=hybrid&model=main) | MERCI | NEGATIVE | 19 |
| 11 | 10-25 | [IAEILLIIMRTFKVS](http://crdd.osdd.net/raghava/ifnepitope/pep_design.php?sequence=IAEILLIIMRTFKVS&method=hybrid&model=main) | MERCI | NEGATIVE | 20 |
| 12 | 11-26 | [AEILLIIMRTFKVSI](http://crdd.osdd.net/raghava/ifnepitope/pep_design.php?sequence=AEILLIIMRTFKVSI&method=hybrid&model=main) | MERCI | NEGATIVE | 11 |
| 13 | 12-27 | [EILLIIMRTFKVSIW](http://crdd.osdd.net/raghava/ifnepitope/pep_design.php?sequence=EILLIIMRTFKVSIW&method=hybrid&model=main) | MERCI | NEGATIVE | 11 |
| 14 | 13-28 | [ILLIIMRTFKVSIWN](http://crdd.osdd.net/raghava/ifnepitope/pep_design.php?sequence=ILLIIMRTFKVSIWN&method=hybrid&model=main) | MERCI | NEGATIVE | 11 |
| 15 | 14-29 | [LLIIMRTFKVSIWNL](http://crdd.osdd.net/raghava/ifnepitope/pep_design.php?sequence=LLIIMRTFKVSIWNL&method=hybrid&model=main) | MERCI | NEGATIVE | 2 |
| 16 | 20-35 | [TFKVSIWNLDYIINL](http://crdd.osdd.net/raghava/ifnepitope/pep_design.php?sequence=TFKVSIWNLDYIINL&method=hybrid&model=main) | MERCI | NEGATIVE | 1 |
| 17 | 21-36 | [FKVSIWNLDYIINLI](http://crdd.osdd.net/raghava/ifnepitope/pep_design.php?sequence=FKVSIWNLDYIINLI&method=hybrid&model=main) | MERCI | NEGATIVE | 1 |
| 18 | 22-37 | [KVSIWNLDYIINLII](http://crdd.osdd.net/raghava/ifnepitope/pep_design.php?sequence=KVSIWNLDYIINLII&method=hybrid&model=main) | MERCI | NEGATIVE | 7 |
| 19 | 23-38 | [VSIWNLDYIINLIIK](http://crdd.osdd.net/raghava/ifnepitope/pep_design.php?sequence=VSIWNLDYIINLIIK&method=hybrid&model=main) | MERCI | NEGATIVE | 8 |
| 20 | 24-39 | [SIWNLDYIINLIIKN](http://crdd.osdd.net/raghava/ifnepitope/pep_design.php?sequence=SIWNLDYIINLIIKN&method=hybrid&model=main) | MERCI | NEGATIVE | 7 |
| 21 | 25-40 | [IWNLDYIINLIIKNL](http://crdd.osdd.net/raghava/ifnepitope/pep_design.php?sequence=IWNLDYIINLIIKNL&method=hybrid&model=main) | MERCI | NEGATIVE | 9 |
| 22 | 26-41 | [WNLDYIINLIIKNLS](http://crdd.osdd.net/raghava/ifnepitope/pep_design.php?sequence=WNLDYIINLIIKNLS&method=hybrid&model=main) | MERCI | NEGATIVE | 9 |
| 23 | 27-42 | [NLDYIINLIIKNLSK](http://crdd.osdd.net/raghava/ifnepitope/pep_design.php?sequence=NLDYIINLIIKNLSK&method=hybrid&model=main) | MERCI | NEGATIVE | 9 |
| 24 | 28-43 | [LDYIINLIIKNLSKS](http://crdd.osdd.net/raghava/ifnepitope/pep_design.php?sequence=LDYIINLIIKNLSKS&method=hybrid&model=main) | MERCI | NEGATIVE | 8 |
| 25 | 29-44 | [DYIINLIIKNLSKSL](http://crdd.osdd.net/raghava/ifnepitope/pep_design.php?sequence=DYIINLIIKNLSKSL&method=hybrid&model=main) | MERCI | NEGATIVE | 10 |
| 26 | 30-45 | [YIINLIIKNLSKSLT](http://crdd.osdd.net/raghava/ifnepitope/pep_design.php?sequence=YIINLIIKNLSKSLT&method=hybrid&model=main) | MERCI | NEGATIVE | 7 |
| 27 | 31-46 | [IINLIIKNLSKSLTE](http://crdd.osdd.net/raghava/ifnepitope/pep_design.php?sequence=IINLIIKNLSKSLTE&method=hybrid&model=main) | MERCI | NEGATIVE | 4 |
| 28 | 32-47 | [INLIIKNLSKSLTEN](http://crdd.osdd.net/raghava/ifnepitope/pep_design.php?sequence=INLIIKNLSKSLTEN&method=hybrid&model=main) | MERCI | NEGATIVE | 1 |
| 29 | 33-48 | [NLIIKNLSKSLTENK](http://crdd.osdd.net/raghava/ifnepitope/pep_design.php?sequence=NLIIKNLSKSLTENK&method=hybrid&model=main) | MERCI | NEGATIVE | 1 |
| 30 | 34-49 | [LIIKNLSKSLTENKY](http://crdd.osdd.net/raghava/ifnepitope/pep_design.php?sequence=LIIKNLSKSLTENKY&method=hybrid&model=main) | MERCI | NEGATIVE | 1 |
| 31 | 15-30 | [LIIMRTFKVSIWNLD](http://crdd.osdd.net/raghava/ifnepitope/pep_design.php?sequence=LIIMRTFKVSIWNLD&method=hybrid&model=main) | SVM | NEGATIVE | -0.38805174 |
| 32 | 16-31 | [IIMRTFKVSIWNLDY](http://crdd.osdd.net/raghava/ifnepitope/pep_design.php?sequence=IIMRTFKVSIWNLDY&method=hybrid&model=main) | SVM | NEGATIVE | -0.41956694 |
| 33 | 17-32 | [IMRTFKVSIWNLDYI](http://crdd.osdd.net/raghava/ifnepitope/pep_design.php?sequence=IMRTFKVSIWNLDYI&method=hybrid&model=main) | SVM | NEGATIVE | -0.35278002 |
| 34 | 18-33 | [MRTFKVSIWNLDYII](http://crdd.osdd.net/raghava/ifnepitope/pep_design.php?sequence=MRTFKVSIWNLDYII&method=hybrid&model=main) | SVM | NEGATIVE | -0.32950599 |
| 35 | 19-34 | [RTFKVSIWNLDYIIN](http://crdd.osdd.net/raghava/ifnepitope/pep_design.php?sequence=RTFKVSIWNLDYIIN&method=hybrid&model=main) | SVM | NEGATIVE | -0.29121379 |
| 36 | 35-50 | [IIKNLSKSLTENKYS](http://crdd.osdd.net/raghava/ifnepitope/pep_design.php?sequence=IIKNLSKSLTENKYS&method=hybrid&model=main) | SVM | NEGATIVE | -0.91502937 |
| 37 | 36-51 | [IKNLSKSLTENKYSQ](http://crdd.osdd.net/raghava/ifnepitope/pep_design.php?sequence=IKNLSKSLTENKYSQ&method=hybrid&model=main) | SVM | NEGATIVE | -0.85002873 |
| 38 | 37-52 | [KNLSKSLTENKYSQL](http://crdd.osdd.net/raghava/ifnepitope/pep_design.php?sequence=KNLSKSLTENKYSQL&method=hybrid&model=main) | SVM | NEGATIVE | -0.86207691 |
| 39 | 38-53 | [NLSKSLTENKYSQLD](http://crdd.osdd.net/raghava/ifnepitope/pep_design.php?sequence=NLSKSLTENKYSQLD&method=hybrid&model=main) | SVM | NEGATIVE | -0.9509903 |
| 40 | 39-54 | [LSKSLTENKYSQLDE](http://crdd.osdd.net/raghava/ifnepitope/pep_design.php?sequence=LSKSLTENKYSQLDE&method=hybrid&model=main) | SVM | NEGATIVE | -0.83797618 |
| 41 | 40-55 | [SKSLTENKYSQLDEE](http://crdd.osdd.net/raghava/ifnepitope/pep_design.php?sequence=SKSLTENKYSQLDEE&method=hybrid&model=main) | SVM | NEGATIVE | -0.66913241 |
| 42 | 41-56 | [KSLTENKYSQLDEEQ](http://crdd.osdd.net/raghava/ifnepitope/pep_design.php?sequence=KSLTENKYSQLDEEQ&method=hybrid&model=main) | SVM | NEGATIVE | -0.55981215 |
| 43 | 42-57 | [SLTENKYSQLDEEQP](http://crdd.osdd.net/raghava/ifnepitope/pep_design.php?sequence=SLTENKYSQLDEEQP&method=hybrid&model=main) | SVM | NEGATIVE | -0.4988042 |
| 44 | 43-58 | [LTENKYSQLDEEQPM](http://crdd.osdd.net/raghava/ifnepitope/pep_design.php?sequence=LTENKYSQLDEEQPM&method=hybrid&model=main) | SVM | NEGATIVE | -0.56158647 |
| 45 | 44-59 | [TENKYSQLDEEQPME](http://crdd.osdd.net/raghava/ifnepitope/pep_design.php?sequence=TENKYSQLDEEQPME&method=hybrid&model=main) | SVM | NEGATIVE | -0.51177155 |
| 46 | 45-60 | [ENKYSQLDEEQPMEI](http://crdd.osdd.net/raghava/ifnepitope/pep_design.php?sequence=ENKYSQLDEEQPMEI&method=hybrid&model=main) | SVM | NEGATIVE | -0.46717256 |
| 47 | 46-61 | [NKYSQLDEEQPMEID](http://crdd.osdd.net/raghava/ifnepitope/pep_design.php?sequence=NKYSQLDEEQPMEID&method=hybrid&model=main) | SVM | NEGATIVE | -0.76803231 |
| 48 | 47-61 | [KYSQLDEEQPMEID](http://crdd.osdd.net/raghava/ifnepitope/pep_design.php?sequence=KYSQLDEEQPMEID&method=hybrid&model=main) | SVM | NEGATIVE | -0.80689514 |
| 49 | 48-61 | [YSQLDEEQPMEID](http://crdd.osdd.net/raghava/ifnepitope/pep_design.php?sequence=YSQLDEEQPMEID&method=hybrid&model=main) | SVM | NEGATIVE | -0.77797843 |
| 50 | 49-61 | [SQLDEEQPMEID](http://crdd.osdd.net/raghava/ifnepitope/pep_design.php?sequence=SQLDEEQPMEID&method=hybrid&model=main) | SVM | NEGATIVE | -0.75147657 |
| 51 | 50-61 | [QLDEEQPMEID](http://crdd.osdd.net/raghava/ifnepitope/pep_design.php?sequence=QLDEEQPMEID&method=hybrid&model=main) | SVM | NEGATIVE | -0.72475018 |
| 52 | 51-61 | [LDEEQPMEID](http://crdd.osdd.net/raghava/ifnepitope/pep_design.php?sequence=LDEEQPMEID&method=hybrid&model=main) | SVM | NEGATIVE | -0.5599827 |
| 53 | 52-61 | [DEEQPMEID](http://crdd.osdd.net/raghava/ifnepitope/pep_design.php?sequence=DEEQPMEID&method=hybrid&model=main) | SVM | NEGATIVE | -0.16514591 |
| ORF7a Protein | | | | | |
| 1 | 0-15 | [MKIILFLALITLATC](http://crdd.osdd.net/raghava/ifnepitope/pep_design.php?sequence=MKIILFLALITLATC&method=hybrid&model=main) | MERCI | NEGATIVE | 35 |
| 2 | 1-16 | [KIILFLALITLATCE](http://crdd.osdd.net/raghava/ifnepitope/pep_design.php?sequence=KIILFLALITLATCE&method=hybrid&model=main) | MERCI | NEGATIVE | 28 |
| 3 | 2-17 | [IILFLALITLATCEL](http://crdd.osdd.net/raghava/ifnepitope/pep_design.php?sequence=IILFLALITLATCEL&method=hybrid&model=main) | MERCI | NEGATIVE | 19 |
| 4 | 3-18 | [ILFLALITLATCELY](http://crdd.osdd.net/raghava/ifnepitope/pep_design.php?sequence=ILFLALITLATCELY&method=hybrid&model=main) | MERCI | NEGATIVE | 8 |
| 5 | 4-19 | [LFLALITLATCELYH](http://crdd.osdd.net/raghava/ifnepitope/pep_design.php?sequence=LFLALITLATCELYH&method=hybrid&model=main) | MERCI | NEGATIVE | 5 |
| 6 | 5-20 | [FLALITLATCELYHY](http://crdd.osdd.net/raghava/ifnepitope/pep_design.php?sequence=FLALITLATCELYHY&method=hybrid&model=main) | MERCI | NEGATIVE | 4 |
| 7 | 6-21 | [LALITLATCELYHYQ](http://crdd.osdd.net/raghava/ifnepitope/pep_design.php?sequence=LALITLATCELYHYQ&method=hybrid&model=main) | MERCI | NEGATIVE | 2 |
| 8 | 7-22 | [ALITLATCELYHYQE](http://crdd.osdd.net/raghava/ifnepitope/pep_design.php?sequence=ALITLATCELYHYQE&method=hybrid&model=main) | MERCI | NEGATIVE | 1 |
| 9 | 46-61 | [HPLADNKFALTCFST](http://crdd.osdd.net/raghava/ifnepitope/pep_design.php?sequence=HPLADNKFALTCFST&method=hybrid&model=main) | MERCI | NEGATIVE | 1 |
| 10 | 47-62 | [PLADNKFALTCFSTQ](http://crdd.osdd.net/raghava/ifnepitope/pep_design.php?sequence=PLADNKFALTCFSTQ&method=hybrid&model=main) | MERCI | NEGATIVE | 1 |
| 11 | 48-63 | [LADNKFALTCFSTQF](http://crdd.osdd.net/raghava/ifnepitope/pep_design.php?sequence=LADNKFALTCFSTQF&method=hybrid&model=main) | MERCI | NEGATIVE | 1 |
| 12 | 49-64 | [ADNKFALTCFSTQFA](http://crdd.osdd.net/raghava/ifnepitope/pep_design.php?sequence=ADNKFALTCFSTQFA&method=hybrid&model=main) | MERCI | NEGATIVE | 1 |
| 13 | 50-65 | [DNKFALTCFSTQFAF](http://crdd.osdd.net/raghava/ifnepitope/pep_design.php?sequence=DNKFALTCFSTQFAF&method=hybrid&model=main) | MERCI | NEGATIVE | 1 |
| 14 | 51-66 | [NKFALTCFSTQFAFA](http://crdd.osdd.net/raghava/ifnepitope/pep_design.php?sequence=NKFALTCFSTQFAFA&method=hybrid&model=main) | MERCI | NEGATIVE | 1 |
| 15 | 52-67 | [KFALTCFSTQFAFAC](http://crdd.osdd.net/raghava/ifnepitope/pep_design.php?sequence=KFALTCFSTQFAFAC&method=hybrid&model=main) | MERCI | NEGATIVE | 1 |
| 16 | 89-104 | [QEEVQELYSPIFLIV](http://crdd.osdd.net/raghava/ifnepitope/pep_design.php?sequence=QEEVQELYSPIFLIV&method=hybrid&model=main) | MERCI | NEGATIVE | 1 |
| 17 | 90-105 | [EEVQELYSPIFLIVA](http://crdd.osdd.net/raghava/ifnepitope/pep_design.php?sequence=EEVQELYSPIFLIVA&method=hybrid&model=main) | MERCI | NEGATIVE | 6 |
| 18 | 91-106 | [EVQELYSPIFLIVAA](http://crdd.osdd.net/raghava/ifnepitope/pep_design.php?sequence=EVQELYSPIFLIVAA&method=hybrid&model=main) | MERCI | NEGATIVE | 10 |
| 19 | 92-107 | [VQELYSPIFLIVAAI](http://crdd.osdd.net/raghava/ifnepitope/pep_design.php?sequence=VQELYSPIFLIVAAI&method=hybrid&model=main) | MERCI | NEGATIVE | 17 |
| 20 | 93-108 | [QELYSPIFLIVAAIV](http://crdd.osdd.net/raghava/ifnepitope/pep_design.php?sequence=QELYSPIFLIVAAIV&method=hybrid&model=main) | MERCI | NEGATIVE | 23 |
| 21 | 94-109 | [ELYSPIFLIVAAIVF](http://crdd.osdd.net/raghava/ifnepitope/pep_design.php?sequence=ELYSPIFLIVAAIVF&method=hybrid&model=main) | MERCI | NEGATIVE | 26 |
| 22 | 95-110 | [LYSPIFLIVAAIVFI](http://crdd.osdd.net/raghava/ifnepitope/pep_design.php?sequence=LYSPIFLIVAAIVFI&method=hybrid&model=main) | MERCI | NEGATIVE | 37 |
| 23 | 96-111 | [YSPIFLIVAAIVFIT](http://crdd.osdd.net/raghava/ifnepitope/pep_design.php?sequence=YSPIFLIVAAIVFIT&method=hybrid&model=main) | MERCI | NEGATIVE | 36 |
| 24 | 97-112 | [SPIFLIVAAIVFITL](http://crdd.osdd.net/raghava/ifnepitope/pep_design.php?sequence=SPIFLIVAAIVFITL&method=hybrid&model=main) | MERCI | NEGATIVE | 53 |
| 25 | 98-113 | [PIFLIVAAIVFITLC](http://crdd.osdd.net/raghava/ifnepitope/pep_design.php?sequence=PIFLIVAAIVFITLC&method=hybrid&model=main) | MERCI | NEGATIVE | 57 |
| 26 | 99-114 | [IFLIVAAIVFITLCF](http://crdd.osdd.net/raghava/ifnepitope/pep_design.php?sequence=IFLIVAAIVFITLCF&method=hybrid&model=main) | MERCI | NEGATIVE | 50 |
| 27 | 100-115 | [FLIVAAIVFITLCFT](http://crdd.osdd.net/raghava/ifnepitope/pep_design.php?sequence=FLIVAAIVFITLCFT&method=hybrid&model=main) | MERCI | NEGATIVE | 24 |
| 28 | 101-116 | [LIVAAIVFITLCFTL](http://crdd.osdd.net/raghava/ifnepitope/pep_design.php?sequence=LIVAAIVFITLCFTL&method=hybrid&model=main) | MERCI | NEGATIVE | 27 |
| 29 | 102-117 | [IVAAIVFITLCFTLK](http://crdd.osdd.net/raghava/ifnepitope/pep_design.php?sequence=IVAAIVFITLCFTLK&method=hybrid&model=main) | MERCI | NEGATIVE | 24 |
| 30 | 103-118 | [VAAIVFITLCFTLKR](http://crdd.osdd.net/raghava/ifnepitope/pep_design.php?sequence=VAAIVFITLCFTLKR&method=hybrid&model=main) | MERCI | NEGATIVE | 21 |
| 31 | 104-119 | [AAIVFITLCFTLKRK](http://crdd.osdd.net/raghava/ifnepitope/pep_design.php?sequence=AAIVFITLCFTLKRK&method=hybrid&model=main) | MERCI | NEGATIVE | 20 |
| 32 | 105-120 | [AIVFITLCFTLKRKT](http://crdd.osdd.net/raghava/ifnepitope/pep_design.php?sequence=AIVFITLCFTLKRKT&method=hybrid&model=main) | MERCI | NEGATIVE | 19 |
| 33 | 106-121 | [IVFITLCFTLKRKTE](http://crdd.osdd.net/raghava/ifnepitope/pep_design.php?sequence=IVFITLCFTLKRKTE&method=hybrid&model=main) | MERCI | NEGATIVE | 14 |
| 34 | 107-121 | [VFITLCFTLKRKTE](http://crdd.osdd.net/raghava/ifnepitope/pep_design.php?sequence=VFITLCFTLKRKTE&method=hybrid&model=main) | MERCI | NEGATIVE | 7 |
| 35 | 108-121 | [FITLCFTLKRKTE](http://crdd.osdd.net/raghava/ifnepitope/pep_design.php?sequence=FITLCFTLKRKTE&method=hybrid&model=main) | MERCI | NEGATIVE | 3 |
| 36 | 8-23 | [LITLATCELYHYQEC](http://crdd.osdd.net/raghava/ifnepitope/pep_design.php?sequence=LITLATCELYHYQEC&method=hybrid&model=main) | SVM | NEGATIVE | -0.3964923 |
| 37 | 9-24 | [ITLATCELYHYQECV](http://crdd.osdd.net/raghava/ifnepitope/pep_design.php?sequence=ITLATCELYHYQECV&method=hybrid&model=main) | SVM | NEGATIVE | -0.43659053 |
| 38 | 10-25 | [TLATCELYHYQECVR](http://crdd.osdd.net/raghava/ifnepitope/pep_design.php?sequence=TLATCELYHYQECVR&method=hybrid&model=main) | SVM | NEGATIVE | -0.36373807 |
| 39 | 11-26 | [LATCELYHYQECVRG](http://crdd.osdd.net/raghava/ifnepitope/pep_design.php?sequence=LATCELYHYQECVRG&method=hybrid&model=main) | SVM | NEGATIVE | -0.33801788 |
| 40 | 12-27 | [ATCELYHYQECVRGT](http://crdd.osdd.net/raghava/ifnepitope/pep_design.php?sequence=ATCELYHYQECVRGT&method=hybrid&model=main) | SVM | NEGATIVE | -0.32486214 |
| 41 | 13-28 | [TCELYHYQECVRGTT](http://crdd.osdd.net/raghava/ifnepitope/pep_design.php?sequence=TCELYHYQECVRGTT&method=hybrid&model=main) | SVM | NEGATIVE | -0.41382925 |
| 42 | 14-29 | [CELYHYQECVRGTTV](http://crdd.osdd.net/raghava/ifnepitope/pep_design.php?sequence=CELYHYQECVRGTTV&method=hybrid&model=main) | SVM | NEGATIVE | -0.43674413 |
| 43 | 15-30 | [ELYHYQECVRGTTVL](http://crdd.osdd.net/raghava/ifnepitope/pep_design.php?sequence=ELYHYQECVRGTTVL&method=hybrid&model=main) | SVM | NEGATIVE | -0.12761662 |
| 44 | 16-31 | [LYHYQECVRGTTVLL](http://crdd.osdd.net/raghava/ifnepitope/pep_design.php?sequence=LYHYQECVRGTTVLL&method=hybrid&model=main) | SVM | NEGATIVE | -0.28981723 |
| 45 | 17-32 | [YHYQECVRGTTVLLK](http://crdd.osdd.net/raghava/ifnepitope/pep_design.php?sequence=YHYQECVRGTTVLLK&method=hybrid&model=main) | SVM | NEGATIVE | -0.4155203 |
| 46 | 18-33 | [HYQECVRGTTVLLKE](http://crdd.osdd.net/raghava/ifnepitope/pep_design.php?sequence=HYQECVRGTTVLLKE&method=hybrid&model=main) | SVM | NEGATIVE | -0.61298453 |
| 47 | 19-34 | [YQECVRGTTVLLKEP](http://crdd.osdd.net/raghava/ifnepitope/pep_design.php?sequence=YQECVRGTTVLLKEP&method=hybrid&model=main) | SVM | NEGATIVE | -0.5557322 |
| 48 | 20-35 | [QECVRGTTVLLKEPC](http://crdd.osdd.net/raghava/ifnepitope/pep_design.php?sequence=QECVRGTTVLLKEPC&method=hybrid&model=main) | SVM | NEGATIVE | -0.60870163 |
| 49 | 21-36 | [ECVRGTTVLLKEPCS](http://crdd.osdd.net/raghava/ifnepitope/pep_design.php?sequence=ECVRGTTVLLKEPCS&method=hybrid&model=main) | SVM | NEGATIVE | -0.60061069 |
| 50 | 22-37 | [CVRGTTVLLKEPCSS](http://crdd.osdd.net/raghava/ifnepitope/pep_design.php?sequence=CVRGTTVLLKEPCSS&method=hybrid&model=main) | SVM | NEGATIVE | -0.38145153 |
| 51 | 23-38 | [VRGTTVLLKEPCSSG](http://crdd.osdd.net/raghava/ifnepitope/pep_design.php?sequence=VRGTTVLLKEPCSSG&method=hybrid&model=main) | SVM | NEGATIVE | -0.54745555 |
| 52 | 24-39 | [RGTTVLLKEPCSSGT](http://crdd.osdd.net/raghava/ifnepitope/pep_design.php?sequence=RGTTVLLKEPCSSGT&method=hybrid&model=main) | SVM | NEGATIVE | -0.35721508 |
| 53 | 25-40 | [GTTVLLKEPCSSGTY](http://crdd.osdd.net/raghava/ifnepitope/pep_design.php?sequence=GTTVLLKEPCSSGTY&method=hybrid&model=main) | SVM | NEGATIVE | -0.59513677 |
| 54 | 26-41 | [TTVLLKEPCSSGTYE](http://crdd.osdd.net/raghava/ifnepitope/pep_design.php?sequence=TTVLLKEPCSSGTYE&method=hybrid&model=main) | SVM | NEGATIVE | -0.94974393 |
| 55 | 27-42 | [TVLLKEPCSSGTYEG](http://crdd.osdd.net/raghava/ifnepitope/pep_design.php?sequence=TVLLKEPCSSGTYEG&method=hybrid&model=main) | SVM | NEGATIVE | -0.8623141 |
| 56 | 28-43 | [VLLKEPCSSGTYEGN](http://crdd.osdd.net/raghava/ifnepitope/pep_design.php?sequence=VLLKEPCSSGTYEGN&method=hybrid&model=main) | SVM | NEGATIVE | -0.8350547 |
| 57 | 29-44 | [LLKEPCSSGTYEGNS](http://crdd.osdd.net/raghava/ifnepitope/pep_design.php?sequence=LLKEPCSSGTYEGNS&method=hybrid&model=main) | SVM | NEGATIVE | -0.84337517 |
| 58 | 30-45 | [LKEPCSSGTYEGNSP](http://crdd.osdd.net/raghava/ifnepitope/pep_design.php?sequence=LKEPCSSGTYEGNSP&method=hybrid&model=main) | SVM | NEGATIVE | -1.0849232 |
| 59 | 31-46 | [KEPCSSGTYEGNSPF](http://crdd.osdd.net/raghava/ifnepitope/pep_design.php?sequence=KEPCSSGTYEGNSPF&method=hybrid&model=main) | SVM | NEGATIVE | -0.7436134 |
| 60 | 32-47 | [EPCSSGTYEGNSPFH](http://crdd.osdd.net/raghava/ifnepitope/pep_design.php?sequence=EPCSSGTYEGNSPFH&method=hybrid&model=main) | SVM | NEGATIVE | -0.57136791 |
| 61 | 33-48 | [PCSSGTYEGNSPFHP](http://crdd.osdd.net/raghava/ifnepitope/pep_design.php?sequence=PCSSGTYEGNSPFHP&method=hybrid&model=main) | SVM | NEGATIVE | -0.29086599 |
| 62 | 34-49 | [CSSGTYEGNSPFHPL](http://crdd.osdd.net/raghava/ifnepitope/pep_design.php?sequence=CSSGTYEGNSPFHPL&method=hybrid&model=main) | SVM | NEGATIVE | -0.25374648 |
| 63 | 35-50 | [SSGTYEGNSPFHPLA](http://crdd.osdd.net/raghava/ifnepitope/pep_design.php?sequence=SSGTYEGNSPFHPLA&method=hybrid&model=main) | SVM | NEGATIVE | -0.35004221 |
| 64 | 36-51 | [SGTYEGNSPFHPLAD](http://crdd.osdd.net/raghava/ifnepitope/pep_design.php?sequence=SGTYEGNSPFHPLAD&method=hybrid&model=main) | SVM | NEGATIVE | -0.40490226 |
| 65 | 37-52 | [GTYEGNSPFHPLADN](http://crdd.osdd.net/raghava/ifnepitope/pep_design.php?sequence=GTYEGNSPFHPLADN&method=hybrid&model=main) | SVM | NEGATIVE | -0.25973356 |
| 66 | 38-53 | [TYEGNSPFHPLADNK](http://crdd.osdd.net/raghava/ifnepitope/pep_design.php?sequence=TYEGNSPFHPLADNK&method=hybrid&model=main) | SVM | NEGATIVE | -0.15354697 |
| 67 | 39-54 | [YEGNSPFHPLADNKF](http://crdd.osdd.net/raghava/ifnepitope/pep_design.php?sequence=YEGNSPFHPLADNKF&method=hybrid&model=main) | SVM | NEGATIVE | -0.15237586 |
| 68 | 40-55 | [EGNSPFHPLADNKFA](http://crdd.osdd.net/raghava/ifnepitope/pep_design.php?sequence=EGNSPFHPLADNKFA&method=hybrid&model=main) | SVM | POSITIVE | 0.032174937 |
| 69 | 41-56 | [GNSPFHPLADNKFAL](http://crdd.osdd.net/raghava/ifnepitope/pep_design.php?sequence=GNSPFHPLADNKFAL&method=hybrid&model=main) | SVM | NEGATIVE | -0.070722692 |
| 70 | 42-57 | [NSPFHPLADNKFALT](http://crdd.osdd.net/raghava/ifnepitope/pep_design.php?sequence=NSPFHPLADNKFALT&method=hybrid&model=main) | SVM | NEGATIVE | -0.045004676 |
| 71 | 43-58 | [SPFHPLADNKFALTC](http://crdd.osdd.net/raghava/ifnepitope/pep_design.php?sequence=SPFHPLADNKFALTC&method=hybrid&model=main) | SVM | POSITIVE | 0.11946523 |
| 72 | 44-59 | [PFHPLADNKFALTCF](http://crdd.osdd.net/raghava/ifnepitope/pep_design.php?sequence=PFHPLADNKFALTCF&method=hybrid&model=main) | SVM | POSITIVE | 0.34694402 |
| 73 | 45-60 | [FHPLADNKFALTCFS](http://crdd.osdd.net/raghava/ifnepitope/pep_design.php?sequence=FHPLADNKFALTCFS&method=hybrid&model=main) | SVM | POSITIVE | 0.18226704 |
| 74 | 53-68 | [FALTCFSTQFAFACP](http://crdd.osdd.net/raghava/ifnepitope/pep_design.php?sequence=FALTCFSTQFAFACP&method=hybrid&model=main) | SVM | POSITIVE | 0.26179696 |
| 75 | 54-69 | [ALTCFSTQFAFACPD](http://crdd.osdd.net/raghava/ifnepitope/pep_design.php?sequence=ALTCFSTQFAFACPD&method=hybrid&model=main) | SVM | NEGATIVE | -0.31590597 |
| 76 | 55-70 | [LTCFSTQFAFACPDG](http://crdd.osdd.net/raghava/ifnepitope/pep_design.php?sequence=LTCFSTQFAFACPDG&method=hybrid&model=main) | SVM | NEGATIVE | -0.48992135 |
| 77 | 56-71 | [TCFSTQFAFACPDGV](http://crdd.osdd.net/raghava/ifnepitope/pep_design.php?sequence=TCFSTQFAFACPDGV&method=hybrid&model=main) | SVM | NEGATIVE | -0.46976195 |
| 78 | 57-72 | [CFSTQFAFACPDGVK](http://crdd.osdd.net/raghava/ifnepitope/pep_design.php?sequence=CFSTQFAFACPDGVK&method=hybrid&model=main) | SVM | NEGATIVE | -0.58399382 |
| 79 | 58-73 | [FSTQFAFACPDGVKH](http://crdd.osdd.net/raghava/ifnepitope/pep_design.php?sequence=FSTQFAFACPDGVKH&method=hybrid&model=main) | SVM | NEGATIVE | -0.48484981 |
| 80 | 59-74 | [STQFAFACPDGVKHV](http://crdd.osdd.net/raghava/ifnepitope/pep_design.php?sequence=STQFAFACPDGVKHV&method=hybrid&model=main) | SVM | NEGATIVE | -0.37827618 |
| 81 | 60-75 | [TQFAFACPDGVKHVY](http://crdd.osdd.net/raghava/ifnepitope/pep_design.php?sequence=TQFAFACPDGVKHVY&method=hybrid&model=main) | SVM | NEGATIVE | -0.18175308 |
| 82 | 61-76 | [QFAFACPDGVKHVYQ](http://crdd.osdd.net/raghava/ifnepitope/pep_design.php?sequence=QFAFACPDGVKHVYQ&method=hybrid&model=main) | SVM | POSITIVE | 0.073164413 |
| 83 | 62-77 | [FAFACPDGVKHVYQL](http://crdd.osdd.net/raghava/ifnepitope/pep_design.php?sequence=FAFACPDGVKHVYQL&method=hybrid&model=main) | SVM | POSITIVE | 0.1328959 |
| 84 | 63-78 | [AFACPDGVKHVYQLR](http://crdd.osdd.net/raghava/ifnepitope/pep_design.php?sequence=AFACPDGVKHVYQLR&method=hybrid&model=main) | SVM | NEGATIVE | -0.22336182 |
| 85 | 64-79 | [FACPDGVKHVYQLRA](http://crdd.osdd.net/raghava/ifnepitope/pep_design.php?sequence=FACPDGVKHVYQLRA&method=hybrid&model=main) | SVM | POSITIVE | 0.034314679 |
| 86 | 65-80 | [ACPDGVKHVYQLRAR](http://crdd.osdd.net/raghava/ifnepitope/pep_design.php?sequence=ACPDGVKHVYQLRAR&method=hybrid&model=main) | SVM | NEGATIVE | -0.20191633 |
| 87 | 66-81 | [CPDGVKHVYQLRARS](http://crdd.osdd.net/raghava/ifnepitope/pep_design.php?sequence=CPDGVKHVYQLRARS&method=hybrid&model=main) | SVM | NEGATIVE | -0.19746149 |
| 88 | 67-82 | [PDGVKHVYQLRARSV](http://crdd.osdd.net/raghava/ifnepitope/pep_design.php?sequence=PDGVKHVYQLRARSV&method=hybrid&model=main) | SVM | NEGATIVE | -0.27635207 |
| 89 | 68-83 | [DGVKHVYQLRARSVS](http://crdd.osdd.net/raghava/ifnepitope/pep_design.php?sequence=DGVKHVYQLRARSVS&method=hybrid&model=main) | SVM | NEGATIVE | -0.1444906 |
| 90 | 69-84 | [GVKHVYQLRARSVSP](http://crdd.osdd.net/raghava/ifnepitope/pep_design.php?sequence=GVKHVYQLRARSVSP&method=hybrid&model=main) | SVM | NEGATIVE | -0.20120938 |
| 91 | 70-85 | [VKHVYQLRARSVSPK](http://crdd.osdd.net/raghava/ifnepitope/pep_design.php?sequence=VKHVYQLRARSVSPK&method=hybrid&model=main) | SVM | NEGATIVE | -0.49008738 |
| 92 | 71-86 | [KHVYQLRARSVSPKL](http://crdd.osdd.net/raghava/ifnepitope/pep_design.php?sequence=KHVYQLRARSVSPKL&method=hybrid&model=main) | SVM | NEGATIVE | -0.54086825 |
| 93 | 72-87 | [HVYQLRARSVSPKLF](http://crdd.osdd.net/raghava/ifnepitope/pep_design.php?sequence=HVYQLRARSVSPKLF&method=hybrid&model=main) | SVM | NEGATIVE | -0.41641536 |
| 94 | 73-88 | [VYQLRARSVSPKLFI](http://crdd.osdd.net/raghava/ifnepitope/pep_design.php?sequence=VYQLRARSVSPKLFI&method=hybrid&model=main) | SVM | NEGATIVE | -0.44028952 |
| 95 | 74-89 | [YQLRARSVSPKLFIR](http://crdd.osdd.net/raghava/ifnepitope/pep_design.php?sequence=YQLRARSVSPKLFIR&method=hybrid&model=main) | SVM | NEGATIVE | -0.49328652 |
| 96 | 75-90 | [QLRARSVSPKLFIRQ](http://crdd.osdd.net/raghava/ifnepitope/pep_design.php?sequence=QLRARSVSPKLFIRQ&method=hybrid&model=main) | SVM | NEGATIVE | -0.67552507 |
| 97 | 76-91 | [LRARSVSPKLFIRQE](http://crdd.osdd.net/raghava/ifnepitope/pep_design.php?sequence=LRARSVSPKLFIRQE&method=hybrid&model=main) | SVM | NEGATIVE | -0.69180092 |
| 98 | 77-92 | [RARSVSPKLFIRQEE](http://crdd.osdd.net/raghava/ifnepitope/pep_design.php?sequence=RARSVSPKLFIRQEE&method=hybrid&model=main) | SVM | NEGATIVE | -0.66328064 |
| 99 | 78-93 | [ARSVSPKLFIRQEEV](http://crdd.osdd.net/raghava/ifnepitope/pep_design.php?sequence=ARSVSPKLFIRQEEV&method=hybrid&model=main) | SVM | NEGATIVE | -0.88198451 |
| 100 | 79-94 | [RSVSPKLFIRQEEVQ](http://crdd.osdd.net/raghava/ifnepitope/pep_design.php?sequence=RSVSPKLFIRQEEVQ&method=hybrid&model=main) | SVM | NEGATIVE | -0.87735206 |
| ORF8 Protein | | | | | |
| 1 | 20-35 | [SLQSCTQHQPYVVDD](http://crdd.osdd.net/raghava/ifnepitope/pep_design.php?sequence=SLQSCTQHQPYVVDD&method=hybrid&model=main) | MERCI | POSITIVE | 1 |
| 2 | 21-36 | [LQSCTQHQPYVVDDP](http://crdd.osdd.net/raghava/ifnepitope/pep_design.php?sequence=LQSCTQHQPYVVDDP&method=hybrid&model=main) | MERCI | POSITIVE | 1 |
| 3 | 22-37 | [QSCTQHQPYVVDDPC](http://crdd.osdd.net/raghava/ifnepitope/pep_design.php?sequence=QSCTQHQPYVVDDPC&method=hybrid&model=main) | MERCI | POSITIVE | 1 |
| 4 | 23-38 | [SCTQHQPYVVDDPCP](http://crdd.osdd.net/raghava/ifnepitope/pep_design.php?sequence=SCTQHQPYVVDDPCP&method=hybrid&model=main) | MERCI | POSITIVE | 1 |
| 5 | 24-39 | [CTQHQPYVVDDPCPI](http://crdd.osdd.net/raghava/ifnepitope/pep_design.php?sequence=CTQHQPYVVDDPCPI&method=hybrid&model=main) | MERCI | POSITIVE | 1 |
| 6 | 25-40 | [TQHQPYVVDDPCPIH](http://crdd.osdd.net/raghava/ifnepitope/pep_design.php?sequence=TQHQPYVVDDPCPIH&method=hybrid&model=main) | MERCI | POSITIVE | 1 |
| 7 | 26-41 | [QHQPYVVDDPCPIHF](http://crdd.osdd.net/raghava/ifnepitope/pep_design.php?sequence=QHQPYVVDDPCPIHF&method=hybrid&model=main) | MERCI | POSITIVE | 1 |
| 8 | 27-42 | [HQPYVVDDPCPIHFY](http://crdd.osdd.net/raghava/ifnepitope/pep_design.php?sequence=HQPYVVDDPCPIHFY&method=hybrid&model=main) | MERCI | POSITIVE | 1 |
| 9 | 28-43 | [QPYVVDDPCPIHFYS](http://crdd.osdd.net/raghava/ifnepitope/pep_design.php?sequence=QPYVVDDPCPIHFYS&method=hybrid&model=main) | MERCI | POSITIVE | 1 |
| 10 | 29-44 | [PYVVDDPCPIHFYSK](http://crdd.osdd.net/raghava/ifnepitope/pep_design.php?sequence=PYVVDDPCPIHFYSK&method=hybrid&model=main) | MERCI | POSITIVE | 1 |
| 11 | 30-45 | [YVVDDPCPIHFYSKW](http://crdd.osdd.net/raghava/ifnepitope/pep_design.php?sequence=YVVDDPCPIHFYSKW&method=hybrid&model=main) | MERCI | POSITIVE | 1 |
| 12 | 31-46 | [VVDDPCPIHFYSKWY](http://crdd.osdd.net/raghava/ifnepitope/pep_design.php?sequence=VVDDPCPIHFYSKWY&method=hybrid&model=main) | MERCI | POSITIVE | 1 |
| 13 | 32-47 | [VDDPCPIHFYSKWYI](http://crdd.osdd.net/raghava/ifnepitope/pep_design.php?sequence=VDDPCPIHFYSKWYI&method=hybrid&model=main) | MERCI | POSITIVE | 1 |
| 14 | 33-48 | [DDPCPIHFYSKWYIR](http://crdd.osdd.net/raghava/ifnepitope/pep_design.php?sequence=DDPCPIHFYSKWYIR&method=hybrid&model=main) | MERCI | POSITIVE | 1 |
| 15 | 34-49 | [DPCPIHFYSKWYIRV](http://crdd.osdd.net/raghava/ifnepitope/pep_design.php?sequence=DPCPIHFYSKWYIRV&method=hybrid&model=main) | MERCI | POSITIVE | 1 |
| 16 | 35-50 | [PCPIHFYSKWYIRVG](http://crdd.osdd.net/raghava/ifnepitope/pep_design.php?sequence=PCPIHFYSKWYIRVG&method=hybrid&model=main) | MERCI | POSITIVE | 1 |
| 17 | 36-51 | [CPIHFYSKWYIRVGA](http://crdd.osdd.net/raghava/ifnepitope/pep_design.php?sequence=CPIHFYSKWYIRVGA&method=hybrid&model=main) | MERCI | POSITIVE | 1 |
| 18 | 37-52 | [PIHFYSKWYIRVGAR](http://crdd.osdd.net/raghava/ifnepitope/pep_design.php?sequence=PIHFYSKWYIRVGAR&method=hybrid&model=main) | MERCI | POSITIVE | 1 |
| 19 | 38-53 | [IHFYSKWYIRVGARK](http://crdd.osdd.net/raghava/ifnepitope/pep_design.php?sequence=IHFYSKWYIRVGARK&method=hybrid&model=main) | MERCI | POSITIVE | 1 |
| 20 | 39-54 | [HFYSKWYIRVGARKS](http://crdd.osdd.net/raghava/ifnepitope/pep_design.php?sequence=HFYSKWYIRVGARKS&method=hybrid&model=main) | MERCI | POSITIVE | 1 |
| 21 | 0-15 | [MKFLVFLGIITTVAA](http://crdd.osdd.net/raghava/ifnepitope/pep_design.php?sequence=MKFLVFLGIITTVAA&method=hybrid&model=main) | MERCI | NEGATIVE | 23 |
| 22 | 1-16 | [KFLVFLGIITTVAAF](http://crdd.osdd.net/raghava/ifnepitope/pep_design.php?sequence=KFLVFLGIITTVAAF&method=hybrid&model=main) | MERCI | NEGATIVE | 24 |
| 23 | 2-17 | [FLVFLGIITTVAAFH](http://crdd.osdd.net/raghava/ifnepitope/pep_design.php?sequence=FLVFLGIITTVAAFH&method=hybrid&model=main) | MERCI | NEGATIVE | 18 |
| 24 | 3-18 | [LVFLGIITTVAAFHQ](http://crdd.osdd.net/raghava/ifnepitope/pep_design.php?sequence=LVFLGIITTVAAFHQ&method=hybrid&model=main) | MERCI | NEGATIVE | 15 |
| 25 | 4-19 | [VFLGIITTVAAFHQE](http://crdd.osdd.net/raghava/ifnepitope/pep_design.php?sequence=VFLGIITTVAAFHQE&method=hybrid&model=main) | MERCI | NEGATIVE | 13 |
| 26 | 5-20 | [FLGIITTVAAFHQEC](http://crdd.osdd.net/raghava/ifnepitope/pep_design.php?sequence=FLGIITTVAAFHQEC&method=hybrid&model=main) | MERCI | NEGATIVE | 13 |
| 27 | 6-21 | [LGIITTVAAFHQECS](http://crdd.osdd.net/raghava/ifnepitope/pep_design.php?sequence=LGIITTVAAFHQECS&method=hybrid&model=main) | MERCI | NEGATIVE | 9 |
| 28 | 7-22 | [GIITTVAAFHQECSL](http://crdd.osdd.net/raghava/ifnepitope/pep_design.php?sequence=GIITTVAAFHQECSL&method=hybrid&model=main) | MERCI | NEGATIVE | 4 |
| 29 | 8-23 | [IITTVAAFHQECSLQ](http://crdd.osdd.net/raghava/ifnepitope/pep_design.php?sequence=IITTVAAFHQECSLQ&method=hybrid&model=main) | MERCI | NEGATIVE | 1 |
| 30 | 47-62 | [RVGARKSAPLIELCV](http://crdd.osdd.net/raghava/ifnepitope/pep_design.php?sequence=RVGARKSAPLIELCV&method=hybrid&model=main) | MERCI | NEGATIVE | 1 |
| 31 | 48-63 | [VGARKSAPLIELCVD](http://crdd.osdd.net/raghava/ifnepitope/pep_design.php?sequence=VGARKSAPLIELCVD&method=hybrid&model=main) | MERCI | NEGATIVE | 1 |
| 32 | 49-64 | [GARKSAPLIELCVDE](http://crdd.osdd.net/raghava/ifnepitope/pep_design.php?sequence=GARKSAPLIELCVDE&method=hybrid&model=main) | MERCI | NEGATIVE | 1 |
| 33 | 50-65 | [ARKSAPLIELCVDEA](http://crdd.osdd.net/raghava/ifnepitope/pep_design.php?sequence=ARKSAPLIELCVDEA&method=hybrid&model=main) | MERCI | NEGATIVE | 1 |
| 34 | 51-66 | [RKSAPLIELCVDEAG](http://crdd.osdd.net/raghava/ifnepitope/pep_design.php?sequence=RKSAPLIELCVDEAG&method=hybrid&model=main) | MERCI | NEGATIVE | 1 |
| 35 | 52-67 | [KSAPLIELCVDEAGS](http://crdd.osdd.net/raghava/ifnepitope/pep_design.php?sequence=KSAPLIELCVDEAGS&method=hybrid&model=main) | MERCI | NEGATIVE | 1 |
| 36 | 53-68 | [SAPLIELCVDEAGSK](http://crdd.osdd.net/raghava/ifnepitope/pep_design.php?sequence=SAPLIELCVDEAGSK&method=hybrid&model=main) | MERCI | NEGATIVE | 2 |
| 37 | 54-69 | [APLIELCVDEAGSKS](http://crdd.osdd.net/raghava/ifnepitope/pep_design.php?sequence=APLIELCVDEAGSKS&method=hybrid&model=main) | MERCI | NEGATIVE | 2 |
| 38 | 55-70 | [PLIELCVDEAGSKSP](http://crdd.osdd.net/raghava/ifnepitope/pep_design.php?sequence=PLIELCVDEAGSKSP&method=hybrid&model=main) | MERCI | NEGATIVE | 2 |
| 39 | 56-71 | [LIELCVDEAGSKSPI](http://crdd.osdd.net/raghava/ifnepitope/pep_design.php?sequence=LIELCVDEAGSKSPI&method=hybrid&model=main) | MERCI | NEGATIVE | 1 |
| 40 | 57-72 | [IELCVDEAGSKSPIQ](http://crdd.osdd.net/raghava/ifnepitope/pep_design.php?sequence=IELCVDEAGSKSPIQ&method=hybrid&model=main) | MERCI | NEGATIVE | 1 |
| 41 | 58-73 | [ELCVDEAGSKSPIQY](http://crdd.osdd.net/raghava/ifnepitope/pep_design.php?sequence=ELCVDEAGSKSPIQY&method=hybrid&model=main) | MERCI | NEGATIVE | 1 |
| 42 | 64-79 | [AGSKSPIQYIDIGNY](http://crdd.osdd.net/raghava/ifnepitope/pep_design.php?sequence=AGSKSPIQYIDIGNY&method=hybrid&model=main) | MERCI | NEGATIVE | 1 |
| 43 | 65-80 | [GSKSPIQYIDIGNYT](http://crdd.osdd.net/raghava/ifnepitope/pep_design.php?sequence=GSKSPIQYIDIGNYT&method=hybrid&model=main) | MERCI | NEGATIVE | 1 |
| 44 | 66-81 | [SKSPIQYIDIGNYTV](http://crdd.osdd.net/raghava/ifnepitope/pep_design.php?sequence=SKSPIQYIDIGNYTV&method=hybrid&model=main) | MERCI | NEGATIVE | 1 |
| 45 | 67-82 | [KSPIQYIDIGNYTVS](http://crdd.osdd.net/raghava/ifnepitope/pep_design.php?sequence=KSPIQYIDIGNYTVS&method=hybrid&model=main) | MERCI | NEGATIVE | 1 |
| 46 | 68-83 | [SPIQYIDIGNYTVSC](http://crdd.osdd.net/raghava/ifnepitope/pep_design.php?sequence=SPIQYIDIGNYTVSC&method=hybrid&model=main) | MERCI | NEGATIVE | 1 |
| 47 | 69-84 | [PIQYIDIGNYTVSCL](http://crdd.osdd.net/raghava/ifnepitope/pep_design.php?sequence=PIQYIDIGNYTVSCL&method=hybrid&model=main) | MERCI | NEGATIVE | 1 |
| 48 | 70-85 | [IQYIDIGNYTVSCLP](http://crdd.osdd.net/raghava/ifnepitope/pep_design.php?sequence=IQYIDIGNYTVSCLP&method=hybrid&model=main) | MERCI | NEGATIVE | 1 |
| 49 | 71-86 | [QYIDIGNYTVSCLPF](http://crdd.osdd.net/raghava/ifnepitope/pep_design.php?sequence=QYIDIGNYTVSCLPF&method=hybrid&model=main) | MERCI | NEGATIVE | 1 |
| 50 | 9-24 | [ITTVAAFHQECSLQS](http://crdd.osdd.net/raghava/ifnepitope/pep_design.php?sequence=ITTVAAFHQECSLQS&method=hybrid&model=main) | SVM | NEGATIVE | -0.34599481 |
| 51 | 10-25 | [TTVAAFHQECSLQSC](http://crdd.osdd.net/raghava/ifnepitope/pep_design.php?sequence=TTVAAFHQECSLQSC&method=hybrid&model=main) | SVM | NEGATIVE | -0.50923047 |
| 52 | 11-26 | [TVAAFHQECSLQSCT](http://crdd.osdd.net/raghava/ifnepitope/pep_design.php?sequence=TVAAFHQECSLQSCT&method=hybrid&model=main) | SVM | NEGATIVE | -0.79521234 |
| 53 | 12-27 | [VAAFHQECSLQSCTQ](http://crdd.osdd.net/raghava/ifnepitope/pep_design.php?sequence=VAAFHQECSLQSCTQ&method=hybrid&model=main) | SVM | NEGATIVE | -0.91794256 |
| 54 | 13-28 | [AAFHQECSLQSCTQH](http://crdd.osdd.net/raghava/ifnepitope/pep_design.php?sequence=AAFHQECSLQSCTQH&method=hybrid&model=main) | SVM | NEGATIVE | -0.91555362 |
| 55 | 14-29 | [AFHQECSLQSCTQHQ](http://crdd.osdd.net/raghava/ifnepitope/pep_design.php?sequence=AFHQECSLQSCTQHQ&method=hybrid&model=main) | SVM | NEGATIVE | -0.80384925 |
| 56 | 15-30 | [FHQECSLQSCTQHQP](http://crdd.osdd.net/raghava/ifnepitope/pep_design.php?sequence=FHQECSLQSCTQHQP&method=hybrid&model=main) | SVM | NEGATIVE | -0.91281832 |
| 57 | 16-31 | [HQECSLQSCTQHQPY](http://crdd.osdd.net/raghava/ifnepitope/pep_design.php?sequence=HQECSLQSCTQHQPY&method=hybrid&model=main) | SVM | NEGATIVE | -1.2227109 |
| 58 | 17-32 | [QECSLQSCTQHQPYV](http://crdd.osdd.net/raghava/ifnepitope/pep_design.php?sequence=QECSLQSCTQHQPYV&method=hybrid&model=main) | SVM | NEGATIVE | -1.2520744 |
| 59 | 18-33 | [ECSLQSCTQHQPYVV](http://crdd.osdd.net/raghava/ifnepitope/pep_design.php?sequence=ECSLQSCTQHQPYVV&method=hybrid&model=main) | SVM | NEGATIVE | -1.1534793 |
| 60 | 19-34 | [CSLQSCTQHQPYVVD](http://crdd.osdd.net/raghava/ifnepitope/pep_design.php?sequence=CSLQSCTQHQPYVVD&method=hybrid&model=main) | SVM | NEGATIVE | -0.9826804 |
| 61 | 40-55 | [FYSKWYIRVGARKSA](http://crdd.osdd.net/raghava/ifnepitope/pep_design.php?sequence=FYSKWYIRVGARKSA&method=hybrid&model=main) | SVM | NEGATIVE | -0.83731364 |
| 62 | 41-56 | [YSKWYIRVGARKSAP](http://crdd.osdd.net/raghava/ifnepitope/pep_design.php?sequence=YSKWYIRVGARKSAP&method=hybrid&model=main) | SVM | NEGATIVE | -0.70146455 |
| 63 | 42-57 | [SKWYIRVGARKSAPL](http://crdd.osdd.net/raghava/ifnepitope/pep_design.php?sequence=SKWYIRVGARKSAPL&method=hybrid&model=main) | SVM | NEGATIVE | -0.38389667 |
| 64 | 43-58 | [KWYIRVGARKSAPLI](http://crdd.osdd.net/raghava/ifnepitope/pep_design.php?sequence=KWYIRVGARKSAPLI&method=hybrid&model=main) | SVM | POSITIVE | 0.13856453 |
| 65 | 44-59 | [WYIRVGARKSAPLIE](http://crdd.osdd.net/raghava/ifnepitope/pep_design.php?sequence=WYIRVGARKSAPLIE&method=hybrid&model=main) | SVM | POSITIVE | 0.25834615 |
| 66 | 45-60 | [YIRVGARKSAPLIEL](http://crdd.osdd.net/raghava/ifnepitope/pep_design.php?sequence=YIRVGARKSAPLIEL&method=hybrid&model=main) | SVM | POSITIVE | 0.084149621 |
| 67 | 46-61 | [IRVGARKSAPLIELC](http://crdd.osdd.net/raghava/ifnepitope/pep_design.php?sequence=IRVGARKSAPLIELC&method=hybrid&model=main) | SVM | NEGATIVE | -0.070383232 |
| 68 | 59-74 | [LCVDEAGSKSPIQYI](http://crdd.osdd.net/raghava/ifnepitope/pep_design.php?sequence=LCVDEAGSKSPIQYI&method=hybrid&model=main) | SVM | NEGATIVE | -0.61038954 |
| 69 | 60-75 | [CVDEAGSKSPIQYID](http://crdd.osdd.net/raghava/ifnepitope/pep_design.php?sequence=CVDEAGSKSPIQYID&method=hybrid&model=main) | SVM | NEGATIVE | -0.60879365 |
| 70 | 61-76 | [VDEAGSKSPIQYIDI](http://crdd.osdd.net/raghava/ifnepitope/pep_design.php?sequence=VDEAGSKSPIQYIDI&method=hybrid&model=main) | SVM | NEGATIVE | -0.49934503 |
| 71 | 62-77 | [DEAGSKSPIQYIDIG](http://crdd.osdd.net/raghava/ifnepitope/pep_design.php?sequence=DEAGSKSPIQYIDIG&method=hybrid&model=main) | SVM | NEGATIVE | -0.50283076 |
| 72 | 63-78 | [EAGSKSPIQYIDIGN](http://crdd.osdd.net/raghava/ifnepitope/pep_design.php?sequence=EAGSKSPIQYIDIGN&method=hybrid&model=main) | SVM | NEGATIVE | -0.51801783 |
| 73 | 72-87 | [YIDIGNYTVSCLPFT](http://crdd.osdd.net/raghava/ifnepitope/pep_design.php?sequence=YIDIGNYTVSCLPFT&method=hybrid&model=main) | SVM | NEGATIVE | -0.57916048 |
| 74 | 73-88 | [IDIGNYTVSCLPFTI](http://crdd.osdd.net/raghava/ifnepitope/pep_design.php?sequence=IDIGNYTVSCLPFTI&method=hybrid&model=main) | SVM | NEGATIVE | -0.65580111 |
| 75 | 74-89 | [DIGNYTVSCLPFTIN](http://crdd.osdd.net/raghava/ifnepitope/pep_design.php?sequence=DIGNYTVSCLPFTIN&method=hybrid&model=main) | SVM | NEGATIVE | -0.65330953 |
| 76 | 75-90 | [IGNYTVSCLPFTINC](http://crdd.osdd.net/raghava/ifnepitope/pep_design.php?sequence=IGNYTVSCLPFTINC&method=hybrid&model=main) | SVM | NEGATIVE | -0.73537646 |
| 77 | 76-91 | [GNYTVSCLPFTINCQ](http://crdd.osdd.net/raghava/ifnepitope/pep_design.php?sequence=GNYTVSCLPFTINCQ&method=hybrid&model=main) | SVM | NEGATIVE | -0.80376701 |
| 78 | 77-92 | [NYTVSCLPFTINCQE](http://crdd.osdd.net/raghava/ifnepitope/pep_design.php?sequence=NYTVSCLPFTINCQE&method=hybrid&model=main) | SVM | NEGATIVE | -0.95263022 |
| 79 | 78-93 | [YTVSCLPFTINCQEP](http://crdd.osdd.net/raghava/ifnepitope/pep_design.php?sequence=YTVSCLPFTINCQEP&method=hybrid&model=main) | SVM | NEGATIVE | -0.86105799 |
| 80 | 79-94 | [TVSCLPFTINCQEPK](http://crdd.osdd.net/raghava/ifnepitope/pep_design.php?sequence=TVSCLPFTINCQEPK&method=hybrid&model=main) | SVM | NEGATIVE | -0.92469062 |
| 81 | 80-95 | [VSCLPFTINCQEPKL](http://crdd.osdd.net/raghava/ifnepitope/pep_design.php?sequence=VSCLPFTINCQEPKL&method=hybrid&model=main) | SVM | NEGATIVE | -1.1091712 |
| 82 | 81-96 | [SCLPFTINCQEPKLG](http://crdd.osdd.net/raghava/ifnepitope/pep_design.php?sequence=SCLPFTINCQEPKLG&method=hybrid&model=main) | SVM | NEGATIVE | -0.88107346 |
| 83 | 82-97 | [CLPFTINCQEPKLGS](http://crdd.osdd.net/raghava/ifnepitope/pep_design.php?sequence=CLPFTINCQEPKLGS&method=hybrid&model=main) | SVM | NEGATIVE | -0.69699019 |
| 84 | 83-98 | [LPFTINCQEPKLGSL](http://crdd.osdd.net/raghava/ifnepitope/pep_design.php?sequence=LPFTINCQEPKLGSL&method=hybrid&model=main) | SVM | NEGATIVE | -0.64714543 |
| 85 | 84-99 | [PFTINCQEPKLGSLV](http://crdd.osdd.net/raghava/ifnepitope/pep_design.php?sequence=PFTINCQEPKLGSLV&method=hybrid&model=main) | SVM | NEGATIVE | -0.43853441 |
| 86 | 85-100 | [FTINCQEPKLGSLVV](http://crdd.osdd.net/raghava/ifnepitope/pep_design.php?sequence=FTINCQEPKLGSLVV&method=hybrid&model=main) | SVM | NEGATIVE | -0.52951322 |
| 87 | 86-101 | [TINCQEPKLGSLVVR](http://crdd.osdd.net/raghava/ifnepitope/pep_design.php?sequence=TINCQEPKLGSLVVR&method=hybrid&model=main) | SVM | NEGATIVE | -0.40040293 |
| 88 | 87-102 | [INCQEPKLGSLVVRC](http://crdd.osdd.net/raghava/ifnepitope/pep_design.php?sequence=INCQEPKLGSLVVRC&method=hybrid&model=main) | SVM | NEGATIVE | -0.4230422 |
| 89 | 88-103 | [NCQEPKLGSLVVRCS](http://crdd.osdd.net/raghava/ifnepitope/pep_design.php?sequence=NCQEPKLGSLVVRCS&method=hybrid&model=main) | SVM | NEGATIVE | -0.54464709 |
| 90 | 89-104 | [CQEPKLGSLVVRCSF](http://crdd.osdd.net/raghava/ifnepitope/pep_design.php?sequence=CQEPKLGSLVVRCSF&method=hybrid&model=main) | SVM | NEGATIVE | -0.33822692 |
| 91 | 90-105 | [QEPKLGSLVVRCSFY](http://crdd.osdd.net/raghava/ifnepitope/pep_design.php?sequence=QEPKLGSLVVRCSFY&method=hybrid&model=main) | SVM | NEGATIVE | -0.32576961 |
| 92 | 91-106 | [EPKLGSLVVRCSFYE](http://crdd.osdd.net/raghava/ifnepitope/pep_design.php?sequence=EPKLGSLVVRCSFYE&method=hybrid&model=main) | SVM | NEGATIVE | -0.38114835 |
| 93 | 92-107 | [PKLGSLVVRCSFYED](http://crdd.osdd.net/raghava/ifnepitope/pep_design.php?sequence=PKLGSLVVRCSFYED&method=hybrid&model=main) | SVM | NEGATIVE | -0.42359864 |
| 94 | 93-108 | [KLGSLVVRCSFYEDF](http://crdd.osdd.net/raghava/ifnepitope/pep_design.php?sequence=KLGSLVVRCSFYEDF&method=hybrid&model=main) | SVM | NEGATIVE | -0.15904329 |
| 95 | 94-109 | [LGSLVVRCSFYEDFL](http://crdd.osdd.net/raghava/ifnepitope/pep_design.php?sequence=LGSLVVRCSFYEDFL&method=hybrid&model=main) | SVM | NEGATIVE | -0.11849465 |
| 96 | 95-110 | [GSLVVRCSFYEDFLE](http://crdd.osdd.net/raghava/ifnepitope/pep_design.php?sequence=GSLVVRCSFYEDFLE&method=hybrid&model=main) | SVM | NEGATIVE | -0.19381552 |
| 97 | 96-111 | [SLVVRCSFYEDFLEY](http://crdd.osdd.net/raghava/ifnepitope/pep_design.php?sequence=SLVVRCSFYEDFLEY&method=hybrid&model=main) | SVM | NEGATIVE | -0.31529964 |
| 98 | 97-112 | [LVVRCSFYEDFLEYH](http://crdd.osdd.net/raghava/ifnepitope/pep_design.php?sequence=LVVRCSFYEDFLEYH&method=hybrid&model=main) | SVM | NEGATIVE | -0.31306183 |
| 99 | 98-113 | [VVRCSFYEDFLEYHD](http://crdd.osdd.net/raghava/ifnepitope/pep_design.php?sequence=VVRCSFYEDFLEYHD&method=hybrid&model=main) | SVM | NEGATIVE | -0.41437692 |
| 100 | 99-114 | [VRCSFYEDFLEYHDV](http://crdd.osdd.net/raghava/ifnepitope/pep_design.php?sequence=VRCSFYEDFLEYHDV&method=hybrid&model=main) | SVM | NEGATIVE | -0.63932543 |
| 101 | 100-115 | [RCSFYEDFLEYHDVR](http://crdd.osdd.net/raghava/ifnepitope/pep_design.php?sequence=RCSFYEDFLEYHDVR&method=hybrid&model=main) | SVM | NEGATIVE | -0.63932543 |
| 102 | 101-116 | [CSFYEDFLEYHDVRV](http://crdd.osdd.net/raghava/ifnepitope/pep_design.php?sequence=CSFYEDFLEYHDVRV&method=hybrid&model=main) | SVM | NEGATIVE | -0.59967292 |
| 103 | 102-117 | [SFYEDFLEYHDVRVV](http://crdd.osdd.net/raghava/ifnepitope/pep_design.php?sequence=SFYEDFLEYHDVRVV&method=hybrid&model=main) | SVM | NEGATIVE | -0.46440246 |
| 104 | 103-118 | [FYEDFLEYHDVRVVL](http://crdd.osdd.net/raghava/ifnepitope/pep_design.php?sequence=FYEDFLEYHDVRVVL&method=hybrid&model=main) | SVM | NEGATIVE | -0.39041266 |
| 105 | 104-119 | [YEDFLEYHDVRVVLD](http://crdd.osdd.net/raghava/ifnepitope/pep_design.php?sequence=YEDFLEYHDVRVVLD&method=hybrid&model=main) | SVM | NEGATIVE | -0.22270417 |
| 106 | 105-120 | [EDFLEYHDVRVVLDF](http://crdd.osdd.net/raghava/ifnepitope/pep_design.php?sequence=EDFLEYHDVRVVLDF&method=hybrid&model=main) | SVM | POSITIVE | 0.30645803 |
| 107 | 106-121 | [DFLEYHDVRVVLDFI](http://crdd.osdd.net/raghava/ifnepitope/pep_design.php?sequence=DFLEYHDVRVVLDFI&method=hybrid&model=main) | SVM | POSITIVE | 0.63927229 |
| 108 | 107-121 | [FLEYHDVRVVLDFI](http://crdd.osdd.net/raghava/ifnepitope/pep_design.php?sequence=FLEYHDVRVVLDFI&method=hybrid&model=main) | SVM | POSITIVE | 0.21940945 |
| 109 | 108-121 | [LEYHDVRVVLDFI](http://crdd.osdd.net/raghava/ifnepitope/pep_design.php?sequence=LEYHDVRVVLDFI&method=hybrid&model=main) | SVM | POSITIVE | 0.11750113 |
| 110 | 109-121 | [EYHDVRVVLDFI](http://crdd.osdd.net/raghava/ifnepitope/pep_design.php?sequence=EYHDVRVVLDFI&method=hybrid&model=main) | SVM | NEGATIVE | -0.0028019844 |
| 111 | 110-121 | [YHDVRVVLDFI](http://crdd.osdd.net/raghava/ifnepitope/pep_design.php?sequence=YHDVRVVLDFI&method=hybrid&model=main) | SVM | NEGATIVE | -0.034862224 |
| 112 | 111-121 | [HDVRVVLDFI](http://crdd.osdd.net/raghava/ifnepitope/pep_design.php?sequence=HDVRVVLDFI&method=hybrid&model=main) | SVM | NEGATIVE | -0.028663302 |
| 113 | 112-121 | [DVRVVLDFI](http://crdd.osdd.net/raghava/ifnepitope/pep_design.php?sequence=DVRVVLDFI&method=hybrid&model=main) | SVM | POSITIVE | 0.36411223 |
| ORF10 Protein | | | | | |
| 1 | 18-33 | [CRMNSRNYIAQVDVV](http://crdd.osdd.net/raghava/ifnepitope/pep_design.php?sequence=CRMNSRNYIAQVDVV&method=hybrid&model=main) | MERCI | POSITIVE | 1 |
| 2 | 19-34 | [RMNSRNYIAQVDVVN](http://crdd.osdd.net/raghava/ifnepitope/pep_design.php?sequence=RMNSRNYIAQVDVVN&method=hybrid&model=main) | MERCI | POSITIVE | 1 |
| 3 | 20-35 | [MNSRNYIAQVDVVNF](http://crdd.osdd.net/raghava/ifnepitope/pep_design.php?sequence=MNSRNYIAQVDVVNF&method=hybrid&model=main) | MERCI | POSITIVE | 1 |
| 4 | 21-36 | [NSRNYIAQVDVVNFN](http://crdd.osdd.net/raghava/ifnepitope/pep_design.php?sequence=NSRNYIAQVDVVNFN&method=hybrid&model=main) | MERCI | POSITIVE | 1 |
| 5 | 22-37 | [SRNYIAQVDVVNFNL](http://crdd.osdd.net/raghava/ifnepitope/pep_design.php?sequence=SRNYIAQVDVVNFNL&method=hybrid&model=main) | MERCI | POSITIVE | 1 |
| 6 | 23-38 | [RNYIAQVDVVNFNLT](http://crdd.osdd.net/raghava/ifnepitope/pep_design.php?sequence=RNYIAQVDVVNFNLT&method=hybrid&model=main) | MERCI | POSITIVE | 1 |
| 7 | 24-38 | [NYIAQVDVVNFNLT](http://crdd.osdd.net/raghava/ifnepitope/pep_design.php?sequence=NYIAQVDVVNFNLT&method=hybrid&model=main) | MERCI | POSITIVE | 1 |
| 8 | 25-38 | [YIAQVDVVNFNLT](http://crdd.osdd.net/raghava/ifnepitope/pep_design.php?sequence=YIAQVDVVNFNLT&method=hybrid&model=main) | MERCI | POSITIVE | 1 |
| 9 | 4-19 | [NVFAFPFTIYSLLLC](http://crdd.osdd.net/raghava/ifnepitope/pep_design.php?sequence=NVFAFPFTIYSLLLC&method=hybrid&model=main) | MERCI | NEGATIVE | 1 |
| 10 | 5-20 | [VFAFPFTIYSLLLCR](http://crdd.osdd.net/raghava/ifnepitope/pep_design.php?sequence=VFAFPFTIYSLLLCR&method=hybrid&model=main) | MERCI | NEGATIVE | 3 |
| 11 | 6-21 | [FAFPFTIYSLLLCRM](http://crdd.osdd.net/raghava/ifnepitope/pep_design.php?sequence=FAFPFTIYSLLLCRM&method=hybrid&model=main) | MERCI | NEGATIVE | 3 |
| 12 | 7-22 | [AFPFTIYSLLLCRMN](http://crdd.osdd.net/raghava/ifnepitope/pep_design.php?sequence=AFPFTIYSLLLCRMN&method=hybrid&model=main) | MERCI | NEGATIVE | 3 |
| 13 | 8-23 | [FPFTIYSLLLCRMNS](http://crdd.osdd.net/raghava/ifnepitope/pep_design.php?sequence=FPFTIYSLLLCRMNS&method=hybrid&model=main) | MERCI | NEGATIVE | 3 |
| 14 | 9-24 | [PFTIYSLLLCRMNSR](http://crdd.osdd.net/raghava/ifnepitope/pep_design.php?sequence=PFTIYSLLLCRMNSR&method=hybrid&model=main) | MERCI | NEGATIVE | 4 |
| 15 | 10-25 | [FTIYSLLLCRMNSRN](http://crdd.osdd.net/raghava/ifnepitope/pep_design.php?sequence=FTIYSLLLCRMNSRN&method=hybrid&model=main) | MERCI | NEGATIVE | 4 |
| 16 | 11-26 | [TIYSLLLCRMNSRNY](http://crdd.osdd.net/raghava/ifnepitope/pep_design.php?sequence=TIYSLLLCRMNSRNY&method=hybrid&model=main) | MERCI | NEGATIVE | 4 |
| 17 | 12-27 | [IYSLLLCRMNSRNYI](http://crdd.osdd.net/raghava/ifnepitope/pep_design.php?sequence=IYSLLLCRMNSRNYI&method=hybrid&model=main) | MERCI | NEGATIVE | 1 |
| 18 | 13-28 | [YSLLLCRMNSRNYIA](http://crdd.osdd.net/raghava/ifnepitope/pep_design.php?sequence=YSLLLCRMNSRNYIA&method=hybrid&model=main) | MERCI | NEGATIVE | 1 |
| 19 | 14-29 | [SLLLCRMNSRNYIAQ](http://crdd.osdd.net/raghava/ifnepitope/pep_design.php?sequence=SLLLCRMNSRNYIAQ&method=hybrid&model=main) | MERCI | NEGATIVE | 1 |
| 20 | 0-15 | [MGYINVFAFPFTIYS](http://crdd.osdd.net/raghava/ifnepitope/pep_design.php?sequence=MGYINVFAFPFTIYS&method=hybrid&model=main) | SVM | NEGATIVE | -0.27034709 |
| 21 | 1-16 | [GYINVFAFPFTIYSL](http://crdd.osdd.net/raghava/ifnepitope/pep_design.php?sequence=GYINVFAFPFTIYSL&method=hybrid&model=main) | SVM | NEGATIVE | -0.35400636 |
| 22 | 2-17 | [YINVFAFPFTIYSLL](http://crdd.osdd.net/raghava/ifnepitope/pep_design.php?sequence=YINVFAFPFTIYSLL&method=hybrid&model=main) | SVM | NEGATIVE | -0.11803847 |
| 23 | 3-18 | [INVFAFPFTIYSLLL](http://crdd.osdd.net/raghava/ifnepitope/pep_design.php?sequence=INVFAFPFTIYSLLL&method=hybrid&model=main) | SVM | POSITIVE | 0.10765496 |
| 24 | 15-30 | [LLLCRMNSRNYIAQV](http://crdd.osdd.net/raghava/ifnepitope/pep_design.php?sequence=LLLCRMNSRNYIAQV&method=hybrid&model=main) | SVM | NEGATIVE | -0.24865935 |
| 25 | 16-31 | [LLCRMNSRNYIAQVD](http://crdd.osdd.net/raghava/ifnepitope/pep_design.php?sequence=LLCRMNSRNYIAQVD&method=hybrid&model=main) | SVM | NEGATIVE | -0.51541261 |
| 26 | 17-32 | [LCRMNSRNYIAQVDV](http://crdd.osdd.net/raghava/ifnepitope/pep_design.php?sequence=LCRMNSRNYIAQVDV&method=hybrid&model=main) | SVM | NEGATIVE | -0.47281731 |
| 27 | 26-38 | [IAQVDVVNFNLT](http://crdd.osdd.net/raghava/ifnepitope/pep_design.php?sequence=IAQVDVVNFNLT&method=hybrid&model=main) | SVM | NEGATIVE | -0.65050672 |
| 28 | 27-38 | [AQVDVVNFNLT](http://crdd.osdd.net/raghava/ifnepitope/pep_design.php?sequence=AQVDVVNFNLT&method=hybrid&model=main) | SVM | NEGATIVE | -0.65891706 |
| 29 | 28-38 | [QVDVVNFNLT](http://crdd.osdd.net/raghava/ifnepitope/pep_design.php?sequence=QVDVVNFNLT&method=hybrid&model=main) | SVM | NEGATIVE | -0.84914375 |
| 30 | 29-38 | [VDVVNFNLT](http://crdd.osdd.net/raghava/ifnepitope/pep_design.php?sequence=VDVVNFNLT&method=hybrid&model=main) | SVM | NEGATIVE | -0.64692783 |
