## Supplementary material for "Designing of a next generation multiepitope based vaccine (MEV) against SARS-COV-2: Immunoinformatics and *in silico* approaches": Supplementary Table 9.docx

Supplementary Table 9. Linear B-cell epitopes in the final MEV construct.

| B-cell Epitopes | Position | Antigenicity |
| --- | --- | --- |
| LVDFQVTIAEAAYG | 113 | 1.0 |
| GPGPGMFHLVDFQV | 368 | 0.8 |
| PIHFYSKWGPGPGP | 419 | 0.6 |
| AAYFHLVDFQVTIA | 108 | 1.3 |
| LACFVLAAVYRIGP | 336 | 1.0 |
| PGPVTLACFVLAAV | 331 | 1.1 |
| GPGMFHLVDFQVTI | 370 | 1.2 |
| IGKCSTRGRKCCRR | 30 | 1.2 |
| TRGRKCCRRKKEAA | 35 | 1.0 |
| LPKEEQIGKCSTRG | 24 | 0.7 |
| MWSFNAAYFHLVDF | 103 | 1.1 |
| GPGPGESELVIGAV | 348 | 0.5 |
| VGPGPGFLLVTLAI | 287 | 0.6 |
| YIAQVGPGPGFLLV | 283 | 0.5 |
| PYVVDDPCPIAAYD | 173 | 0.5 |
| PLADNKFALTCAAY | 142 | 1.2 |
| GPGPKLGSLVVRCS | 429 | 0.7 |
| ELVIGAVILRGHLG | 355 | 0.8 |
| SKWAAYPIHFYSKW | 195 | 0.8 |
| WSFNAAYFHLVDFQ | 104 | 1.5 |
| WGPGPGPKLGSLVV | 426 | 0.5 |
| VIGAVILRGHLGPG | 357 | 1.0 |
| NLVIGFLFLTWGPG | 317 | 1.1 |
| AYDPNFKDQVILLN | 259 | 1.1 |
| PCPIHFYSKWAAYP | 188 | 0.9 |
| NAAYFHLVDFQVTI | 107 | 1.3 |
| GMFHLVDFQVTIAE | 327 | 1.1 |
| GESELVIGAVILRG | 352 | 0.5 |
| AVYRIGPGPGESEL | 343 | 0.5 |
| AQVGPGPGFLLVTL | 285 | 0.8 |
| DDPCPIHFYSKWGP | 415 | 1.1 |
| GPKLGSLVVRCSFY | 431 | 1.0 |
| FHPLAAYHPLADNK | 134 | 0.6 |
| SELVIGAVILRGHL | 354 | 0.5 |
| GLEQWNLVIGFLFL | 312 | 1.0 |
| YDDPCPIHFYSKWA | 185 | 0.9 |
| FNAAYFHLVDFQVT | 106 | 1.4 |
| LRGHLGPGPGMFHL | 363 | 0.9 |
| SKWGPGPGPKLGSL | 424 | 0.6 |
| GPGVVDDPCPIHFY | 410 | 0.5 |
| HFYSKWAAYPIHFY | 192 | 0.9 |
| DDPCPIAAYDDPCP | 177 | 0.8 |
| LAAYHPLADNKFAL | 137 | 0.5 |
| CPIHFYSKWGPGPG | 418 | 0.6 |
| DDPCPIHFYSKWAA | 186 | 0.8 |
| AYDDPCPIHFYSKW | 184 | 0.8 |
| YGNYTVSCLPFTIA | 215 | 1.6 |
| AYGTYEGNSPFHPL | 124 | 0.7 |
| AEAAYGTYEGNSPF | 121 | 0.6 |
| YSKWGPGPGPKLGS | 423 | 0.5 |
| ESELVIGAVILRGH | 353 | 0.6 |
| FVLAAVYRIGPGPG | 339 | 0.6 |
| PGPGPVTLACFVLA | 329 | 0.9 |
| PKEEQIGKCSTRGR | 25 | 1.3 |
| PGPGPKLGSLVVRC | 428 | 0.7 |
| PIHFYSKWAAYPIH | 190 | 1.0 |
| DPCPIAAYDDPCPI | 178 | 1.0 |
| VDDPCPIAAYDDPC | 176 | 0.5 |
| YHPLADNKFALTCA | 140 | 1.1 |
| AAYHPLADNKFALT | 138 | 0.6 |
| VDDPCPIHFYSKWG | 414 | 0.8 |
| LSPRWYFYAAYDPN | 250 | 1.1 |
| YTVSCLPFTIAAYL | 218 | 1.3 |
| FAFACPDGVAAYHQ | 159 | 0.7 |
| LAAVYRIGPGPGES | 341 | 05 |
| IAAYDDPCPIHFYS | 182 | 0.6 |
| RGHLGPGPGMFHLV | 364 | 1.1 |
| AAYDPNFKDQVILL | 258 | 1.2 |
| NFKDQVILLNAAYC | 263 | 0.8 |
| QPYVVDDPCPIAAY | 172 | 0.5 |
| VTLACFVLAAVYRI | 334 | 0.9 |
| EQWNLVIGFLFLTW | 314 | 1.2 |
| LEQWNLVIGFLFLT | 313 | 0.9 |
| AYLPFTINCQEPKL | 229 | 1.0 |
| AAYLPFTINCQEPK | 228 | 1.0 |
| PCPIHFYSKWGPGP | 417 | 0.9 |
| GPGESELVIGAVIL | 350 | 0.5 |
| ACFVLAAVYRIGPG | 337 | 0.9 |
| KWGPGPGPKLGSLV | 425 | 0.6 |
| LPFTIAAYLPFTIN | 223 | 1.2 |
| AVLSCLPKEEQIGK | 19 | 0.5 |
| GVVDDPCPIHFYSK | 412 | 0.6 |
| TIAAYLPFTINCQE | 226 | 1.1 |
| DPCPIHFYSKWAAY | 187 | 0.9 |
| EAAYGTYEGNSPFH | 122 | 0.6 |
| PGPKLGSLVVRCSF | 430 | 0.8 |
| IGPGPGESELVIGA | 347 | 0.9 |
| LVIGFLFLTWGPGP | 318 | 1.0 |
| VVDDPCPIHFYSKW | 413 | 0.5 |
| IGAVILRGHLGPGP | 358 | 1.2 |
| RWYFYAAYDPNFKD | 253 | 1.2 |
| VSCLPFTIAAYLPF | 220 | 1.3 |
