## Supplementary material for "Designing of a next generation multiepitope based vaccine (MEV) against SARS-COV-2: Immunoinformatics and *in silico* approaches": Supplementary Table 10.docx

Supplementary Table 10. Conformational B-cell epitopes in the final MEV construct.

| Conformational B-cell Epitopes | Position of epitopes on MEV 3D Structure |
| --- | --- |
| A:T151, A:C152, A:A153, A:A154, A:Y155, A:S156, A:Q158, A:C163, A:P164, A:D165, A:G166, A:V167, A:A168, A:A169, A:Y170, A:H171, A:Q172, A:P173 | 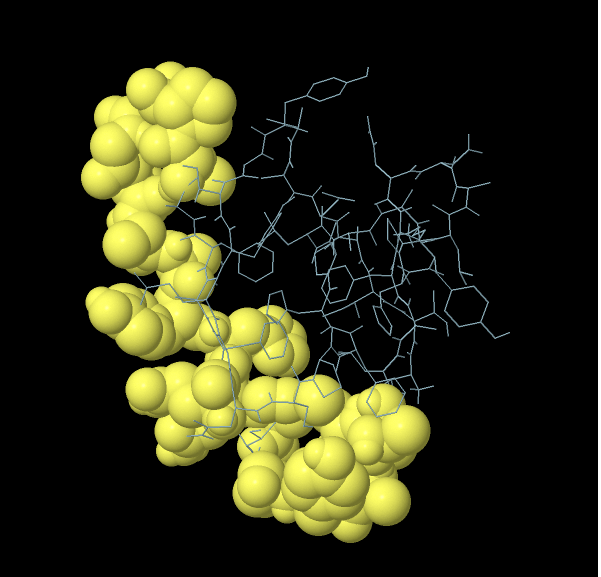 |
| A:L137, A:A138, A:A139, A:H141, A:P142, A:L143, A:A144, A:D145, A:N146 | 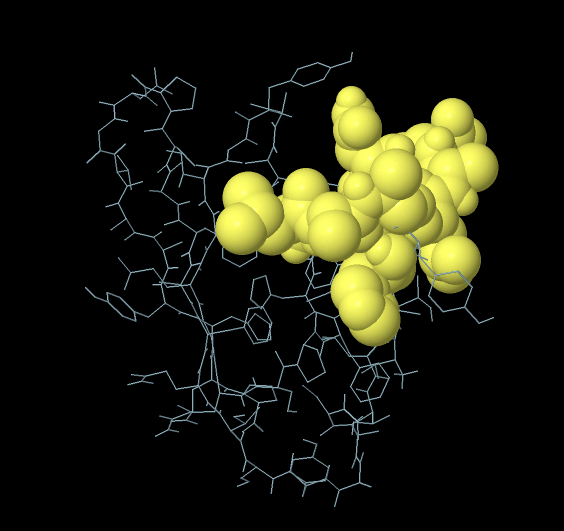 |
| A:A123, A:A124, A:Y125 | 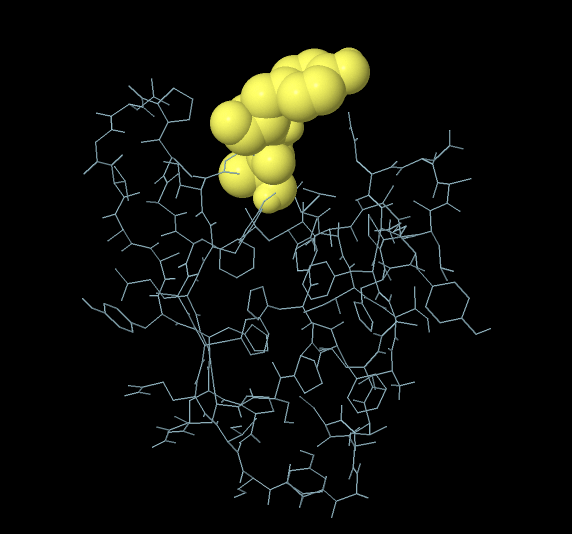 |
