## Supplementary material for "Designing of a next generation multiepitope based vaccine (MEV) against SARS-COV-2: Immunoinformatics and *in silico* approaches": Supplementary Table 11.docx

Supplementary Table 11. Codon optimized nucleotide sequence of MEV construct for cloning in *E. coli* strain K12. Green color bold sequence at 5` site (N-terminal) is representing XhoI restriction enzyme site, while maroon color bold sequence at 3` site (C-terminal) is representing HindIII restriction enzyme site.

**CTCGAG**GGTATCATCAACACCCTGCAGAAATACTACTGCCGTGTTCGTGGTGGTCG	TTGCGCTGTTCTGTCTTGCCTGCCGAAAGAAGAACAGATCGGTAAATGCT	CTACCCGTGGTCGTAAATGCTGCCGTCGTAAAAAAGAAGCTGCTGCTAAA TTCCTGCTGGTTACCCTGGCTATCCTGACCGCTCTGGCTGCTTACTTCCG TCTGTTCGCTCGTACCCGTTCTATGTGGTCTGCTGCTTACCGTCTGTTCG CTCGTACCCGTTCTATGTGGTCTTTCGCTGCTTACCTGTTCGCTCGTACC CGTTCTATGTGGTCTTTCAACGCTGCTTACTTCCACCTGGTTGACTTCCA GGTTACCATCGCTGAAGCTGCTTACGGTACCTACGAAGGTAACTCTCCGT TCCACCCGCTGGCTGCTTACCACCCGCTGGCTGACAACAAATTCGCTCTG ACCTGCGCTGCTTACTCTACCCAGTTCGCTTTCGCTTGCCCGGACGGTGT TGCTGCTTACCACCAGCCGTACGTTGTTGACGACCCGTGCCCGATCGCTG CTTACGACGACCCGTGCCCGATCCACTTCTACTCTAAATGGGCTGCTTAC CCGATCCACTTCTACTCTAAATGGTACATCCGTGTTGCTGCTTACGGCAA CTACACCGTGAGCTGCCTGCCGTTCACCATCGCTGCTTACCTGCCGTTCA CCATCAACTGCCAGGAACCGAAACTGGCTGCTTACAAAATGAAAGACCTG TCTCCGCGTTGGTACTTCTACGCTGCTTACGACCCGAACTTCAAAGACCA GGTTATCCTGCTGAACGCTGCTTACTGCCGTATGAACTCTCGTAACTACA TCGCTCAGGTTGGGCCGGGGCCGGGCTTCCTGCTGGTTACTCTGGCTATC CTGACCGCTCTGCGTCTGTGCGGTCCGGGTCCGGGTCTGGAACAGTGGAA CCTGGTTATCGGTTTCCTGTTCCTGACCTGGGGTCCGGGTCCGGGTCCGG TTACCCTGGCTTGCTTCGTTCTGGCTGCTGTTTACCGTATCGGTCCGGGT CCGGGTGAATCTGAACTGGTTATCGGTGCTGTTATCCTGCGTGGTCACCT GGGTCCGGGTCCGGGTATGTTCCACCTGGTTGACTTCCAGGTTACCATCG CTGAAATCCTGGGTCCGGGTCCGGGTGACGGCGTTAAACACGTGTACCAG CTGAGAGCGCGTTCTGTAAGCGGTCCGGGTCCGGGTGTTGTTGACGACCC GTGCCCGATCCACTTCTACTCTAAATGGGGTCCGGGTCCGGGTCCGAAAC TGGGTTCTCTGGTTGTTCGTTGCTCTTTCTACGAAGACGGTCCGGGTCCG GGTGCTCTGCTGCTGCTGGACCGTCTGAACCAGCTGGAATCTAAAATGGG GCCGGGGCCGGGCTTCTTCGGTATGTCGCGTATCGGTATGGAAGTTACCC CGTCTGGTGGTCCGGGTCCGGGTTGCCGTATGAACTCTCGTAACTACATC GCTCAGGTTGACGTT**AAGCTT**
