## Supplementary figures and images for "Designing of a next generation multiepitope based vaccine (MEV) against SARS-COV-2: Immunoinformatics and *in silico* approaches"

### Supply Figure 1.tif

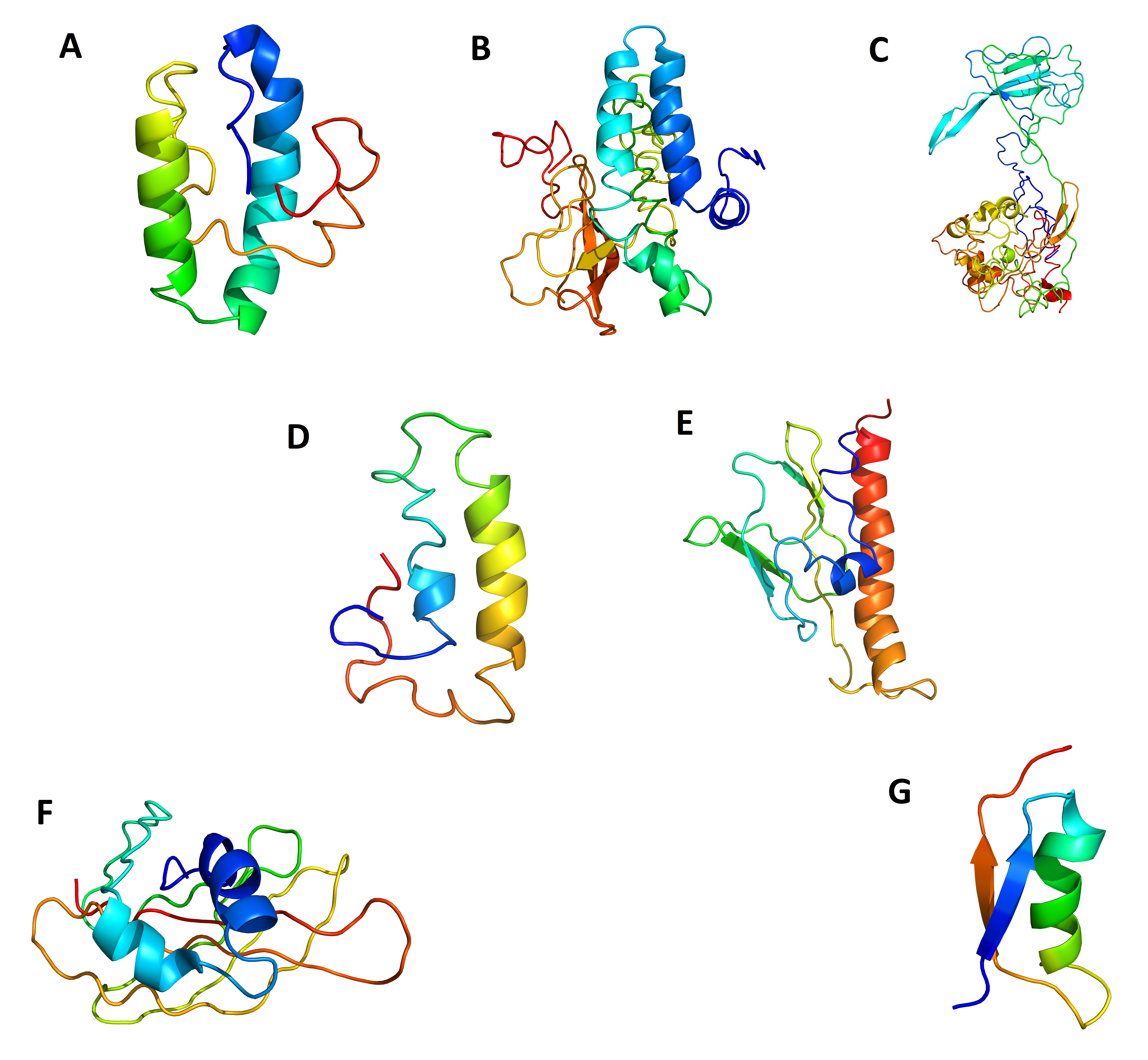

### Supply Figure 2.tif

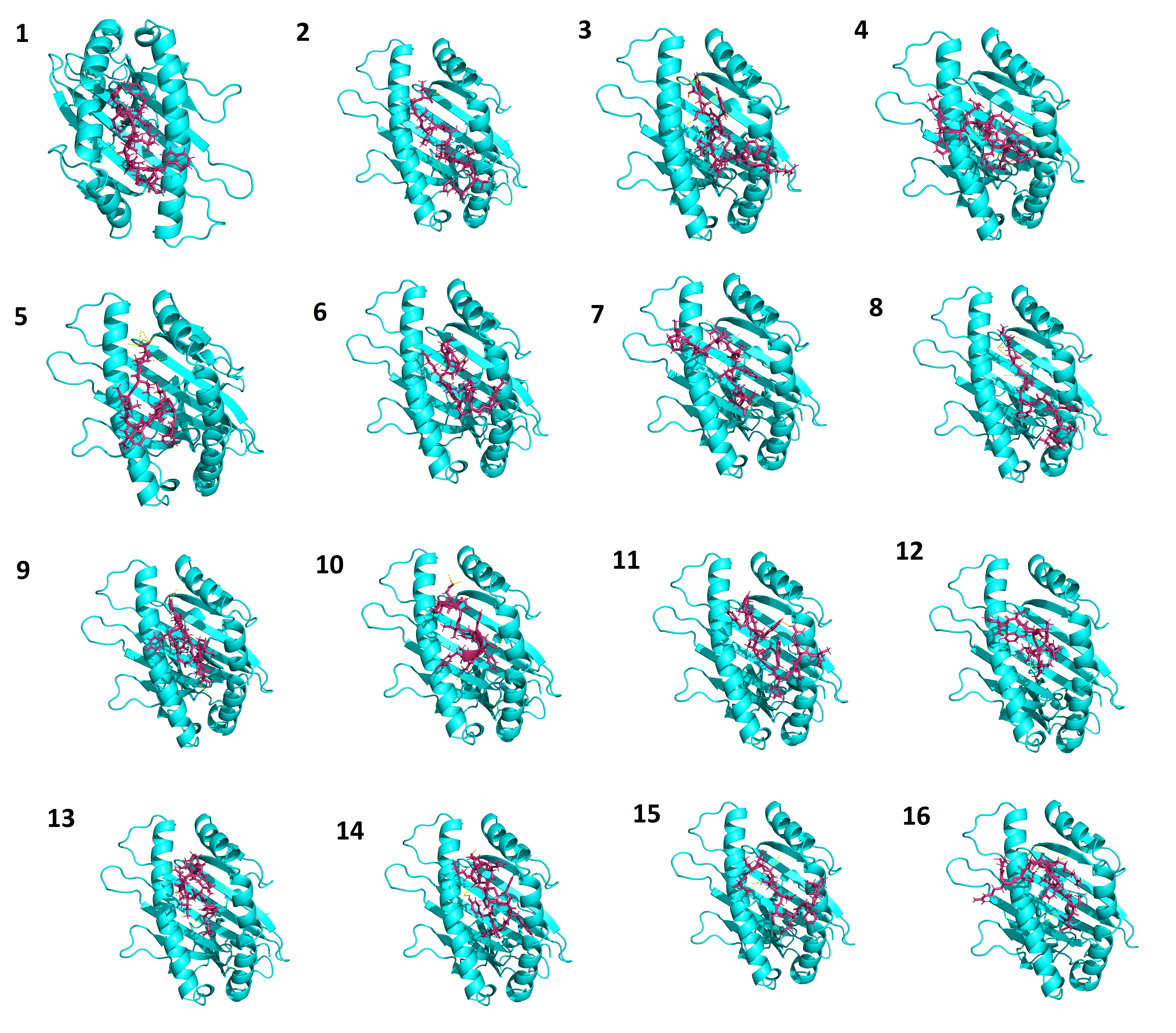

### Supply Figure 3.tif

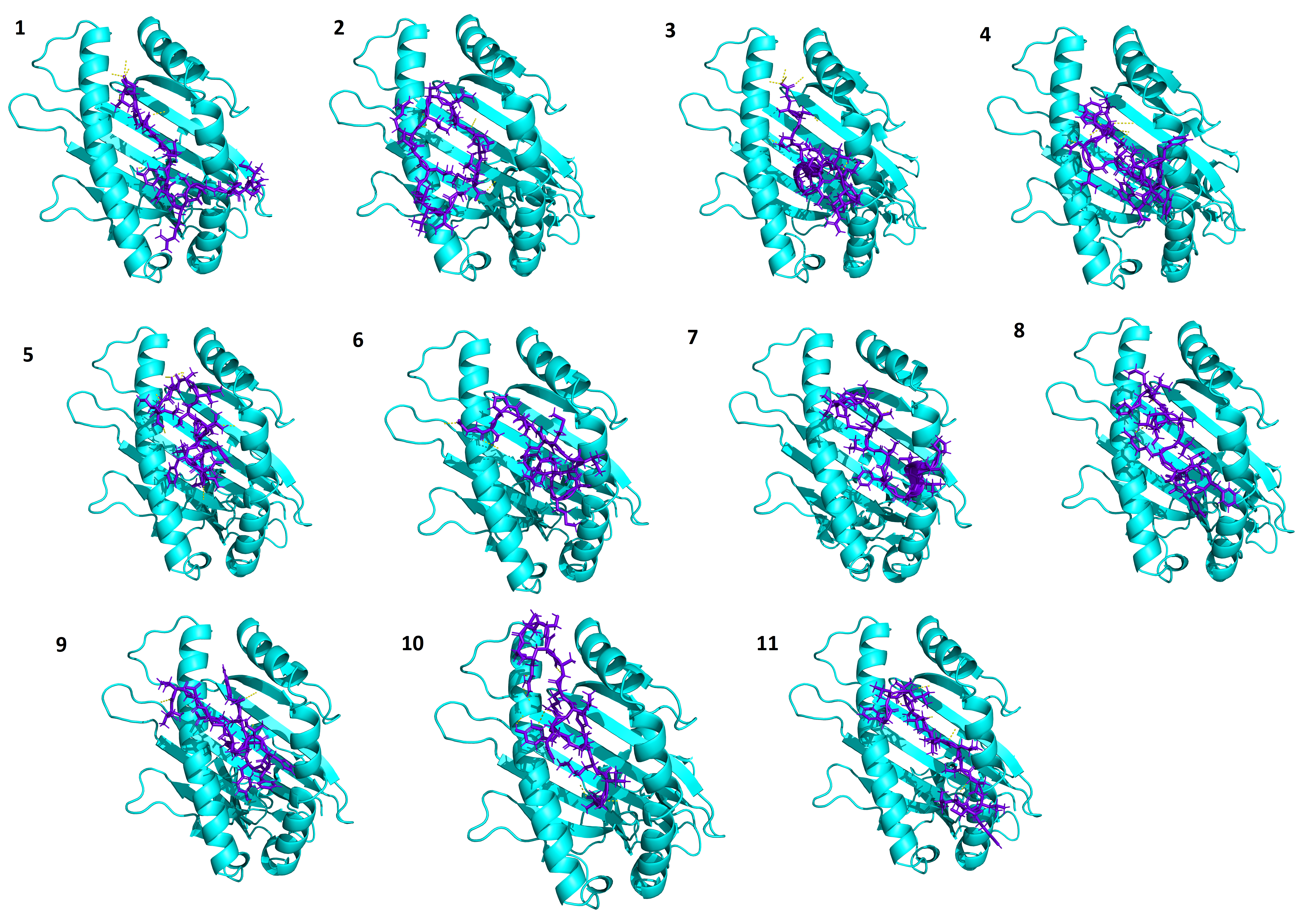

### Supply Figure 4.tif

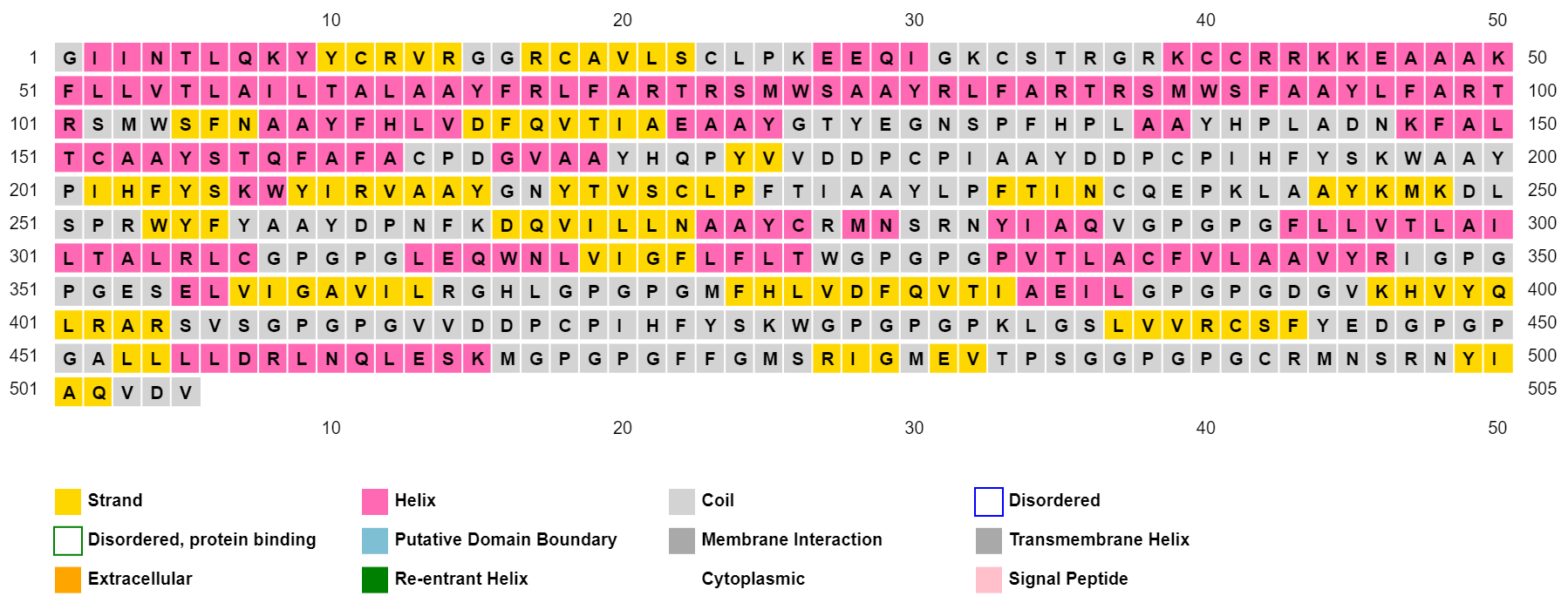

### Supply Figure 5.tif

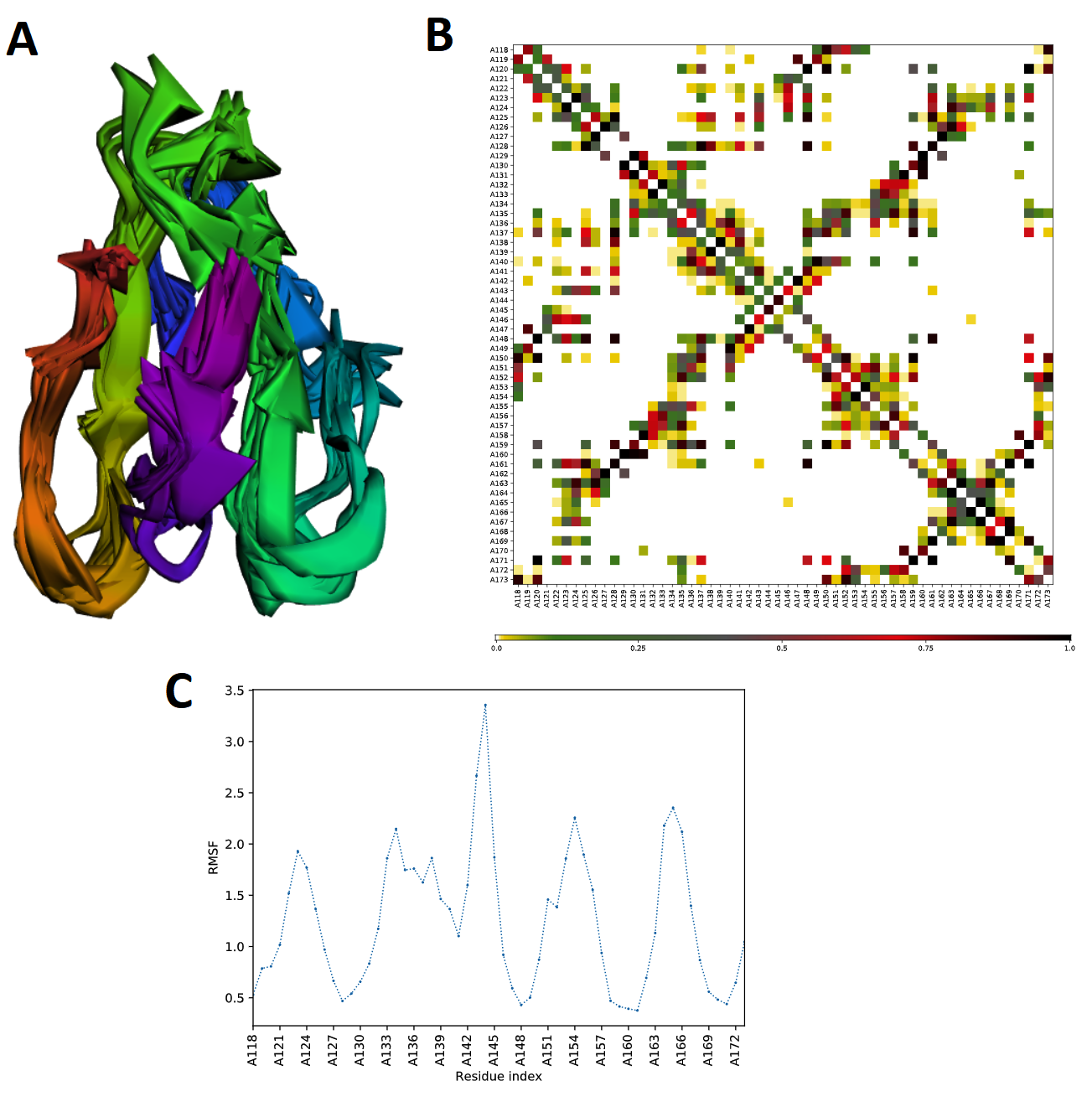
